# High throughput screening of eukaryotic release factor 1 variants to enhance noncanonical amino acid incorporation

**DOI:** 10.1101/2025.09.10.675417

**Authors:** Briana R. Lino, James A. Van Deventer

**Affiliations:** Chemical and Biological Engineering Department, Tufts University, Medford, Massachusetts 02155, USA; Biomedical Engineering Department, Tufts University, Medford, Massachusetts 02155, USA

**Keywords:** Noncanonical amino acids, genetic code expansion, eukaryotic release factor 1, yeast

## Abstract

Noncanonical amino acids (ncAAs) enable diversification of protein functions, but the efficiency of genetic code expansion (GCE) in eukaryotes is hindered by competition between suppressor tRNAs and release factors. Prior work has identified eukaryotic release factor 1 (eRF1) mutants that improve ncAA incorporation, suggesting that screens for improved variants may lead to further enhancements. Here, we developed a high-throughput system to screen eRF1 mutants in *Saccharomyces cerevisiae* where eRF1 mutants are coexpressed on a plasmid alongside genomically encoded, wild-type eRF1. This strategy enabled recovery of live cells expressing eRF1 variants that enhance ncAA incorporation, even with mutants known to severely affect cell viability in the absence of WT eRF1 expression. We prepared and screened a million-member library of randomly mutated eRF1 variants for clones exhibiting improved ncAA integration phenotypes. Deep sequencing revealed a diverse set of enriched mutations across all three major domains of eRF1. Interestingly, several enriched mutations identified here are also found in naturally occurring eRF1 homologs from species that recode canonical stop codons. When eRF1 variants were combined with yeast knockout strains also known to enhance ncAA incorporation, this resulted in further improvements to efficiency, highlighting the complementarity of release factor engineering to other GCE enhancement strategies. This work demonstrates that high-throughput engineering of the eukaryotic translational apparatus is a powerful approach to identify previously unknown solutions for enhancing ncAA incorporation, with implications for elucidating and precisely manipulating the molecular functions of essential translational machinery.

## MAIN

Integrating noncanonical amino acids (ncAAs) into proteins via genetic code expansion is a versatile methodology for diversifying the chemical and structural properties of proteins. Examples of proteins accessible by incorporation of ncAAs include proteins with precisely defined post-translational modifications^1–3^, proteins that crosslink to targets spontaneously or with light^4–8^, and proteins containing bioorthogonal handles that enable preparation of precisely defined conjugates^9–14^. Substantial progress has been made to engineer improved ncAA incorporation in response to stop codons in both prokaryotic and eukaryotic cells, focused primarily on engineering the aminoacyl-tRNA synthetases of orthogonal translation systems (OTSs), with some notable efforts to engineer tRNAs of OTSs, prokaryotic ribosomes, and prokaryotic elongation factors^15–27^. However, ncAA incorporation frequently remains inefficient relative to wild type (WT) protein translation because encoding ncAAs in response to stop codons must outcompete the native cellular machinery responsible for terminating translation. In *E. coli*, this problem has been addressed at the amber (UAG) stop codon by eliminating this codon from the genome and deleting the gene encoding release factor one (RF1), the only bacterial release factor that recognizes the amber codon^28–34^. Recently, Isaacs and coworkers reported an engineered variant of bacterial release factor 2 that facilitates more efficient readthrough of opal stop codons in *E. coli*^35^, demonstrating that rational engineering of prokaryotic release factors can overcome native termination constraints in support of improved ncAA incorporation. In eukaryotes, termination of translation is carried out by an omnipotent set of release factors, preventing implementation of the codon compression and release factor deletion strategy used in prokaryotes. Thus, addressing ncAA incorporation inefficiencies associated with termination of translation must be accomplished via altering the properties of the termination machinery.

Eukaryotic release factor 1 (eRF1) recognizes all three stop codons (UAA, UAG, UGA) and thus plays a central role in mediating termination of translation. A substantial body of literature describes functions of native and mutant eRF1 variants from a broad range of organisms^36–47^. Prior work has demonstrated that mutations to the SUP45 gene, which encodes eRF1, results in variable levels of stop codon readthrough frequencies at amber, opal, and ochre stop codons^43,46,48^; mutations in these studies were derived from both rational and high throughput approaches. Studies in mammalian cell expression systems have directly evaluated the effects of eRF1 variants on ncAA incorporation efficiency, demonstrating that point mutations^44,49–52^ and deletions^49^ to human eRF1 result in enhanced ncAA incorporation.

Interestingly, despite SUP45 being an essential gene, eRF1 exhibits substantial evolutionary plasticity, with naturally occurring variants supporting alternative genetic codes in which stop codons are reassigned to sense codons^53,54^. In ciliates and kinetoplastids, limited substitutions within eRF1 confer distinct stop-codon specificities, including selective recognition or reassignment of UGA, UAA, and UAG^39,54–58^. Similarly, mitochondrial release factors have evolved specialized functions that enable peptide release at noncanonical codons, reassigning stop codons for readthrough events^59^. These examples of alternate code organisms demonstrate that stop codon recognition by eRF1 is inherently tunable and can be modulated through relatively small sequence changes. This natural malleability provides a strong biological precedent for engineering eRF1 variants to attenuate translation termination, thereby enabling the use of stop codon readthrough for applications in synthetic biology such as genetic code expansion. Building on this foundation, we sought to implement a high throughput screening system to discover variants of *S. cerevisiae* eRF1 that enhance ncAA incorporation.

The system we established uses plasmid-based expression of eRF1 variants alongside the wild-type genomic copy of eRF1, in contrast to prior systems that eliminate wild-type eRF1 expression^43,49,60^. We found that coexpression of WT and mutant eRF1 facilitates evaluation of a wide range of variants and growth-based recovery of cells exhibiting enhanced ncAA incorporation phenotypes, even in cases where eRF1 mutants of interest are known to substantially reduce cell viability^37,47,61–63^. Combining this coexpression system with yeast display reporters and fluorescent protein reporters facilitated flow cytometric evaluation of ncAA incorporation efficiency and fidelity in response to the amber stop codon^64^. We then constructed and screened a million-member random mutagenesis library of eRF1 variants for enhanced ncAA incorporation using fluorescence-activated cell sorting. Deep sequencing of screening outputs revealed a broad set of enriched mutations, including at positions that have not been identified in prior eRF1 mutagenesis studies (to the best of our knowledge) as well as at positions where known mutations lead to improved stop codon readthrough. Flow cytometric analysis confirmed that both clones isolated from sorted libraries (containing multiple mutations) and point mutants identified during deep sequencing exhibit improvement in ncAA incorporation relative to controls. Finally, these variants could be combined with yeast strains with gene knockouts known to enhance ncAA incorporation to result in further enhancement of ncAA incorporation^65,66^. Overall, high throughput screening enabled unbiased engineering of a key component of the eukaryotic translation apparatus for enhanced ncAA incorporation; mutations identified here may also shed light on fundamental mechanisms of termination of translation in future studies. Focused screening with a single component of the translational apparatus is a powerful, complementary approach to genome engineering and genome-wide screening approaches to enhancing ncAA incorporation. This simple but powerful approach to release factor engineering is a versatile tool for pushing the limits of ncAA incorporation strategies and may also facilitate identification of critical properties of the translation apparatus that cannot be identified using rational approaches.

## RESULTS

### Parallel expression of eRF1 mutants to enable screening for enhanced ncAA incorporation

Prior work has shown that eRF1 mutations that promote aberrant stop codon readthrough also compromise cell viability^37,47,61–63^. To limit these effects, we sought to implement a system in yeast where wild-type eRF1 is expressed from genomic SUP45 and a second copy of the gene is introduced on a plasmid (also under control of the native promoter). We surmised that mutations to the plasmid-based copy could result in phenotypic effects on ncAA incorporation while still enabling cell propagation; this would enable evaluation and screening of eRF1 variant collections without the need for plasmid shuffling or conditional expression strategies^42,43,49^. To determine whether enhancement of ncAA incorporation can be observed with plasmid-based eRF1 variant expression, we acquired a subset of plasmids encoding SUP45 mutants (S30P, N58A, I222S, and P174Q) known to exhibit improved TAG codon readthrough from the work of Merritt et al^43^. Plasmids encoding eRF1 mutants were co-transformed into yeast alongside single-plasmid systems to evaluate ncAA incorporation (Fig. 1A). The single-plasmid systems^64^ encode dual fluorescent reporter genes to enable evaluation of ncAA incorporation efficiency and fidelity and an orthogonal translation system (OTS; LeuOmeRS for yeast display reporters or TyrOmeRS for fluorescent protein reporters), for incorporating the ncAA *O*-methyl-L-tyrosine (OmeY) in response to the amber codon. The yeast strain RJY100 (MATa AGA1::GAL1-AGA1::URA3 ura3-52 trp1::KanMX leu2-Δ200 his3-Δ200 pep4::HIS3 prbd1.6R can1 GAL) is a yeast display strain and is used in conjunction with a display-based reporter system (FAPB2.3.6L1TAG) for evaluating ncAA incorporation via labeling N-terminal and C-terminal epitope tags. BY4741 (MATa his3Δ1 leu2Δ0 met15Δ0 ura3Δ0) is commonly employed in yeast genetics and is used here in conjunction with dual fluorescent reporters encoding N-terminal (BFP) and C-terminal (GFP) fluorescent proteins flanking a linker containing a stop codon for ncAA incorporation (BXGTAG and BXGAltTAG; these reporters differ in the location of the TAG codon for ncAA incorporation, with ncAA incorporation at BXGTAG exhibiting higher efficiencies than ncAA incorporation at BXGAltTAG). A summary of the combinations of strains and reporters used in this section is provided in Table 1.

**Figure 1.**
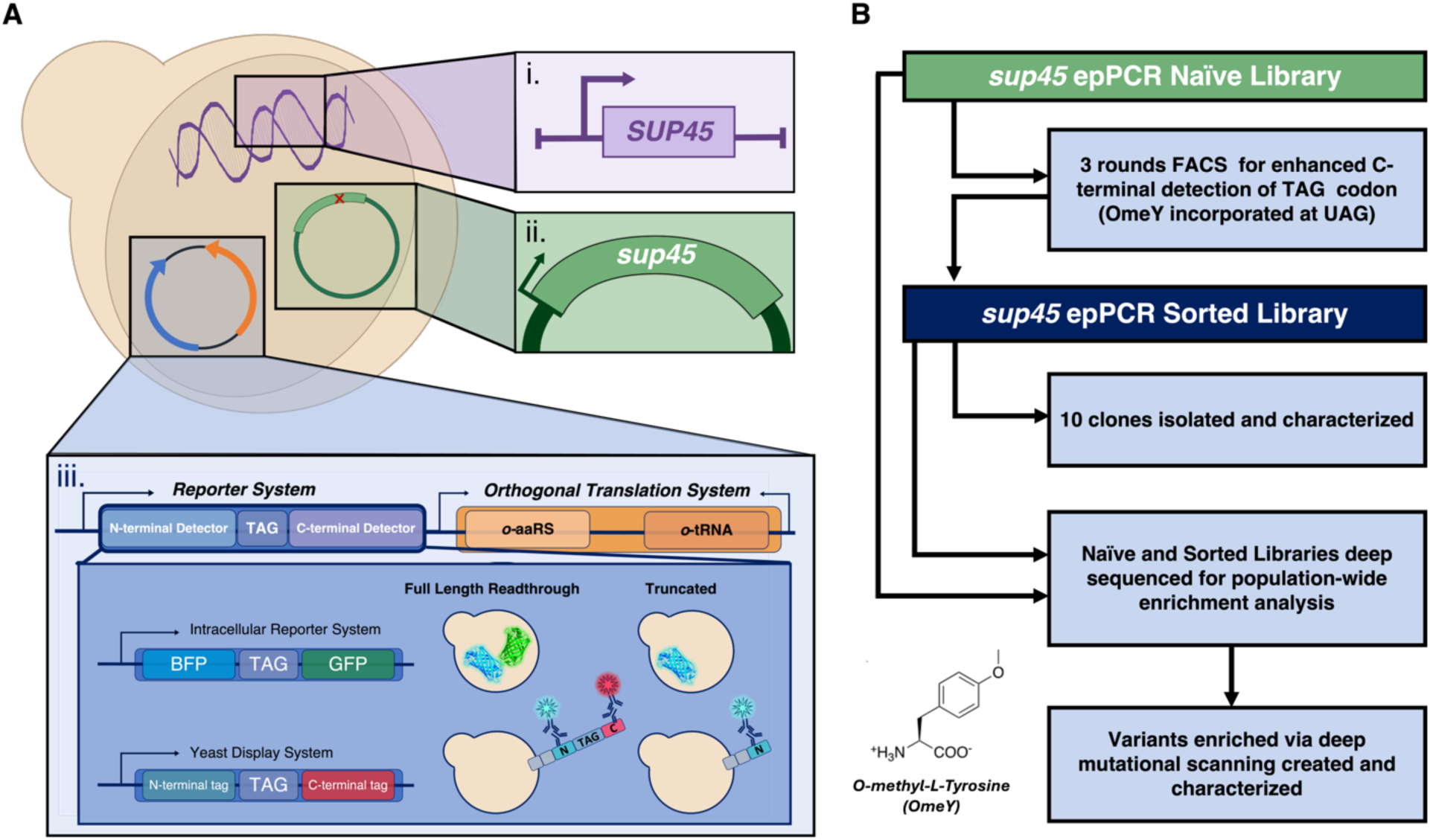
Overview of eRF1 experimentation. A) Schematic of *Saccharomyces cerevisiae* with (i) active endogenous SUP45 transformed with (ii) mutant SUP45 plasmid and (iii) single plasmid reporter/orthogonal translation system. B) Experimental workflow: eRF1 random mutagenesis (epPCR) library is screened via FACS for enhanced stop codon readthrough (OmeY incorporation at UAG codon). Sorted library is characterized via flow cytometry and sequencing of 10 clones (Table 2, Fig. S5), then both naïve and sorted libraries are deep sequenced and analyzed via deep mutational scanning (Fig. 2, S6-S8) guiding the identification of enriched variants for further phenotypic characterization (Fig. 3).

**Table 1.**
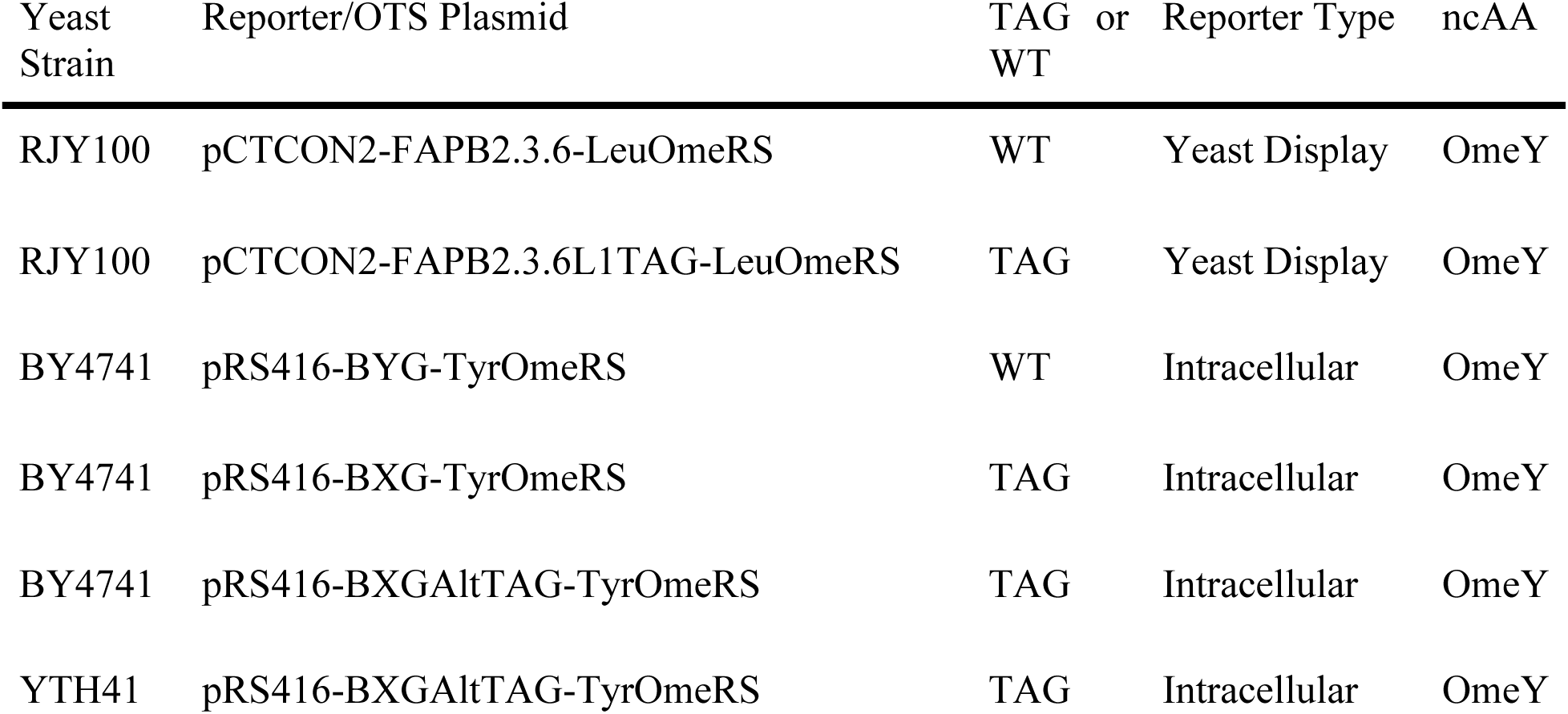
Single Plasmid Reporter Systems & Yeast Strain Combinations used in this work.

**Table 2.** Mutations found in subset of clones sorted for enhanced ncAA incorporation.

| Clone | Mutations |
| --- | --- |
| CR3B.6 | S20P, F187L, D232G, E258G |
| CR3T.2 | S61P*, D141G, A203T |
| CR3T.4 | C124R, K194E, F298L |
| CR3T.6 | S20P |
| CR3T.7 | R25G, N58S*, F288L, D362N, S421P* |
| CR3T.9 | R25G, V68A**, V165A, E366G* |
| BR2.2 | S70F**, K103T |
| BR2.4 | S30P**, I222L* |
| BR2.9 | L69S, S70P*, E280G, V412A |
| BR3T.4 | N64D, K105R, P116L |

\*Previously reported position for enhanced stop codon readthrough
\*\*Previously reported position & mutation for enhanced stop codon readthrough

We used flow cytometry experiments to evaluate changes in ncAA incorporation in strains coexpressing wild-type (genome) and mutant (plasmid-based) eRF1 variants. We evaluated relative readthrough efficiency (RRE) and increases in full-length fluorescent protein levels in the strains RJY100 and BY4741 transformed as listed in Table 1 (Fig. S1). Possible effects from plasmid-based expression of eRF1 were controlled for by comparing cells transformed with plasmids encoding eRF1 mutants with cells transformed with a plasmid encoding for wild-type eRF1 and the corresponding reporter system. RRE is calculated from dual-fluorescent reporter data by measuring the ratio of C-terminal detection (full length readthrough) to N-terminal detection (both full-length and truncated protein) of samples containing a TAG reporter. This is further divided by the ratio of C-terminal detection to N-terminal detection with the corresponding WT reporter; both of these sets of measurements are made in biological triplicate. For all 3 reporter + strain combinations evaluated here, RRE increased in cells transformed with the eRF1 variants S30P and N58A in comparison to wild-type controls. Changes in RRE were more pronounced with the yeast display reporter, where RRE is increased relative to controls for all four eRF1 variants tested. This may be attributable to the use of LeuOmeRS with the display reporter, which we have previously reported to exhibit more efficient OmeY incorporation than TyrOmeRS (used with the fluorescent reporter)^67^. In the case of fluorescent reporter BXGTAG, we have observed that the inherently high RRE of the system tends to limit the extent to which RRE values can be improved further^64,66,67^. Since RRE data cannot be subjected to conventional statistical analysis^66^, we evaluated changes in the median fluorescence intensity (MFI) of full-length protein detection using a one-way analysis of variance (ANOVA) (Fig. S1B). Under the conditions used here, we observe statistically significant increases in fluorescence for cells expressing the S30P and N58A variants with the yeast display reporter, with median fluorescent intensity (MFI) values increasing by 27% and 25% respectively, relative to WT. However, the fluorescence levels observed for the other conditions evaluated here did not reach statistical significance. We attribute this to the retention of WT eRF1 expression in the strains and surmise that the phenotypic effects of expressing a plasmid-based eRF1 mutant alongside WT eRF1 are less pronounced than when the mutant is expressed alone.

Although the changes in ncAA incorporation efficiency appear to be somewhat modest, we conducted model cell sorting experiments to determine if these changes were sufficient to support enrichments of improved variants using simple live cell recovery. We used BY4741 cells transformed with fluorescent reporters and attempted to isolate cells containing plasmids encoding either S30P or N58A mutations from a large excess of cells transformed with plasmids encoding wild-type eRF1. After inducing individual variants separately, we prepared 1:10 and 1:100 variant:WT ratios of cells cotransformed with either BXGTAG or BXGAltTAG reporters. Cell sorting was performed to isolate individual cells exhibiting the highest levels of GFP fluorescence (presumably corresponding to high levels of full-length reporter construct) which were then grown to saturation. After dilution and induction, the enriched populations were subjected to flow cytometry analysis. Two-dimensional dot plots of analyzed samples revealed improvements in full length detection relative to control WT eRF1 populations with the corresponding reporters (Fig S2A). To estimate enrichments at the genetic level, we isolated and sequenced ten clones from each recovered 1:100 population (Fig. S2B). In three out of four populations, we observed a 10-20 fold increase of variant eRF1 clones. These enrichments demonstrate that the plasmid-based eRF1 mutagenesis system supports enrichment for variants that support improved ncAA incorporation using standard cell sorting and growth recovery approaches. Overall, flow cytometry analysis and model enrichment experiments indicate that parallel, plasmid-based expression of eRF1 variants is sufficient to identify variants that support enhanced ncAA incorporation.

### Screening a randomly mutagenized eRF1 library for variants that enhance ncAA incorporation

Having demonstrated that the parallel expression approach is suitable for eRF1 engineering, we sought to conduct high throughput screening to identify additional eRF1 variants that enhance ncAA incorporation. To survey a broad range of candidate mutations, we performed error-prone PCR (epPCR) on the full WT SUP45 gene. Mutagenized DNA encoding SUP45 and a separately linearized vector backbone for SUP45 expression were co-electroporated into *S. cerevisiae* strain RJY100 along with the single plasmid system yeast display reporter/OTS plasmid (Table 1)^64^. We also attempted to prepare eRF1 libraries in cells transformed with single plasmid systems encoding fluorescent reporters^64^, but our electroporations yielded much lower numbers of transformants than for libraries prepared with cells containing the display reporter; we did not pursue eRF1 mutant libraries with fluorescent reporters further. Colony counting of plated library samples indicated that the mutagenized SUP45 library contained approximately 10^6^ transformants. Flow cytometry analysis indicated that the naïve library exhibited a broader range of fluorescence levels than levels observed in a control population transformed with the plasmid encoding WT eRF1, preliminarily suggesting some level of phenotypic diversity (Fig. S3A). Individual colonies were isolated from the library and 10 clones were subjected to full-plasmid sequencing to investigate genetic diversity. The clones contained between 3 and 9 nucleotide mutations, with an average of approximately 4 nucleotide mutations per clone, not all of which resulted in amino acid changes (Fig S3B, C). Deep sequencing of the naïve library indicated a broad distribution of mutations throughout the SUP45 gene, confirming that the overall mutagenesis scheme was successful (Fig. S3D, Extended Data File—excel file of DMS data). Despite potential overrepresentation of some individual clones in the library, the extensive diversity present within the library made it suitable for high-throughput screening.

We performed three rounds of fluorescence-activated cell sorting (FACS) to screen for clones exhibiting enhanced ncAA incorporation in response to UAG. The gating strategies used for all three rounds of positive sorts are shown in Fig S4A, and flow cytometric analysis of enriched populations are shown in Fig S4B. We used multiple gates in Rounds 1 and 3 of sorting in an attempt to maximize the diversity of clones with improved ncAA incorporation phenotypes isolated from the library. As sorting progressed, flow cytometry analysis indicated modest but detectable increases in the levels of C-terminal detection (c-Myc) relative to wild-type eRF1 (Fig. S4B), suggesting enrichment of clones exhibiting improved ncAA incorporation. We next isolated individual clones from three of the enriched populations, sequenced them, and evaluated ncAA incorporation via flow cytometry (Fig. S5, Table 2). Two-dimensional dot plots revealed multiple clones exhibiting putatively enhanced ncAA incorporation efficiencies, indicated by rightward shifts in c-Myc detection levels in comparison to controls (Fig. S5; see especially the boxed flow cytometry dot plots). Full plasmid sequencing of isolated clones (Table 2) revealed a diverse set of individual clones, most of which contained two or more amino acid mutations. Several of the amino acid mutations identified within these clones are identical to previously reported mutations known to enhance stop codon readthrough with near-cognate tRNA anticodons. In addition, we identified mutations at positions where amino acid substitutions that enhance stop codon readthrough are known, but the amino acid changes we observed have not been previously reported^43,45,46,49,68–70^. Both the presence of these types of mutations and the observed phenotypic enhancements suggested that our screening approach yielded variants with enhanced ncAA incorporation properties. In addition, many of the mutations present in these individual clones are at positions where no prior mutants that enhance stop codon suppression have been reported (to the best of our knowledge), motivating more detailed analysis.

To more fully elucidate the range of mutations enriched across the SUP45 gene, we subjected the naïve library and two tracks of sorted library populations (BR3T and CR3T) to deep sequencing (Fig. 2, S6-S8). In our analysis, we focused on highly enriched mutations resulting in amino acid point mutations in eRF1 because we used Illumina-based sequencing of four ∼300bp fragments (SUP45 length: ∼1.3 kb); this prevented coevolution analysis across the whole gene. Numerous mutations observed in the naïve library are depleted following enrichments, consistent with the expectation that most random mutations are neutral or deleterious to protein function^71,72^. However, at a broad subset of eRF1 positions, we observed one or more enriched mutations (Fig. 2C, D, Fig. S6-S8). These positions, key functional motifs of eRF1, and positions mutated in individual clones (Fig. 3, S5, Table 2) are depicted in a structure of WT *S. cerevisiae* eRF1 (Fig. 2C; PDB ID: 4CRN) and summarized in a heatmap of data from selected positions (Fig. 2D, S6). Enriched mutations are distributed throughout the protein structure, even though only the N-terminal domain is directly responsible for stop codon recognition^38,46,48^. The N-terminal domain harbors three out of four highly conserved eRF1 motifs: The GTS loop (28-30), TASNIKS motif (55-61), and the YxCxxxF motif (122-128). Several enriched mutations in this domain are found within these motifs (<u>G28A</u>, <u>S30P</u>, Q46E, E52K, <u>N58H</u>, <u>I59V</u>, <u>S61P</u>, N64D, K105R, P116L, <u>Y122N</u>, <u>C124R</u>, <u>N126S</u>). Interestingly, enrichments of variants in the other two domains (F169V, N213T, A227D, F239I, Y255N, Y255D, L272W, D310Y, G381A, G405A) are comparable or higher than the enrichments of N-terminal domain mutants (Fig. 2B, 2D). This raises the possibility that mutations to the middle domain (polypeptide chain release) and C-terminal domain (interactions with eRF3) are capable of enhancing ncAA incorporation. In one of the two enriched populations analyzed, we even observed enrichment of the mutation Q182E (Fig. S8). This position is part of the highly conserved GGQ motif, which has a direct role in the polypeptide chain release mechanism^37,73^.

**Figure 2.**
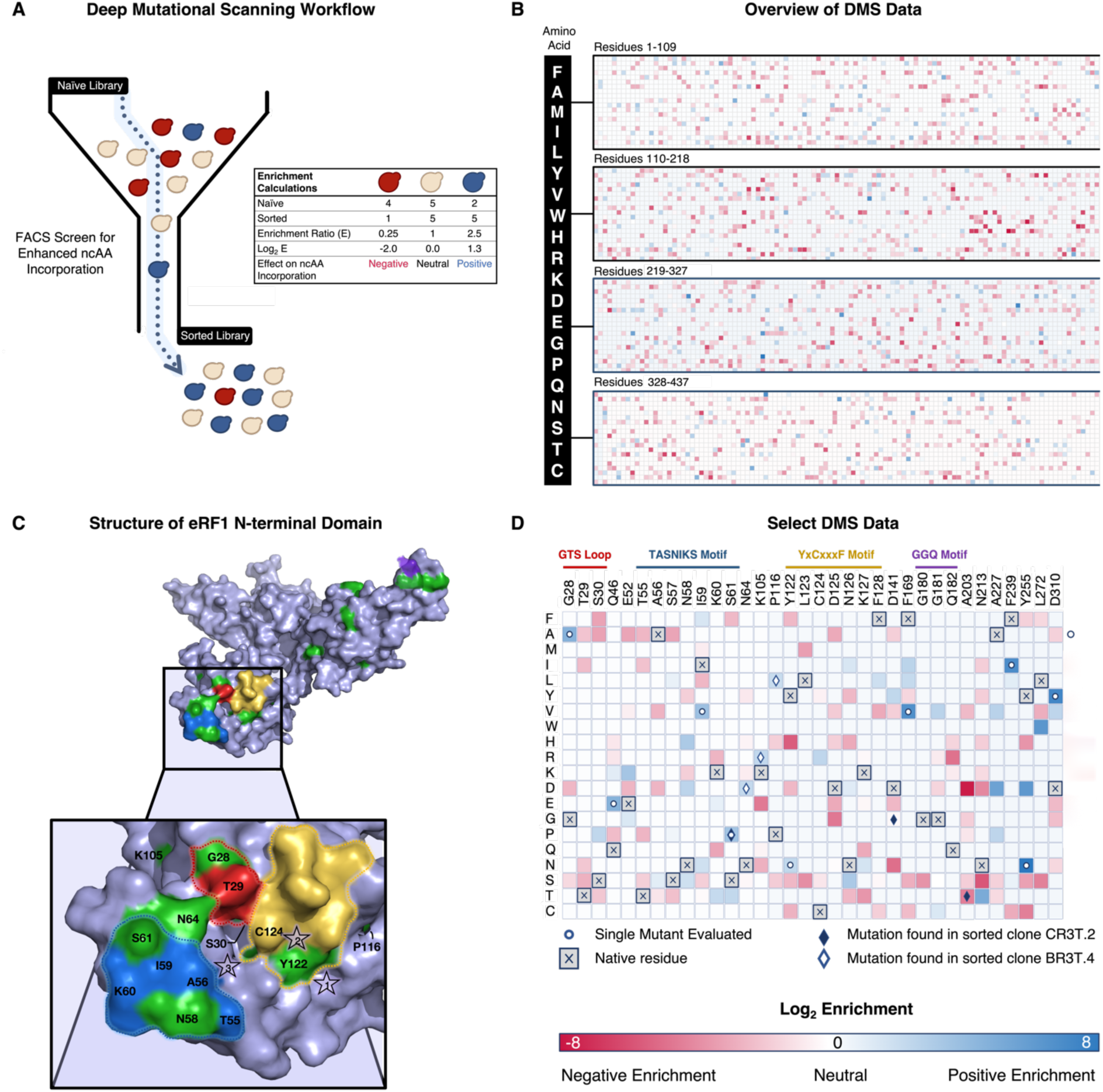
Deep mutational scanning of eRF1 for enhanced ncAA incorporation. A) Overview of workflow to identify enriched mutations based on a screen for enhanced ncAA incorporation. A library of eRF1 mutants is deep sequenced before and after flow cytometry-based screens for enhanced ncAA incorporation. Enrichment ratios and corresponding Log_2_ values are calculated to determine effect of mutation on ncAA incorporation. For a given mutation, Log_2_ (Enrichment) less than zero indicates that the mutation was deenriched after screening, Log_2_ (Enrichment) of zero indicates no change in mutational frequency after screening, and Log_2_ (Enrichment) greater than zero indicates enrichment of the mutation after screening. B) Log_2_ (Enrichment) values for mutations across entire SUP45 gene for Track BR3T. C) Overview of structure of *S. cerevisiae* eRF1 (PDB Id: 4CRN) with detailed view of the stop codon binding site. Highly conserved motifs are outlined as follows: TASNIKS 55-61 (blue), GTS 28-30 loop (red), YxCxxxF 122-128 (yellow) GGQ 180-182 (purple). Green residues highlight positions of interest according to the DMS data in panel C. Stars are numbered according to stop codon nucleotide positions. D) Highest positive values of enrichment and areas of interest on Track BR3T graphed using Morpheus^114^. Highly conserved eRF1 motifs in *Saccharomyces cerevisiae* are noted above the grid. Deep sequencing of Track CR3T for these positions can be found in Figures S6B, S8).

**Figure 3.**
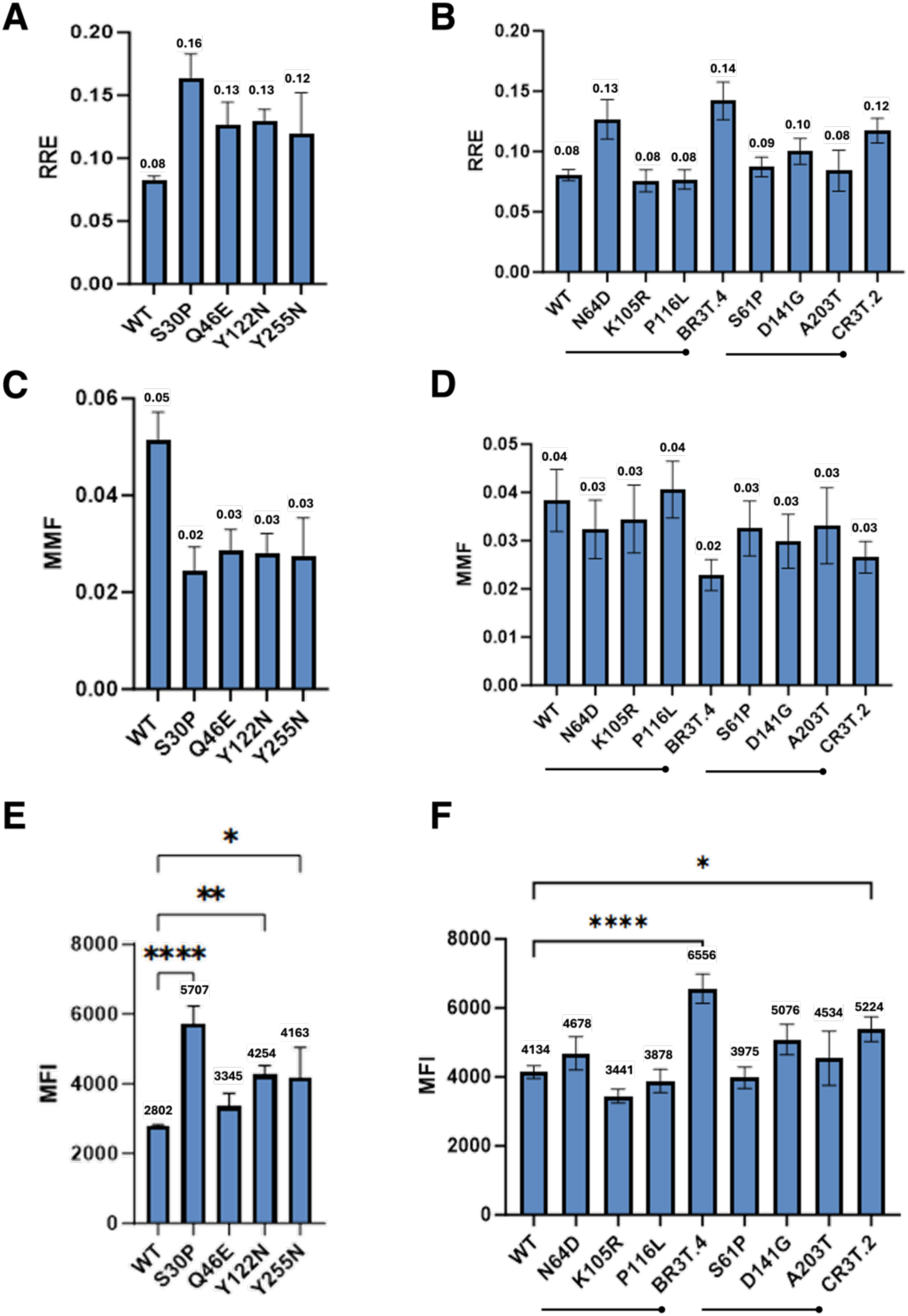
Analysis of effects of eRF1 mutants on ncAA incorporation efficiency and fidelity. A selection of variants enriched in the DMS analysis (S30P, Q46E, Y122N, and Y255N) as well as two multi-mutant clones isolated from sorted libraries alongside their individual components were evaluated for readthrough efficiency (RRE), fidelity (MMF), and stop codon readthrough levels (MFI). Sorted clone BR3T.4 contains mutations N64D, K105R, and P116L. Sorted clone CR3T.2 contains mutations S61P, D141G, and A203T. Each sorted clone is shown alongside the the corresponding single-point mutations contained in the triple mutant. A-B) Relative Readthrough Efficiency (RRE) defined as (TAG C-terminus detection/ TAG N-terminus detection)/(WT C-terminus detection/ WT N-terminus detection) for top enriched variants (left) and two sorted clones, BR3T.4 and CR3T.2, along with RRE and MMF for each of the corresponding single mutations (right). C-D) Maximum Misincorporation Frequency (MMF) defined as (RRE_-ncAA_)/ (RRE_+ncAA_) for top enriched variants (left) and two sorted clones, BR3T.4 and CR3T.2, with each of their corresponding single mutations (right). E-F) Median Fluorescence Intensity (MFI) of C-terminal (GFP) detection in N-terminal (BFP) positive cells. One-way ANOVA statistical analysis shown with the following p-value assignments: **** p < 0.0001,*** p < 0.001, ** p < 0.01, * p ≤ 0.05. Y255N was analyzed in biological duplicate and remaining clones were analyzed in biological triplicate. As discussed in the main text, no statistical analyses are reported for RRE and MMF determinations because they are not compatible with conventional statistical analyses.

Within our data, we observed several mutations previously identified in literature: S30P, I32F, V68A, D110G^43^, N126D^44^, and N126S^74^. A Y122F substitution has been shown to enhance stop codon readthrough in prior studies^44,68,75^. In our enrichment data, we observe an increase in Y122N, but a reduction of Y122F. Mutation E52D (E55D in human numbering) has been widely reported to enhance ncAA incorporation efficiency in mammalian cells^44,49–52,68,76–82^. Consistent with these studies, we observed improved ncAA incorporation in E52D relative to WT eRF1 in our system (Fig. S11). However, despite this validation of improvement in yeast, E52D was de-enriched in our deep sequencing datasets compared to other mutations at the same site: both E52K and E52G appeared to be enriched to some extent. It is crucial to highlight that our screening was conducted under distinct conditions from experiments performed in prior mutational studies of eRF1, including parallel expression of eRF1 mutants alongside WT eRF1 and evaluation of ncAA incorporation in response to the amber codon rather than nonsense suppression events with endogenous tRNAs and canonical amino acids. Furthermore, some of the best-characterized mutants reported previously were identified via rational design and targeted testing rather than high throughput screens. In addition to observing distinct mutations at previously identified positions of interest, we identified highly enriched mutations at numerous positions that have not been reported, except in the case of highly divergent eRF1 sequences (to the best of our knowledge; see Table S1 and further discussion below). However, it is important to recognize that most sorted clones harbor multiple mutations, some of which may co-occur with the beneficial mutations rather than being directly implicated in the observed phenotype. Altogether, these results demonstrate the power of our high throughput screening approach while motivating investigation of the effects of individual mutations on ncAA incorporation efficiencies.

### Functional impact of eRF1 variants on ncAA incorporation

To further our understanding of the effects of specific mutations, we investigated the impact of individual, highly enriched eRF1 variants on ncAA incorporation. We identified ten single mutants enriched via deep sequencing analysis (G28A, Q46E, I59V, S61P, Y122N, F169V, F239I, Y255N, D310Y, S365F) and prepared the plasmid-encoded eRF1 variants prior to evaluation of ncAA incorporation efficiencies of the resulting clones. We also attempted to evaluate an eRF1 variant containing the enriched Q182E mutation, but we were unable to reproducibly propagate cells expressing this variant, even with a genomic copy of WT eRF1 intact. Post-translational methylation of the GGQ motif stabilizes eRF1 and promotes its release function, suggesting that disruptions to this modification could enhance stop codon readthrough^41,69,83–86^. The enrichment of a charged residue (i.e. Q182E) in selection screens supports a potential role in modulating methylation; however, such mutations may also destabilize eRF1, leading to loss of cell viability which highlights the trade-off between enhanced incorporation and structural instability. This is consistent with previous reports that changes to the highly conserved GGQ motif resulted in the generation of unstable strains^36, 37, 63^.

We first conducted flow cytometry evaluations with each of the ten clones transformed into the BY4741 yeast strain alongside the AltTAG reporter pRS416-BXGAltTAG-TyrOmeRS (Fig. S9; see Materials and Methods for further details). After induction in the presence of 1 mM OmeY, inspection of flow cytometry dot plots indicated that cells transformed with the mutants Q46E, Y122N, and Y255N exhibited apparent improvements in ncAA incorporation compared to a WT control shown via a shift in C-terminal detection along the x-axis (Fig. S9A). Further analysis of the median fluorescence intensity levels of full-length reporter indicated that most of the mutants tested exhibited modestly increased levels of full-length protein in comparison to controls, with some mutants enabling statistically significant increases under these conditions (Fig. S9B).

The Q46E, Y122N, and Y255N single mutants, the previously characterized S30P mutant, widely reported E52D variant^44,49,51,52,79,87^, and two individual clones exhibiting enhanced ncAA incorporation (BR3T.4 and CR3T.2; Fig. 2) were then evaluated for relative readthrough efficiency (RRE) and maximum misincorporation frequency (MMF) (Fig. 3A-D, Fig. S10, Fig. S11)^88^. Calculations of RRE and MMF indicated that all of the clones evaluated exhibited higher readthrough efficiencies and lower misincorporation events than the corresponding WT control.

To statistically analyze these observations, we performed one-way ANOVA of the median fluorescence intensity (MFI) values using the WT control for comparison (Fig. 3E, F; as we have noted in prior work, RRE and MMF values cannot be directly evaluated for statistical significance^66^). For all clones evaluated except Q46E, the analysis indicates a statistically significant increase in full-length reporter construct detection (S30P, Q46E, Y122N, and Y255N exhibited 104%, 19%, 52%, and 49% increases in MFI respectively). Interestingly, clones BR3T.4 and CR3T.2, which each contain 3 amino acid mutations, exhibit similar levels of ncAA incorporation to the S30P clone, with the other point mutations resulting in lower levels of improvement. Five of the six mutations in these clones (N64D, S61P, K105R, P116L, A203T) were modestly enriched (Log_2_ enrichment values ranging from 2.00 – 2.60) in our deep sequencing analysis, whereas D141G was mildly de-enriched (Log_2_ enrichment of -0.60). In this set of experiments, BR3T.4 exhibited a 59% increase in MFI relative to WT, whereas the mutations within that clone (N64D, K105R, and P116L) had MFI changes of +13%, -17%, and -6% respectively. CR3T.2 had a 26% increase in MFI relative to WT, whereas the mutations within that clone (S61P, D141G, and A203T) had MFI changes of -4%, +23%, and +10% respectively. These observations suggest that the combined effects of multiple mutations can substantially exceed the impact of individual substitutions, indicating that the context-dependent interactions between amino acid changes contribute to enhanced ncAA incorporation. Overall, investigating individual mutants identified during screening revealed that point mutations can exhibit a range of effects on ncAA incorporation. The varied properties of point mutations and clones containing multiple point mutations that we identified highlight the value of the high throughput screening approach we used here to identify clones with desirable phenotypes of interest.

### Investigating the effects of eRF1 variants on ncAA incorporation in different genomic contexts

Since our screening platform and the characterizations above all utilize parallel expression of WT eRF1 (endogenous genomic locus) and mutant expression (plasmid), we also investigated ncAA incorporation with eRF1 variants (1) after shutting down expression of endogenous eRF1; and (2) in yeast strains with knockouts with already-enhanced ncAA incorporation due to previously identified gene deletions. To investigate the effects of eRF1 variants in the absence of endogenously expressed eRF1, we used a strain encoding SUP45 expression under the control of a Tet-Off promoter (Fig. 4A). The expression of eRF1 in haploid strain YTH41 (derivative of CML476, *MAT***a** ura3-52 leu2-Δ1 his3-Δ200 GAL2 LEU2::CVp (TetR’-SSN6) trp1::tTA sup45::kanMX4-tetO7-SUP45; gift from Dr. Tobias von der Haar) can be shut off by supplementation of media with doxycycline, enabling propagation of strains during expression of WT eRF1, but eliminating WT eRF1 expression prior to ncAA incorporation assays (Fig. 4A)^89^. We were able to transform and propagate YTH41 with AltTAG reporter pRS416-BXGAltTAG-TyrOmeRS and each of three plasmid-encoded variants of eRF1: WT eRF1, eRF1 N58A (point mutation evaluated in Figs. S1 and S2 exhibiting enhanced TAG codon readthrough), and the individual clone BR3T.4 (top-performing individual clone evaluated in Figs. 3, S5-S7). The eRF1 variants were still expressed from the native promoter, but due to the lack of an available Leu selection marker, the variants were cloned into a pRS413 vector backbone (His selection marker). Even in the absence of Dox, these transformants exhibited slow growth phenotypes, presumably due to reduced viability as a result of placing wild-type SUP45 under the control of a nonnative promoter (this reduction in viability hindered transformation and characterization of additional eRF1 variants in YTH41). Nonetheless, we were able to conduct flow cytometry assays to evaluate ncAA incorporation efficiency. Consistent with our experiments in BY4741, constitutive expression of eRF1 variants in YTH41 led to increased levels of full-length reporter even in the absence of Dox (73% increase in MFI of N58A and 50% increase in MFI of BR3T.4 relative to the WT control within BY4741). Upon treatment of cells with 10 µg/mL Dox at the beginning of reporter protein induction (16 hours prior to analysis), the amount of full-length reporter detected increases further with N58A exhibiting a 148% increase in MFI relative to WT plasmid-based eRF1, and BR3T.4. exhibiting a 181% increase in MFI relative to WT plasmid-based eRF1 (Fig. 4B, Fig. S12). These results indicate that the eRF1 variants substantially enhance ncAA incorporation efficiencies in the absence of endogenously expressed eRF1.

**Figure 4.**
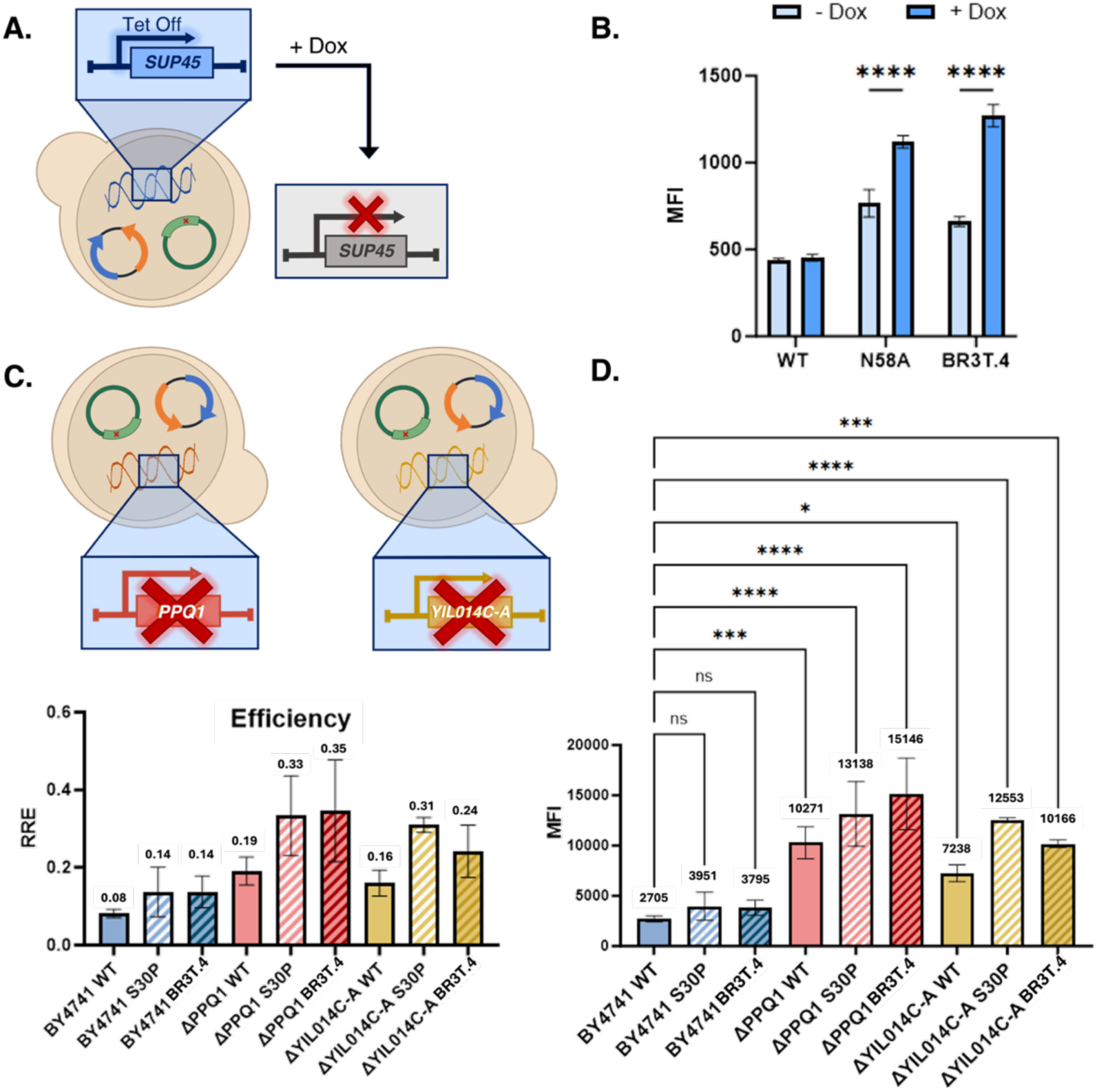
Investigation of mutant eRF1 activities in different cell strains. A) Schematic of the Tet-off strain YTH41 shutting off endogenous SUP45 in the presence of doxycycline, enabling evaluation of the phenotype of eRF1 variants expressed from plasmids. B) Investigation of ncAA incorporation in YTH41 transformed with WT, N58A, or BR3T.4 (isolated triple mutant with N64D, K105R, and P116L) and AltTAG reporter. Median fluorescence intensity (MFI) of full-length reporter protein in BFP-positive cells. Inductions of reporters were performed in the presence of 1 mM OmeY and either in the presence or absence of 10 μg/mL doxycycline (dox). Two-way ANOVA statistical analysis shown with the following p-value assignments: ****p ≤ 0.0001. Wild Type clones were analyzed in biological duplicate and mutant clones were analyzed in biological triplicate. Flow cytometry dot plots can be found in Figure S13. C) Schematic of the BY4741 derived knockout strains ΔPPQ1 and ΔYIL014C-A which were analyzed in tandem with eRF1 variants for enhanced ncAA incorporation phenotypes. Relative readthrough efficiencies (RRE) of cells transformed with wild type (WT) eRF1, S30P (single mutant clone) and BR3T.4 in yeast strains BY4741, BY4741ΔPPQ1, and BY4741ΔYIL014C-A. D) Median Fluorescence Intensity (MFI) of C-terminal (GFP) detection in BFP-positive cells. One-way ANOVA statistical analysis shown with the following p-value assignments: ****p ≤ 0.0001, *** p ≤ 0.001, * p ≤ 0.05, ns p > 0.05. For panels (C) and (D), three biological replicates were used for each condition investigated. As discussed in the main text, no statistical analyses are reported for RRE and MMF determinations because they are not compatible with conventional statistical analyses.

We also investigated the effects of plasmid-based expression of eRF1 variants in yeast strains containing gene knockouts that enhance ncAA incorporation. Previously, we have reported that BY4741ΔPPQ1^66^ and BY4741ΔYIL014C-A^65^ each exhibit improved ncAA incorporation in comparison to the parent BY4741 strain. Here we investigated whether combining these knockouts with eRF1 variants would further enhance ncAA incorporation and amplify the impact of subtle improvements that are not individually statistically significant. Each of the knockout strains and a BY4741 control strain were transformed with AltTAG reporter pRS416-BXGAltTAG-TyrOmeRS and either WT eRF1, S30P (point mutation evaluated in Fig. S1 and S2 exhibiting enhanced TAG codon readthrough), or the individual clone BR3T.4 (top-performing individual clone evaluated in Figs. 3, S5). As expected, when comparing a control WT plasmid-based eRF1 across strains, the knockout strains exhibited enhanced ncAA incorporation (280% increase in ΔPPQ1 MFI and 168% increase in ΔYIL014C-A MFI, compared to the parent strain). NcAA incorporation was further improved when the knockout strains were transformed with eRF1 variants S30P and BR3T.4 (Fig. 4C, 4D, S13). BY4741 exhibited a 46% MFI increase when transformed with S30P relative to WT, and a 40% increase with BR3T.4 relative to WT. In ΔPPQ1, S30P and BR3T.4 exhibited a 28% and 48% increase respectively relative to WT within the same knockout (KO) strain, and the combined KO + mutants resulted in a 386% and 460% increase in MFI respectively, relative to the BY4741 WT control. In ΔYIL014C-A, S30P and BR3T.4 exhibited a 73% and 40% increase respectively relative to WT within the same knockout (KO) strain, and the combined KO + mutants resulted in a 364% and 276% increase in MFI respectively, relative to the BY4741 WT control. A one-way ANOVA of the MFI data (Fig. 4D; see Materials and Methods for details) indicated that BY4741 transformed with eRF1 variants did not exhibit significant enhancements in this set of experiments. We attribute the lack of significance in this experiment to the subtle effects of introducing plasmid-based eRF1 variants into BY4741, which were overshadowed by the more pronounced impacts of gene deletions on ncAA incorporation. The high error associated with this small number of replicates hindered the ability to detect significance using ANOVA. However, upon performing a separate experiment with 12 biological replicates instead of 3 per condition, statistical analysis indicated significant changes in reporter levels, confirming the enhancement of ncAA incorporation by S30P and BR3T.4 in BY4741 (Fig. S13C). Analysis of the RREs and reporter levels in the knockout strains indicate that the introduction of the eRF1 variants enhances ncAA incorporation beyond the levels achieved by the strains alone (Fig. 4C-D, Fig. S13). The further improvement of ncAA incorporation when combining eRF1 variants and gene knockouts suggests that these two approaches are complementary to one another, and could even operate by distinct pathways or mechanisms (though further experimentation would be needed to confirm this hypothesis). Our data indicates that eRF1 variants can drive enhancements to ncAA incorporation as the sole release factor present in the cell as well as within the context of strains already known to support elevated levels of ncAA incorporation.

### Validation of ncAA incorporation in response to amber stop codon with mass spectrometry and flow cytometry experiments

To ensure our that systems are in fact incorporating a ncAA of interest in response to the amber (UAG) codon and not exhibiting enhanced fluorescent phenotypes due to near-cognate suppression events^90–92^, we evaluated the presence of *O-*methyl-L-tyrosine (OmeY) in tryptic digests of a secreted version of Donkey1.1 H54TAG via MALDI-MS (Fig. S14). This construct was grown and induced in cells alongside a plasmid-based eRF1 variant (either WT, Q46E, or the sorted clone BR3T.4) to determine whether eRF1 mutants result in detectable levels of aberrant cAA incorporation in response to the stop codon. Across conditions, m/z values corresponding to OmeY incorporation were readily detected, but no signals consistent with near-cognate cAA misincorporation were detected in the presence of eRF1 variants.

To further confirm ncAA incorporation in response to a UAG codon, we used click chemistry experiments on the yeast surface to evaluate the incorporation of the ncAA *p*-propargyloxy-L-phenylalanine (OPG). Following co-transformation of a set eRF1 variants alongside the single-plasmid yeast display reporter/OTS pCTCON2-FAPB2.3.6L1TAG-LeuOmeRS, cells were induced in the presence of either OPG or a non-clickable control amino acid (OmeY). Cells were subsequently labelled via copper-catalyzed azide-alkyne cycloaddition (CuAAC) with an azide-functionalized biotin probe, enabling detection by a fluorescently labelled antibody (PE anti-Biotin). For all cells transformed with eRF1 variants, the biotin detection was ∼1000X higher in samples induced with OPG relative to the OmeY control (Fig. S15). The strong enrichment of signal under clickable conditions, coupled with minimal background in the control, confirms selective incorporation of the ncAA and argues against substantial contributions from near-cognate suppression. These data indicate that the enhanced fluorescence observed in reporter systems with engineered eRF1 variants reflects improved ncAA incorporation, rather than reduced fidelity, supporting a model in which modulation of termination primarily shifts the competitive balance between release factors and suppressor tRNAs without appreciably affecting near-cognate suppression.

## DISCUSSION

In this study, we established a high throughput screening approach to identify variants of eukaryotic release factor 1 (eRF1) that enhance noncanonical amino acid (ncAA) incorporation. Our approach is distinct from prior work, where eRF1 variants were evaluated for the ability to promote aberrant stop codon readthrough facilitated by endogenous tRNAs (noncognate with respect to stop codons), but not suppression with a cognate suppressor tRNA. Another distinguishing feature of our approach is the plasmid-based encoding of eRF1 variants alongside the expression of endogenous WT eRF1. This enabled the identification of a broad array of mutations that enhance ncAA incorporation, even in cases where mutations were previously shown to compromise cell viability^37,43,93^. However, we note that with parallel expression we could not study all point mutations of interest, such as the Q182E mutation within the highly conserved GGQ motif (discussed above). Within these limitations, both direct isolation of individual clones from the library (containing multiple mutations) and identification of enriched point mutations via deep sequencing yielded variants that support enhanced ncAA incorporation, indicating the versatility of our approach for screening eRF1 variants of interest. We found a broad set of mutations, ranging from previously described amino acid changes to mutations at positions not previously investigated in the context of UAG readthrough events.

Our findings affirm the importance of the N-terminal domain of eRF1 while also highlighting opportunities to investigate the roles of the other eRF1 domains in stop codon readthrough. Consistent with prior studies, we identified mutations in the GTS loop, YxCxxxF motif, and GGQ motif that were enriched during screening. When it was possible to characterize the properties of variants, we typically observed ∼1.5 to 2.0-fold increases in full-length protein levels indicative of enhanced ncAA incorporation efficiency. Isolated clones, which included combinations of mutations in these motifs with other mutations (either in the N-terminal domain or elsewhere) exhibited closer to 2.0-fold increases in ncAA incorporation efficiency. Interestingly, while point mutations located in the middle and C-terminal domains conferred measurable improvements in ncAA incorporation, these enhancements were generally smaller than those observed for individual N-terminal domain variants. Nevertheless, the enrichment and modest functional benefits associated with these middle and C-terminal mutations, together with the properties of individual multi-mutation clones, suggest that combining substitutions across domains could produce additive or even synergistic effects on stop codon readthrough and ncAA incorporation. The tools of protein engineering and design provide numerous opportunities to evaluate the effects of combinatorial mutagenesis of eRF1^94–97^.

In addition to laboratory-prepared libraries based on observations from eRF1 mutagenesis, naturally occurring eRF1 homologs from alternative code organisms provide valuable insights into how stop codon readthrough and ncAA incorporation may be enhanced in higher order eukaryotes. Many of these organisms encode for divergent eRF1 variants (Fig. S16) that facilitate selective readthrough of canonical stop codons, a trait that could inform future diversification strategies for ncAA incorporation. Such selective suppression may reflect co-evolution of these divergent eRF1 variants with distinct local codon contexts flanking termination sites, as neighboring sense codons are known to modulate release factor engagement, suggesting that systematic assessment of UAG readthrough across varied codon contexts in the background of the evolved eRF1 mutants discovered here posits an important future direction^39,54–59^. Several mutations found in these variants were identified in our screens, as summarized in Table S1^39,40,45,54,55,57,58,93,98–104^. Parallel expression of these divergent proteins alongside wild-type eRF1 could overcome limitations observed in prior characterizations of these release factors^43,98,100^. Expanding the range of eRF1 functions in yeast and other eukaryotes will require new tools to assess readthrough at codons beyond UAG; this is crucial for understanding eRF1’s broader role as an omnipotent release factor. For example, our recent work in *S. cerevisiae* enabling ncAA incorporation at UGA, along with similar approaches for UAA and rare cognate codons in mammalian cells, provides a foundation for further exploration of selective readthrough mechanisms^49,105–107^. The approaches described in this work make it possible to conduct comprehensive studies that elucidate specific eRF1 features that govern codon selectivity and advance our fundamental understanding of this essential translation process.

Finally, our findings have potential uses in synthetic biology and biomanufacturing. Eukaryotic release factors with altered stop codon recognition capabilities are a required component of genomically recoded eukaryotic organisms. The ongoing genome engineering work of the Sc2.0 project has led to substantial progress in establishing an “amberless” yeast strain; alteration of eRF1 codon recognition is complementary to the Sc2.0 work as it is expected to reduce or eliminate termination events at UAG. It may be necessary to evaluate many eRF1 variants to identify ones that support both ncAA incorporation in response to UAG while also maintaining a high level of organismal fitness. The dual reporters used in this study and others can support such efforts. In biomanufacturing settings, engineered release factors may be valuable tools for enhancing the production yields of proteins containing ncAAs. Since heterologous protein expression does not necessarily require maintenance of cell viability, it may even be possible to induce the expression of eRF1 variants that reduce viability in host cells only when biosynthesis of ncAA-containing proteins is occurring. Overall, this work expands the potential for engineering eRF1 to improve ncAA incorporation, with broad implications for both fundamental understanding of protein biosynthesis and bioengineering applications of genetic code expansion.

## MATERIALS & METHODS

### Materials

Q5 2× Master Mix and restriction enzymes were purchased from New England Biolabs (NEB). Synthetic oligonucleotides for cloning and sequencing were purchased from GENEWIZ (Azenta Life Sciences). Pellet Paint NF co-precipitant was purchased from Millipore Sigma. *E. coli* plasmid purification was performed using Epoch Life Science GenCatch Plasmid DNA Mini-Prep Kits. Yeast plasmid DNA purification was performed using Zymoprep Yeast Plasmid Miniprep II kits. Gel extraction and purification was performed using an Epoch Life Science GenCatch Gel Extraction Kit. Chemically competent yeast and plasmid DNA transformations were prepared using Zymo Research Frozen-EZ Yeast Transformation II kits. *O*-Methyl-L-tyrosine (OmeY) was purchased from Chem-Impex International. O-Propargyl-L-tyrosine hydrochloride (OPG) was purchased from Iris Biotech. PE anti-Biotin antibody was purchased from BioLegend. Mass-spec grade Promega Trypin Gold was purchased from Fisher Scientific. Reagents for CuAAC (copper-catalyzed azide–alkyne cycloaddition): Biotin-PEG_3_-Azide, and THPTA were purchased from Click Chemistry Tools; (+)-Sodium L-ascorbate, aminoguanidine hydrochloride, copper sulfate pentahydrate, EDTA, dimethylformamide, and DMSO were purchased from Sigma-Aldrich. SDS-PAGE analysis was done with 4–12% Bis-Tris mini gels and SimplyBlue Safestain. Whole plasmid sequencing was performed by Plasmidsaurus using Oxford Nanopore Technology (Eugene, OR). Sanger sequencing and deep sequencing (AmpExpress) were performed by Quintara Biosciences (Cambridge, MA). MALDI-MS was performed by the Koch Institute Biopolymers and Proteomics Core (Cambridge, MA). All *S. cerevisiae* strains used in this work are described in Table 3.

**Table 3.** Genotypes of yeast strains used in this study.

| Yeast Strain | Genotype |
| --- | --- |
| RJY100 | MATa AGA1::GAL1-AGA1::URA3 ura3-52 trp1::KanMX leu2-Δ200 his3-Δ200 pep4::HIS3 prbd1.6R can1 GAL |
| BY4741 | MATa his3Δ1 leu2Δ0 met15Δ0 ura3Δ0 |
| BY4741ΔPPQ1 | MATa his3Δ1 leu2Δ0 met15Δ0 ura3Δ0 ppq1Δ::KanMX |
| BY4741ΔYIL014C-A | MATa his3Δ1 leu2Δ0 met15Δ0 ura3Δ0 yil014c-aΔ::KanMX |
| YTH41 | derivative of CML476; MATa ura3-52 leu2-Δ1 his3-Δ200 GAL2 LEU2::CVp(TetR'-SSN6) trp1::tTA sup45::kanMX4-tetO7-SUP45 |

**Table 4:**
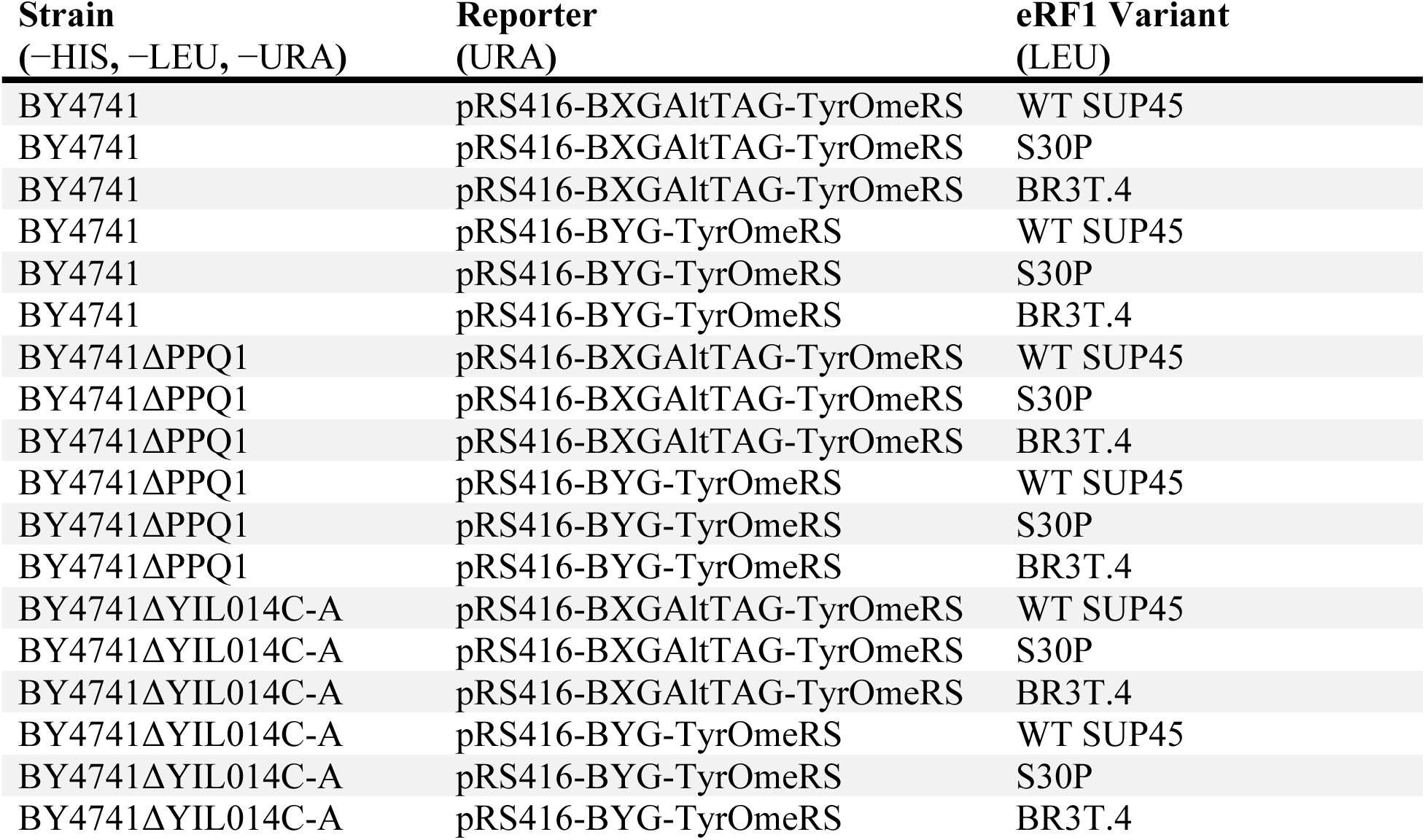
Transformed strains used to evaluate ncAA incorporation efficiency in combination with gene knockouts.

| **Strain**<br>(–HIS, –LEU, –URA) | **Reporter**<br>(URA) | **eRF1 Variant**<br>(LEU) |
| --- | --- | --- |
| BY4741 | pRS416-BXGAltTAG-TyrOmeRS | WT SUP45 |
| BY4741 | pRS416-BXGAltTAG-TyrOmeRS | S30P |
| BY4741 | pRS416-BXGAltTAG-TyrOmeRS | BR3T.4 |
| BY4741 | pRS416-BYG-TyrOmeRS | WT SUP45 |
| BY4741 | pRS416-BYG-TyrOmeRS | S30P |
| BY4741 | pRS416-BYG-TyrOmeRS | BR3T.4 |
| BY4741ΔPPQ1 | pRS416-BXGAltTAG-TyrOmeRS | WT SUP45 |
| BY4741ΔPPQ1 | pRS416-BXGAltTAG-TyrOmeRS | S30P |
| BY4741ΔPPQ1 | pRS416-BXGAltTAG-TyrOmeRS | BR3T.4 |
| BY4741ΔPPQ1 | pRS416-BYG-TyrOmeRS | WT SUP45 |
| BY4741ΔPPQ1 | pRS416-BYG-TyrOmeRS | S30P |
| BY4741ΔPPQ1 | pRS416-BYG-TyrOmeRS | BR3T.4 |
| BY4741ΔYIL014C-A | pRS416-BXGAltTAG-TyrOmeRS | WT SUP45 |
| BY4741ΔYIL014C-A | pRS416-BXGAltTAG-TyrOmeRS | S30P |
| BY4741ΔYIL014C-A | pRS416-BXGAltTAG-TyrOmeRS | BR3T.4 |
| BY4741ΔYIL014C-A | pRS416-BYG-TyrOmeRS | WT SUP45 |
| BY4741ΔYIL014C-A | pRS416-BYG-TyrOmeRS | S30P |
| BY4741ΔYIL014C-A | pRS416-BYG-TyrOmeRS | BR3T.4 |

### Media and ncAA Liquid Stock Preparation

Liquid and solid media were prepared as previously described^88,108^. SD-SCAA and SG-SCAA media was prepared in the following combinations: without tryptophan (TRP), leucine (LEU), and uracil (URA) (for experiments with RJY100 transformed with derivatives of plasmids pCTCON2 and pTH353 described below); without TRP and URA (for experiments with RJY100 transformed with derivatives of pCTCON2); without URA and LEU (for experiments with BY4741 transformed with derivatives of plasmids pRS416 and pTH353 described below); without URA (for experiments with BY4741 transformed with derivatives of pRS416); or without histidine (HIS) and URA (for experiments with YTH41 transformed with derivatives of plasmids pRS413 and pRS416 described below). *O*-methyl-L-tyrosine (OmeY) was prepared at a concentration of 50 mM L-isomer as previously described^88^.

### Plasmid Design and Construction

All single-plasmid systems (SPSs), which encode both a dual reporter and an orthogonal translation system on the same plasmid (pCTCON2-FAPB2.3.6L1TAG-LeuOmeRS (yeast display), pRS416-BXG-TyrOmeRS (intracellular), and pRS416-BXGaltTAG-TyrOmeRS (intracellular reporter)) have been previously described^64,109^.

The following plasmid-encoded eRF1 variants under the control of the native *S. cerevisiae SUP45* promoter were obtained from the laboratory of Tobias von der Haar via Addgene: Wild Type (pTH353-SUP45WT Addgene #29371), S30P (pTH399-SUP45-S30P Addgene #29372), N58A (pTH371-SUP45-N58A Addgene #29378), I222S (pTH397-SUP45-I222S Addgene #29388), and P174Q (pTH375-SUP45-P174Q Addgene #29383)^43^. Addgene entries contain the sequences of these plasmids.

To prepare plasmids, pRS413-SUP45-WT, pRS413-SUP45-N58A, and pRS413-SUP45-BR3T.4, plasmids pRS413^110^, pTH353-SUP45WT, pTH371-SUP45-N58A, and pTH353-SUP45-BR3T.4, were digested using SalI-HF and XbaI restriction enzymes. For each reaction, 10 μg of plasmid DNA was combined with 10 μL of 10× Cutsmart, 3 μL of SalI-HF, 3 μL of XbaI and sterile water to a final volume of 100 μL. Reactions were prepared in separate microcentrifuge tubes, thoroughly mixed by pipetting, and incubated for ∼4 hours at 37 °C. The digested products were run on a 0.8% agarose gel. For pRS413 the band corresponding with the backbone (4913 bp) was extracted, and for the eRF1 WT and variant constructs, the insert bands (2999 bp) were extracted. Insert and backbone DNA was purified using the Epoch Life Science GenCatch Gel Extraction Kit. Backbone and insert DNA was ligated at a 1:5 molar ratio for each construct. For each 20 μL reaction, 2 μL of 10× T4 DNA ligase buffer, 100 ng of backbone DNA (33.05 fmol), a 5× molar ratio of insert DNA (165.25 fmol), 1 μL of T4 DNA ligase, and sterile water to a final volume of 20 μL was added to a PCR strip tube and mixed thoroughly by pipetting. Each reaction was then incubated at room temperature for 10 min followed by a heat inactivation step at 65 °C for 10 min. Reactions were then chilled on ice for at least 5 min before transforming 5 μL of each reaction into 50 μL of competent *E. coli* strain DHαZ1 and plating on LB + 50 μg/ mL ampicillin. *E. coli* plasmid purification was performed according to manufacturer’s instructions using Epoch Life Science GenCatch Plasmid DNA Mini-Prep Kit then sent for Sanger Sequencing at Quintara Biosciences using primers Seq_his_SUP45_fwd and Seq_his_SUP45_rev to confirm successful incorporation of the eRF1 inserts. Sequences of constructs pRS413-SUP45-WT, pRS413-SUP45-N58A, and pRS413-SUP45-BR3T.4 are available in the supplementary information.

### Yeast Transformations, Propagation, and Induction

Transformations were performed according to the manufacturer’s protocol using Zymo competent *S. cerevisiae* RJY100, BY4741, or YTH41. Cells were plated on solid SD-SCAA media with the appropriate drop-out mixes as described above and grown with shaking at 30 °C for 2-4 days until colonies were visible. All quantitative experiments were performed in biological triplicate, unless otherwise noted. For RRE and MMF measurements, three separate transformants were inoculated for each experiment and grown in 5 mL SD-SCAA media using the same drop out mixes used in solid media for plating after transformation. Liquid cultures were supplemented with 100 IU penicillin and 100 μg/mL streptomycin. Once saturated, liquid cultures were diluted to an OD600 of 1 in fresh media and grown 4-8 hours to mid log phase (OD 2-5) with shaking at 30 °C. Cells were once again diluted to an OD of 1, pelleted at 2400 × g for 5 min, resuspended in induction media (SG-SCAA) with the appropriate drop out mix supplemented with 1 mM final concentration of L-isomer OmeY and grown for 16 hours at 20 °C with shaking.

### Reporter System Validation Experiments

RJY100 cells were transformed with pTH399-SUP45-S30P, pTH371-SUP45-N58A, pTH397-SUP45-I222S, pTH375-SUP45-P174Q, or WT eRF1 (backbone pTH353) alongside a SPS yeast display reporter with (pCTCON2-FAPB2.3.6L1TAG-LeuOmeRS) and without (pCTCON2-FAPB2.3.6-LeuOmeRS) a TAG sequence. BY4741 cells were transformed with pTH399-SUP45-S30P, pTH371-SUP45-N58A, pTH397-SUP45-I222S, pTH375-SUP45-P174Q, or WT eRF1 (backbone pTH353) alongside a SPS BXG or BXGaltTAG intracellular reporter, either with a TAG in the reporter construct (pRS416-TyrOmeRS-BXG) & pRS416-TyrOmeRS-BXGaltTAG), or without a TAG in the reporter construct (pRS416-TyrOmeRS-BYG). Transformed cells were propagated and induced as described above. Samples with yeast display reporters were labelled for flow cytometry as previously described^67^ using the following antibodies at a 1:500 dilution in PBSA: mouse anti-HA (BioLegend), mouse anti-c-Myc (Exalpha), goat anti-mouse Alexa Fluor 488 (Invitrogen), and goat anti-chicken Alexa Fluor 647 (Invitrogen). Samples with intracellular reporters were washed 3 times with PBSA prior to flow cytometry. For all samples, 2 million cells were prepared for flow cytometry using an Attune NxT flow cytometer (Life Technologies), of which 10,000 events were collected. RRE values were calculated for each sample as described previously^88^(Fig. S1).

### Model enrichments

BY4741 cells were transformed with pTH399-SUP45-S30P, pTH371-SUP45-N58A, or pTH353-SUP45WT alongside each of two intracellular reporter/OTSs (pRS416-BXGTAG-TyrOmeRS and pRS416-BXGAltTAG-TyrOmeRS). Samples were propagated and induced as described under “Yeast Transformations, Propagation, and Induction”. Following an overnight induction, mixtures of 1:10 and 1:100 variant:WT ratios were created for each reporter. For 1:10 mixtures, volumes corresponding to 500,000 cells of mutant eRF1 were combined with volumes corresponding to 5 million WT cells. In 1:100 mixtures, volumes corresponding to 50,000 cells of mutant eRF1 were combined with volumes corresponding to 5 million WT cells. Each of these mixtures were aliquoted into 1.5 mL microcentrifuge tubes, diluted to 1 mL of PBSA then centrifuged for 30 sec at 15,871 × g. The supernatants were aspirated and pellets were washed by resuspending in 1 mL PBSA, centrifuging at 15,871 × g for 30 sec, aspirating the supernatant, and repeating for a total of 3 washes. Samples were first characterized by Flow Cytometry using an Attune NxT Flow Cytometer (Thermo Fisher) then screened for enhanced ncAA incorporation via Fluorescence Activated Cell Sorting (FACS) using an S3e Cell Sorter (BioRad) with gates set to collect samples with high levels of C-terminal (GFP) detection. Approximately 2500 cells were collected per sample. Sorted samples were collected in 5 mL tubes with 1 mL SD-SCAA (−LEU −URA) supplemented with penicillin-streptomycin. Immediately following sorts, an additional 1 mL of media was rinsed along the inside of the collection tube. Following all sorts, recovery cultures were transferred to 14 mL culture tubes and additional media was added for a total of 5 mL. Recovered sorts were grown for 3 days in 30 °C shaking at 300 rpm until saturated, then propagated and induced as described above prior to being subjected to flow cytometry analysis. FCS files were analyzed using FlowJo as previously described^88^. In addition to flow cytometry characterizations, DNA from each of the recovered sorts (from the 1:100 mutant:WT starting ratios) was extracted and purified using a Zymoprep Yeast Plasmid Miniprep II kit. The miniprepped DNA was then transformed into chemically competent *E. coli* strain DH5αZ1: 50 μL of cells and the entire ∼30 μL yeast miniprep DNA output was combined in a 1.5 mL microcentrifuge tube per recovery DNA sample, mixed well by pipetting, and incubated on ice for 30 min. Tubes were then subjected to a 45 s heat shock at 42 °C followed by an additional 5 min incubation on ice. 500 μL of SOC media was then added to each tube, pipetted well to mix, then the tubes were incubated in 37 °C shaking for 1 hour. Following the 1 hour incubation, 200-500 μL of cells were plated on LB with 50 μg/mL ampicillin and incubated in 37 °C for 16 hours. 10 colonies were inoculated and miniprepped from each sample and sent for Sanger Sequencing (Quintara Biosciences) using the SUP45_Seq_Fwd primer to determine post-sort ratios of mutant: WT eRF1. Fold enrichments were calculated by dividing the post sort ratio of variant:WT by the starting ratios (1:100). Post sort ratios were determined by the number of clones sequenced containing the mutant eRF1 divided by 10 (total clones sequenced).

### Error-Prone PCR (epPCR) Library Construction

The mutant eRF1 library was generated via error prone PCR (epPCR) using a wild-type SUP45 open reading frame as a template. epPCR was performed by combining 1 μL template DNA (0.7 ng/ μL), 2.5 μL of 10 μM forward primer (SUP45_Cloning_Fwd), 2.5 μL of 10 μM reverse primer (SUP45_Cloning_Rev), 1 μL of 10mM dNTP, 5 μL of 20 μM dPTP, 5 μL of 20 μM 8-oxo-dGTP, 1 μL Taq DNA Polymerase, 5 μL 10× Taq reaction buffer, and 27 μL sterile water for a final reaction volume of 50 μL. The reaction was run on a thermocycler at 95 °C for 500 s followed by 16 cycles of 95 °C for 45 sec, 60 °C for 30 sec, and 72 °C for 117 sec. Once the cycles were complete, the reaction underwent a final extension for 10 min at 72 °C followed by a final hold at 4 °C until they were removed from the thermocycler. The amplified product was run on a 1% agarose gel, the correctly sized band was cut out of the gel, and the DNA purified using the Epoch Life Science GenCatch Gel Extraction Kit.

Following epPCR, the purified mutagenized gene was further amplified via PCR. The amplification process was performed by combining 100 ng of the epPCR mutated DNA template, 4 μL 100 μM forward primer, 4 μL of 100 μM reverse primer, 8 μL of 10 mM dNTP, 8 μL Taq DNA Polymerase, 40 μL 10× Taq Buffer, and sterile water to a final volume of 400 μL. The 400 μL reaction was mixed and then separated into 8 50 μL PCR tubes prior to running on the thermocycler. The PCR was run as follows: 95 °C for 180 sec followed by 30 cycles of 95 °C for 45 sec, 57 °C for 30 sec, and 72 °C for 117 sec. Once the cycles were complete, the reactions underwent a final extension for 10 min at 72 °C followed by a final hold at 4 °C until removed from thermocycler. The amplified product was run on a 1% agarose gel, the correctly sized band was cut out of the gel, and the DNA purified using the Epoch Life Science GenCatch Gel Extraction Kit.

The pTH353 vector was subjected to a two-step restriction enzyme digest: first with SpeI, followed by digestion with BamHI and BglII. The following DNA masses were precipitated using Pellet Paint NF Co-Precipitant according to manufacturer’s protocols: 4 μg mutagenized SUP45 insert, 1 μg digested vector backbone, 1 μg reporter plasmid (pCTCON2-FAPB2.3.6L1TAG-LeuOmeRS). A control sample containing only the digested backbone DNA and reporter plasmid (no insert) was also pelleted.

Preparation of electrocompetent *S. cerevisiae* strain RJY100 (MATa AGA1::GAL1-AGA1::URA3 ura3-52 trp1::KanMX leu2-Δ200 his3-Δ200 pep4::HIS3 prbd1.6R can1 GAL) and electroporation protocols were performed as described previously^64,108^. The number of transformants was estimated by plating and colony counting as previously described^64,108^. Library growth and long-term storage was performed as previously described^87^. Naïve library containing pCTCON2-FAPB2.3.6L1TAG-LeuOmeRS (yeast display) reporter induced and labelled prior to flow cytometry characterization as described in the “reporter system validation experiments” section. FCS files were analyzed using FlowJo as previously described^88^. 10 clones from the naïve library were subjected to full plasmid sequencing (Plasmidsaurus).

### Fluorescence Activated Cell Sorting (FACS)

For each round of sorting, 10 million induced cells were aliquoted into 1.5 mL microcentrifuge tubes, diluted to 1 mL of PBSA then centrifuged for 30 sec at 15,871 × g. The supernatants were aspirated and pellets were washed by resuspending in 1 mL PBSA, centrifuging at 15,871 × g for 30 sec, aspirating the supernatant, and repeating for a total of 3 washes. Cells were then labelled with 100 μL of PBSA with a 1:500 dilution of primary antibodies: mouse anti-HA (BioLengend) and chicken anti-c-Myc (Exalpha) and incubated for 30 min on a rotary wheel at room temperature. Following incubation, cells were diluted to 1 mL of PBSA then centrifuged for 30 sec at 15,871 × g. The supernatants were aspirated and pellets were washed by resuspending in 1 mL PBSA, centrifuging at 15,871 × g for 30 sec, aspirating the supernatant, and repeating for a total of 2 washes. Cells were then labelled with 100 μL of PBSA with a 1:500 dilution of secondary antibodies: goat anti-mouse Alexa Fluor 488 (Invitrogen), and goat anti-chicken Alexa Fluor 647 (Invitrogen) and incubated for 15 min on ice. Following incubation, cells were diluted to 1 mL of PBSA then centrifuged for 30 sec at 15,871 × g. The supernatants were aspirated and pellets were washed once by resuspending in 1 mL PBSA, centrifuging at 15,871 × g for 30 sec, aspirating the supernatant. Samples were sorted using S3e Cell Sorter (BioRad). The number of cells used in each round of sorting was equal to or larger than 10× the library size. Gating strategies can be found in Fig. S4A. Sorted samples were recovered in 5 mL of SD-SCAA-W-L-U supplemented with penicillin-streptomycin for ∼3 days at 30 °C with shaking.

### Sorted Clone Characterizations

Recovered sorts were subjected to yeast miniprep to isolate DNA which was then transformed into *E. coli*. When grown on solid media in the presence of 50 μg/mL ampicillin, individual clones were able to be isolated containing either a SUP45 mutant plasmid or a reporter/OTS plasmid. DNA from 40 colonies was isolated and subjected to a restriction analysis with BamHI and XbaI and run on a 1% agarose gel. 10 samples with bands running at a size corresponding to eRF1 plasmids were subjected to full plasmid sequencing (Plasmidsaurus) and transformed into RJY100 alongside a yeast display reporter/OTS (pCTCON2-FAPB2.3.6L1TAG-LeuOmeRS) prior to flow cytometry characterization (Fig. S5).

### Single Mutant Construction and Characterization

Single mutants were generated via site-directed mutagenesis of Wild Type SUP45 open reading frame (pTH353-SUP45WT Addgene #29371). The pTH353 vector underwent a two-step restriction enzyme digest first with BamHI then with BglII. The SUP45 insert was then amplified with mutagenic oligos and Q5 polymerase. The following primers were used for single mutant cloning: SUP45_Cloning_Fwd, SUP45_Cloning_Rev, P2_variant, and P3_variant. P2_variant and P3_variant full names and sequences are described in Table S2. The variants created include: G28A, Q46E, I59V, S61P, Y122N, F169V, Q182E, F239I, Y255N, D310Y, and G381A. PCRs were run as follows: 30s initialization at 98 °C followed by 35 cycles of 10 s at 98 °C, 30 s at 61 °C, and 1 min at °C. Reactions were then subjected to a final extension of 4 min at 72 °C with a final hold at 4 °C.

DNA fragments were analyzed via gel electrophoresis, extracted, purified, and assembled via Gibson Assembly prior to being transformed into *E. coli* strain DH5αZ1. DNA extracted from individual colonies was sequenced verified via Sanger sequencing (Quintara Biosciences) using primers SUP45_Seq_Fwd and SUP45_Seq_Rev to confirm the desired mutations, then transformed into *S. cerevisiae* strain BY4741 (MATa his3Δ1 leu2Δ0 met15Δ0 ura3Δ0) alongside the BXGaltTAG intracellular reporter/OTS.

Sequence-verified single mutants were propagated and induced as described above, then washed in PBSA prior to being subjected to flow cytometry using an Attune NxT Flow Cytometer (Life Science Technologies). Flow cytometry FCS files were analyzed using FlowJo. RRE and MMF for S30P, Q46E, E52D, Y122N, and Y255 were calculated along with sorted clones BR3T.4 and CR3T.2 (Fig. 3A-D)^26^. The median fluorescence intensity of C-terminal detection in N-terminal positive cells of S30P, Q46E, Y122N, E52D, Y255, BR3T.4 and CR3T.2 was analyzed in Graphpad (Prism) and subjected to a one-way ANOVA statistical analysis (Fig. 3E-F).

### Deep Mutational Scanning Analysis

Prior to deep sequencing, naïve (inp) and two sorted (sel) libraries: Track B round 3 sort (BR3T) and Track C round 3 sort (CR3T) were amplified via Q5 PCR to generate 4 ∼300 bp fragments with Illumina adapter sequences. PCR was performed by combining 1 μL template DNA, 2.5 μL 10 μM forward primer (Frag1_Fwd, Frag2_Fwd, Frag3_Fwd, or Frag4_Fwd) and 2.5 μL 10 μM reverse primer (Frag1_Rev, Frag2_Rev, Frag3_Rev, or Frag4_Rev) per respective fragment, 25 μL 2× Q5 Master Mix, and 19 μL water for a 50 μL total reaction volume. Sequences of all primers can be found in Table S2. Reactions were performed on a thermocycler at 98 °C for 30 sec followed by 30 cycles of 98 °C for 10 s, 58 °C for 30 sec, 72 °C for 15 sec. Once cycles were complete, reactions underwent a 2 min final extension at 72 °C followed by a final hold at 4 °C until removed. DNA fragments were then analyzed on a 2.5% agarose gel run at 130 V for 35 min. DNA fragments containing adapter sequences were extracted and purified using the GenCatch gel extraction kit then sent to Quintara Biosciences for deep sequencing via AmpExpress Illumina sequencing.

Analysis of raw sequencing files was performed using Geneious Prime 2023.1.2. Reads were paired with a 300 bp insert size. Paired end reads were then trimmed using BBDuk with a Kmer Length of 27, trimming low quality reads with a minimum quality score of 30. Short reads (<150bp) were discarded. Trimmed files were mapped to reference *S. cerevisiae* SUP45, Gene ID: 852440. Once assembled to reference, single nucleotide variations were called and the following data was exported for each SNV: CDS Codon Number, Variant Raw Frequency (C_v_), Coverage (C_wt_), and Amino Acid Change. The exported csv files were used to calculate enrichment scores using Equation 1^111,112^ ^113^. A nominal value of 0.5 was added to raw counts to avoid errors caused by dividing by zero in instances where specific variants did not occur^112,113^. Similarly, an average coverage value (per fragment) was used in instances where there was no variant data.

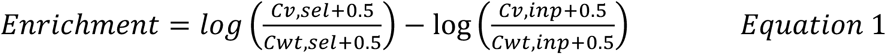

### Evaluation of ncAA incorporation efficiency in YTH41

*S. cerevisiae* strain YTH41 strain (derivative of CML476; ura3-52 leu2-Δ1 his3-Δ200 GAL2 LEU2::CVp(TetR’-SSN6) trp1::tTA sup45::kanMX4-tetO7-SUP45) was obtained from the von der Haar Lab (University of Kent, England, United Kingdom)^89^. pRS413-SUP45-WT, pRS413-SUP45-N58A, and pRS413-SUP45-BR3T.4 were each transformed alongside pRS416-BXGAltTAG-TyrOmeRS in YTH41. Transformation, propagation, and induction was performed as described above with SD-SCAA (−HIS −URA) for growth and SG-SCAA (−HIS −URA) for induction. At the induction step, samples were supplemented with either 0 or 10 µg/mL of doxycycline, supplemented with a final concentration of 1 mM OmeY, and incubated overnight at 20 °C with shaking at 300 rpm. After the overnight incubation, samples were then washed in PBSA and then subjected to flow cytometry using an Attune NxT Flow Cytometer (Life Science Technologies). Flow cytometry FCS files were analyzed using FlowJo (Fig. S12). The median fluorescence intensity of C-terminal detection in N-terminal positive cells of each sample was analyzed in Graphpad Prism and subjected to a one-way ANOVA statistical analysis (Fig. 4B).

### Evaluation of ncAA Incorporation Efficiency in Knockout Strains

To evaluate eRF1 expression across a subset of knockout strains, transformations specified in Table 3 were performed and plated on solid SD-SCAA (−LEU –URA). Four colonies from each plated transformation were inoculated in 5 mL liquid SD-SCAA (−LEU –URA) with 50 µL of 100X penicillin-streptomycin. Cultures were grown for 3 days at 30 °C with shaking at 300 rpm. Once saturated, liquid cultures were diluted to an OD600 of 1 in fresh media and grown 4-8 hours to mid log phase (OD600 2-5) with shaking at 30 °C. Cells were once again diluted to an OD600 of 1, pelleted at 2400 × *g* for 5 min, resuspended in induction media (SG-SCAA) supplemented with 1 mM final concentration of L-isomer OmeY and grown for 16 hours at 20 °C with shaking. After the overnight incubation, ODs were measured and volumes corresponding to 2 million cells per sample were aliquoted into a 96 well plate. Samples were diluted once to a final volume of 200 µL PBSA, centrifuged at 2400 × *g* for 5 min and decanted. Samples were then washed 3 more times in 200 µL PBSA, centrifuging at 2400 × *g* for 2 min and decanting between each wash. Samples were then subjected to flow cytometry and analyzed as previously described^88^.

### Protein Expression and Purification for MALDI-MS

Yeast strain RJY100 was transformed with protein secretion vector pCHA-Fc-Sup-Donkey1.1-H54TAG-TyrAcFRS, as well as an eRF1 plasmid (either WT, Q46E, or BR3T.4) as described above, and plated on SD-SCAA (−TRP −LEU –URA). One colony from each plate was inoculated in 5 mL liquid minimal media supplemented with penicillin-streptomycin and grown shaking at 30 °C for 3 days. Once saturated, the 5 mL culture was diluted to ∼OD 1 in 50 mL YPD then grown overnight shaking at 30 °C until the OD reaches 6-8. The entire 50 mL culture was then centrifuged at 2400 × *g* for 10 min and decanted. Cell pellet was then resuspended in 100 mL of YPG induction media supplemented with penicillin-streptomycin and 1 mM final concentration of L-isomer OmeY and induced for 4 days at 20 °C with shaking. The 100 mL culture was then centrifuged at 3000 × *g* for 10 min. Once pelleted, the culture was decanted into a filter cup for vacuum filtration and collection of the protein-containing supernatant. Proteins were then purified prior to undergoing a tryptic digest and ziptip cleanup as previously described^88^. Purified peptide fragments were flash frozen and sent on dry ice to the Koch Institute Biopolymers and Proteomics Core for MALDI mass spectrometry. The expected trypsinized mass for the wildtype Donkey1.1 peptide containing the H54 position was 2234.1 Da (amino acid sequence GLEWVSAISGSGGSTYYADSVK). The expected mass for the trypsinized peptide containing OmeY encoded at the H54TAG position was 2324.2 Da (amino acid sequence GLEWVSAISG(OmeY)GGSTYYADSVK) with an observed mass of 2324.8 when cultured and induced in the presence of WT eRF1, or 2325.0 in the presence of eRF1 variants Q46E and BR3T.4. The molecular weight of OmeY is 195.2 Da and the molecular weight of serine (the WT residue at position H54) is 105.1, thus the difference in mass between the wildtype and the OmeY incorporated peptide is 90.1 Da which is reflected in the expected mass for the trypsinized peptide.

### Click Chemistry on the yeast surface

RJY100 was transformed with pCTCON2-FAPB2.36L1TAG-LeuOmeRS alongside either plasmid-based WT or mutant eRF1 variants (S30P, Q46E, Y122N, Y255N, BR3T.4, and CR3T.2). Cells were transformed and propagated in biological triplicate as described above prior to inducing overnight in SG-SCAA (−TRP −LEU –URA) supplemented with penicillin-streptomycin and either 1 mM *O*-methyl-L-tyrosine or 1 mM *p*-propargyloxy-L-phenylalanine (OPG) at 20 °C with shaking. From the induced cultures, 2 million cells were aliquoted from each sample into a 96 well plate. Cells were then washed 3 times in 200 µL room temperature PBSA prior to resuspension in 220 µL 1X PBS pH 7.4. The following reagents were then added to each well for biotin click chemistry: 1.25 µL 20mM biotin-alkyne, 3.8 µL CuSO_4_/THPTA, 12.5 µL aminoguanidine, 12.5 µL sodium ascorbate. Cells were incubated with all listed reagents for a minimum of 15 min and a maximum of ∼25 minutes. For the purposes of this experiment, the goal was to qualitatively assess whether click chemistry happened or not in response to OPG or OmeY so precise timing for extent of reaction calculations was not performed. After the incubation period, cells were washed twice in 4 °C PBSA then labelled for flow cytometry. Full length readthrough was detected via c-Myc labelled with 1:500 dilutions of chicken anti-cMyc and goat anti-chicken AF 647. Biotin detection (resulting from successful click chemistry of biotin onto azide-containing ncAA) was labelled for via a 1:500 dilution of PE-anti biotin. Flow cytometry analysis enabled simultaneous evaluation of readthrough efficiency and click-dependent labelling, allowing qualitative discrimination between samples incorporating the clickable ncAA (OPG) and the non-clickable control (OmeY).

### Phylogenetic analysis of homologous eRF1

A standard global alignment was performed using Geneious Prime® 2025.0.3 on homologous peptide chain release factors in the following organisms, each listed alongside their UniProtKB ascension number: *S. cerevisiae* P12385, *S. pombe* P79063, *P. anserina* O59948, *D. melanogaster* Q9VPH7, *X. laevis* P35615, *H. sapiens* P62495, *O. cuniculus* P62497, *M. musculus* Q8BWY3, *B. taurus* Q0VCX5, *P. abelii* Q5R4C7, *T. brucei* Q9NAX8, *M. barkeri* A0A0E3R3Y4, *C. elegans* O44568, *M. capricolum* P71496, *S. citri* Q14QE1, *E. crassus* A0AAD1XDL0, *E. coli* prfA A0AAI9GXY5, *E. coli* prfB P07012, *P. tetraurelia* Q965E7, *L. striatus* Q5CD84, *S. lemnae* Q9BMM0, *T. thermophilia* Q9U8U5, *S. strix* A0A5J4USS5. The consensus sequence was determined by majority, excluding gaps (Table S1). A standard global alignment tree was additionally created in Geneious Prime® 2025.0.3 with the same organisms (Fig S16).

## Acknowledgements

We are grateful to J.T. Stieglitz, A. Rezhdo, and R.L. Hershman for advice and discussions, T. von der Haar from the University of Kent for providing yeast strain YTH41, and R. Said and V. Trivedi for advice on deep sequencing analysis. This work was supported by a grant from the National Institute of General Medical Sciences of the National Institutes of Health (R35GM133471 to J.A.V.), and the Bright Futures Assistant Professorship at Tufts University (to J.A.V.). The content is solely the responsibility of the authors and does not necessarily represent the official views of the National Institutes of Health or Tufts University.

## Supplementary Information

Supplementary Figures 1-16, Table S1, and Table S2.

## Supplementary Data 1

Excel file of deep sequencing analysis.

