## Supplementary Information for "High throughput screening of eukaryotic release factor 1 variants to enhance noncanonical amino acid incorporation"

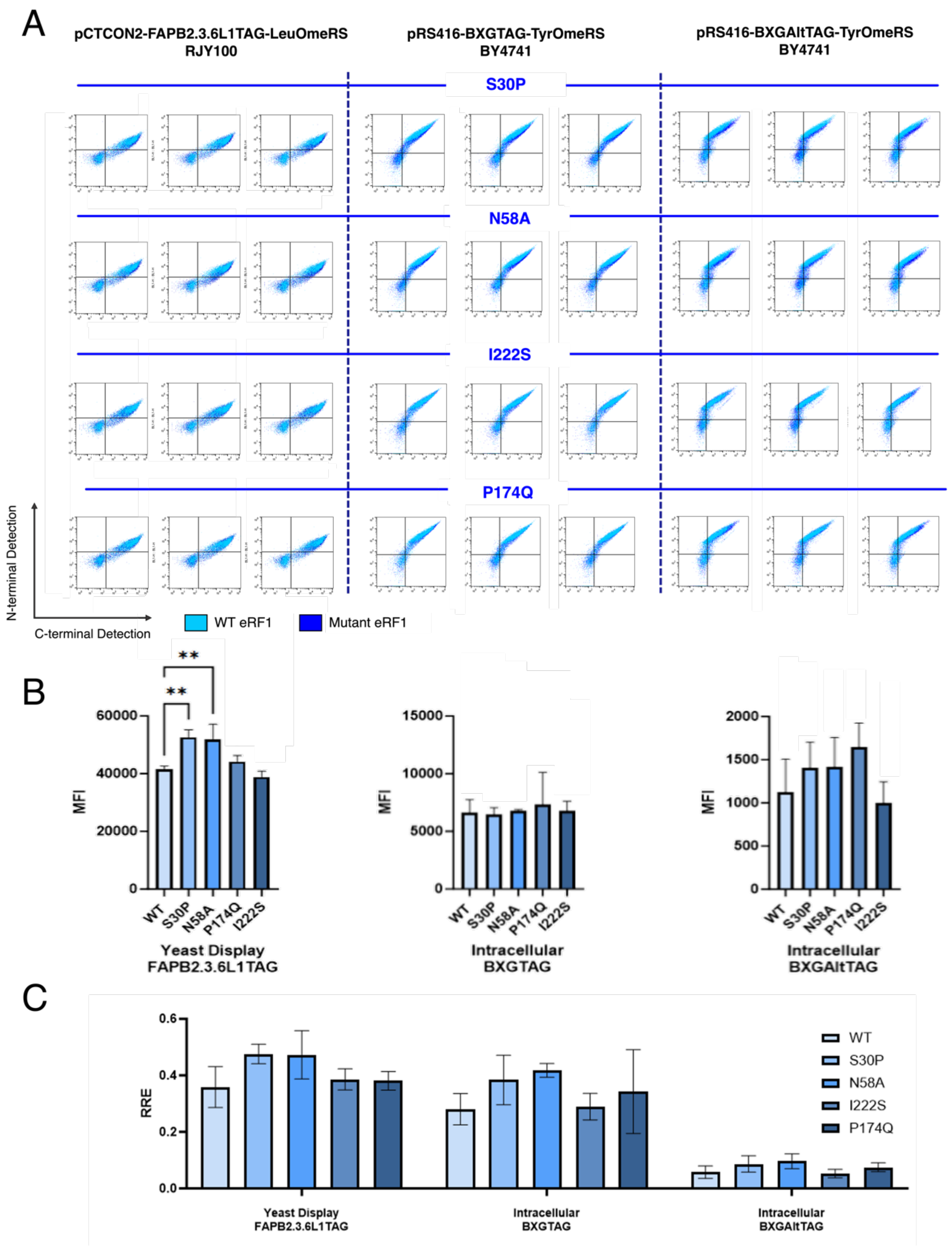


**Figure S1. Single mutant characterizations across reporter systems.** Four *S. cerevisiae* eRF1 variants previously reported to have improved TAG codon readthrough and a WT eRF1 control were expressed on a plasmid-based second copy of eRF1 alongside one of three fluorescent reporter systems, all of which contain an orthogonal translation system for OmeY incorporation. Fluorescent detection indicating levels of full-length protein was analyzed via flow cytometry. A)

The left-most panel shows flow plots from a yeast display reporter (pCTCON2-FAPB2.3.6L1TAG-LeuOmeRS) co-transformed with each of the variants (S30P, N58A, I222S, P147Q; yeast numbering) in *S. cerevisiae* strain RJY100. The middle and right-most panels utilize intracellular reporters with a TAG codon inserted at two different positions within the linker between Blue Fluorescent Protein (BFP) and Green Fluorescent Protein (GFP) transformed into *S. cerevisiae* strain BY4741. Each sample was run in biological triplicate, with three dot plots per sample shown side by side. The WT eRF1 flow plots (light blue) are overlaid on the mutant eRF1 plots (dark blue) for each reporter. B) Median fluorescence intensity (MFI) of full-length reporter proteins in N-terminal positive cells. One-way ANOVA statistical analysis shown with the following p-value assignments: ** $p \leq 0.01$, ns $p > 0.05$. C) Relative readthrough efficiencies (RRE), calculated as previously described[1], of WT and mutant (S30P, N58A, I222S, P174Q) eRF1 plasmids using a yeast display and 2 different intracellular reporter systems.

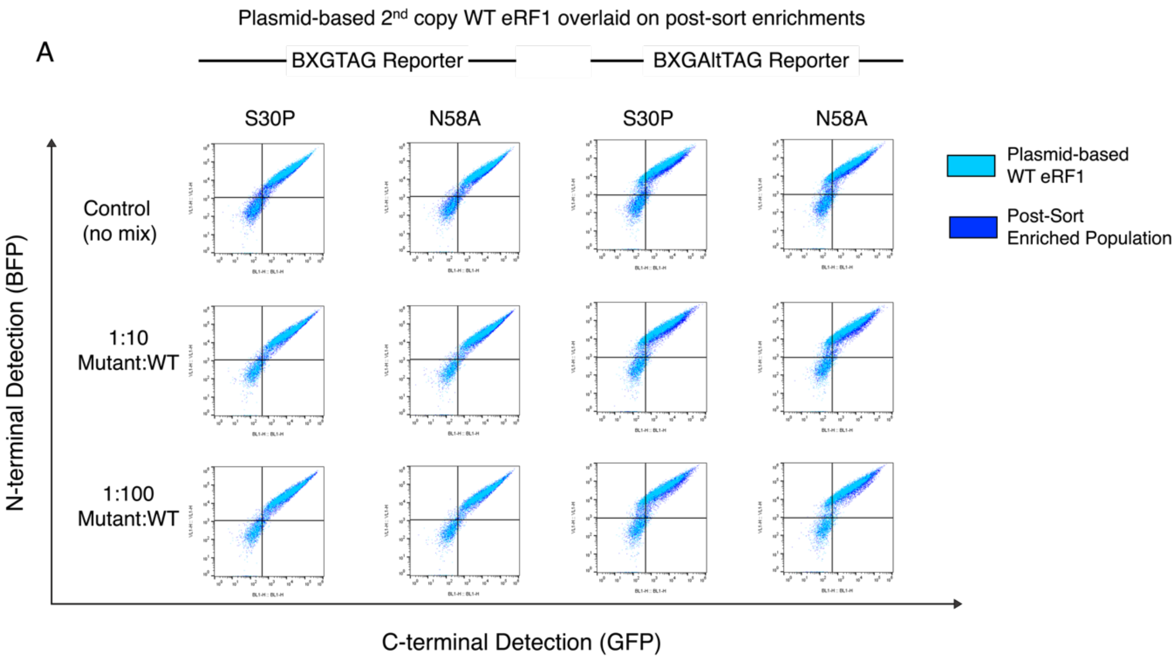


B

| | **S30P BXGTAG** | **S30P BXGAltTAG** | **N58A BXGTAG** | **N58A BXGAltTAG** |
|---|---|---|---|---|
| Starting Ratio | 1/100 | 1/100 | 1/100 | 1/100 |
| Post-Sort Colonies with specified mutation | 1/10 | 2/10 | 0/10 | 2/10 |
| Estimated Fold Enrichment | 10X | 20X | -- | 20X |

**Figure S2: Model enrichment analysis of plasmid-based 2nd copies of eRF1.** A) Columns 1 and 2 show flow cytometry data using intracellular BXG reporter, columns 3 and 4 using intracellular BXGaltTAG reporter. Top Row: Flow cytometry characterizations of WT eRF1 (light blue) overlaid on variant (S30P or N58A) eRF1 (dark blue). Middle Row: Post-sort flow cytometry characterizations of WT eRF1 (light blue) overlaid on the sorted populations recovered from 1:10 mixtures of variant (S30P or N58A) to WT eRF1 (dark blue). Bottom row: Post-sort flow cytometry characterizations of WT eRF1 (light blue) overlaid on the sorted populations recovered from 1:100 mixtures of variant (S30P or N58A) to WT eRF1 (dark blue). X-axis represents C-terminal detection, Y-axis represents N-terminal detection. B) Table of FACS model enrichment data with mutants S30P and N58A using two intracellular reporters (BXGTAG and BXGAltTAG). A mix of mutant:WT at a 1:100 ratio was sorted using FACS for enhanced stop codon readthrough. 10 clones from recovered populations were sequenced to calculate the estimated fold enrichment.

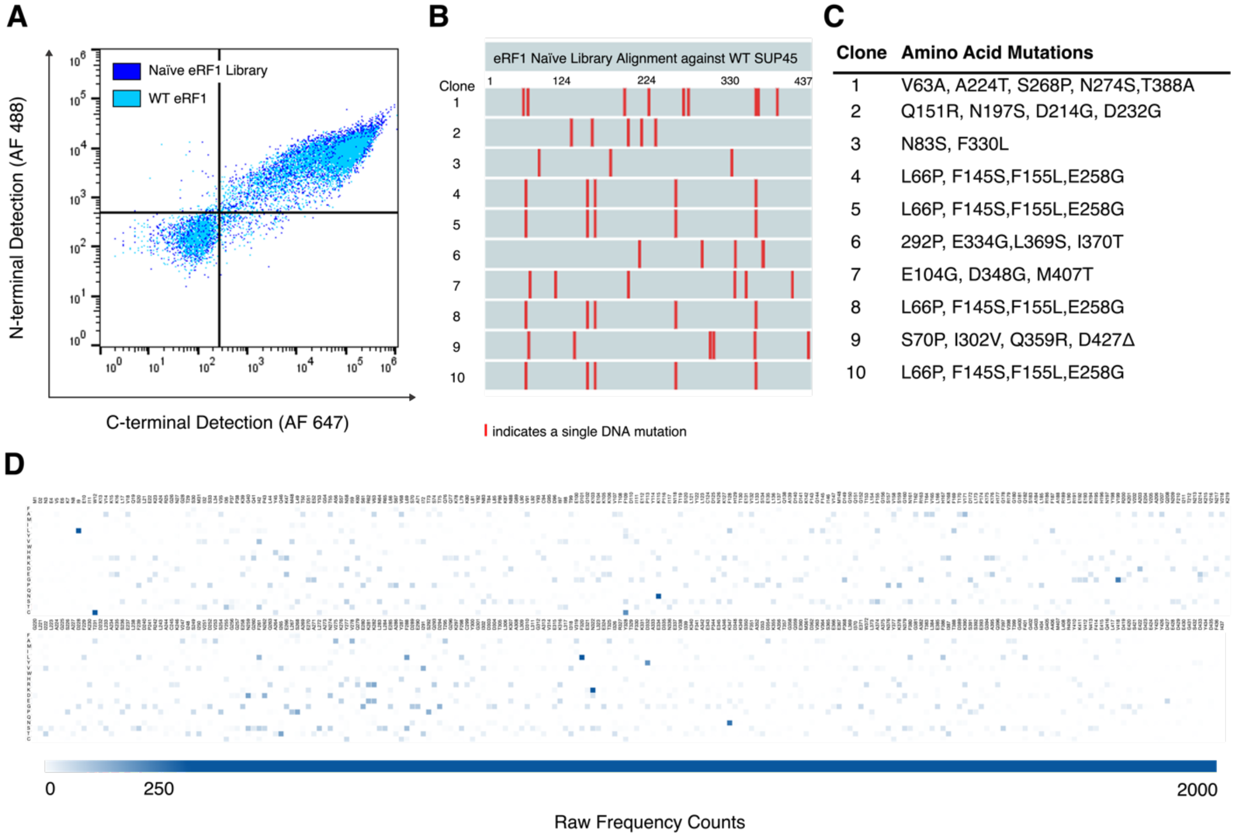


**Figure S3. Naïve Library Characterizations** A) Flow cytometry analysis of naïve eRF1 library (dark blue) in comparison to WT eRF1 (light blue) co-expressed with a yeast display reporter/OTS (pCTCON2-FAPB.2.3.6L1TAG-LeuOmeRS). On the two-dimensional dot plots, the N-terminal epitope detection levels are plotted on the Y-axis and the C-terminal epitope detection levels are plotted on the X-axis. B) Sequence alignment of 10 clones isolated from the naïve library against a WT *SUP45* open reading frame (ORF). Red lines indicate single nucleotide mutations. C) Amino acid mutations present in the 10 isolated clones. D) Raw frequency counts of amino acid mutations present in naïve library across full *SUP45* ORF obtained from deep sequencing analysis. Frequency values plotted on a gradient heatmap from 0 (white) to 2000 (dark blue). Average coverage per gene fragment as follows: Residues 1-109: 12810, Residues 110-219: 13402, Residues 220-329: 16475, Residues 330-437: 8737.

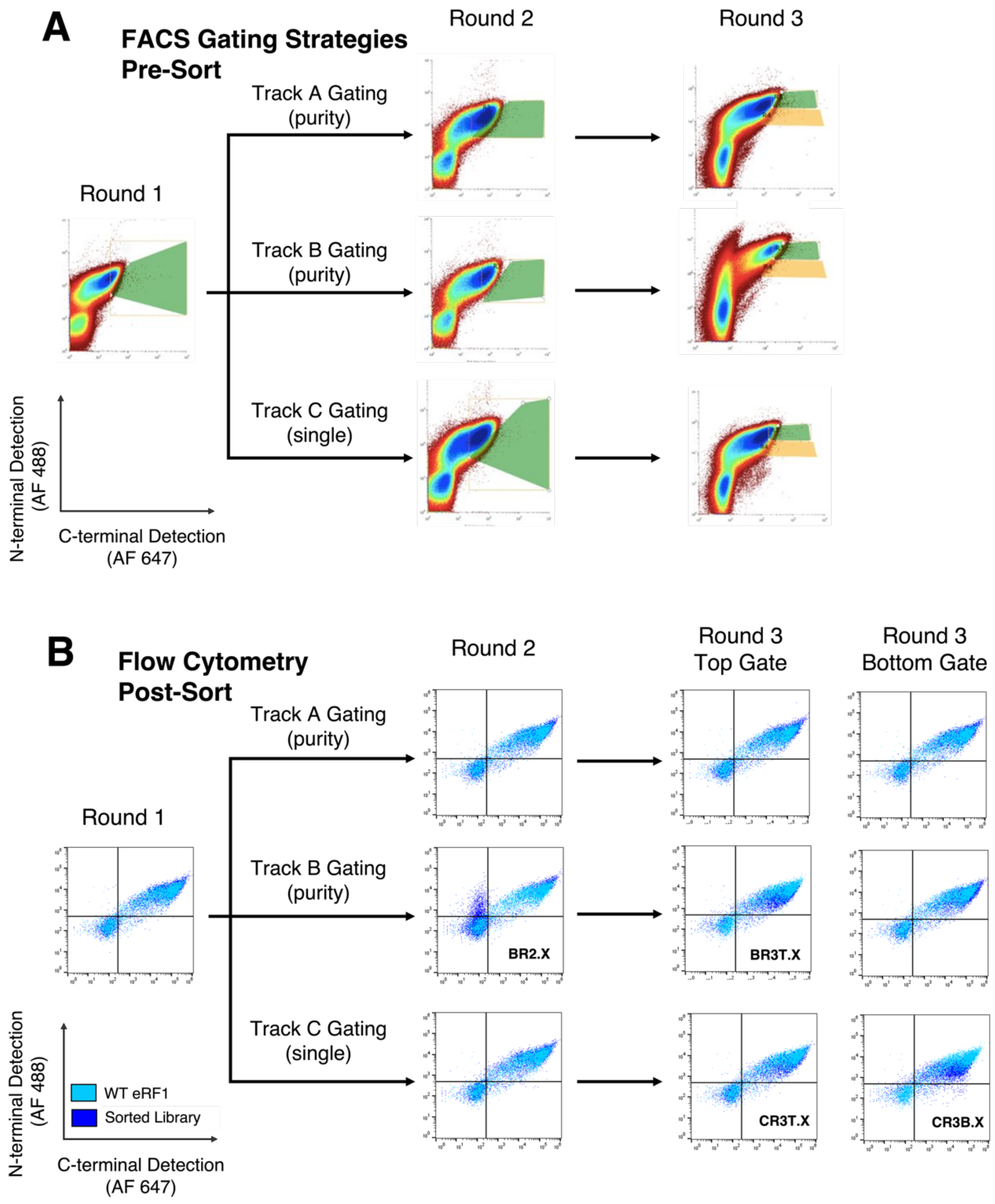


**Figure S4. Screen for enhanced ncAA incorporation.** A) Gating strategies for each round of FACS. Green or yellow shaded polygons indicate sort gates. After round 1, sorting was performed using three gating strategies labelled Track A (purity sort), Track B (purity sort), or Track C (single sort). X-axis represents C-terminal detection (AF 647), Y-axis represents N-terminal detection (AF 488). Labels indicate which track the sorted clones were derived from (BR2.X, BR3T.X, CR3T.X, CR3B.X) with X representing the individual clone number. B) Flow cytometry analysis of WT eRF1 (light blue) overlaid on library population (dark blue) after each round of sorting according to the gating strategies in panel A. X-axis indicates C-terminal detection (AF 647), Y-axis represents N-terminal detection (AF488). Round 3 top gate references the green gate in round 3 of Panel A, bottom gate references yellow gate in round 3 of Panel A.

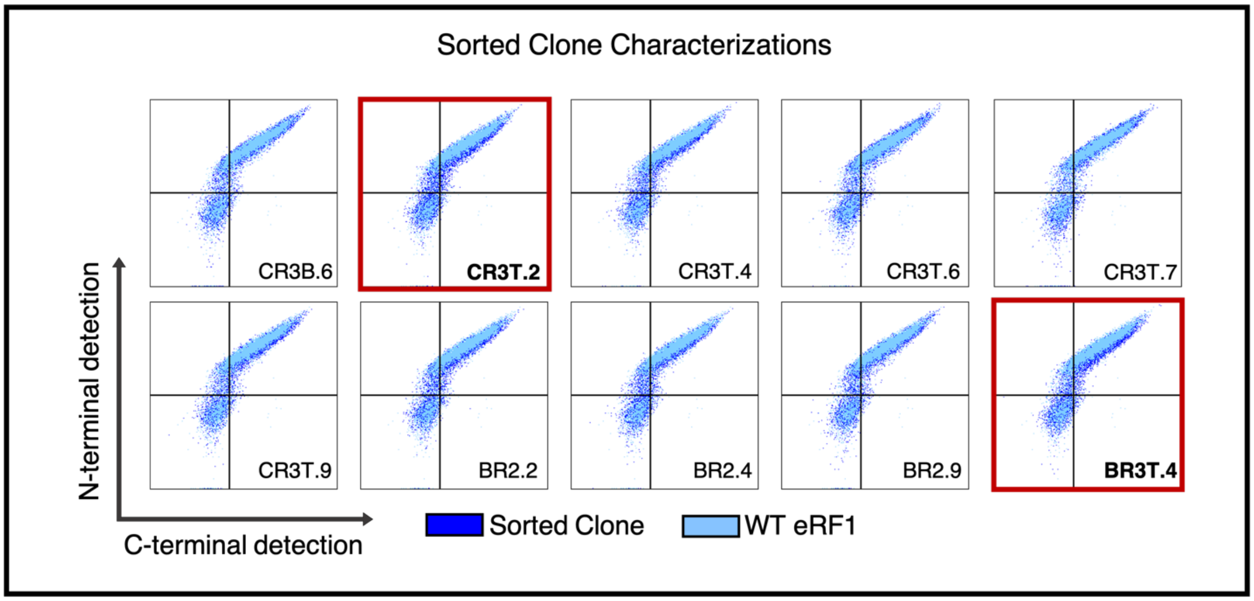


**Figure S5. Sorted clone phenotypes.** Flow cytometry characterizations of single clones isolated from FACS for enhanced ncAA incorporation. See Fig. S4 for sorting strategies resulting in each clone. WT eRF1 data (light blue) overlaid on specified sorted clone (dark blue). Y-axis measures N-terminal detection, X-axis measures C-terminal detection indicating full length TAG codon readthrough. Dot plots boxed in red indicate clones that were further characterized in subsequent experiments.

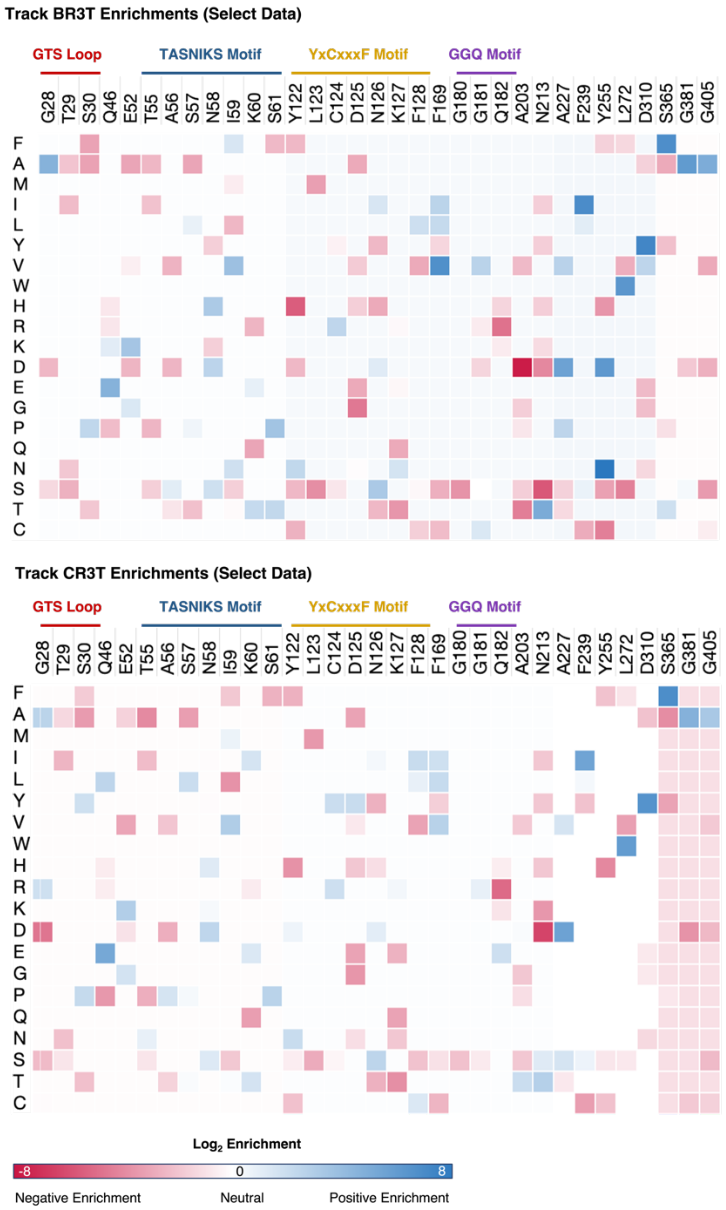


**Figure S6. Log$_2$ enrichment values per mutation in select positions of the SUP45 gene for the sorting tracks B and C after 3 rounds of sorting.** Heatmaps graphed using Morpheus[2]. Enrichment values range from Log$_2$ of enrichment = -8 to 8, with negative enrichments shown in red, and positive enrichments in blue. Single letter amino acid codes of mutation are labelled on the y-axis, and WT residues are labelled on the x-axis. The WT residue at each position is excluded from enrichment analysis (see Materials and Methods for details); the value plotted for the WT residue follows Equation 1 where $C_v = 0$ and $C_{wt}$ = average coverage of the fragment.

$$Enrichment = log\left(\frac{Cv,sel+0.5}{Cwt,sel+0.5}\right) - \log\left(\frac{Cv,inp+0.5}{Cwt,inp+0.5}\right) \qquad Equation\ 1$$

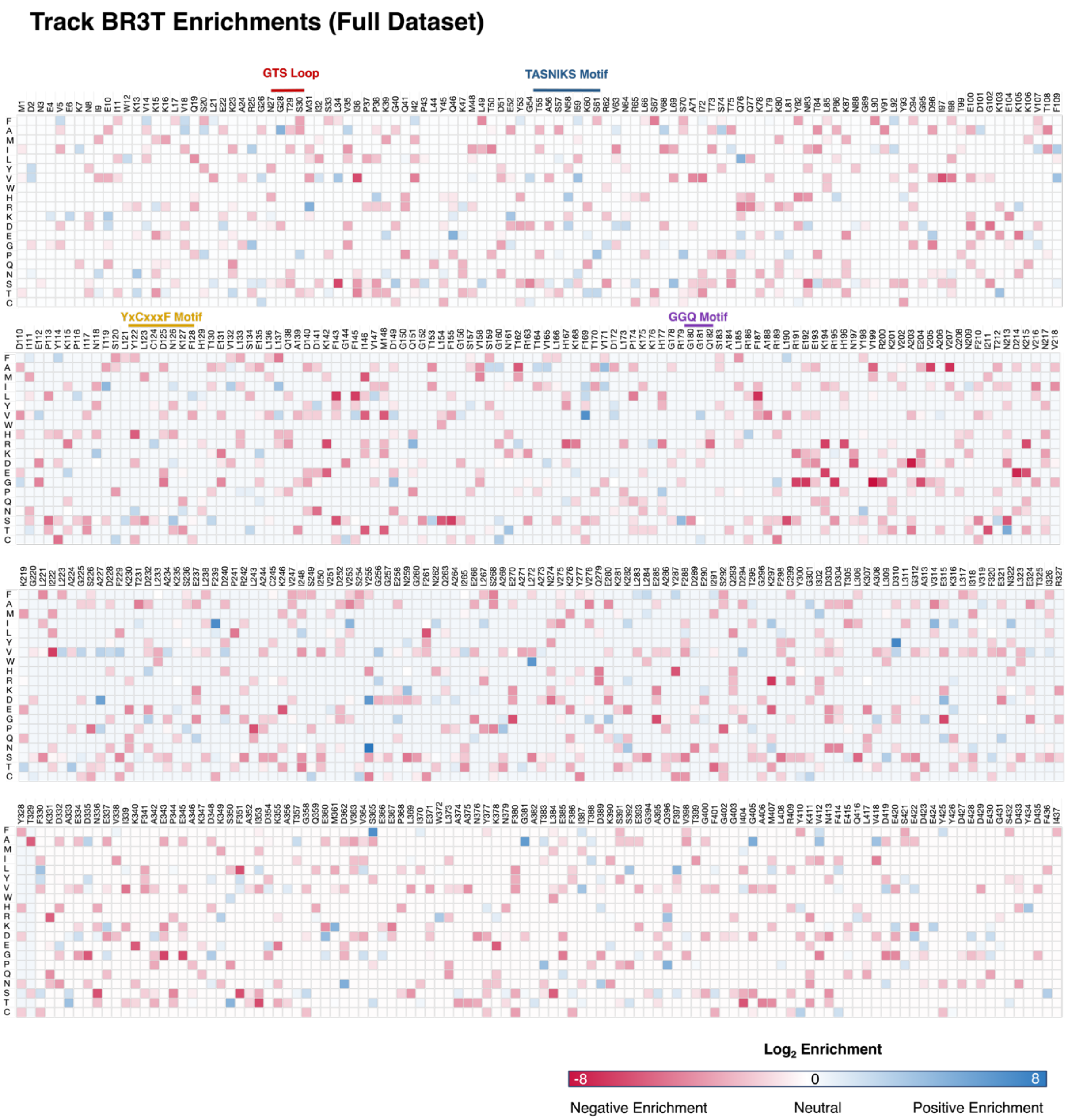


**Figure S7. Log₂ enrichment values per mutation across entire SUP45 gene for sorting track B after 3 rounds of sorting.** Heatmaps graphed using Morpheus. Enrichment values range from Log2 of enrichment = -8 to 8, with negative enrichments shown in red, and positive enrichments in blue. Single letter amino acid codes of mutations on the y-axis, and WT residues labelled on the x-axis. The SUP45 gene was sequenced in four segments to accommodate its large size, resulting in varied coverage values that depend on the sequencing reads obtained in each Illumina sequencing run. In some cases, this resulted in varying average coverage values (see Figure S8).

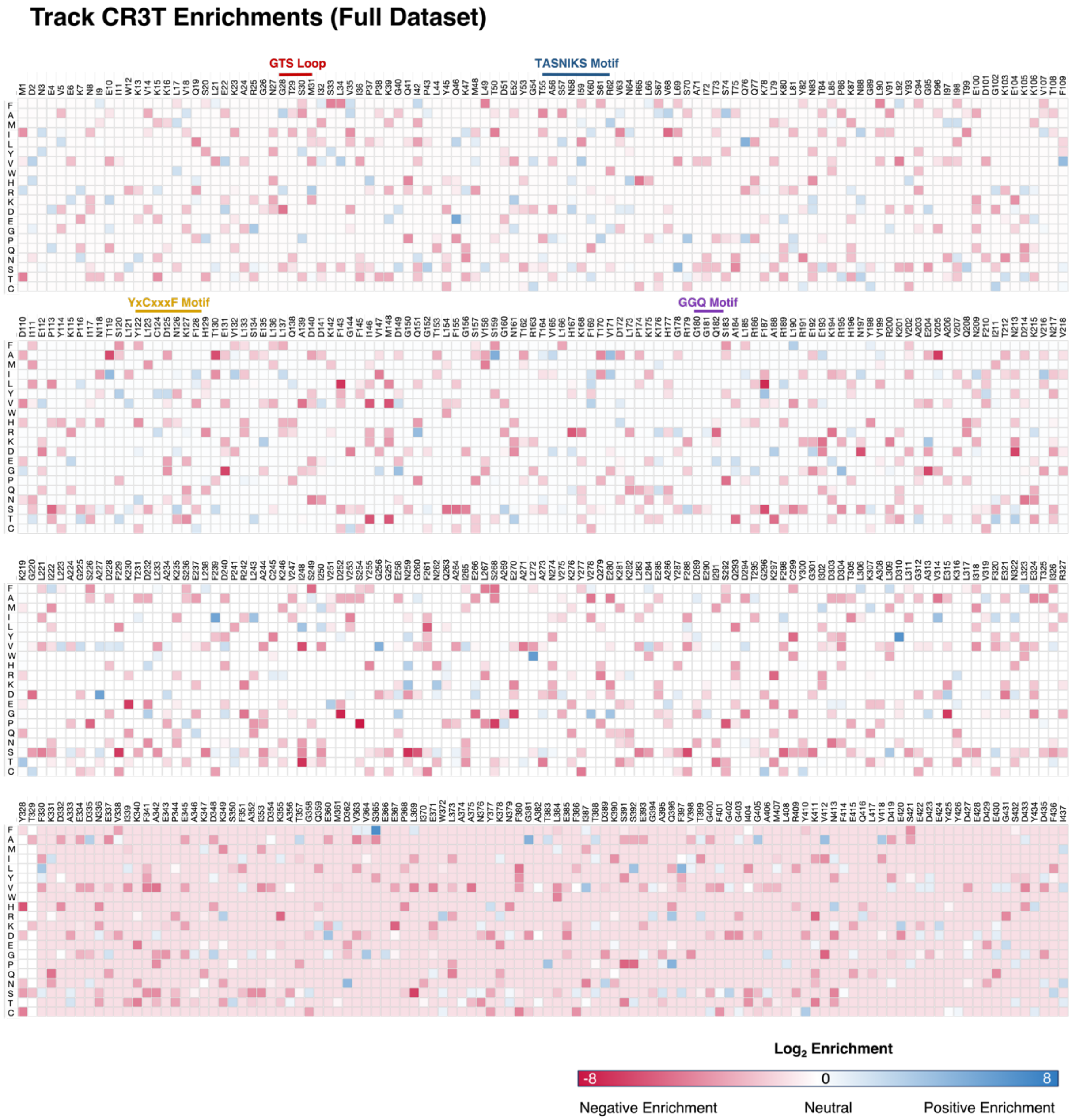


**Figure S8. $Log_2$ enrichment values per mutation across entire SUP45 gene for sorting track C after 3 rounds of sorting.** Heatmaps graphed using Morpheus. Enrichment values range from $Log_2$ of enrichment = -8 to 8, with negative enrichments shown in red, and positive enrichments in blue. Single letter amino acid codes of mutation are labelled on the y-axis, and WT residues are labelled on the x-axis. Raw enrichment values are normalized to the coverage at each position. As a result, mutants in the last fragment of track CR3T appear to be largely de-enriched due to higher average coverage in this fragment compared to the other fragments. The data for the last fragment is plotted with the same gradient as data from the other 3 domains (these domains all have similar coverage values to one another). The SUP45 gene was sequenced in four fragments to accommodate its large size, resulting in varied coverage values that depend on the sequencing reads obtained in each Illumina sequencing run.

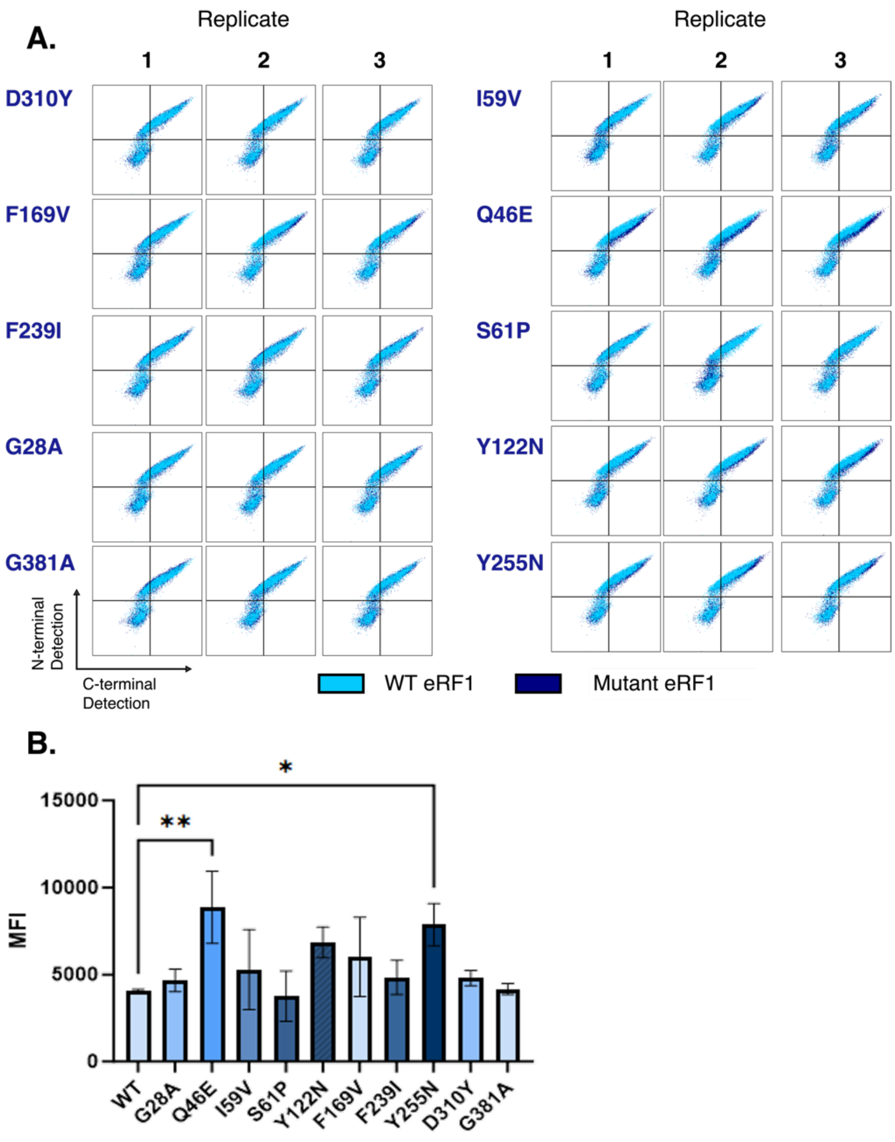


**Figure S9. Single mutant characterizations.** Flow cytometry analysis of single mutants generated and transformed alongside the intracellular BXGAltTAG reporter run in biological triplicate. A) Dot plots of WT eRF1 (light blue) overlaid on variant eRF1 (dark blue). X-axis represents C-terminal detection, Y-axis represents N-terminal detection. B) Median Fluorescence Intensity (MFI) values of GFP (C-terminal) detection in BFP (N-terminal) positive cells. One-way ANOVA statistical analysis shown with the following p-value assignments: ** $p < 0.01$, * $p < 0.05$. All clones were analyzed in biological triplicate.

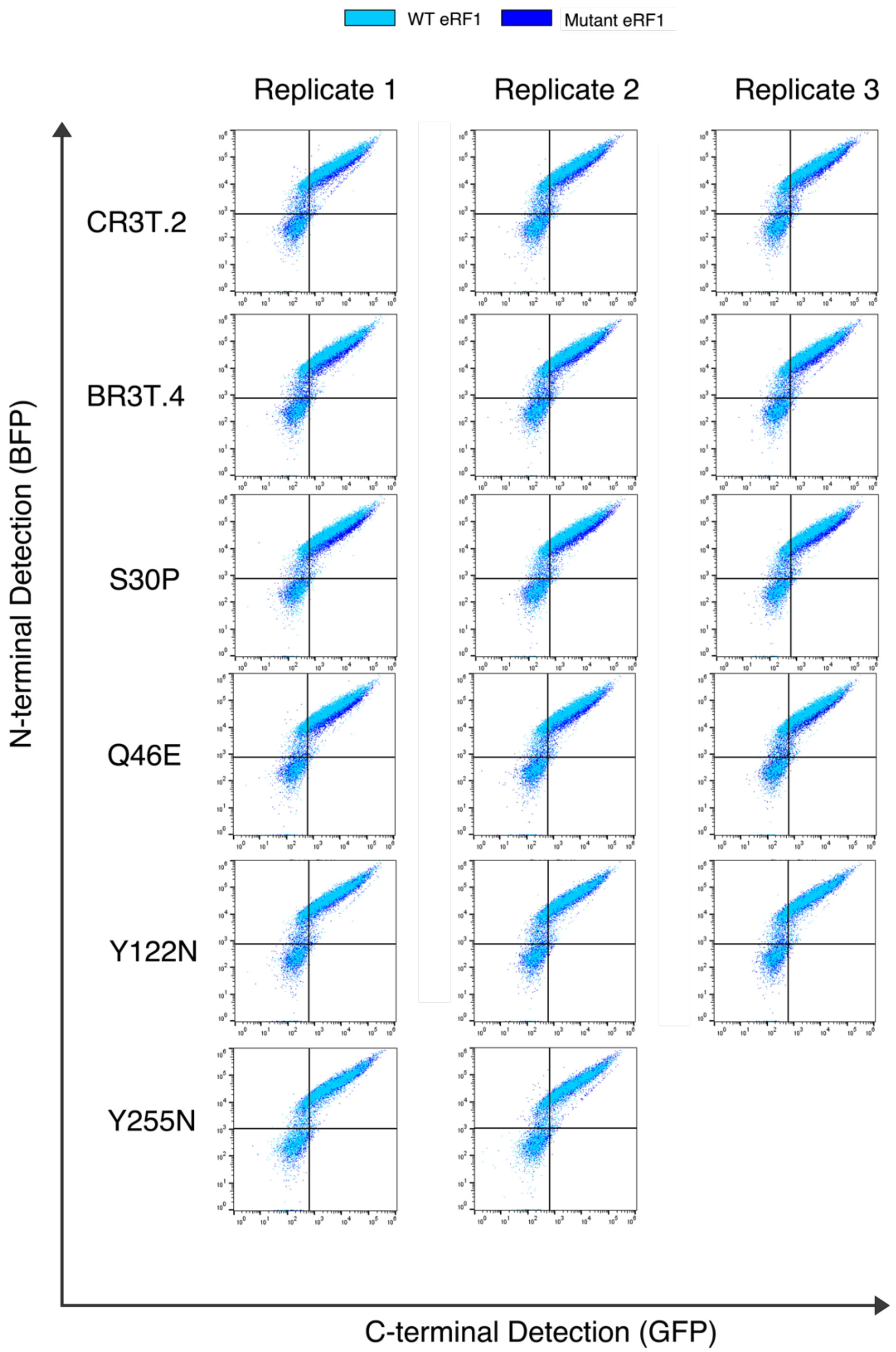


**Figure S10. Characterizations of eRF1 variants exhibiting enhanced ncAA incorporation.** Flow cytometry analysis of WT eRF1 (light blue) overlaid on 2 sorted clones (CR3T.2, BR3T.4) and 4 clones containing point mutations (S30P, Q46E, Y122N, Y255N) (dark blue). X-axis represents C-terminal detection, Y-axis represent N-terminal detection. Analysis of WT, CR3T.2, BR3T.4, S30P, Q46E, and Y122N clones performed in biological triplicate. Y255N analysis performed in biological duplicate due to failed induction of third replicate.

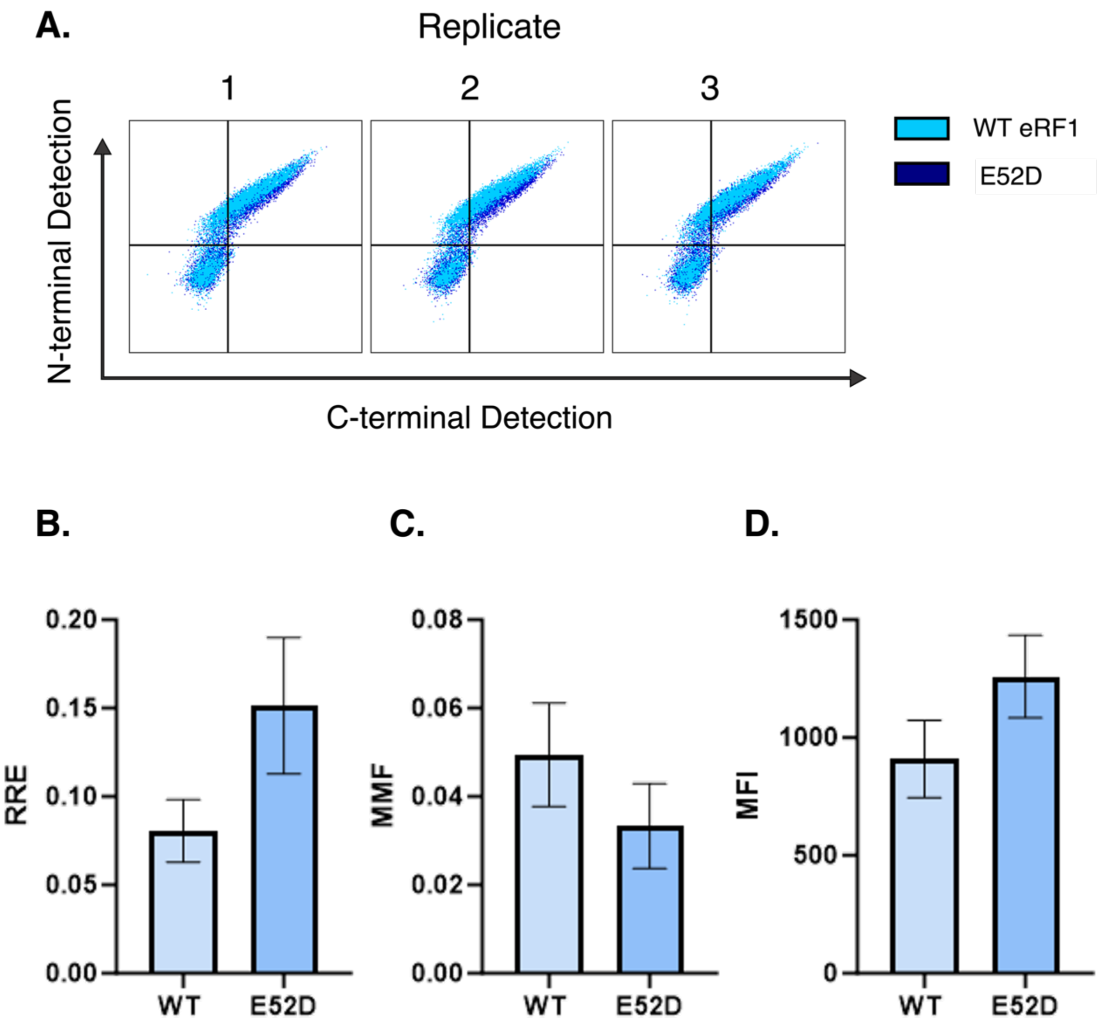


**Figure S11. Characterizations of E52D mutation.** Flow cytometry analysis of E52D and WT 2nd copy eRF1 transformed alongside the intracellular BXGaltTAG reporter run in biological triplicate. A) Dot plots of WT eRF1 (light blue) overlaid on E52D (dark blue). X-axis represents C-terminal detection, y-axis represents N-terminal detection. B) Relative Readthrough Efficiency (RRE) of E52D and WT eRF1. C) Maximum misincorporation frequency (MMF) of E52D and WT eRF1. D) Median Fluorescence Intensity (MFI) values of GFP (C-terminal) detection in BFP (N-terminal) positive cells for both WT eRF1 and E52D. A paired t-test statistical analysis was performed and no significance was attributed.

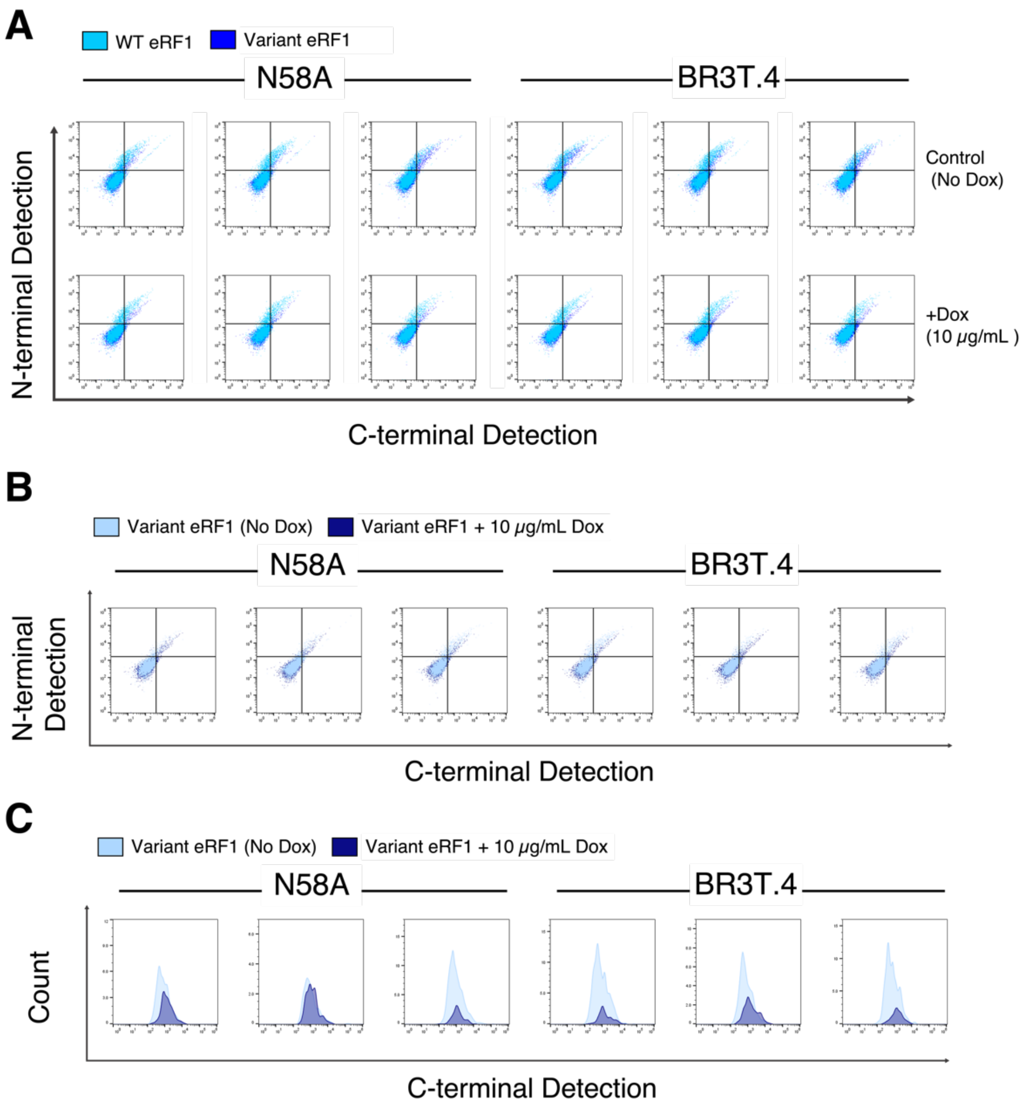


**Figure S12. Analysis of eRF1 mutants in the absence of endogenous eRF1 expression.** A) Flow cytometry analysis of WT eRF1 (cyan) overlaid on single mutant N58A or sorted clone BR3T.4 (cobalt) transformed into tet-repressible eRF1 yeast strain YTH41 alongside intracellular BXGaltTAG reporter/OTS. Analysis performed in the presence (bottom row; endogenous eRF1 shut off) and absence (top row; endogenous eRF1 co-expressed) of 10 µg/mL doxycycline. Data collected in biological triplicate, all three replicates shown. X-axis represents C-terminal detection, Y-axis represents N-terminal detection. B) Flow cytometry analysis of single mutant N58A or sorted clone BR3T.4 in the presence (dark indigo) or absence (light blue) of 10 µg/mL doxycycline. Data collected in biological triplicate, all three replicates shown. X-axis represents C-terminal detection, y-axis represents N-terminal detection. C) Flow cytometry histograms of single mutant N58A or sorted clone BR3T.4 (C-terminal fluorescence levels of cells positive for N-terminal fluorescence) in the presence (dark indigo) or absence (light blue) of 10 µg/mL doxycycline. Data collected in biological triplicate, all three replicates shown. X-axis represents C-terminal fluorescence levels, y-axis represents event counts at a specific fluorescence level.

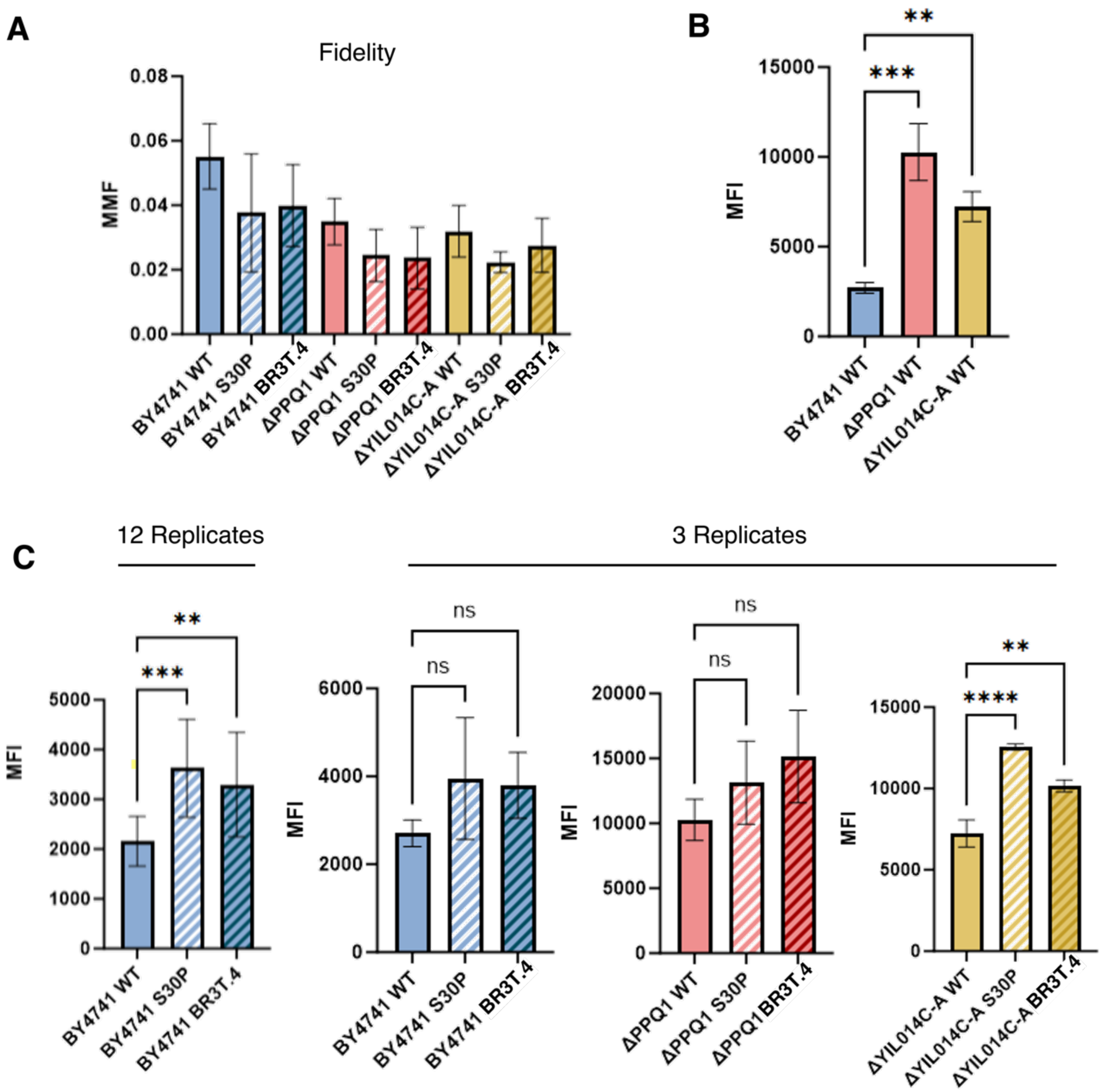


**Figure S13. Investigation of eRF1 mutants in various cell strains.** A) Maximum misincorporation frequency (MMF) of OmeY incorporation in cells transformed with plasmid-based 2$^{nd}$ copy of wild type eRF1 (WT), S30P (single mutant clone) or BR3T.4 (sorted clone) in yeast strains BY4741, BY4741ΔPPQ1, and BY4741ΔYIL014C-A. B) Median Fluorescence Intensity (MFI) of C-terminal (GFP) fluorescence values in BFP-positive cells comparing WT eRF1 across three yeast knockout strains known to enhance ncAA incorporation. One-way ANOVA statistical analysis shown with the following p-value assignments: *** $p \leq 0.001$, ** $p \leq 0.01$. C) Median Fluorescence Intensity (MFI) of C-terminal (GFP) fluorescence values in BFP-positive cells comparing eRF1 variants (S30P and BR3T.4) to WT eRF1 across three yeast knockout strains known to enhance ncAA incorporation. One-way ANOVA statistical analysis shown with the following p-value assignments: ****$p \leq 0.0001$, *** $p \leq 0.001$, * $p \leq 0.05$, ns $p > 0.05$. Panels A-C were analyzed in biological triplicate, except for the first graph in panel C, which included 11 WT replicates and 12 replicates of S30P and BR3T.4, demonstrating increased significance in ANOVA tests with additional replicates.

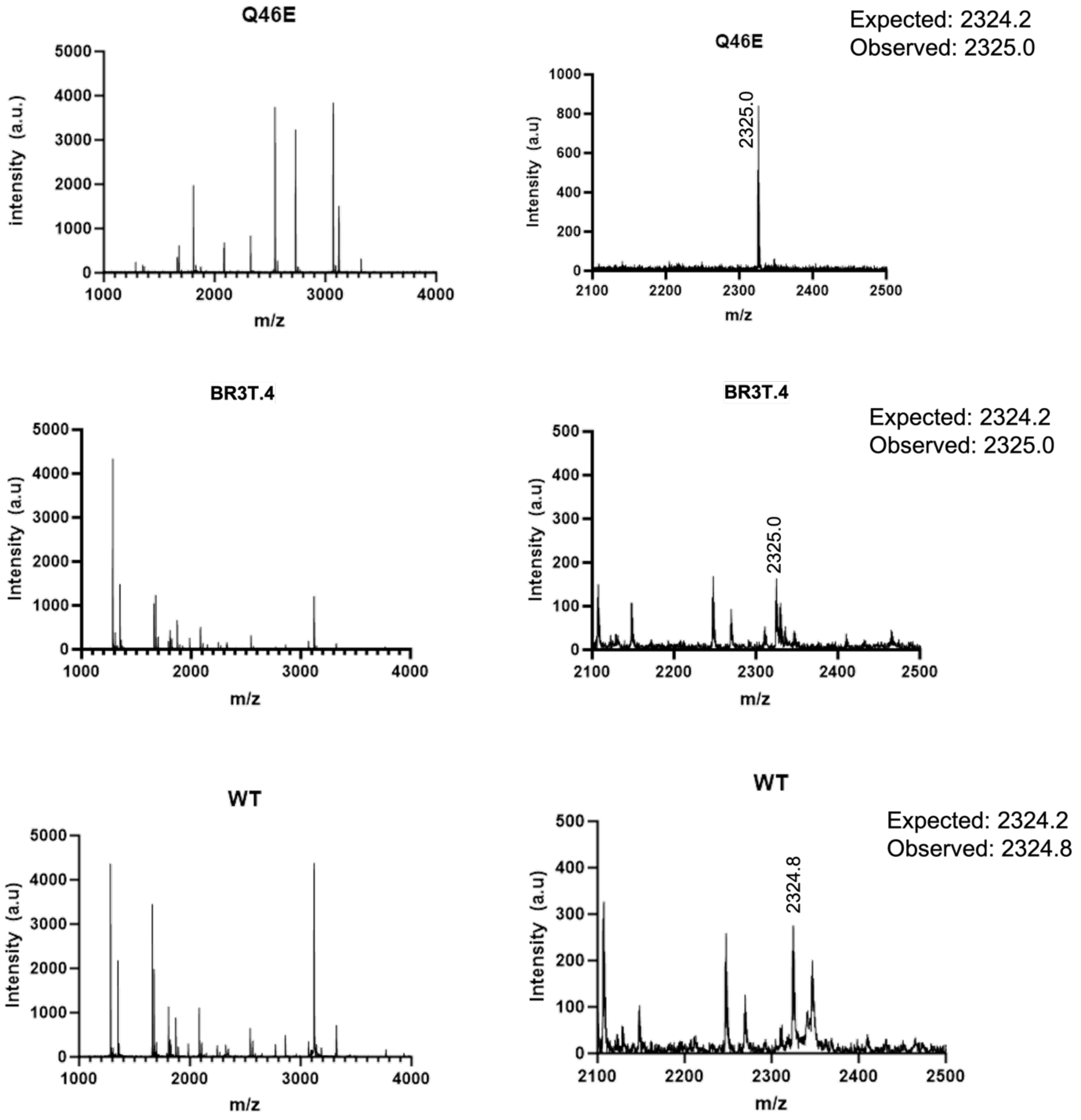


**Fig S14. MALDI mass spectrometry data of tryptic-digested TAG containing reporter scFv (Donkey 1.1 H54-TAG) with O-methyl-L-Tyrosine inserted at the TAG position.** Scfvs were expressed under three conditions; in the presence of plasmid-based WT eRF1, eRF1 with a Q46E mutation, and eRF1 sorted clone BR3T.4 (mutations N64D, K105R, and P116L).

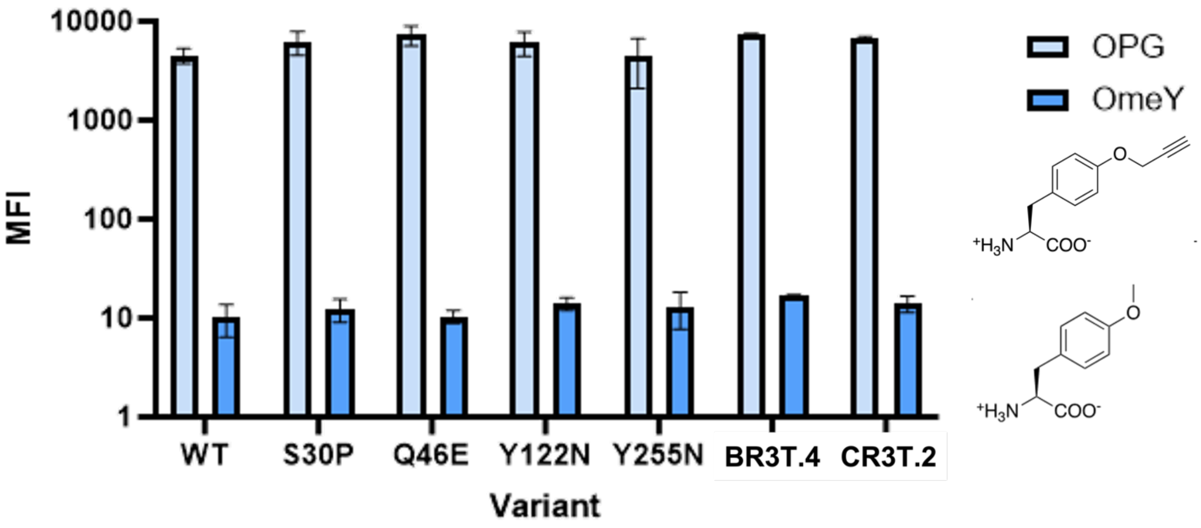


**Fig S15. Click chemistry confirmation of specific ncAA incorporation.** MFI of PE-anti-biotin detection (click chemistry) for C-myc-positive cells (chicken anti-cmyc AF647 positive cells). Cells induced in presence of alkyne-containing *p*-propargyloxy-L-phenylalanine (OPG) that enables copper-catalyzed azide-alkyne click chemistry (CuAAC) or *O*-methyl-L-tyrosine (OmeY) control that is not amenable to click chemistry due to lack of alkyne group. Experiments performed in biological triplicate.

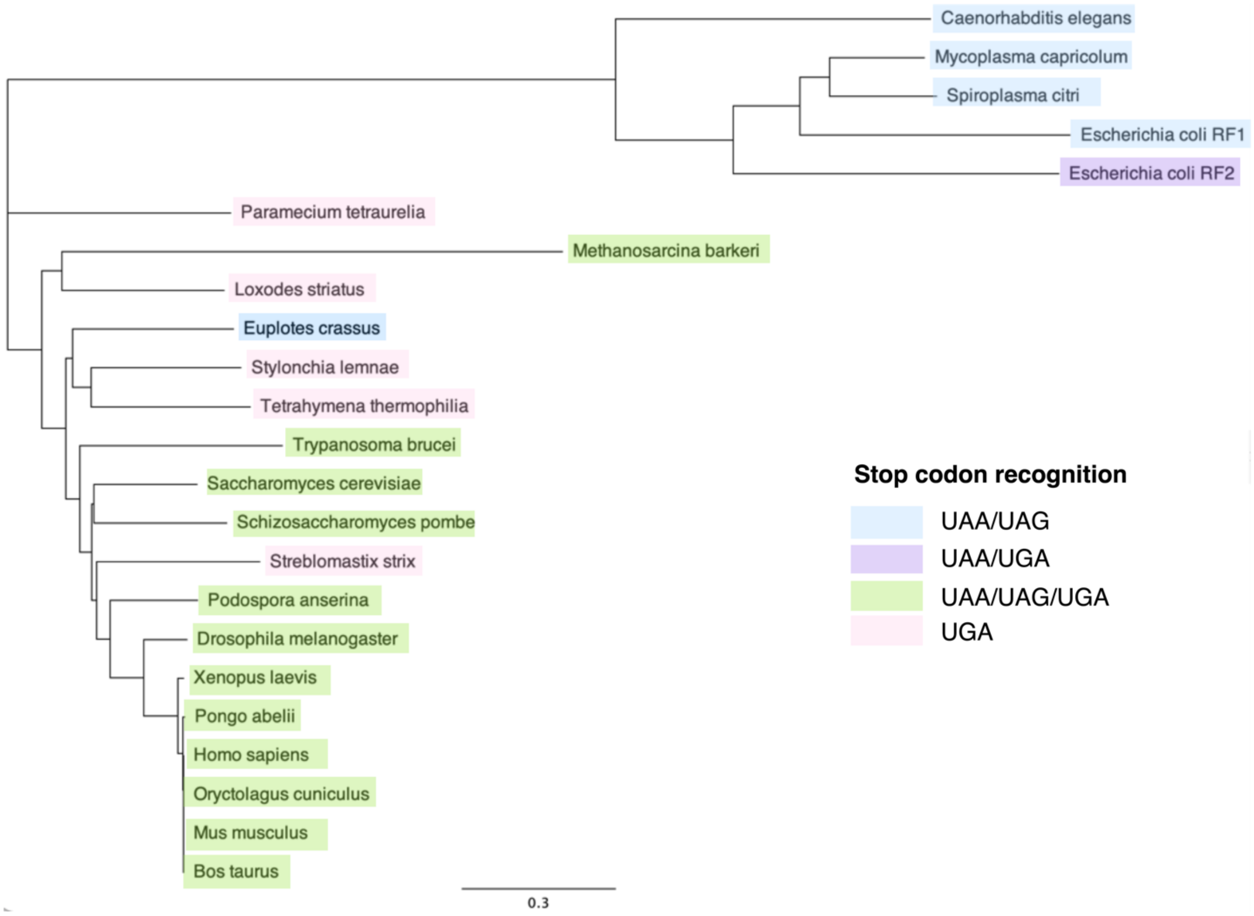


**Figure S16. Standard Global Alignment Tree of organisms with homologous eRF1.** Tree of organisms listed in Table S1, created in Geneious Prime® 2025.0.3[3].

| Stop codon recognition | Gene | Organism | 20 | 25 | 28 | 29 | 30 | 31 | 46 | 52 | 55 | 56 | 57 | 58 | 59 | 60 | 61 | 64 | 67 | 68 | 69 | 70 | 74 | 76 | 103 | 105 | 109 | 116 | 119 | 120 | 121 | 122 | 123 | 124 | 125 | 126 | 127 | 128 | 141 |
|---|---|---|---|---|---|---|---|---|---|---|---|---|---|---|---|---|---|---|---|---|---|---|---|---|---|---|---|---|---|---|---|---|---|---|---|---|---|---|---|
| UAA/UAG/UGA | SUP45 | *Saccharomyces cerevisiae** | S | R | G | T | S | M | Q | E | T | A | S | N | I | K | S | N | S | V | L | S | S | Q | K | K | F | P | T | S | L | Y | L | C | D | N | K | F | D |
| UAA/UAG/UGA | sup45 | *Schizosaccharomyces pombe* | Q | H | G | T | S | M | S | E | T | A | S | N | I | K | S | N | S | V | L | S | S | R | K | R | I | P | T | S | Q | Y | L | C | D | N | K | F | Q |
| UAA/UAG/UGA | SU2 | *Podospora anserina* | R | R | G | T | S | M | A | E | T | A | S | N | I | K | S | N | S | V | L | S | S | Q | K | R | I | P | T | S | L | Y | L | C | D | N | K | F | Q |
| UAA/UAG/UGA | eRF1 | *Drosophila melanogaster* | S | R | G | T | S | M | S | E | T | A | S | N | I | K | S | N | S | V | L | G | S | Q | K | K | I | P | T | S | L | Y | L | C | D | N | K | F | N |
| UAA/UAG/UGA | etf1 | *Xenopus laevis* | S | R | G | T | S | M | A | E | T | A | S | N | I | K | S | N | S | V | L | G | S | Q | K | K | I | P | T | S | L | Y | L | C | D | N | K | F | S |
| UAA/UAG/UGA | ETF1 | *Homo sapiens* | S | R | G | T | S | M | A | E | T | A | S | N | I | K | S | N | S | V | L | G | S | Q | K | K | I | P | T | S | L | Y | L | C | D | N | K | F | S |
| UAA/UAG/UGA | ETF1 | *Oryctolagus cuniculus* | S | R | G | T | S | M | A | E | T | A | S | N | I | K | S | N | S | V | L | G | S | Q | K | K | I | P | T | S | L | Y | L | C | D | N | K | F | S |
| UAA/UAG/UGA | Etf1 | *Mus musculus* | S | R | G | T | S | M | A | E | T | A | S | N | I | K | S | N | S | V | L | G | S | Q | K | K | I | P | T | S | L | Y | L | C | D | N | K | F | S |
| UAA/UAG/UGA | ETF1 | *Bos taurus* | S | R | G | T | S | M | A | E | T | A | S | N | I | K | S | N | S | V | L | G | S | Q | K | K | I | P | T | S | L | Y | L | C | D | N | K | F | S |
| UAA/UAG/UGA | ETF1 | *Pongo abelii* | S | R | G | T | S | M | A | E | T | A | S | N | I | K | S | N | S | V | L | G | S | Q | K | K | I | P | T | S | L | Y | L | C | D | N | K | F | S |
| UAA/UAG/UGA | ERF1 | *Trypanosoma brucei* | M | R | G | T | S | V | V | E | T | A | S | N | I | K | S | N | S | V | Q | S | A | L | K | K | L | P | R | S | L | Y | L | C | D | N | K | F | E |
| UAA/UAG/UGA | prf1 | *Methanosarcina barkeri* | - | - | - | - | - | - | - | E | Q | A | S | G | I | K | S | S | N | V | Q | G | F | L | R | G | E | P | D | Y | I | Y | R | C | D | S | V | F | S |
| UAA/UAG | mtRF1 | *Caenorhabditis elegans* | Q | Q | P | T | S | - | S | S | G | E | K | F | V | K | T | D | A | Q | C | Q | A | P | T | I | K | K | E | V | - | - | - | - | D | V | E | S | - |
| UAA, UAG | prfA | *Mycoplasma capricolum* | A | Q | - | - | - | - | Q | E | I | L | K | D | I | K | K | E | K | E | K | A | E | V | D | E | L | E | D | K | - | - | - | - | - | - | - | - | - |
| UAA, UAG | prfA | *Spiroplasma citri* | R | L | - | - | - | - | N | P | I | V | N | D | V | K | K | S | K | E | Q | A | E | V | D | E | L | T | - | Y | - | - | - | - | - | S | E | I | - |
| UAA, UAG | eRF1 | *Euplotes crassus* | N | R | G | T | S | M | Q | E | T | A | S | H | I | K | S | N | A | V | M | S | S | K | S | K | I | P | T | F | K | Y | I | C | D | N | K | F | E |
| UAA/UAG | prfA | *Escherichia coli* | K | H | - | - | - | - | Q | A | I | A | D | Q | E | R | F | L | E | Y | A | Q | D | S | M | E | D | E | - | - | - | - | - | - | - | - | - | - | - |
| UAA/UGA | prfB | *Escherichia coli* | R | R | L | D | - | - | N | P | - | - | - | D | V | W | N | E | Q | A | L | G | S | L | - | - | V | T | E | A | V | - | - | - | - | - | - | A | - |
| UGA | eRF1-1 | *Paramecium tetraurelia* | T | R | G | T | S | V | T | Q | E | A | A | S | I | K | D | N | S | V | Q | E | A | I | G | K | V | P | L | S | L | Y | F | C | D | P | Q | F | P |
| UGA | eRF1 | *Loxodes striatus* | Y | K | G | T | S | L | N | A | R | A | Q | N | I | K | S | V | A | V | Q | D | S | K | T | K | E | A | T | F | K | F | L | C | E | N | T | F | D |
| UGA | ERF1 | *Stylonychia lemnae* | K | K | G | T | S | M | G | E | A | A | Q | N | I | K | S | T | S | V | I | T | S | K | T | K | F | P | Q | F | M | Y | F | C | G | G | K | F | D |
| UGA | ERF1 | *Tetrahymena thermophilia* | R | R | G | T | S | M | T | E | K | A | T | N | I | K | D | N | S | V | Q | D | S | L | K | I | I | P | T | S | L | Y | F | C | D | S | K | F | P |
| UGA | eRF1 | *Streblomastix strix* | M | R | G | T | S | M | N | E | S | A | K | N | I | K | S | N | S | V | L | D | S | Q | - | K | I | A | T | S | L | Y | L | C | D | F | R | F | D |
| | | ***Consensus*** | ***S*** | ***R*** | ***G*** | ***T*** | ***S*** | ***M*** | ***A*** | ***E*** | ***T*** | ***A*** | ***S*** | ***N*** | ***I*** | ***K*** | ***S*** | ***N*** | ***S*** | ***V*** | ***L*** | ***G*** | ***S*** | ***Q*** | ***K*** | ***K*** | ***I*** | ***P*** | ***T*** | ***S*** | ***L*** | ***Y*** | ***L*** | ***C*** | ***D*** | ***N*** | ***K*** | ***F*** | ***S*** |
| | | | . | : | : | : | : | : | . | : | . | : | : | : | : | : | : | : | : | : | : | . | : | . | . | : | . | : | : | : | : | : | : | : | : | : | : | : | . |
| | | Mutations found in: | | | | | | | | | | | | | | | | | | | | | | | | | | | | | | | | | | | | | |
| | | Sorted clones | P | G | | | P | | | | | | | S | | | P | D | | A | S | F/P | | | T | R | | L | | | | | | R | | | | | G |
| | | Track B DMS | P/A | G | A | | P | R | E/K | K | | S | L/P | H/D/S | V/N/F | T/E | P/T | D | | A | S | P | A | L/P/E | T | R | I/V | L | I | C | S | N | | R | | S/D/I | N | L | Y |
| | | Track C DMS | P | G/S | A/R | | P/Y | R | E/L | K/G | N | P | L | D/H/S | V/M | T/E/I | P | I/H/D | A | A | S | P/T | A | L/P | T | R/T | V | T/L | I | Y/T | V/S | N/D | | R/Y | Y | S/D | R | I/C/L | V |

| Stop codon recognition | Gene | Organism | 151 | 159 | 165 | 169 | 180 | 181 | 182 | 187 | 194 | 203 | 213 | 222 | 227 | 232 | 239 | 253 | 255 | 256 | 258 | 272 | 280 | 288 | 298 | 310 | 314 | 315 | 360 | 361 | 362 | 365 | 366 | 381 | 396 | 397 | 405 | 412 | 421 |
|---|---|---|---|---|---|---|---|---|---|---|---|---|---|---|---|---|---|---|---|---|---|---|---|---|---|---|---|---|---|---|---|---|---|---|---|---|---|---|---|
| UAA/UAG/UGA | SUP45 | *Saccharomyces cerevisiae** | Q | S | V | F | G | G | Q | F | K | A | N | I | A | D | F | V | Y | G | E | L | E | F | F | D | V | E | E | M | D | S | E | G | Q | F | G | V | S |
| UAA/UAG/UGA | sup45 | *Schizosaccharomyces pombe* | H | S | V | F | G | G | Q | F | K | A | D | V | A | E | F | V | Y | G | D | L | E | F | Y | Q | V | E | E | M | E | S | S | G | Q | F | G | L | S |
| UAA/UAG/UGA | SU2 | *Podospora anserina* | N | S | I | F | G | G | Q | F | K | A | N | V | A | D | F | V | Y | G | E | L | E | F | I | E | V | E | E | M | E | S | Q | G | Q | F | G | V | - |
| UAA/UAG/UGA | eRF1 | *Drosophila melanogaster* | N | Q | V | F | G | G | Q | F | K | A | N | I | A | E | F | V | Y | G | E | L | E | F | Y | E | V | E | E | M | E | E | S | G | Q | F | G | V | - |
| UAA/UAG/UGA | etf1 | *Xenopus laevis* | S | Q | V | F | G | G | Q | F | K | A | G | V | A | E | F | I | Y | G | E | L | E | F | Y | E | V | E | E | H | E | E | S | G | Q | F | G | V | G |
| UAA/UAG/UGA | ETF1 | *Homo sapiens* | S | Q | V | F | G | G | Q | F | K | A | G | V | A | E | F | I | Y | G | E | L | E | F | Y | E | V | E | E | H | E | E | S | G | Q | F | G | V | - |
| UAA/UAG/UGA | ETF1 | *Oryctolagus cuniculus* | S | Q | V | F | G | G | Q | F | K | A | G | V | A | E | F | I | Y | G | E | L | E | F | Y | E | V | E | E | H | E | E | S | G | Q | F | G | V | - |
| UAA/UAG/UGA | Etf1 | *Mus musculus* | S | Q | V | F | G | G | Q | F | K | A | G | V | A | E | F | I | Y | G | E | L | E | F | Y | E | V | E | E | H | E | E | S | G | Q | F | G | V | - |
| UAA/UAG/UGA | ETF1 | *Bos taurus* | S | Q | V | F | G | G | Q | F | K | A | G | V | A | E | F | I | Y | G | E | L | E | F | Y | E | V | E | E | H | E | E | S | G | Q | F | G | V | - |
| UAA/UAG/UGA | ETF1 | *Pongo abelii* | S | Q | V | F | G | G | Q | F | K | A | G | V | A | E | F | I | Y | G | E | L | E | F | Y | E | V | E | E | H | E | E | S | G | Q | F | G | V | G |
| UAA/UAG/UGA | ERF1 | *Trypanosoma brucei* | S | C | K | F | G | G | Q | F | R | A | N | V | A | V | F | V | H | P | D | L | E | F | Y | E | V | E | T | R | N | E | Q | G | Q | F | G | L | E |
| UAA/UAG/UGA | prf1 | *Methanosarcina barkeri* | R | V | V | L | G | G | Q | F | A | G | V | L | S | E | L | T | Y | T | E | L | Q | F | V | E | V | D | A | L | E | D | A | N | Q | L | A | T | - |
| UAA/UAG | mtRF1 | *Caenorhabditis elegans* | - | P | V | C | G | G | Q | F | L | A | Q | V | I | E | R | - | N | D | R | P | P | - | G | R | K | E | K | L | Q | S | K | R | E | F | Q | L | - |
| UAA, UAG | prfA | *Mycoplasma capricolum* | - | P | V | I | G | G | D | F | L | E | E | Y | F | D | K | T | A | K | R | M | I | - | G | R | L | P | K | N | A | T | G | R | Q | V | D | I | - |
| UAA, UAG | prfA | *Spiroplasma citri* | - | P | V | I | G | G | D | F | L | E | E | F | F | D | K | T | S | K | R | I | I | - | G | R | L | P | K | D | A | T | G | R | Q | I | D | I | - |
| UAA, UAG | eRF1 | *Euplotes crassus* | A | Q | T | F | G | G | Q | F | K | A | D | V | A | D | F | V | Y | G | E | L | E | F | I | E | C | E | D | F | D | K | C | G | Q | F | G | I | K |
| UAA/UAG | prfA | *Escherichia coli RF1* | - | P | A | - | G | G | D | F | L | E | S | Y | A | D | K | T | S | Q | R | L | I | - | G | R | L | P | R | N | L | S | G | R | E | V | D | I | - |
| UAA/UGA | prfB | *Escherichia coli RF2* | - | F | S | Y | G | G | T | W | L | E | E | I | I | D | R | F | S | G | R | V | I | - | G | R | I | P | K | S | D | W | G | D | D | - | D | L | - |
| UGA | eRF1-1 | *Paramecium tetraurelia* | N | Q | I | F | G | G | Q | F | K | C | E | V | A | D | F | I | Y | G | E | L | E | F | V | E | I | K | Q | F | V | V | K | G | Q | F | G | M | - |
| UGA | eRF1 | *Loxodes striatus* | N | Q | I | F | G | G | Q | F | K | A | N | V | A | E | L | V | Y | G | E | L | E | F | V | E | V | E | E | L | N | E | R | G | Q | F | C | V | E |
| UGA | ERF1 | *Stylonychia lemnae* | N | Q | I | I | G | G | Q | F | K | A | N | V | A | E | L | V | Y | G | E | L | E | F | I | E | V | E | D | M | D | A | E | G | Q | F | G | V | V |
| UGA | ERF1 | *Tetrahymena thermophilia* | Q | Q | V | F | G | G | Q | F | K | C | Q | V | G | E | F | V | Y | G | E | L | E | F | V | L | I | E | E | L | E | E | V | G | Q | F | G | V | - |
| UGA | eRF1 | *Streblomastix strix* | S | Q | I | F | G | G | Q | F | K | A | D | V | A | E | L | V | Y | G | E | L | E | F | Y | S | V | Q | E | L | L | N | K | G | Q | F | G | L | M |
| | | ***Consensus*** | ***S*** | ***Q*** | ***V*** | ***F*** | ***G*** | ***G*** | ***Q*** | ***F*** | ***K*** | ***A*** | ***X*** | ***V*** | ***A*** | ***E*** | ***F*** | ***V*** | ***Y*** | ***G*** | ***E*** | ***L*** | ***E*** | ***F*** | ***Y*** | ***E*** | ***V*** | ***E*** | ***E*** | ***H*** | ***E*** | ***E*** | ***S*** | ***G*** | ***Q*** | ***F*** | ***G*** | ***V*** | ***X*** |
| | | | . | : | : | : | * | * | : | : | : | : | | : | : | : | : | . | : | : | : | : | : | : | . | : | : | : | : | . | . | . | . | : | : | : | : | : | |
| | | Mutations found in: | | | | | | | | | | | | | | | | | | | | | | | | | | | | | | | | | | | | | |
| | | Sorted clones | | | A | | | | E | L | E | T | | L | | G | | | | | G | | G | L | L | | | | | | N | | G | | | | | A | P |
| | | Track B DMS | R | A | A/D | V/I/L | | V/C | | | | | T | | D/V/P | G/H | I/L/S | I | N/D | V/R | G | W | G | L | L | Y/V | I | D/Q | D/G | K | N | F/T | G/D | A/S | P/R | L/S | A | A/F | P/Y |
| | | Track C DMS | R | A | A/D | V/I/L | | R | E | Y | E | T | T/S | F | D/V/S | G/H | I/S | D | | V | G/D | W | G | | I | Y | I | D | D/G | K | N | F | G | A | P/R | L/S | A | A | P |

**Table S1. Global alignment of homologous release factors.** *Numbering according to *S. cerevisiae* eRF1. Consenus sequence determined by majority with the following symbols to indicate % identity: . < 30% identity, : >30%<100%, *100% identity (excluding gaps) X= no consensus. Accession numbers as follows: *S. cerevisiae* P12385, *S. pombe* P79063, *P. anserina* O59948, *D. melanogaster* Q9VPH7, *X. laevis* P35615, *H. sapiens* P62495, *O. cuniculus* P62497, *M. musculus* Q8BWY3, *B. taurus* Q0VCX5, *P. abelii* Q5R4C7, *T. brucei* Q9NAX8, *M. barkeri* A0A0E3R3Y4, *C. elegans* O44568, *M. capricolum* P71496, *S. citri* Q14QE1, *E. crassus* A0AAD1XDL0, *E. coli* prfA A0AAI9GXY5, *E. coli* prfB P07012, *P. tetraurelia* Q965E7, *L. striatus* Q5CD84, *S. lemnae* Q9BMM0, *T. thermophilia* Q9U8U5, *S. strix* A0A5J4USS5. Consensus residues highlighted as follows: red- positively charged, blue- negatively charged, green- hydrophobic, yellow- polar uncharged, purple- special cases, black- variant (no consensus) residues found in sorted clones.

**Table S2: Primers used in this study**

| Experiment | Label | Sequence (5’→ 3’) |
|---|---|---|
| Library Creation & Single Mutant Cloning | SUP45_Cloning_Fwd (BglII) | CAATAATGGATAACGAGGTTGAAAAAAATATTGAGATCTGGAAGG |
| | SUP45_Cloning_Rev (SpeI) | AAATGAAATCATAGTCGGATCCTTCATCTTCGTCATAATATTCATCC |
| Sanger Sequencing | SUP45_Seq_Fwd | CTGTTGGTGTGGCCTTAACG |
| | SUP45_Seq_Rev | CACGGTCCTCTAAACCCACTATG |
| Deep Sequencing | Frag1_Fwd | TCCCTACACGACGCTCTTCCGATCTTTAAGACTACAGAAATAGACAAAGG |
| | Frag1_Rev | GTTCAGACGTGTGCTCTTCCGATCTTGTGTTGATAGGTTTGTAAGGTTCG |
| | Frag2_Fwd | TCCCTACACGACGCTCTTCCGATCTGGTAAAGAAAAAAAGGTCACTTTTG |
| | Frag2_Rev | GTTCAGACGTGTGCTCTTCCGATCTCGGTCTTAAAGTCAGCAGAACCAGC |
| | Frag3_Fwd | TCCCTACACGACGCTCTTCCGATCTCTAATGACAAAGTCAATGTTAAGGG |
| | Frag3_Rev | GTTCAGACGTGTGCTCTTCCGATCTTTATAACCTCATTATCCTCGGCATC |
| | Frag4_Fwd | TCCCTACACGACGCTCTTCCGATCTAAAATTTGGAAACTATCAGATATAC |

| | | |
|---|---|---|
| | Frag4_Rev | GTTCAGACGTGTGCTCTTCCGATCTTTTTGATTCGATTTTTTTCTCCCCC |
| Single Mutant Cloning | P2_G28A | CTAAGGAAATCATAGAAGTAGCATTACCTCTAGCTTTTTCTAAAGATTGG |
| | P3_G28A | CCAATCTTTAGAAAAAGCTAGAGGTAATGCTACTTCTATGATTTCCTTAG |
| | P2_Q46E | CCATATTCATCTGTTAACATTTTTTCGTACAGTGGAATTTGACCCTTAG |
| | P3_Q46E | CTAAGGGTCAAATTCCACTGTACGAAAAAATGTTAACAGATGAATATGG |
| | P2_I59V | GATAAAACGGAAAGACGATTAACCCTAGATTTAACATTCGAGGCAGTACCATATTC |
| | P3_I59V | GAATATGGTACTGCCTCGAATGTTAAATCTAGGGTTAATCGTCTTTCCGTTTTATC |
| | P2_S61P | GCAGATAAAACGGAAAGACGATTAACCCTAGGTTTAATATTCGAGGCAGTACC |
| | P3_S61P | GGTACTGCCTCGAATATTAAACCTAGGGTTAATCGTCTTTCCGTTTTATCTGC |
| | P2_Y122N | CTGTATGAAATTTGTTATCACACAAATTTAAGGATGTGTTGATAGGTTTGTAAGG |
| | P3_Y122N | CCTTACAAACCTATCAACACATCCTTAAATTTGTGTGATAACAAATTTCATACAG |

| | | |
|---|---|---|
| | P2_F169V | CCATGCTTTTTGGCAG ATCGACAGTAACTTTAT GTAAAACAGTTCTCGTA TTACCGG |
| | P3_F169V | CCGGTAATACGAGAACT GTTTTACATAAAGTTAC TGTCGATCTGCCAAAAA AGCATGG |
| | P2_Q182E | CTCTTAAACGAGCAAAA CGAAGCGCAGATTCACC ACCTCTACCATGCTTTT TTGGC |
| | P3_Q182E | GCCAAAAAAGCATGGT AGAGGTGGTGAATCTGC GCTTCGTTTTGCTCGTTT AAGAG |
| | P2_F239I | GGAAATAACCTTACATG CTAGTCTTGGATCGATT AATTCAGATTTAGCCAA ATCGG |
| | P3_F239I | CCGATTTGGCTAAATCT GAATTAATCGATCCAAG ACTAGCATGTAAGGTTA TTTCC |
| | P2_Y255N | CGATAGCCTGGTTGAAA CCGTTTTCACCACCATT AGAAACATCCACGATG GAAATAACC |
| | P3_Y255N | GGTTATTTCCATCGTGG ATGTTTCTAATGGTGGT GAAAACGGTTTCAACCA GGCTATCG |
| | P2_D310Y | CAATTAATTTTTCGACT GCACCTAAATACAATGC CTTTAAAGTATCATCTA TACC |
| | P3_D310Y | GGTATAGATGATACTTT AAAGGCATTGTATTTAG GTGCAGTCGAAAAATTA ATTG |

| | | |
|---|---|---|
| | P2_G381A | |
| | P3_G381A | |
| Sanger Sequencing His-modified Variants | Seq_his_SUP45_fwd | CCATTTTGTAATTTCGT GTCGTTTCTATTATG |
| | Seq_his_SUP45_rev | CGAAAATCATTTAATTG GTGGTGCTGC |

**eRF1 Sequences**

**PTH353-SUP45-WT**
*SUP45* Promoter *SUP45-WT*

TCGCGCGTTTCGGTGATGACGGTGAAAACCTCTGACACATGCAGCTCCCGGAGACG
GTCACAGCTTGTCTGTAAGCGGATGCCGGGAGCAGACAAGCCCGTCAGGGCGCGTC
AGCGGGTGTTGGCGGGTGTCGGGGCTGGCTTAACTATGCGGCATCAGAGCAGATTG
TACTGAGAGTGCACCATATCGACTACGTCGTAAGGCCGTTTCTGACAGAGTAAAATT
CTTGAGGGAACTTTCACCATTATGGGAAATGGTTCAAGAAGGTATTGACTTAAACTC
CATCAAATGGTCAGGTCATTGAGTGTTTTTTATTTGTTGTATTTTTTTTTTTTTAGAGA
AAATCCTCCAATATCAAATTAGGAATCGTAGTTTCATGATTTTCTGTTACACCTAACT
TTTTGTGTGGTGCCCTCCTCCTTGTCAATATTAATGTTAAAGTGCAATTCTTTTTCCTT
ATCACGTTGAGCCATTAGTATCAATTTGCTTACCTGTATTCCTTTACTATCCTCCTTTT
TCTCCTTCTTGATAAATGTATGTAGATTGCGTATATAGTTTCGTCTACCCTATGAACA
TATTCCATTTTGTAATTTCGTGTCGTTTCTATTATGAATTTCATTTATAAAGTTTATGT
ACAAATATCATAAAAAAAGAGAATCTTTTTAAGCAAGGATTTTCTTAACTTCTTCGG
CGACAGCATCACCGACTTCGGTGGTACTGTTGGAACCACCTAAATCACCAGTTCTGA
TACCTGCATCCAAAACCTTTTTAACTGCATCTTCAATGGCCTTACCTTCTTCAGGCAA
GTTCAATGACAATTTCAACATCATTGCAGCAGACAAGATAGTGGCGATAGGGTCAA
CCTTATTCTTTGGCAAATCTGGAGCAGAACCGTGGCATGGTTCGTACAAACCAAATG
CGGTGTTCTTGTCTGGCAAAGAGGCCAAGGACGCAGATGGCAACAAACCCAAGGAA
CCTGGGATAACGGAGGCTTCATCGGAGATGATATCACCAAACATGTTGCTGGTGATT
ATAATACCATTTAGGTGGGTTGGGTTCTTAACTAGGATCATGGCGGCAGAATCAATC
AATTGATGTTGAACCTTCAATGTAGGGAATTCGTTCTTGATGGTTTCCTCCACAGTTT
TTCTCCATAATCTTGAAGAGGCCAAAACATTAGCTTTATCCAAGGACCAAATAGGCA
ATGGTGGCTCATGTTGTAGGGCCATGAAAGCGGCCATTCTTGTGATTCTTTGCACTTC
TGGAACGGTGTATTGTTCACTATCCCAAGCGACACCATCACCATCGTCTTCCTTTCTC
TTACCAAAGTAAATACCTCCCACTAATTCTCTGACAACAACGAAGTCAGTACCTTTA
GCAAATTGTGGCTTGATTGGAGATAAGTCTAAAAGAGAGTCGGATGCAAAGTTACA

TGGTCTTAAGTTGGCGTACAATTGAAGTTCTTTACGGATTTTTAGTAAACCTTGTTCA
GGTCTAACACTACCGGTACCCCATTTAGGACCACCCACAGCACCTAACAAAACGGC
ATCAACCTTCTTGGAGGCTTCCAGCGCCTCATCTGGAAGTGGGACACCTGTAGCATC
GATAGCAGCACCACCAATTAAATGATTTTCGAAATCGAACTTGACATTGGAACGAA
CATCAGAAATAGCTTTAAGAACCTTAATGGCTTCGGCTGTGATTTCTTGACCAACGT
GGTCACCTGGCAAAACGACGATCTTCTTAGGGGCAGACATAGGGGCAGACATTAGA
ATGGTATATCCTTGAAATATATATATATATTGCTGAAATGTAAAAGGTAAGAAAAGT
TAGAAAGTAAGACGATTGCTAACCACCTATTGGAAAAAACAATAGGTCCTTAAATA
ATATTGTCAACTTCAAGTATTGTGATGCAAGCATTTAGTCATGAACGCTTCTCTATTC
TATATGAAAAGCCGGTTCCGGCCTCTCACCTTTCCTTTTTCTCCCAATTTTTCAGTTG
AAAAAGGTATATGCGTCAGGCGACCTCTGAAATTAACAAAAAATTTCCAGTCATCG
AATTTGATTCTGTGCGATAGCGCCCCTGTGTGTTCTCGTTATGTTGAGGAAAAAAAT
AATGGTTGCTAAGAGATTCGAACTCTTGCATCTTACGATACCTGAGTATTCCCACAG
TTAACTGCGGTCAAGATATTTCTTGAATCAGGCGCCTTAGACCGCTCGGCCAAACAA
CCAATTACTTGTTGAGAAATAGAGTATAATTATCCTATAAATATAACGTTTTTGAAC
ACACATGAACAAGGAAGTACAGGACAATTGATTTTGAAGAGAATGTGGATTTTGAT
GTAATTGTTCCGATTCCATTTTTAATAAGGCAATAATATTAGGTATGTGGATATACTA
GAAGTTCTCCTCGAGGGTCGATATGCGGTGTGAAATACCGCACAGATGCGTAAGGA
GAAAATACCGCATCAGGAAATTGTAAACGTTAATATTTTGTTAAAATTCGCGTTAAA
TTTTTGTTAAATCAGCTCATTTTTTAACCAATAGGCCGAAATCGGCAAAATCCCTTAT
AAATCAAAAGAATAGACCGAGATAGGGTTGAGTGTTGTTCCAGTTTGGAACAAGAG
TCCACTATTAAAGAACGTGGACTCCAACGTCAAAGGGCGAAAAACCGTCTATCAGG
GCGATGGCCCACTACGTGAACCATCACCCTAATCAAGTTTTTTGGGGTCGAGGTGCC
GTAAAGCACTAAATCGGAACCCTAAAGGGAGCCCCCGATTTAGAGCTTGACGGGGA
AAGCCGGCGAACGTGGCGAGAAAGGAAGGGAAGAAAGCGAAAGGAGCGGGCGCTA
GGGCGCTGGCAAGTGTAGCGGTCACGCTGCGCGTAACCACCACACCCGCCGCGCTT
AATGCGCCGCTACAGGGCGCGTCGCGCCATTCGCCATTCAGGCTGCGCAACTGTTGG
GAAGGGCGATCGGTGCGGGCCTCTTCGCTATTACGCCAGCTGGCGAAGGGGGGATG
TGCTGCAAGGCGATTAAGTTGGGTAACGCCAGGGTTTTCCCAGTCACGACGTTGTAA
AACGACGGCCAGTGAATTGTAATACGACTCACTATAGGGCGAATTGGAGCTCCACC
GCGGTGGCGGCCGCTCTAGACAATACGAAGGAACGATCCCGCGTCAGCAGATGTCA
CAATTTGATTATTTCATTGTCACATGCTTTTTGACTCATCCTTCCGCGAAAGGAGAAC
TTTTTACAATGCCGCCGGCTCTGTTAGCGTACCCTTCTACAAGCAGATAGCAGAACA
AACACATGATATATTCAAAAGGTGCAATGTCAGAGAACATATATTGCGCCCCTGTCC
TGTAGACATCAGTCATTTTTCGCGGGACTTGAATGGCGCACCATTATCCTATACAGA
AAACATTAAATTTACTGCAAAATTTTGGGCAAACGCTTGGATAGACTATGAATCCGT
CATGAAGACACTACTTGTAAAATTATATATATCCTTTTTTTTCATGTGCTAGTAAGAA
TTAGCAACTACTTTCATTCATCTTCGACGCAACTTCGAGGAAAGCACCTTTTTACTTG
CTGTAGCTCTATTCTCTTCCCCAACCACCTTTTCCTTTATTCCAAAATTTTTAAAACTT
TTTCTGTTACATTATATAATCTTCTGTCTGAAATGTTTGGATATAACGCCTCTTGATC
CACTTTGTATATGCGTGCTATTTATTTTCAGATTTATAAAGAGTATGAGCGTCATTTA
CATAAATAGCTGAAGTTATTCATGGAAAATACGAAGAGCACGTATGTGAGCCAACA
GAACATTTGACGTAAGACTCTACAATGTGCCAAGAACTGGACAAGTAGAGGACTGA
GAACTTTATTTCAATTCATTGCTCCTTTTTGGTGGCGCTACCTTTAGCGAAGGTCAAT
GATGAATGTGCACATGCTGTCGAAACCAAAAAGCAAATTCTAACCAACTTCAAAAT
GACATAGTCATCTGATATTTCTACTCATTATAGATAGTATGGGAGCCTTGAAACGAA

AAGTAAGTAAAAAGCTGGATATGAGCAGTATGAGGTAGACCTTAGCTACATCATTT
CCCCCAATAGCTGCTGCAAATATCTGGTTAAATTTGTGATTCCATGAAGAGGATAAC
AGACTTGTTAAAAAGCATCCTGTCAAAATCTAATTTTTGAAGGGCAGTATTCAATTC
ATAATTTACTTTAGCTTAGATCCTTCCAATTTATACATGGTATTATAACCAGATCATA
AACTACAATCTGTCGCTACCGCATGTACGAAGAATACTTAAGTCACTTGCTCTCTCA
TCATTTGTACATTTTTCAGTAATACCGTTTGATAGCGCCGTCTTTATTACCCGGATTA
TTCCGTTGACCCTGAATGAAAAATTTTTTCAGAAATCCAGTGCTAAGCGTCAAATCA
ATGAAATACATCACTGTATTTTTAACTGATATACTGTTGGTGTGGCCTTAACGACAC
CTTTATTTCTTAATTCATTTCGGCTTGTCTCCTTATTAAGACTACAGAAATAGACAAA
GGAAATACTTCAATAATGGATAACGAGGTTGAAAAAAATATTGAGATCTGGAAGGT
CAAGAAGTTGGTCCAATCTTTAGAAAAAGCTAGAGGTAATGGTACTTCTATGATTTC
CTTAGTTATTCCTCCTAAGGGTCAAATTCCACTGTACCAAAAAATGTTAACAGATGA
ATATGGTACTGCCTCGAATATTAAATCTAGGGTTAATCGTCTTTCCGTTTTATCTGCT
ATCACTTCCACCCAACAAAAGTTGAAGCTATATAATACTTTGCCCAAGAACGGTTTA
GTTTTATATTGTGGTGATATCATCACTGAAGATGGTAAAGAAAAAAAGGTCACTTTT
GACATCGAACCTTACAAACCTATCAACACATCCTTATATTTGTGTGATAACAAATTT
CATACAGAAGTTCTTTCGGAATTGCTTCAAGCTGACGACAAGTTCGGTTTTATAGTC
ATGGACGGTCAAGGTACTTTGTTTGGTTCTGTGTCCGGTAATACGAGAACTGTTTTA
CATAAATTTACTGTCGATCTGCCAAAAAAGCATGGTAGAGGTGGTCAATCTGCGCTT
CGTTTTGCTCGTTTAAGAGAAGAAAAAAGACATAATTATGTGAGAAAGGTCGCCGA
AGTTGCTGTTCAAAATTTTATTACTAATGACAAAGTCAATGTTAAGGGTTTAATTTTA
GCTGGTTCTGCTGACTTTAAGACCGATTTGGCTAAATCTGAATTATTCGATCCAAGA
CTAGCATGTAAGGTTATTTCCATCGTGGATGTTTCTTATGGTGGTGAAAACGGTTTCA
ACCAGGCTATCGAACTTTCTGCCGAAGCGTTGGCCAATGTCAAGTATGTTCAAGAAA
AGAAATTATTGGAGGCATATTTTGACGAAATTTCCCAGGACACTGGTAAATTCTGTT
ATGGTATAGATGATACTTTAAAGGCATTGGATTTAGGTGCAGTCGAAAAATTAATTG
TTTTCGAAAATTTGGAAACTATCAGATATACATTTAAAGATGCCGAGGATAATGAGG
TTATAAAATTCGCTGAACCAGAAGCCAAGGACAAGTCGTTTGCTATTGACAAAGCTA
CCGGCCAAGAAATGGACGTTGTCTCCGAAGAACCTTTAATTGAATGGCTAGCAGCTA
ACTACAAAAACTTCGGTGCTACCTTGGAATTCATCACAGACAAATCTTCAGAAGGTG
CCCAATTTGTCACAGGTTTTGGTGGTATTGGTGCCATGCTGCGTTACAAAGTTAATTT
TGAACAACTAGTTGATGAATCTGAGGATGAATATTATGACGAAGATGAAGGATCCG
ACTATGATTTCATTTAAATAAATAAAAGGGGGAGAAAAAAATCGAATCAAAAAGAA
TTTAATCACTAGATGCCAGATTTAAATTAAATTCGCTTTTAATTTTTTGTACAATATA
ATATATACTTGGTAAACCTTTTGCTCTATATTGAGCTAATTCCTTTGTTGAAAGTACA
TAGTGGGTTTAGAGGACCGTGTATATTACGTAGAAAATACAGTGAAAGGAGAGTTT
CTCTTCAAAAGCCTCGACGGTATCGATAAGCTTATCGATACCGTCGACCTCGAGGGG
GGGCCCGGTACCCAGCTTTTGTTCCCTTTAGTGAGGGTTAATTCCGAGCTTGGCGTA
ATCATGGTCATAGCTGTTTCCTGTGTGAAATTGTTATCCGCTCACAATTCCACACAAC
ATAGGAGCCGGAAGCATAAAGTGTAAAGCCTGGGGTGCCTAATGAGTGAGGTAACT
CACATTAATTGCGTTGCGCTCACTGCCCGCTTTCCAGTCGGGAAACCTGTCGTGCCA
GCTGCATTAATGAATCGGCCAACGCGCGGGGAGAGGCGGTTTGCGTATTGGGCGCT
CTTCCGCTTCCTCGCTCACTGACTCGCTGCGCTCGGTCGTTCGGCTGCGGCGAGCGG
TATCAGCTCACTCAAAGGCGGTAATACGGTTATCCACAGAATCAGGGGATAACGCA
GGAAAGAACATGTGAGCAAAAGGCCAGCAAAAGGCCAGGAACCGTAAAAAGGCCG
CGTTGCTGGCGTTTTTCCATAGGCTCGGCCCCCCTGACGAGCATCACAAAAATCGAC

GCTCAAGTCAGAGGTGGCGAAACCCGACAGGACTATAAAGATACCAGGCGTTCCCC
CCTGGAAGCTCCCTCGTGCGCTCTCCTGTTCCGACCCTGCCGCTTACCGGATACCTGT
CCGCCTTTCTCCCTTCGGGAAGCGTGGCGCTTTCTCAATGCTCACGCTGTAGGTATCT
CAGTTCGGTGTAGGTCGTTCGCTCCAAGCTGGGCTGTGTGCACGAACCCCCCGTTCA
GCCCGACCGCTGCGCCTTATCCGGTAACTATCGTCTTGAGTCCAACCCGGTAAGACA
CGACTTATCGCCACTGGCAGCAGCCACTGGTAACAGGATTAGCAGAGCGAGGTATG
TAGGCGGTGCTACAGAGTTCTTGAAGTGGTGGCCTAACTACGGCTACACTAGAAGG
ACAGTATTTGGTATCTGCGCTCTGCTGAAGCCAGTTACCTTCGGAAAAAGAGTTGGT
AGCTCTTGATCCGGCAAACAAACCACCGCTGGTAGCGGTGGTTTTTTTGTTTGCAAG
CAGCAGATTACGCGCAGAAAAAAAGGATCTCAAGAAGATCCTTTGATCTTTTCTACG
GGGTCTGACGCTCAGTGGAACGAAAACTCACGTTAAGGGATTTTGGTCATGAGATTA
TCAAAAAGGATCTTCACCTAGATCCTTTTAAATTAAAAATGAAGTTTTAAATCAATC
TAAAGTATATATGAGTAAACTTGGTCTGACAGTTACCAATGCTTAATCAGTGAGGCA
CCTATCTCAGCGATCTGTCTATTTCGTTCATCCATAGTTGCCTGACTGCCCGTCGTGT
AGATAACTACGATACGGGAGGGCTTACCATCTGGCCCCAGTGCTGCAATGATACCG
CGAGACCCACGCTCACCGGCTCCAGATTTATCAGCAATAAACCAGCCAGCCGGAAG
GGCCGAGCGCAGAAGTGGTCCTGCAACTTTATCCGCCTCCATCCAGTCTATTAATTG
TTGCCGGGAAGCTAGAGTAAGTAGTTCGCCAGTTAATAGTTTGCGCAACGTTGTTGC
CATTGCTACAGGCATCGTGGTGTCACGCTCGTCGTTTGGTATGGCTTCATTCAGCTCC
GGTTCCCAACGATCAAGGCGAGTTACATGATCCCCCATGTTGTGAAAAAAAGCGGTT
AGCTCCTTCGGTCCTCCGATCGTTGTCAGAAGTAAGTTGGCCGCAGTGTTATCACTC
ATGGTTATGGCAGCACTGCATAATTCTCTTACTGTCATGCCATCCGTAAGATGCTTTT
CTGTGACTGGTGAGTACTCAACCAAGTCATTCTGAGAATAGTGTATGCGGCGACCGA
GTTGCTCTTGCCCGGCGTCAATACGGGATAATACCGCGCCACATAGCAGAACTTTAA
AAGTGCTCATCATTGGAAAACGTTCTTCGGGGCGAAAACTCTCAAGGATCTTACCGC
TGTTGAGATCCAGTTCGATGTAACCCACTCGTGCACCCAACTGATCTTCAGCATCTTT
TACTTTCACCAGCGTTTCTGGGTGAGCAAAAACAGGAAGGCAAAATGCCGCAAAAA
AGGGAATAAGGGCGACACGGAAATGTTGAATACTCATACTCTTCCTTTTTCAATATT
ATTGAAGCATTTATCAGGGTTATTGTCTCATGAGCGGATACATATTTGAATGTATTTA
GAAAAATAAACAAATAGGGGTTCCGCGCACATTTCCCCGAAAAGTGCCACCTGGGT
CCTTTTCATCACGTGCTATAAAAATAATTATAATTTAAATTTTTTAATATAAATATAT
AAATTAAAAATAGAAAGTAAAAAAAGAAATTAAAGAAAAAATAGTTTTTGTTTTCC
GAAGATGTAAAAGACTCTAGGGGGATCGCCAACAAATACTACCTTTTATCTTGCTCT
TCCTGCTCTCAGGTATTAATGCCGAATTGTTTCATCTTGTCTGTGTAGAAGACCACAC
ACGAAAATCCTGTGATTTTACATTTTACTTATCGTTAATCGAATGTATATCTATTTAA
TCTGCTTTTCTTGTCTAATAAATATATATGTAAAGTACGCTTTTTGTTGAAATTTTTTA
AACCTTTGTTTATTTTTTTTTCTTCATTCCGTAACTCTTCTACCTTCTTTATTTACTTTC
TAAAATCCAAATACAAAACATAAAAATAAATAAACACAGAGTAAATTCCCAAATTA
TTCCATCATTAAAAGATACGAGGCGCGTGTAAGTTACAGGCAAGCGATCCGTCCTAA
GAAACCATTATTATCATGACATTAACCTATAAAAATAGGCGTATCACGAGGCCCTTT
CGTC

**PTH353-SUP45-CR3B.6**

*SUP45* Promoter *SUP45*-***CR3B.6***

CAATACGGGATAATACCGCGCCACATAGCAGAACTTTAAAAGTGCTCATCATTGGA
AAACGTTCTTCGGGGCGAAAACTCTCAAGGATCTTACCGCTGTTGAGATCCAGTTCG
ATGTAACCCACTCGTGCACCCAACTGATCTTCAGCATCTTTTACTTTCACCAGCGTTT
CTGGGTGAGCAAAAACAGGAAGGCAAAATGCCGCAAAAAAGGGAATAAGGGCGAC
ACGGAAATGTTGAATACTCATACTCTTCCTTTTTCAATATTATTGAAGCATTTATCAG
GGTTATTGTCTCATGAGCGGATACATATTTGAATGTATTTAGAAAAATAAACAAATA
GGGGTTCCGCGCACATTTCCCCGAAAAGTGCCACCTGGGTCCTTTTCATCACGTGCT
ATAAAAATAATTATAATTTAAATTTTTTAATATAAATATATAAATTAAAAATAGAAA
GTAAAAAAAGAAATTAAAGAAAAAATAGTTTTTGTTTTCCGAAGATGTAAAAGACT
CTAGGGGGATCGCCAACAAATACTACCTTTTATCTTGCTCTTCCTGCTCTCAGGTATT
AATGCCGAATTGTTTCATCTTGTCTGTGTAGAAGACCACACACGAAAATCCTGTGAT
TTTACATTTTACTTATCGTTAATCGAATGTATATCTATTTAATCTGCTTTTCTTGTCTA
ATAAATATATATGTAAAGTACGCTTTTTGTTGAAATTTTTTAAACCTTTGTTTATTTTT
TTTCTTCATTCCGTAACTCTTCTACCTTCTTTATTTACTTTCTAAAATCCAAATACAAA
ACATAAAAATAAATAAACACAGAGTAAATTCCCAAATTATTCCATCATTAAAAGAT
ACGAGGCGCGTGTAAGTTACAGGCAAGCGATCCGTCCTAAGAAACCATTATTATCAT
GACATTAACCTATAAAAATAGGCGTATCACGAGGCCCTTTCGTCTCGCGCGTTTCGG
TGATGACGGTGAAAACCTCTGACACATGCAGCTCCCGGAGACGGTCACAGCTTGTCT
GTAAGCGGATGCCGGGAGCAGACAAGCCCGTCAGGGCGCGTCAGCGGGTGTTGGCG
GGTGTCGGGGCTGGCTTAACTATGCGGCATCAGAGCAGATTGTACTGAGAGTGCAC
CATATCGACTACGTCGTTAAGGCCGTTTCTGACAGAGTAAAATTCTTGAGGGAACTT
TCACCATTATGGGAAATGGTTCAAGAAGGTATTGACTTAAACTCCATCAAATGGTCA
GGTCATTGAGTGTTTTTTATTTGTTGTATTTTTTTTTTTTTAGAGAAAATCCTCCAATA
TATAAATTAGGAATCATAGTTTCATGATTTTCTGTTACACCTAACTTTTTGTGTGGTG
CCCTCCTCCTTGTCAATATTAATGTTAAAGTGCAATTCTTTTTCCTTATCACGTTGAG
CCATTAGTATCAATTTGCTTACCTGTATTCCTTTACATCCTCCTTTTTCTCCTTCTTGA
TAAATGTATGTAGATTGCGTATATAGTTTCGTCTACCCTATGAACATATTCCATTTTG
TAATTTCGTGTCGTTTCTATTATGAATTTCATTTATAAAGTTTATGTACAAATATCAT
AAAAAAAGAGAATCTTTTTAAGCAAGGATTTTCTTAACTTCTTCGGCGACAGCATCA
CCGACTTCGGTGGTACTGTTGGAACCACCTAAATCACCAGTTCTGATACCTGCATCC
AAAACCTTTTTAACTGCATCTTCAATGGCCTTACCTTCTTCAGGCAAGTTCAATGACA
ATTTCAACATCATTGCAGCAGACAAGATAGTGGCGATAGGGTTGACCTTATTCTTTG
GCAAATCTGGAGCAGAACCGTGGCATGGTTCGTACAAACCAAATGCGGTGTTCTTGT
CTGGCAAAGAGGCCAAGGACGCAGATGGCAACAAACCCAAGGAACCTGGGATAAC
GGAGGCTTCATCGGAGATGATATCACCAAACATGTTGCTGGTGATTATAATACCATT
TAGGTGGGTTGGGTTCTTAACTAGGATCATGGCGGCAGAATCAATCAATTGATGTTG
AACCTTCAATGTAGGGAATTCGTTCTTGATGGTTTCCTCCACAGTTTTTCTCCATAAT
CTTGAAGAGGCCAAAACATTAGCTTTATCCAAGGACCAAATAGGCAATGGTGGCTC
ATGTTGTAGGGCCATGAAAGCGGCCATTCTTGTGATTCTTTGCACTTCTGGAACGGT
GTATTGTTCACTATCCCAAGCGACACCATCACCATCGTCTTCCTTTCTCTTACCAAAG
TAAATACCTCCCACTAATTCTCTGACAACAACGAAGTCAGTACCTTTAGCAAATTGT
GGCTTGATTGGAGATAAGTCTAAAAGAGAGTCGGATGCAAAGTTACATGGTCTTAA
GTTGGCGTACAATTGAAGTTCTTTACGGATTTTTAGTAAACCTTGTTCAGGTCTAACA
CTACCGGTACCCCATTTAGGACCACCCACAGCACCTAACAAAACGGCATCAGCCTTC
TTGGAGGCTTCCAGCGCCTCATCTGGAAGTGGAACACCTGTAGCATCGATAGCAGCA
CCACCAATTAAATGATTTCGAAATCGAACTTGACATTGGAACGAACATCAGAAATA

GCTTTAAGAACCTTAATGGCTTCGGCTGTGATTTCTTGACCAACGTGGTCACCTGGC
AAAACGACGATCTTCTTAGGGGCAGACATTAGAATGGTATATCCTTGAAATATATAT
ATATATATTGCTGAAATGTAAAAGGTAAGAAAAGTTAGAAAGTAAGACGATTGCTA
ACCACCTATTGGAAAAAACAATAGGTCCTTAAATAATATTGTCAACTTCAAGTATTG
TGATGCAAGCATTTAGTCATGAACGCTTCTCTATTCTATATGAAAAGCCGGTTCCGG
CGCTCTCACCTTTCCTTTTTCTCCCAATTTTTCAGTTGAAAAAGGTATATGCGTCAGG
CGACCTCTGAAATTAACAAAAAATTTCCAGTCATCGAATTTGATTCTGTGCGATAGC
GCCCCTGTGTGTTCTCGTTATGTTGAGGAAAAAAATAATGGTTGCTAAGAGATTCGA
ACTCTTGCATCTTACGATACCTGAGTATTCCCACAGTTAACTGCGGTCAAGATATTTC
TTGAATCAGGCGCCTTAGACCGCTCGGCCAAACAACCAATTACTTGTTGAGAAATAG
AGTATAATTATCCTATAAATATAACGTTTTTGAACACACATGAACAAGGAAGTACAG
GACAATTGATTTTGAAGAGAATGTGGATTTTGATGTAATTGTTGGGATTCCATTTTTA
ATAAGGCAATAATATTAGGTATGTAGATATACTAGAAGTTCTCCTCGACCGGTCGAT
ATGCGGTGTGAAATACCGCACAGATGCGTAAGGAGAAAATACCGCATCAGGAAATT
GTAAGCGTTAATATTTTGTTAAAATTCGCGTTAAATTTTTGTTAAATCAGCTCATTTT
TTAACCAATAGGCCGAAATCGGCAAAATCCCTTATAAATCAAAAGAATAGACCGAG
ATAGGGTTGAGTGTTGTTCCAGTTTGGAACAAGAGTCCACTATTAAAGAACGTGGAC
TCCAACGTCAAAGGGCGAAAAACCGTCTATCAGGGCGATGGCCCACTACGTGAACC
ATCACCCTAATCAAGTTTTTTGGGGTCGAGGTGCCGTAAAGCACTAAATCGGAACCC
TAAAGGGAGCCCCCGATTTAGAGCTTGACGGGGAAAGCCGGCGAACGTGGCGAGAA
AGGAAGGGAAGAAAGCGAAAGGAGCGGGCGCTAGGGCGCTGGCAAGTGTAGCGGT
CACGCTGCGCGTAACCACCACACCCGCCGCGCTTAATGCGCCGCTACAGGGCGCGT
CCATTCGCCATTCAGGCTGCGCAACTGTTGGGAAGGGCGATCGGTGCGGGCCTCTTC
GCTATTACGCCAGCTGGCGAAAGGGGGATGTGCTGCAAGGCGATTAAGTTGGGTAA
CGCCAGGGTTTTCCCAGTCACGACGTTGTAAAACGACGGCCAGTGAATTGTAATACG
ACTCACTATAGGGCGAATTGGAGCTCCACCGCGGTGGCGGCCGCTCTAGACAATAC
GAAGGAACGATCCCGCGTCAGCAGATGTCACAATTTGATTATTTCATTGTCACATGC
TTTTTGACTCATCCTTCCGCGAAAGGAGAACTTTTTACAATGCCGCCGGCTCTGTTAG
CGTACCCTTCTACAAGCAGATAGCAGAACAAACACATGATATATTCAAAAGGTGCA
ATGTCAGAGAACATATATTGCGCCCCTGTCCTGTAGACATCAGTCATTTTTCGCGGG
ACTTGAATGGCGCACCATTATCCTATACAGAAAACATTAAATTTACTGCAAAATTTT
GGGCAAACGCTTGGATAGACTATGAATCCGTCATGAAGACACTACTTGTAAAATTAT
ATATATCCTTTTTTTTCATGTGCTAGTAAGAATTAGCAACTACTTTCATTCATCTTCG
ACGCAACTTCGAGGAAAGCACCTTTTTACTTGCTGTAGCTCTATTCTCTTCCCCAACC
ACCTTTTCCTTTATTCCAAAATTTTTAAAACTTTTTCTGTTACATTATATAATCTTCTG
TCTGAAATGTTTGGATATAACGCCTCTTGATCCACTTTGTATATGCGTGCTATTTATT
TTCAGATTTATAAAGAGTATGAGCGTCATTTACATAAATAGCTGAAGTTATTCATGG
AAAATACGAAGAGCACGTATGTGAGCCAACAGAACATTTGACGTAAGACTCTACAA
TGTGCCAAGAACTGGACAAGTAGAGGACTGAGAACTTTATTTCAATTCATTGCTCCT
TTTTGGTGGCGCTACCTTTAGCGAAGGTCAATGATGAATGTGCACATGCTGTCGAAA
CCAAAAAGCAAATTCTAACCAACTTCAAAATGACATAGTCATCTGATATTTCTACTC
ATTATAGATAGTATGGGAGCCTTGAAACGAAAAGTAAGTAAAAAGCTGGATATGAG
CAGTATGAGGTAGACCTTAGCTACATCATTTCCCCCAATAGCTGCTGCAAATATCTG
GTTAAATTTGTGATTCCATGAAGAGGATAACAGACTTGTTAAAAAGCATCCTGTCAA
AATCTAATTTTTGAAGGGCAGTATTCAATTCATAATTTACTTTAGCTTAGATCCTTCC
AATTTATACATGGTATTATAACCAGATCATAAACTACAATCTGTCGCTACCGCATGT

ACGAAGAATACTTAAGTCACTTGCTCTCTCATCATTTGTACATTTTTCAGTAATACCG
TTTGATAGCGCCGTCTTTATTACCCGGATTATTCCGTTGACCCTGAATGAAAAATTTT
TTCAGAAATCCAGTGCTAAGCGTCAAATCAATGAAATACATCACTGTATTTTTAACT
GATATACTGTTGGTGTGGCCTTAACGACACCTTTATTTCTTAATTCATTTCGGCTTGT
CTCCTTATTAAGACTACAGAAATAGACAAAGGAAATACTTCAATAATGGATAACGA
GGTTGAAAAAAATATTGAGATCTGGAAGGTCAAGAAGTTGGTCCAA**C**CTTTAGAAA
AAGCTAGAGGTAATGGTACTTCTATGATTTCCTTAGTTATTCCTCCTAAGGGTCAAAT
TCCACTGTACCAAAAAATGTTAACAGATGAATATGGTACTGCCTCGAATATTAAATC
TAGGGTTAATCGTCTTTCCGTTTTATCTGCTATCACTTCCACCCAACAAAAGTTGAAG
CTATATAATACTTTGCCCAAGAACGGTTTAGTTTTATATTGTGGTGATATCATCACTG
AAGATGGTAAAGAAAAAAAGGTCACTTTTGACATCGAACCTTACAAACCTATCAAC
ACATCCTTATATTTGTGTGATAACAAATTTCATACAGAAGTTCTTTCGGAATTGCTTC
AAGCTGACGACAAGTTCGGTTTTATAGTCATGGACGGTCAAGGTACTTTGTTTGGTT
CTGTGTCCGGTAATACGAGAACTGTTTTACATAAATTTACTGTCGATCTGCCAAAAA
AGCATGGTAGAGGTGGTCAATCTGCGCTTCGT**C**TTGCTCGTTTAAGAGAAGAAAAA
AGACATAATTATGTGAGAAAGGTCGCCGAAGTTGCTGTTCAAAATTTTATTACTAAT
GACAAAGTCAATGTTAAGGGTTTAATTTTAGCTGGTTCTGCTGACTTTAAGACCG**G**T
TTGGCTAAATCTGAATTATTCGATCCAAGACTAGCATGTAAGGTTATTTCCATCGTG
GATGTTTCTTATGGTGGTG**G**AAACGGTTTCAACCAGGCTATCGAACTTTCTGCCGAA
GCGTTGGCCAATGTCAAGTATGTTCAAGAAAAGAAATTATTGGAGGCATATTTTGAC
GAAATTTCCCAGGACACTGGTAAATTCTGTTATGGTATAGATGATACTTTAAAGGCA
TTGGATTTAGGTGCAGTCGAAAAATTAATTGTTTTCGAAAATTTGGAAACTATCAGA
TATACATTTAAAGATGCCGAGGATAATGAGGTTATAAAATTCGCTGAACCAGAAGC
CAAGGACAAGTCGTTTGCTATTGACAAAGCTACCGGCCAAGAAATGGACGTTGTCTC
CGAAGAACCTTTAATTGAATGGCTAGCAGCTAACTACAAAAACTTCGGTGCTACCTT
GGAATTCATCACAGACAAATCTTCAGAAGGTGCCCAATTTGTCACAGGTTTTGGTGG
TATTGGTGCCATGCTGCGTTACAAAGTTAATTTTGAACAACTAGTTGATGAATCTGA
GGATGAATATTATGACGAAGATGAAGGATCCGACTATGATTTCATTTAAATAAATAA
AAGGGGGAGAAAAAAATCGAATCAAAAAGAATTTAATCACTAGATGCCAGATTTAA
ATTAAATTCGCTTTTAATTTTTTGTACAATATAATATATACTTGGTAAACCTTTTGCT
CTATATTGAGCTAATTCCTTTGTTGAAAGTACATAGTGGGTTTAGAGGACCGTGTAT
ATTACGTAGAAAATACAGTGAAAGGAGAGTTTCTCTTCAAAAGCCTCGACGGTATC
GATAAGCTTATCGATACCGTCGACCTCGAGGGGGGGCCCGGTACCAGCTTTTGTTCC
CTTTAGTGAGGGTTAATTTCGAGCTTGGCGTAATCATGGTCATAGCTGTTTCCTGTGT
GAAATTGTTATCCGCTCACAATTCCACACAACATACGAGCCGGAAGCATAAAGTGT
AAAGCCTGGGGTGCCTAATGAGTGAGCTAACTCACATTAATTGCGTTGCGCTCACTG
CCCGCTTTCCAGTCGGGAAACCTGTCGTGCCAGCTGCATTAATGAATCGGCCAACGC
GCGGGGAGAGGCGGTTTGCGTATTGGGCGCTCTTCCGCTTCCTCGCTCACTGACTCG
CTGCGCTCGGTCGTTCGGCTGCGGCGAGCGGTATCAGCTCACTCAAAGGCGGTAATA
CGGTTATCCACAGAATCAGGGGATAACGCAGGAAAGAACATGTGAGCAAAAGGCC
AGCAAAAGGCCAGGAACCGTAAAAAGGCCGCGTTGCTGGCGTTTTTCCATAGGCTC
CGCCCCCCTGACGAGCATCACAAAAATCGACGCTCAAGTCAGAGGTGGCGAAACCC
GACAGGACTATAAAGATACCAGGCGTTTCCCCCTGGAAGCTCCCTCGTGCGCTCTCC
TGTTCCGACCCTGCCGCTTACCGGATACCTGTCCGCCTTTCTCCCTTCGGGAAGCGTG
GCGCTTTCTCATAGCTCACGCTGTAGGTATCTCAGTTCGGTGTAGGTCGTTCGCTCCA
AGCTGGGCTGTGTGCACGAACCCCCCGTTCAGCCCGACCGCTGCGCCTTATCCGGTA

ACTATCGTCTTGAGTCCAACCCGGTAAGACACGACTTATCGCCACTGGCAGCAGCCA
CTGGTAACAGGATTAGCAGAGCGAGGTATGTAGGCGGTGCTACAGAGTTCTTGAAG
TGGTGGCCTAACTACGGCTACACTAGAAGAACAGTATTTGGTATCTGCGCTCTGCTG
AAGCCAGTTACCTTCGGAAAAAGAGTTGGTAGCTCTTGATCCGGCAAACAAACCAC
CGCTGGTAGCGGTGGTTTTTTTGTTTGCAAGCAGCAGATTACGCGCAGAAAAAAAGG
ATCTCAAGAAGATCCTTTGATCTTTTCTACGGGGTCTGACGCTCAGTGGAACGAAAA
CTCACGTTAAGGGATTTTGGTCATGAGATTATCAAAAAGGATCTTCACCTAGATCCT
TTTAAATTAAAAATGAAGTTTTAAATCAATCTAAAGTATATATGAGTAAACTTGGTC
TGACAGTTACCAATGCTTAATCAGTGAGGCACCTATCTCAGCGATCTGTCTATTTCGT
TCATCCATAGTTGCCTGACTCCCCGTCGTGTAGATAACTACGATACGGGAGGGCTTA
CCATCTGGCCCCAGTGCTGCAATGATACCGCGAGACCCACGCTCACCGGCTCCAGAT
TTATCAGCAATAAACCAGCCAGCCGGAAGGGCCGAGCGCAGAAGTGGTCCTGCAAC
TTTATCCGCCTCCATCCAGTCTATTAATTGTTGCCGGGAAGCTAGAGTAAGTAGTTC
GCCAGTTAATAGTTTGCGCAACGTTGTTGCCATTGCTACAGGCATCGTGGTGTCACG
CTCGTCGTTTGGTATGGCTTCATTCAGCTCCGGTTCCCAACGATCAAGGCGAGTTAC
ATGATCCCCCATGTTGTGCAAAAAAGCGGTTAGCTCCTTCGGTCCTCCGATCGTTGT
CAGAAGTAAGTTGGCCGCAGTGTTATCACTCATGGTTATGGCAGCACTGCATAATTC
TCTTACTGTCATGCCATCCGTAAGATGCTTTTCTGTGACTGGTGAGTACTCAACCAAG
TCATTCTGAGAATAGTGTATGCGGCGACCGAGTTGCTCTTGCCCGGCGT

**PTH353-SUP45-CR3T.2**
*SUP45* Promoter *SUP45-**CR3T.2***

CTTGCCCGGCGTCAATACGGGATAATACCGCGCCACATAGCAGAACTTTAAAAGTG
CTCATCATTGGAAAACGTTCTTCGGGGCGAAAACTCTCAAGGATCTTACCGCTGTTG
AGATCCAGTTCGATGTAACCCACTCGTGCACCCAACTGATCTTCAGCATCTTTTACTT
TCACCAGCGTTTCTGGGTGAGCAAAAACAGGAAGGCAAAATGCCGCAAAAAAGGG
AATAAGGGCGACACGGAAATGTTGAATACTCATACTCTTCCTTTTTCAATATTATTG
AAGCATTTATCAGGGTTATTGTCTCATGAGCGGATACATATTTGAATGTATTTAGAA
AAATAAACAAATAGGGGTTCCGCGCACATTTCCCCGAAAAGTGCCACCTGGGTCCTT
TTCATCACGTGCTATAAAAATAATTATAATTTAAATTTTTTAATATAAATATATAAAT
TAAAAATAGAAGTAAAAAAAGAAATTAAAGAAAAAATAGTTTTTGTTTTCCGAAG
ATGTAAAAGACTCTAGGGGGATCGCCAACAAATACTACCTTTTATCTTGCTCTTCCT
GCTCTCAGGTATTAATGCCGAATTGTTTCATCTTGTCTGTGTAGAAGACCACACACG
AAAATCCTGTGATTTTACATTTTACTTATCGTTAATCGAATGTATATCTATTTAATCT
GCTTTTCTTGTCTAATAAATATATATGTAAAGTACGCTTTTTGTTGAAATTTTTTAAA
CCTTTGTTTATTTTTTTTCTTCATTCCGTAACTCTTCTACCTTCTTTATTTACTTTCTAA
AATCCAAATACAAAACATAAAAATAAATAAACACAGAGTAAATTCCCAAATTATTC
CATCATTAAAAGATACGAGGCGCGTGTAAGTTACAGGCAAGCGATCCGTCCTAAGA
AACCATTATTATCATGACATTAACCTATAAAAATAGGCGTATCACGAGGCCCTTTCG
TCTCGCGCGTTTCGGTGATGACGGTGAAAACCTCTGACACATGCAGCTCCCGGAGAC
GGTCACAGCTTGTCTGTAAGCGGATGCCGGGAGCAGACAAGCCCGTCAGGGCGCGT
CAGCGGGTGTTGGCGGGTGTCGGGGCTGGCTTAACTATGCGGCATCAGAGCAGATT
GTACTGAGAGTGCACCATATCGACTACGTCGTTAAGGCCGTTTCTGACAGAGTAAAA
TTCTTGAGGGAACTTTCACCATTATGGGAAATGGTTCAAGAAGGTATTGACTTAAAC
TCCATCAAATGGTCAGGTCATTGAGTGTTTTTTATTTGTTGTATTTTTTTTTTTTTAGA

GAAAATCCTCCAATATATAAATTAGGAATCATAGTTTCATGATTTTCTGTTACACCTA
ACTTTTTGTGTGGTGCCCTCCTCCTTGTCAATATTAATGTTAAAGTGCAATTCTTTTTC
CTTATCACGTTGAGCCATTAGTATCAATTTGCTTACCTGTATTCCTTTACATCCTCCTT
TTTCTCCTTCTTGATAAATGTATGTAGATTGCGTATATAGTTTCGTCTACCCTATGAA
CATATTCCATTTTGTAATTTCGTGTCGTTTCTATTATGAATTTCATTTATAAAGTTTAT
GTACAAATATCATAAAAAAAGAGAATCTTTTTAAGCAAGGATTTTCTTAACTTCTTC
GGCGACAGCATCACCGACTTCGGTGGTACTGTTGGAACCACCTAAATCACCAGTTCT
GATACCTGCATCCAAAACCTTTTTAACTGCATCTTCAATGGCCTTACCTTCTTCAGGC
AAGTTCAATGACAATTTCAACATCATTGCAGCAGACAAGATAGTGGCGATAGGGTT
GACCTTATTCTTTGGCAAATCTGGAGCAGAACCGTGGCATGGTTCGTACAAACCAAA
TGCGGTGTTCTTGTCTGGCAAAGAGGCCAAGGACGCAGATGGCAACAAACCCAAGG
AACCTGGGATAACGGAGGCTTCATCGGAGATGATATCACCAAACATGTTGCTGGTG
ATTATAATACCATTTAGGTGGGTTGGGTTCTTAACTAGGATCATGGCGGCAGAATCA
ATCAATTGATGTTGAACCTTCAATGTAGGGAATTCGTTCTTGATGGTTTCCTCCACAG
TTTTTCTCCATAATCTTGAAGAGGCCAAAACATTAGCTTTATCCAAGGACCAAATAG
GCAATGGTGGCTCATGTTGTAGGGCCATGAAAGCGGCCATTCTTGTGATTCTTTGCA
CTTCTGGAACGGTGTATTGTTCACTATCCCAAGCGACACCATCACCATCGTCTTCCTT
TCTCTTACCAAAGTAAATACCTCCCACTAATTCTCTGACAACAACGAAGTCAGTACC
TTTAGCAAATTGTGGCTTGATTGGAGATAAGTCTAAAAGAGAGTCGGATGCAAAGTT
ACATGGTCTTAAGTTGGCGTACAATTGAAGTTCTTTACGGATTTTTAGTAAACCTTGT
TCAGGTCTAACACTACCGGTACCCCATTTAGGACCACCCACAGCACCTAACAAAAC
GGCATCAGCCTTCTTGGAGGCTTCCAGCGCCTCATCTGGAAGTGGAACACCTGTAGC
ATCGATAGCAGCACCACCAATTAAATGATTTTCGAAATCGAACTTGACATTGGAACG
AACATCAGAAATAGCTTTAAGAACCTTAATGGCTTCGGCTGTGATTTCTTGACCAAC
GTGGTCACCTGGCAAAACGACGATCTTCTTAGGGGCAGACATTAGAATGGTATATCC
TTGAAATATATATATATATATTGCTGAAATGTAAAAGGTAAGAAAAGTTAGAAAGT
AAGACGATTGCTAACCACCTATTGGAAAAAACAATAGGTCCTTAAATAATATTGTCA
ACTTCAAGTATTGTGATGCAAGCATTTAGTCATGAACGCTTCTCTATTCTATATGAAA
AGCCGGTTCCGGCGCTCTCACCTTTCCTTTTTCTCCCAATTTTTCAGTTGAAAAAGGT
ATATGCGTCAGGCGACCTCTGAAATTAACAAAAAATTTCCAGTCATCGAATTTGATT
CTGTGCGATAGCGCCCCTGTGTGTTCTCGTTATGTTGAGGAAAAAAATAATGGTTGC
TAAGAGATTCGAACTCTTGCATCTTACGATACCTGAGTATTCCCACAGTTAACTGCG
GTCAAGATATTTCTTGAATCAGGCGCCTTAGACCGCTCGGCCAAACAACCAATTACT
TGTTGAGAAATAGAGTATAATTATCCTATAAATATAACGTTTTTGAACACACATGAA
CAAGGAAGTACAGGACAATTGATTTTGAAGAGAATGTGGATTTTGATGTAATTGTTG
GGATTCCATTTTTAATAAGGCAATAATATTAGGTATGTAGATATACTAGAAGTTCTC
CTCGACCGGTCGATATGCGGTGTGAAATACCGCACAGATGCGTAAGGAGAAAATAC
CGCATCAGGAAATTGTAAGCGTTAATATTTTGTTAAAATTCGCGTTAAATTTTTGTTA
AATCAGCTCATTTTTTAACCAATAGGCCGAAATCGGCAAAATCCCTTATAAATCAAA
AGAATAGACCGAGATAGGGTTGAGTGTTGTTCCAGTTTGGAACAAGAGTCCACTATT
AAAGAACGTGGACTCCAACGTCAAAGGGCGAAAAACCGTCTATCAGGGCGATGGCC
CACTACGTGAACCATCACCCTAATCAAGTTTTTTGGGGTCGAGGTGCCGTAAAGCAC
TAAATCGGAACCCTAAAGGGAGCCCCCGATTTAGAGCTTGACGGGGAAAGCCGGCG
AACGTGGCGAGAAAGGAAGGGAAGAAAGCGAAAGGAGCGGGCGCTAGGGCGCTGG
CAAGTGTAGCGGTCACGCTGCGCGTAACCACCACACCCGCCGCGCTTAATGCGCCG
CTACAGGGCGCGTCCATTCGCCATTCAGGCTGCGCAACTGTTGGGAAGGGCGATCG

GTGCGGGCCTCTTCGCTATTACGCCAGCTGGCGAAAGGGGGATGTGCTGCAAGGCG
ATTAAGTTGGGTAACGCCAGGGTTTTCCCAGTCACGACGTTGTAAAACGACGGCCAG
TGAATTGTAATACGACTCACTATAGGGCGAATTGGAGCTCCACCGCGGTGGCGGCC
GCTCTAGACAATACGAAGGAACGATCCCGCGTCAGCAGATGTCACAATTTGATTATT
TCATTGTCACATGCTTTTTGACTCATCCTTCCGCGAAAGGAGAACTTTTTACAATGCC
GCCGGCTCTGTTAGCGTACCCTTCTACAAGCAGATAGCAGAACAAACACATGATATA
TTCAAAAGGTGCAATGTCAGAGAACATATATTGCGCCCCTGTCCTGTAGACATCAGT
CATTTTTCGCGGGACTTGAATGGCGCACCATTATCCTATACAGAAAACATTAAATTT
ACTGCAAAATTTTGGGCAAACGCTTGGATAGACTATGAATCCGTCATGAAGACACTA
CTTGTAAAATTATATATATCCTTTTTTTTCATGTGCTAGTAAGAATTAGCAACTACTT
TCATTCATCTTCGACGCAACTTCGAGGAAAGCACCTTTTTACTTGCTGTAGCTCTATT
CTCTTCCCCAACCACCTTTTCCTTTATTCCAAAATTTTTAAAACTTTTTCTGTTACATT
ATATAATCTTCTGTCTGAAATGTTTGGATATAACGCCTCTTGATCCACTTTGTATATG
CGTGCTATTTATTTTCAGATTTATAAAGAGTATGAGCGTCATTTACATAAATAGCTG
AAGTTATTCATGGAAAATACGAAGAGCACGTATGTGAGCCAACAGAACATTTGACG
TAAGACTCTACAATGTGCCAAGAACTGGACAAGTAGAGGACTGAGAACTTTATTTC
AATTCATTGCTCCTTTTTGGTGGCGCTACCTTTAGCGAAGGTCAATGATGAATGTGC
ACATGCTGTCGAAACCAAAAAGCAAATTCTAACCAACTTCAAAATGACATAGTCAT
CTGATATTTCTACTCATTATAGATAGTATGGGAGCCTTGAAACGAAAAGTAAGTAAA
AAGCTGGATATGAGCAGTATGAGGTAGACCTTAGCTACATCATTTCCCCCAATAGCT
GCTGCAAATATCTGGTTAAATTTGTGATTCCATGAAGAGGATAACAGACTTGTTAAA
AAGCATCCTGTCAAAATCTAATTTTTGAAGGGCAGTATTCAATTCATAATTTACTTTA
GCTTAGATCCTTCCAATTTATACATGGTATTATAACCAGATCATAAACTACAATCTGT
CGCTACCGCATGTACGAAGAATACTTAAGTCACTTGCTCTCTCATCATTTGTACATTT
TTCAGTAATACCGTTTGATAGCGCCGTCTTTATTACCCGGATTATTCCGTTGACCCTG
AATGAAAAATTTTTCAGAAATCCAGTGCTAAGCGTCAAATCAATGAAATACATCAC
TGTATTTTTAACTGATATACTGTTGGTGTGGCCTTAACGACACCTTTATTTCTTAATT
CATTTCGGCTTGTCTCCTTATTAAGACTACAGAAATAGACAAAGGAAATACTTCAAT
AATGGATAACGAGGTTGAAAAAAATATTGAGATCTGGAAGGTCAAGAAGTTGGTCC
AATCTTTAGAAAAAGCTAGAGGTAATGGTACTTCTATGATTTCCTTAGTTATTCCTCC
TAAGGGTCAAATTCCACTGTACCAAAAAATGTTAACAGATGAATATGGTACTGCCTC
GAATATTAAA**C**CTAGGGTTAATCGTCTTTCCGTTTTATCTGCTATCACTTCCACCCAA
CAAAAGTTGAAGCTATATAATACT**C**TGCCCAAGAACGGTTTAGTTTTATA**C**TGTGGT
GATATCATCACTGAAGATGGTAAAGAAAAAAAGGTCACTTTTGACATCGAACCTTA
CAAACCTATCAACACATCCTTATAT**C**TGTGTGATAACAAATTTCATACAGAAGTTCT
TTCGGAATTGCTTCAAGCTGACG**G**CAAGTTCGGTTTTATAGTCATGGACGGTCAAGG
TACTTTGTTTGGTTCTGTGTCCGGTAATACGAGAACTGTTTTACATAAATTTACTGTC
GATCTGCCAAAAAAGCATGGTAGAGGTGGTCAATCTGCGCTTCGTTTTGCTCGTTTA
AGAGAAGAAAAAAGACATAATTATGTGAGAAAGGTC**A**CCGAAGTTGCTGTTCA**G**AA
TTTTATTACTAATGACAAAGTCAATGTTAAGGGTTTAATTTTAGCTGGTTCTGCTGAC
TTTAAGACCGATTTGGCTAAATCTGAATTATTCGATCCAAGACTAGCATGTAAGGTT
ATTTCCATCGTGGATGTTTCTTATGGTGGTGAAAACGGTTTCAACCAGGCTATCGAA
CTTTCTGCCGAAGCGTTGGCCAATGTCAAGTATGTTCAAGAAAAGAAATTATTGGAG
GCATATTTTGACGAAATTTCCCAGGACACTGGTAAATTCTGTTATGGTATAGATGAT
ACTTTAAAGGCATTGGATTTAGGTGCAGTCGAAAAATTAATTGTTTTCGAAAATTTG
GAAACTATCAGATATACATTTAAAGATGCCGAGGATAATGAGGTTATAAAATTCGCT

GAACCAGAAGCCAAGGACAAGTCGTTTGCTATTGACAAAGCTACCGGCCAAGAAAT
GGACGTTGTCTCCGAAGAACCTTTAATTGAATGGCTAGCAGCTAACTACAAAAACTT
CGGTGCTACCTTGGAATTCATCACAGACAAATCTTCAGAAGGTGCCCAATTTGTCAC
AGGTTTTGGTGGTATTGGTGCCATGCTGCGTTACAAAGTTAATTTTGAACAACTAGT
TGATGAATCTGAGGATGAATATTATGACGAAGATGAAGGATCCGACTATGATTTCAT
TTAAATAAATAAAAGGGGGAGAAAAAAATCGAATCAAAAAGAATTTAATCACTAGA
TGCCAGATTTAAATTAAATTCGCTTTTAATTTTTTGTACAATATAATATATACTTGGT
AAACCTTTTGCTCTATATTGAGCTAATTCCTTTGTTGAAAGTACATAGTGGGTTTAGA
GGACCGTGTATATTACGTAGAAAATACAGTGAAAGGAGAGTTTCTCTTCAAAAGCCT
CGACGGTATCGATAAGCTTATCGATACCGTCGACCTCGAGGGGGGGCCCGGTACCA
GCTTTTGTTCCCTTTAGTGAGGGTTAATTTCGAGCTTGGCGTAATCATGGTCATAGCT
GTTTCCTGTGTGAAATTGTTATCCGCTCACAATTCCACACAACATACGAGCCGGAAG
CATAAAGTGTAAAGCCTGGGGTGCCTAATGAGTGAGCTAACTCACATTAATTGCGTT
GCGCTCACTGCCCGCTTTCCAGTCGGGAAACCTGTCGTGCCAGCTGCATTAATGAAT
CGGCCAACGCGCGGGGAGAGGCGGTTTGCGTATTGGGCGCTCTTCCGCTTCCTCGCT
CACTGACTCGCTGCGCTCGGTCGTTCGGCTGCGGCGAGCGGTATCAGCTCACTCAAA
GGCGGTAATACGGTTATCCACAGAATCAGGGGATAACGCAGGAAAGAACATGTGAG
CAAAAGGCCAGCAAAAGGCCAGGAACCGTAAAAAGGCCGCGTTGCTGGCGTTTTTC
CATAGGCTCCGCCCCCCTGACGAGCATCACAAAAATCGACGCTCAAGTCAGAGGTG
GCGAAACCCGACAGGACTATAAAGATACCAGGCGTTTCCCCCTGGAAGCTCCCTCG
TGCGCTCTCCTGTTCCGACCCTGCCGCTTACCGGATACCTGTCCGCCTTTCTCCCTTC
GGGAAGCGTGGCGCTTTCTCATAGCTCACGCTGTAGGTATCTCAGTTCGGTGTAGGT
CGTTCGCTCCAAGCTGGGCTGTGTGCACGAACCCCCCGTTCAGCCCGACCGCTGCGC
CTTATCCGGTAACTATCGTCTTGAGTCCAACCCGGTAAGACACGACTTATCGCCACT
GGCAGCAGCCACTGGTAACAGGATTAGCAGAGCGAGGTATGTAGGCGGTGCTACAG
AGTTCTTGAAGTGGTGGCCTAACTACGGCTACACTAGAAGAACAGTATTTGGTATCT
GCGCTCTGCTGAAGCCAGTTACCTTCGGAAAAAGAGTTGGTAGCTCTTGATCCGGCA
AACAAACCACCGCTGGTAGCGGTGGTTTTTTTGTTTGCAAGCAGCAGATTACGCGCA
GAAAAAAAGGATCTCAAGAAGATCCTTTGATCTTTTCTACGGGGTCTGACGCTCAGT
GGAACGAAAACTCACGTTAAGGGATTTTGGTCATGAGATTATCAAAAAGGATCTTC
ACCTAGATCCTTTTAAATTAAAAATGAAGTTTTAAATCAATCTAAAGTATATATGAG
TAAACTTGGTCTGACAGTTACCAATGCTTAATCAGTGAGGCACCTATCTCAGCGATC
TGTCTATTTCGTTCATCCATAGTTGCCTGACTCCCCGTCGTGTAGATAACTACGATAC
GGGAGGGCTTACCATCTGGCCCCAGTGCTGCAATGATACCGCGAGACCCACGCTCA
CCGGCTCCAGATTTATCAGCAATAAACCAGCCAGCCGGAAGGGCCGAGCGCAGAAG
TGGTCCTGCAACTTTATCCGCCTCCATCCAGTCTATTAATTGTTGCCGGGAAGCTAGA
GTAAGTAGTTCGCCAGTTAATAGTTTGCGCAACGTTGTTGCCATTGCTACAGGCATC
GTGGTGTCACGCTCGTCGTTTGGTATGGCTTCATTCAGCTCCGGTTCCCAACGATCAA
GGCGAGTTACATGATCCCCCATGTTGTGCAAAAAAGCGGTTAGCTCCTTCGGTCCTC
CGATCGTTGTCAGAAGTAAGTTGGCCGCAGTGTTATCACTCATGGTTATGGCAGCAC
TGCATAATTCTCTTACTGTCATGCCATCCGTAAGATGCTTTTCTGTGACTGGTGAGTA
CTCAACCAAGTCATTCTGAGAATAGTGTATGCGGCGACCGAGTTGCT

**PTH353-SUP45-CR3T.4**

*SUP45* Promoter *SUP45-**CR3T.4***

GCTAAATCTGAATTATTCGATCCAAGACTAGCATGTAAGGTTATTTCCATCGTGGAT
GTTTCTTATGGTGGTGAAAACGGTTTCAACCAGGCTATCGAACTTTCTGCCGAAGCG
TTGGCCAATGTCAAGTATGTTCAAGAAAAGAAATTATTGGAGGCATATTTTGACGAA
ATTTCCCAGGACACTGGTAAA**C**TCTGTTATGGTATAGATGATACTTTAAAGGCATTG
GATTTAGGTGCAGTCGAAAAATTAATTGTTTTCGAAAATTTGGAAACTATCAGATAT
ACATTTAAAGATGCCGAGGATAATGAGGTTATAAAATTCGCTGAACCAGAAGCCAA
GGACAAGTCGTTTGCTATTGACAAAGCTACCGGCCAAGAAATGGACGTTGTCTCCGA
AGAACCTTTAATTGAATGGCTAGCAGCTAACTACAAAAACTTCGGTGCTACCTTGGA
ATTCATCACAGACAAATCTTCAGAAGGTGCCCAATTTGTCACAGGTTTTGGTGGTAT
TGGTGCCATGCTGCGTTACAAAGTTAATTTTGAACAACTAGTTGATGAATCTGAGGA
TGAATATTATGACGAAGATGAAGGATCCGACTATGATTTCATTTAAATAAATAAAAG
GGGGAGAAAAAAATCGAATCAAAAAGAATTTAATCACTAGATGCCAGATTTAAATT
AAATTCGCTTTTAATTTTTTGTACAATATAATATATACTTGGTAAACCTTTTGCTCTA
TATTGAGCTAATTCCTTTGTTGAAAGTACATAGTGGGTTTAGAGGACCGTGTATATT
ACGTAGAAAATACAGTGAAAGGAGAGTTTCTCTTCAAAAGCCTCGACGGTATCGAT
AAGCTTATCGATACCGTCGACCTCGAGGGGGGGCCCGGTACCAGCTTTTGTTCCCTT
TAGTGAGGGTTAATTTCGAGCTTGGCGTAATCATGGTCATAGCTGTTTCCTGTGTGA
AATTGTTATCCGCTCACAATTCCACACAACATACGAGCCGGAAGCATAAAGTGTAA
AGCCTGGGGTGCCTAATGAGTGAGCTAACTCACATTAATTGCGTTGCGCTCACTGCC
CGCTTTCCAGTCGGGAAACCTGTCGTGCCAGCTGCATTAATGAATCGGCCAACGCGC
GGGGAGAGGCGGTTTGCGTATTGGGCGCTCTTCCGCTTCCTCGCTCACTGACTCGCT
GCGCTCGGTCGTTCGGCTGCGGCGAGCGGTATCAGCTCACTCAAAGGCGGTAATAC
GGTTATCCACAGAATCAGGGGATAACGCAGGAAAGAACATGTGAGCAAAAGGCCA
GCAAAAGGCCAGGAACCGTAAAAAGGCCGCGTTGCTGGCGTTTTTCCATAGGCTCC
GCCCCCCTGACGAGCATCACAAAAATCGACGCTCAAGTCAGAGGTGGCGAAACCCG
ACAGGACTATAAAGATACCAGGCGTTTCCCCCTGGAAGCTCCCTCGTGCGCTCTCCT
GTTCCGACCCTGCCGCTTACCGGATACCTGTCCGCCTTTCTCCCTTCGGGAAGCGTG
GCGCTTTCTCATAGCTCACGCTGTAGGTATCTCAGTTCGGTGTAGGTCGTTCGCTCCA
AGCTGGGCTGTGTGCACGAACCCCCCGTTCAGCCCGACCGCTGCGCCTTATCCGGTA
ACTATCGTCTTGAGTCCAACCCGGTAAGACACGACTTATCGCCACTGGCAGCAGCCA
CTGGTAACAGGATTAGCAGAGCGAGGTATGTAGGCGGTGCTACAGAGTTCTTGAAG
TGGTGGCCTAACTACGGCTACACTAGAAGAACAGTATTTGGTATCTGCGCTCTGCTG
AAGCCAGTTACCTTCGGAAAAAGAGTTGGTAGCTCTTGATCCGGCAAACAAACCAC
CGCTGGTAGCGGTGGTTTTTTTGTTTGCAAGCAGCAGATTACGCGCAGAAAAAAAGG
ATCTCAAGAAGATCCTTTGATCTTTTCTACGGGGTCTGACGCTCAGTGGAACGAAAA
CTCACGTTAAGGGATTTTGGTCATGAGATTATCAAAAAGGATCTTCACCTAGATCCT
TTTAAATTAAAAATGAAGTTTTAAATCAATCTAAAGTATATATGAGTAAACTTGGTC
TGACAGTTACCAATGCTTAATCAGTGAGGCACCTATCTCAGCGATCTGTCTATTTCGT
TCATCCATAGTTGCCTGACTCCCCGTCGTGTAGATAACTACGATACGGGAGGGCTTA
CCATCTGGCCCCAGTGCTGCAATGATACCGCGAGACCCACGCTCACCGGCTCCAGAT
TTATCAGCAATAAACCAGCCAGCCGGAAGGGCCGAGCGCAGAAGTGGTCCTGCAAC
TTTATCCGCCTCCATCCAGTCTATTAATTGTTGCCGGGAAGCTAGAGTAAGTAGTTC
GCCAGTTAATAGTTTGCGCAACGTTGTTGCCATTGCTACAGGCATCGTGGTGTCACG
CTCGTCGTTTGGTATGGCTTCATTCAGCTCCGGTTCCCAACGATCAAGGCGAGTTAC
ATGATCCCCCATGTTGTGCAAAAAAGCGGTTAGCTCCTTCGGTCCTCCGATCGTTGT
CAGAAGTAAGTTGGCCGCAGTGTTATCACTCATGGTTATGGCAGCACTGCATAATTC

TCTTACTGTCATGCCATCCGTAAGATGCTTTTCTGTGACTGGTGAGTACTCAACCAAG
TCATTCTGAGAATAGTGTATGCGGCGACCGAGTTGCTCTTGCCCGGCGTCAATACGG
GATAATACCGCGCCACATAGCAGAACTTTAAAAGTGCTCATCATTGGAAAACGTTCT
TCGGGGCGAAAACTCTCAAGGATCTTACCGCTGTTGAGATCCAGTTCGATGTAACCC
ACTCAGTGCACCCAACTGATCTTCAGCATCTTTTACTTTCACCAGCGTTTCTGGGTGA
GCAAAAACAGGAAGGCAAAATGCCGCAAAAAAGGGAATAAGGGCGACACGGAAAT
GTTGAATACTCATACTCTTCCTTTTTCAATATTATTGAAGCATTTATCAGGGTTATTG
TCTCATGAGCGGATACATATTTGAATGTATTTAGAAAAATAAACAAATAGGGGTTCC
GCGCACATTTCCCCGAAAAGTGCCACCTGGGTCCTTTTCATCACGTGCTATAAAAAT
AATTATAATTTAAATTTTTTAATATAAATATATAAATTAAAAATAGAAAGTAAAAAA
AGAAATTAAAGAAAAAATAGTTTTTGTTTTCCGAAGATGTAAAAGACTCTAGGGGG
ATCGCCAACAAATACTACCTTTTATCTTGCTCTTCCTGCTCTCAGGTATTAATGCCGA
ATTGTTTCATCTTGTCTGTGTAGAAGACCACACACGAAAATCCTGTGATTTTACATTT
TACTTATCGTTAATCGAATGTATATCTATTTAATCTGCTTTTCTTGTCTAATAAATAT
ATATGTAAAGTACGCTTTTTGTTGAAATTTTTTAAACCTTTGTTTATTTTTTTTTCTTCA
TTCCGTAACTCTTCTACCTTCTTTATTTACTTTCTAAAATCCAAATACAAAACATAAA
AATAAATAAACACAGAGTAAATTCCCAAATTATTCCATCATTAAAAGATACGAGGC
GCGTGTAAGTTACAGGCAAGCGATCCGTCCTAAGAAACCATTATTATCATGACATTA
ACCTATAAAAATAGGCGTATCACGAGGCCCTTTCGTCTCGCGCGTTTCGGTGATGAC
GGTGAAAACCTCTGACACATGCAGCTCCCGGAGACGGTCACAGCTTGTCTGTAAGC
GGATGCCGGGAGCAGACAAGCCCGTCAGGGCGCGTCAGCGGGTGTTGGCGGGTGTC
GGGGCTGGCTTAACTATGCGGCATCAGAGCAGATTGTACTGAGAGTGCACCATATC
GACTACGTCGTTAAGGCCGTTTCTGACAGAGTAAAATTCTTGAGGGAACTTTCACCA
TTATGGGAAATGGTTCAAGAAGGTATTGACTTAAACTCCATCAAATGGTCAGGTCAT
TGAGTGTTTTTTATTTGTTGTATTTTTTTTTTTTTAGAGAAAATCCTCCAATATATAAA
TTAGGAATCATAGTTTCATGATTTTCTGTTACACCTAACTTTTTGTGTGGTGCCCTCC
TCCTTGTCAATATTAATGTTAAAGTGCAATTCTTTTTCCTTATCACGTTGAGCCATTA
GTATCAATTTGCTTACCTGTATTCCTTTACATCCTCCTTTTTCTCCTTCTTGATAAATG
TATGTAGATTGCGTATATAGTTTCGTCTACCCTATGAACATATTCCATTTTGTAATTT
CGTGTCGTTTCTATTATGAATTTCATTTATAAAGTTTATGTACAAATATCATAAAAAA
AGAGAATCTTTTTAAGCAAGGATTTTCTTAACTTCTTCGGCGACAGCATCACCGACT
TCGGTGGTACTGTTGGAACCACCTAAATCACCAGTTCTGATACCTGCATCCAAAACC
TTTTTAACTGCATCTTCAATGGCCTTACCTTCTTCAGGCAAGTTCAATGACAATTTCA
ACATCATTGCAGCAGACAAGATAGTGGCGATAGGGTTGACCTTATTCTTTGGCAAAT
CTGGAGCAGAACCGTGGCATGGTTCGTACAAACCAAATGCGGTGTTCTTGTCTGGCA
AAGAGGCCAAGGACGCAGATGGCAACAAACCCAAGGAACCTGGGATAACGGAGGC
TTCATCGGAGATGATATCACCAAACATGTTGCTGGTGATTATAATACCATTTAGGTG
GGTTGGGTTCTTAACTAGGATCATGGCGGCAGAATCAATCAATTGATGTTGAACCTT
CAATGTAGGGAATTCGTTCTTGATGGTTTCCTCCACAGTTTTTCTCCATAATCTTGAA
GAGGCCAAAACATTAGCTTTATCCAAGGACCAAATAGGCAATGGTGGCTCATGTTGT
AGGGCCATGAAAGCGGCCATTCTTGTGATTCTTTGCACTTCTGGAACGGTGTATTGT
TCACTATCCCAAGCGACACCATCACCATCGTCTTCCTTTCTCTTACCAAAGTAAATAC
CTCCCACTAATTCTCTGACAACAACGAAGTCAGTACCTTTAGCAAATTGTGGCTTGA
TTGGAGATAAGTCTAAAAGAGAGTCGGATGCAAAGTTACATGGTCTTAAGTTGGCG
TACAATTGAAGTTCTTTACGGATTTTTAGTAAACCTTGTTCAGGTCTAACACTACCGG
TACCCCATTTAGGACCACCCACAGCACCTAACAAAACGGCATCAGCCTTCTTGGAGG

CTTCCAGCGCCTCATCTGGAAGTGGAACACCTGTAGCATCGATAGCAGCACCACCA
ATTAAATGATTTTCGAAATCGAACTTGACATTGGAACGAACATCAGAAATAGCTTTA
AGAACCTTAATGGCTTCGGCTGTGATTTCTTGACCAACGTGGTCACCTGGCAAAACG
ACGATCTTCTTAGGGGCAGACATTAGAATGGTATATCCTTGAAATATATATATATAT
ATTGCTGAAATGTAAAAGGTAAGAAAAGTTAGAAAGTAAGACGATTGCTAACCACC
TATTGGAAAAAACAATAGGTCCTTAAATAATATTGTCAACTTCAAGTATTGTGATGC
AAGCATTTAGTCATGAACGCTTCTCTATTCTATATGAAAAGCCGGTTCCGGCGCTCT
CACCTTTCCTTTTTCTCCCAATTTTTCAGTTGAAAAAGGTATATGCGTCAGGCGACCT
CTGAAATTAACAAAAAATTTCCAGTCATCGAATTTGATTCTGTGCGATAGCGCCCCT
GTGTGTTCTCGTTATGTTGAGGAAAAAAATAATGGTTGCTAAGAGATTCGAACTCTT
GCATCTTACGATACCTGAGTATTCCCACAGTTAACTGCGGTCAAGATATTTCTTGAA
TCAGGCGCCTTAGACCGCTCGGCCAAACAACCAATTACTTGTTGAGAAATAGAGTAT
AATTATCCTATAAATATAACGTTTTTGAACACACATGAACAAGGAAGTACAGGACA
ATTGATTTTGAAGAGAATGTGGATTTTGATGTAATTGTTGGGATTCCATTTTTAATAA
GGCAATAATATTAGGTATGTAGATATACTAGAAGTTCTCCTCGACCGGTCGATATGC
GGTGTGAAATACCGCACAGATGCGTAAGGAGAAAATACCGCATCAGGAAATTGTAA
GCGTTAATATTTTGTTAAAATTCGCGTTAAATTTTTGTTAAATCAGCTCATTTTTTAA
CCAATAGGCCGAAATCGGCAAAATCCCTTATAAATCAAAAGAATAGACCGAGATAG
GGTTGAGTGTTGTTCCAGTTTGGAACAAGAGTCCACTATTAAAGAACGTGGACTCCA
ACGTCAAAGGGCGAAAAACCGTCTATCAGGGCGATGGCCCACTACGTGAACCATCA
CCCTAATCAAGTTTTTTGGGGTCGAGGTGCCGTAAAGCACTAAATCGGAACCCTAAA
GGGAGCCCCCGATTTAGAGCTTGACGGGGAAAGCCGGCGAACGTGGCGAGAAAGG
AAGGGAAGAAAGCGAAAGGAGCGGGCGCTAGGGCGCTGGCAAGTGTAGCGGTCAC
GCTGCGCGTAACCACCACACCCGCCGCGCTTAATGCGCCGCTACAGGGCGCGTCCAT
TCGCCATTCAGGCTGCGCAACTGTTGGGAAGGGCGATCGGTGCGGGCCTCTTCGCTA
TTACGCCAGCTGGCGAAAGGGGGATGTGCTGCAAGGCGATTAAGTTGGGTAACGCC
AGGGTTTTCCCAGTCACGACGTTGTAAAACGACGGCCAGTGAATTGTAATACGACTC
ACTATAGGGCGAATTGGAGCTCCACCGCGGTGGCGGCCGCTCTAGACAATACGAAG
GAACGATCCCGCGTCAGCAGATGTCACAATTTGATTATTTCATTGTCACATGCTTTTT
GACTCATCCTTCCGCGAAAGGAGAACTTTTTACAATGCCGCCGGCTCTGTTAGCGTA
CCCTTCTACAAGCAGATAGCAGAACAAACACATGATATATTCAAAAGGTGCAATGT
CAGAGAACATATATTGCGCCCCTGTCCTGTAGACATCAGTCATTTTTCGCGGGACTT
GAATGGCGCACCATTATCCTATACAGAAAACATTAAATTTACTGCAAAATTTTGGGC
AAACGCTTGGATAGACTATGAATCCGTCATGAAGACACTACTTGTAAAATTATATAT
ATCCTTTTTTTTCATGTGCTAGTAAGAATTAGCAACTACTTTCATTCATCTTCGACGC
AACTTCGAGGAAAGCACCTTTTTACTTGCTGTAGCTCTATTCTCTTCCCCAACCACCT
TTTCCTTTATTCCAAAATTTTTAAAACTTTTTCTGTTACATTATATAATCTTCTGTCTG
AAATGTTTGGATATAACGCCTCTTGATCCACTTTGTATATGCGTGCTATTTATTTTCA
GATTTATAAAGAGTATGAGCGTCATTTACATAAATAGCTGAAGTTATTCATGGAAAA
TACGAAGAGCACGTATGTGAGCCAACAGAACATTTGACGTAAGACTCTACAATGTG
CCAAGAACTGGACAAGTAGAGGACTGAGAACTTTATTTCAATTCATTGCTCCTTTTT
GGTGGCGCTACCTTTAGCGAAGGTCAATGATGAATGTGCACATGCTGTCGAAACCA
AAAAGCAAATTCTAACCAACTTCAAAATGACATAGTCATCTGATATTTCTACTCATT
ATAGATAGTATGGGAGCCTTGAAACGAAAAGTAAGTAAAAAGCTGGATATGAGCAG
TATGAGGTAGACCTTAGCTACATCATTTCCCCCAATAGCTGCTGCAAATATCTGGTT
AAATTTGTGATTCCATGAAGAGGATAACAGACTTGTTAAAAAGCATCCTGTCAAAAT

CTAATTTTTGAAGGGCAGTATTCAATTCATAATTTACTTTAGCTTAGATCCTTCCAAT
TTATACATGGTATTATAACCAGATCATAAACTACAATCTGTCGCTACCGCATGTACG
AAGAATACTTAAGTCACTTGCTCTCTCATCATTTGTACATTTTTCAGTAATACCGTTT
GATAGCGCCGTCTTTATTACCCGGATTATTCCGTTGACCCTGAATGAAAAATTTTTC
AGAAATCCAGTGCTAAGCGTCAAATCAATGAAATACATCACTGTATTTTTAACTGAT
ATACTGTTGGTGTGGCCTTAACGACACCTTTATTTCTTAATTCATTTCGGCTTGTCTC
CTTATTAAGACTACAGAAATAGACAAAGGAAATACTTCAATAATGGATAACGAGGT
TGAAAAAAATATTGAGATCTGGAAGGTCAAGAAGTTGGTCCAATCTTTAGAAAAAG
CTAGAGGTAATGGTACTTCTATGATTTCCTTAGTTATTCCTCCTAAGGGTCAAATTCC
ACTGTACCAAAAAATGTTAACAGATGAATATGGTACTGCCTCGAATATTAAATCTAG
GGTTAATCGTCTTTCCGTTTTATCTGCTATCACTTCCACCCAACAAAAGTTGAAGCTA
TATAATACTTTGCCCAAGAACGGTTTAGTTTTATA**C**TGTGGTGATATCATCACTGAA
GATGGTAAAGAAAAAAAGGTCACTTTTGACATCGAACCTTACAA**G**CCTATCAACAC
ATCCTTATATTTG**C**GTGATAACAAATTTCATACAGAAGTTCTTTCGGAATTGCTTCAA
GCTGACGACAAGTTCGGTTTTATAGTCATGGACGGTCAAGGTACTTTGTTTGGTTCT
GTGTCCGGTAATACGAGAACTGTTTTACATAAATTTACTGTCGATCTGCCAAAAAAG
CATGGTAGAGGTGGTCAATCTGCGCTTCGTTTTGCTCGTTTAAGAGAAGAA**G**AAAGA
CATAATTATGTGAGAAAGGTCGC**T**GAAGTTGCTGTTCAAAATTTTATTACTAATGAC
AAAGTCAATGTTAAGGGTTTAATTTTAGCTGGTTCTGCTGACTTTAAGACCGATTTG

**PTH353-SUP45-CR3T.6**

*SUP45* Promoter *SUP45*-***CR3T.6***

GCGTCAATACGGGATAATACCGCGCCACATAGCAGAACTTTAAAAGTGCTCATCATT
GGAAAACGTTCTTCGGGGCGAAAACTCTCAAGGATCTTACCGCTGTTGAGATCCAGT
TCGATGTAACCCACTCGTGCACCCAACTGATCTTCAGCATCTTTTACTTTCACCAGCG
TTTCTGGGTGAGCAAAAACAGGAAGGCAAAATGCCGCAAAAAAGGGAATAAGGGC
GACACGGAAATGTTGAATACTCATACTCTTCCTTTTTCAATATTATTGAAGCATTTAT
CAGGGTTATTGTCTCATGAGCGGATACATATTTGAATGTATTTAGAAAAATAAACAA
ATAGGGGTTCCGCGCACATTTCCCCGAAAAGTGCCACCTGGGTCCTTTTCATCACGT
GCTATAAAAATAATTATAATTTAAATTTTTTAATATAAATATATAAATTAAAAATAG
AAAGTAAAAAAAGAAATTAAAGAAAAAATAGTTTTTGTTTTCCGAAGATGTAAAAG
ACTCTAGGGGGATCGCCAACAAATACTACCTTTTATCTTGCTCTTCCTGCTCTCAGGT
ATTAATGCCGAATTGTTTCATCTTGTCTGTGTAGAAGACCACACACGAAAATCCTGT
GATTTTACATTTTACTTATCGTTAATCGAATGTATATCTATTTAATCTGCTTTTCTTGT
CTAATAAATATATATGTAAAGTACGCTTTTTGTTGAAATTTTTTAAACCTTTGTTTAT
TTTTTTTCTTCATTCCGTAACTCTTCTACCTTCTTTATTTACTTTCTAAAATCCAAATA
CAAAACATAAAAATAAATAAACACAGAGTAAATTCCCAAATTATTCCATCATTAAA
AGATACGAGGCGCGTGTAAGTTACAGGCAAGCGATCCGTCCTAAGAAACCATTATT
ATCATGACATTAACCTATAAAAATAGGCGTATCACGAGGCCCTTTCGTCTCGCGCGT
TTCGGTGATGACGGTGAAAACCTCTGACACATGCAGCTCCCGGAGACGGTCACAGC
TTGTCTGTAAGCGGATGCCGGGAGCAGACAAGCCCGTCAGGGCGCGTCAGCGGGTG
TTGGCGGGTGTCGGGGCTGGCTTAACTATGCGGCATCAGAGCAGATTGTACTGAGA
GTGCACCATATCGACTACGTCGTTAAGGCCGTTTCTGACAGAGTAAAATTCTTGAGG
GAACTTTCACCATTATGGGAAATGGTTCAAGAAGGTATTGACTTAAACTCCATCAAA
TGGTCAGGTCATTGAGTGTTTTTTATTTGTTGTATTTTTTTTTTTTTAGAGAAAATCCT

CCAATATATAAATTAGGAATCATAGTTTCATGATTTTCTGTTACACCTAACTTTTTGT
GTGGTGCCCTCCTCCTTGTCAATATTAATGTTAAAGTGCAATTCTTTTTCCTTATCAC
GTTGAGCCATTAGTATCAATTTGCTTACCTGTATTCCTTTACATCCTCCTTTTTCTCCT
TCTTGATAAATGTATGTAGATTGCGTATATAGTTTCGTCTACCCTATGAACATATTCC
ATTTTGTAATTTCGTGTCGTTTCTATTATGAATTTCATTTATAAAGTTTATGTACAAAT
ATCATAAAAAAAGAGAATCTTTTTAAGCAAGGATTTTCTTAACTTCTTCGGCGACAG
CATCACCGACTTCGGTGGTACTGTTGGAACCACCTAAATCACCAGTTCTGATACCTG
CATCCAAAACCTTTTTAACTGCATCTTCAATGGCCTTACCTTCTTCAGGCAAGTTCAA
TGACAATTTCAACATCATTGCAGCAGACAAGATAGTGGCGATAGGGTTGACCTTATT
CTTTGGCAAATCTGGAGCAGAACCGTGGCATGGTTCGTACAAACCAAATGCGGTGTT
CTTGTCTGGCAAAGAGGCCAAGGACGCAGATGGCAACAAACCCAAGGAACCTGGG
ATAACGGAGGCTTCATCGGAGATGATATCACCAAACATGTTGCTGGTGATTATAATA
CCATTTAGGTGGGTTGGGTTCTTAACTAGGATCATGGCGGCAGAATCAATCAATTGA
TGTTGAACCTTCAATGTAGGGAATTCGTTCTTGATGGTTTCCTCCACAGTTTTTCTCC
ATAATCTTGAAGAGGCCAAAACATTAGCTTTATCCAAGGACCAAATAGGCAATGGT
GGCTCATGTTGTAGGGCCATGAAAGCGGCCATTCTTGTGATTCTTTGCACTTCTGGA
ACGGTGTATTGTTCACTATCCCAAGCGACACCATCACCATCGTCTTCCTTTCTCTTAC
CAAAGTAAATACCTCCCACTAATTCTCTGACAACAACGAAGTCAGTACCTTTAGCAA
ATTGTGGCTTGATTGGAGATAAGTCTAAAAGAGAGTCGGATGCAAAGTTACATGGT
CTTAAGTTGGCGTACAATTGAAGTTCTTTACGGATTTTTAGTAAACCTTGTTCAGGTC
TAACACTACCGGTACCCCATTTAGGACCACCCACAGCACCTAACAAAACGGCATCA
GCCTTCTTGGAGGCTTCCAGCGCCTCATCTGGAAGTGGAACACCTGTAGCATCGATA
GCAGCACCACCAATTAAATGATTTTCGAAATCGAACTTGACATTGGAACGAACATCA
GAAATAGCTTTAAGAACCTTAATGGCTTCGGCTGTGATTTCTTGACCAACGTGGTCA
CCTGGCAAAACGACGATCTTCTTAGGGGCAGACATTAGAATGGTATATCCTTGAAAT
ATATATATATATATTGCTGAAATGTAAAAGGTAAGAAAAGTTAGAAAGTAAGACGA
TTGCTAACCACCTATTGGAAAAAACAATAGGTCCTTAAATAATATTGTCAACTTCAA
GTATTGTGATGCAAGCATTTAGTCATGAACGCTTCTCTATTCTATATGAAAAGCCGG
TTCCGGCGCTCTCACCTTTCCTTTTTCTCCCAATTTTTCAGTTGAAAAAGGTATATGC
GTCAGGCGACCTCTGAAATTAACAAAAAATTTCCAGTCATCGAATTTGATTCTGTGC
GATAGCGCCCCTGTGTGTTCTCGTTATGTTGAGGAAAAAAATAATGGTTGCTAAGAG
ATTCGAACTCTTGCATCTTACGATACCTGAGTATTCCCACAGTTAACTGCGGTCAAG
ATATTTCTTGAATCAGGCGCCTTAGACCGCTCGGCCAAACAACCAATTACTTGTTGA
GAAATAGAGTATAATTATCCTATAAATATAACGTTTTTGAACACACATGAACAAGGA
AGTACAGGACAATTGATTTTGAAGAGAATGTGGATTTTGATGTAATTGTTGGGATTC
CATTTTTAATAAGGCAATAATATTAGGTATGTAGATATACTAGAAGTTCTCCTCGAC
CGGTCGATATGCGGTGTGAAATACCGCACAGATGCGTAAGGAGAAAATACCGCATC
AGGAAATTGTAAGCGTTAATATTTTGTTAAAATTCGCGTTAAATTTTTGTTAAATCAG
CTCATTTTTTAACCAATAGGCCGAAATCGGCAAAATCCCTTATAAATCAAAAGAATA
GACCGAGATAGGGTTGAGTGTTGTTCCAGTTTGGAACAAGAGTCCACTATTAAAGA
ACGTGGACTCCAACGTCAAAGGGCGAAAAACCGTCTATCAGGGCGATGGCCCACTA
CGTGAACCATCACCCTAATCAAGTTTTTTGGGGTCGAGGTGCCGTAAAGCACTAAAT
CGGAACCCTAAAGGGAGCCCCCGATTTAGAGCTTGACGGGGAAAGCCGGCGAACGT
GGCGAGAAAGGAAGGGAAGAAAGCGAAAGGAGCGGGCGCTAGGGCGCTGGCAAGT
GTAGCGGTCACGCTGCGCGTAACCACCACACCCGCCGCGCTTAATGCGCCGCTACA
GGGCGCGTCCATTCGCCATTCAGGCTGCGCAACTGTTGGGAAGGGCGATCGGTGCG

GGCCTCTTCGCTATTACGCCAGCTGGCGAAAGGGGGATGTGCTGCAAGGCGATTA
AGTTGGGTAACGCCAGGGTTTTCCCAGTCACGACGTTGTAAAACGACGGCCAGTGA
ATTGTAATACGACTCACTATAGGGCGAATTGGAGCTCCACCGCGGTGGCGGCCGCTC
TAGACAATACGAAGGAACGATCCCGCGTCAGCAGATGTCACAATTTGATTATTTCAT
TGTCACATGCTTTTTGACTCATCCTTCCGCGAAAGGAGAACTTTTTACAATGCCGCCG
GCTCTGTTAGCGTACCCTTCTACAAGCAGATAGCAGAACAAACACATGATATATTCA
AAAGGTGCAATGTCAGAGAACATATATTGCGCCCCTGTCCTGTAGACATCAGTCATT
TTTCGCGGGACTTGAATGGCGCACCATTATCCTATACAGAAAACATTAAATTTACTG
CAAAATTTTGGGCAAACGCTTGGATAGACTATGAATCCGTCATGAAGACACTACTTG
TAAAATTATATATATCCTTTTTTTTCATGTGCTAGTAAGAATTAGCAACTACTTTCAT
TCATCTTCGACGCAACTTCGAGGAAAGCACCTTTTTACTTGCTGTAGCTCTATTCTCT
TCCCCAACCACCTTTTCCTTTATTCCAAAATTTTTAAAACTTTTTCTGTTACATTATAT
AATCTTCTGTCTGAAATGTTTGGATATAACGCCTCTTGATCCACTTTGTATATGCGTG
CTATTTATTTTCAGATTTATAAAGAGTATGAGCGTCATTTACATAAATAGCTGAAGTT
ATTCATGGAAAATACGAAGAGCACGTATGTGAGCCAACAGAACATTTGACGTAAGA
CTCTACAATGTGCCAAGAACTGGACAAGTAGAGGACTGAGAACTTTATTTCAATTCA
TTGCTCCTTTTTGGTGGCGCTACCTTTAGCGAAGGTCAATGATGAATGTGCACATGCT
GTCGAAACCAAAAAGCAAATTCTAACCAACTTCAAAATGACATAGTCATCTGATATT
TCTACTCATTATAGATAGTATGGGAGCCTTGAAACGAAAAGTAAGTAAAAAGCTGG
ATATGAGCAGTATGAGGTAGACCTTAGCTACATCATTTCCCCCAATAGCTGCTGCAA
ATATCTGGTTAAATTTGTGATTCCATGAAGAGGATAACAGACTTGTTAAAAAGCATC
CTGTCAAAATCTAATTTTTGAAGGGCAGTATTCAATTCATAATTTACTTTAGCTTAGA
TCCTTCCAATTTATACATGGTATTATAACCAGATCATAAACTACAATCTGTCGCTACC
GCATGTACGAAGAATACTTAAGTCACTTGCTCTCTCATCATTTGTACATTTTTCAGTA
ATACCGTTTGATAGCGCCGTCTTTATTACCCGGATTATTCCGTTGACCCTGAATGAAA
AATTTTTTCAGAAATCCAGTGCTAAGCGTCAAATCAATGAAATACATCACTGTATTT
TTAACTGATATACTGTTGGTGTGGCCTTAACGACACCTTTATTTCTTAATTCATTTCG
GCTTGTCTCCTTATTAAGACTACAGAAATAGACAAAGGAAATACTTCAATAATGGAT
AACGAGGTTGAAAAAAATATTGAGATCTGGAAGGTCAAGAAGTTGGTCCAA**C**CTTT
AGAAAAAGCTAGAGGTAATGGTACTTCTATGATTTCCTTAGTTATTCCTCCTAAGGG
TCAAATTCCACTGTACCAAAAAATGTTAACAGATGAATATGGTACTGCCTCGAATAT
TAAATCTAGGGTTAATCGTCTTTCCGTTTTATCTGCTATCACTTCCACCCAACAAAAG
TTGAAGCTATATAATACTTTGCCCAAGAACGGTTTAG_TTTATATTGTGGTGATATCA
TCACTGAAGATGGTAAAGAAAAAAAGGTCACTTTTGACATCGAACCTTACAAACCT
ATCAACACATCCTTATATTTGTGTGATAACAAATTTCATACAGAAGTTCT**C**TCGGAA
TTGCTTCAAGCTGACGACAAGTTCGGTTTTATAGTCATGGACGGTCAAGGTACTTTG
TTTGGTTCTGTGTCCGGTAATACGAGAACTGTTTTACATAAATTTACTGTCGATCTGC
CAAAAAAGCATGGTAGAGGTGGTCAATCTGCGCTTCGTTTTGCTCGTTTA**G**GAGAAG
AAAAAAGACATAATTATGTGAGAAAGGTCGCCGAAGTTGCTGTTCAAAATTTTATTA
CTAATGACAAAGTCAATGTTAAGGGTTTAATTTTAGCTGGTTCTGCTGACTTTAAGA
CCGATTTGGCTAAATCTGAATTATTCGATCCAAGACTAGCATGTAAGGTTATTTCCA
TCGTGGATGTTTCTTA**C**GGTGGTGAAAACGGTTTCAACCAGGCTATCGAACTTTCTG
CCGAAGCGTTGGCCAATGTCAAGTATGTTCAAGAAAAGAA**T**TTATTGGAGGCATATT
TTGACGAAATTTCCCAGGACACTGGTAAATTCTGTTATGGTATAGATGATACTTTAA
AGGCATTGGATTTAGGTGCAGTCGAAAAATTAATTGTTTTCGAAAATTTGGAAACTA
TCAGATATACATTTAAAGATGCCGAGGATAATGAGGTTATAAAATTCGCTGAACCA

GAAGCCAAGGACAAGTCGTTTGCTATTGACAAAGCTACCGGCCAAGAAATGGACGT
TGTCTCCGAAGAACCTTTAATTGAATGGCTAGCAGCTAACTACAAAAACTTCGGTGC
TACCTTGGAATTCATCACAGACAAATCTTCAGAAGGTGCCCAATTTGTCACAGGTTT
TGGTGGTATTGGTGCCATGCTGCGT**C**ACAAAGTTAATTTGAACAACTAGTTGATGA
ATCTGAGGATGAATATTATGACGAAGATGAAGGATCCGACTATGATTTCATTTAAAT
AAATAAAAGGGGGAGAAAAAAATCGAATCAAAAAGAATTTAATCACTAGATGCCA
GATTTAAATTAAATTCGCTTTTAATTTTTTGTACAATATAATATATACTTGGTAAACC
TTTTGCTCTATATTGAGCTAATTCCTTTGTTGAAAGTACATAGTGGGTTTAGAGGACC
GTGTATATTACGTAGAAAATACAGTGAAAGGAGAGTTTCTCTTCAAAAGCCTCGACG
GTATCGATAAGCTTATCGATACCGTCGACCTCGAGGGGGGGCCCGGTACCAGCTTTT
GTTCCCTTTAGTGAGGGTTAATTTCGAGCTTGGCGTAATCATGGTCATAGCTGTTTCC
TGTGTGAAATTGTTATCCGCTCACAATTCCACACAACATACGAGCCGGAAGCATAAA
GTGTAAAGCCTGGGGTGCCTAATGAGTGAGCTAACTCACATTAATTGCGTTGCGCTC
ACTGCCCGCTTTCCAGTCGGGAAACCTGTCGTGCCAGCTGCATTAATGAATCGGCCA
ACGCGCGGGGAGAGGCGGTTTGCGTATTGGGCGCTCTTCCGCTTCCTCGCTCACTGA
CTCGCTGCGCTCGGTCGTTCGGCTGCGGCGAGCGGTATCAGCTCACTCAAAGGCGGT
AATACGGTTATCCACAGAATCAGGGGATAACGCAGGAAAGAACATGTGAGCAAAA
GGCCAGCAAAAGGCCAGGAACCGTAAAAAGGCCGCGTTGCTGGCGTTTTTCCATAG
GCTCCGCCCCCCTGACGAGCATCACAAAAATCGACGCTCAAGTCAGAGGTGGCGAA
ACCCGACAGGACTATAAAGATACCAGGCGTTTCCCCCTGGAAGCTCCCTCGTGCGCT
CTCCTGTTCCGACCCTGCCGCTTACCGGATACCTGTCCGCCTTTCTCCCTTCGGGAAG
CGTGGCGCTTTCTCATAGCTCACGCTGTAGGTATCTCAGTTCGGTGTAGGTCGTTCGC
TCCAAGCTGGGCTGTGTGCACGAACCCCCCGTTCAGCCCGACCGCTGCGCCTTATCC
GGTAACTATCGTCTTGAGTCCAACCCGGTAAGACACGACTTATCGCCACTGGCAGCA
GCCACTGGTAACAGGATTAGCAGAGCGAGGTATGTAGGCGGTGCTACAGAGTTCTT
GAAGTGGTGGCCTAACTACGGCTACACTAGAAGAACAGTATTTGGTATCTGCGCTCT
GCTGAAGCCAGTTACCTTCGGAAAAAGAGTTGGTAGCTCTTGATCCGGCAAACAAA
CCACCGCTGGTAGCGGTGGTTTTTTTGTTTGCAAGCAGCAGATTACGCGCAGAAAAA
AAGGATCTCAAGAAGATCCTTTGATCTTTTCTACGGGGTCTGACGCTCAGTGGAACG
AAAACTCACGTTAAGGGATTTTGGTCATGAGATTATCAAAAAGGATCTTCACCTAGA
TCCTTTTAAATTAAAAATGAAGTTTTAAATCAATCTAAAGTATATATGAGTAAACTT
GGTCTGACAGTTACCAATGCTTAATCAGTGAGGCACCTATCTCAGCGATCTGTCTAT
TTCGTTCATCCATAGTTGCCTGACTCCCCGTCGTGTAGATAACTACGATACGGGAGG
GCTTACCATCTGGCCCCAGTGCTGCAATGATACCGCGAGACCCACGCTCACCGGCTC
CAGATTTATCAGCAATAAACCAGCCAGCCGGAAGGGCCGAGCGCAGAAGTGGTCCT
GCAACTTTATCCGCCTCCATCCAGTCTATTAATTGTTGCCGGGAAGCTAGAGTAAGT
AGTTCGCCAGTTAATAGTTTGCGCAACGTTGTTGCCATTGCTACAGGCATCGTGGTG
TCACGCTCGTCGTTTGGTATGGCTTCATTCAGCTCCGGTTCCCAACGATCAAGGCGA
GTTACATGATCCCCCATGTTGTGCAAAAAAGCGGTTAGCTCCTTCGGTCCTCCGATC
GTTGTCAGAAGTAAGTTGGCCGCAGTGTTATCACTCATGGTTATGGCAGCACTGCAT
AATTCTCTTACTGTCATGCCATCCGTAAGATGCTTTTCTGTGACTGGTGAGTACTCAA
CCAAGTCATTCTGAGAATAGTGTATGCGGCGACCGAGTTGCTCTTGCCCG

**PTH353-SUP45-CR3T.7**

*SUP45* Promoter *SUP45*-***CR3T.7***

CAACTGATCTTCAGCATCTTTTACTTTCACCAGCGTTTCTGGGTGAGCAAAAACAGG
AAGGCAAAATGCCGCAAAAAAGGGAATAAGGGCGACACGGAAATGTTGAATACTC
ATACTCTTCCTTTTTCAATATTATTGAAGCATTTATCAGGGTTATTGTCTCATGAGCG
GATACATATTTGAATGTATTTAGAAAAATAAACAAATAGGGGTTCCGCGCACATTTC
CCCGAAAAGTGCCACCTGGGTCCTTTTCATCACGTGCTATAAAAATAATTATAATTT
AAATTTTTTAATATAAATATATAAATTAAAAATAGAAAGTAAAAAAAGAAATTAAA
GAAAAAATAGTTTTTGTTTTCCGAAGATGTAAAAGACTCTAGGGGGATCGCCAACA
AATACTACCTTTTATCTTGCTCTTCCTGCTCTCAGGTATTAATGCCGAATTGTTTCATC
TTGTCTGTGTAGAAGACCACACACGAAAATCCTGTGATTTTACATTTTACTTATCGTT
AATCGAATGTATATCTATTTAATCTGCTTTTCTTGTCTAATAAATATATATGTAAAGT
ACGCTTTTTGTTGAAATTTTTTAAACCTTTGTTTATTTTTTTTTCTTCATTCCGTAACTCT
TCTACCTTCTTTATTTACTTTCTAAAATCCAAATACAAAACATAAAAATAAATAAAC
ACAGAGTAAATTCCCAAATTATTCCATCATTAAAAGATACGAGGCGCGTGTAAGTTA
CAGGCAAGCGATCCGTCCTAAGAAACCATTATTATCATGACATTAACCTATAAAAAT
AGGCGTATCACGAGGCCCTTTCGTCTCGCGCGTTTCGGTGATGACGGTGAAAACCTC
TGACACATGCAGCTCCCGGAGACGGTCACAGCTTGTCTGTAAGCGGATGCCGGGAG
CAGACAAGCCCGTCAGGGCGCGTCAGCGGGTGTTGGCGGGTGTCGGGGCTGGCTTA
ACTATGCGGCATCAGAGCAGATTGTACTGAGAGTGCACCATATCGACTACGTCGTTA
AGGCCGTTTCTGACAGAGTAAAATTCTTGAGGGAACTTTCACCATTATGGGAAATGG
TTCAAGAAGGTATTGACTTAAACTCCATCAAATGGTCAGGTCATTGAGTGTTTTTTAT
TTGTTGTATTTTTTTTTTTTTTAGAGAAAATCCTCCAATATATAAATTAGGAATCATAG
TTTCATGATTTTCTGTTACACCTAACTTTTTGTGTGGTGCCCTCCTCCTTGTCAATATT
AATGTTAAAGTGCAATTCTTTTTCCTTATCACGTTGAGCCATTAGTATCAATTTGCTT
ACCTGTATTCCTTTACATCCTCCTTTTTCTCCTTCTTGATAAATGTATGTAGATTGCGT
ATATAGTTTCGTCTACCCTATGAACATATTCCATTTTGTAATTTCGTGTCGTTTCTATT
ATGAATTTCATTTATAAAGTTTATGTACAAATATCATAAAAAAAGAGAATCTTTTTA
AGCAAGGATTTTCTTAACTTCTTCGGCGACAGCATCACCGACTTCGGTGGTACTGTT
GGAACCACCTAAATCACCAGTTCTGATACCTGCATCCAAAACCTTTTTAACTGCATC
TTCAATGGCCTTACCTTCTTCAGGCAAGTTCAATGACAATTTCAACATCATTGCAGC
AGACAAGATAGTGGCGATAGGGTTGACCTTATTCTTTGGCAAATCTGGAGCAGAAC
CGTGGCATGGTTCGTACAAACCAAATGCGGTGTTCTTGTCTGGCAAAGAGGCCAAG
GACGCAGATGGCAACAAACCCAAGGAACCTGGGATAACGGAGGCTTCATCGGAGAT
GATATCACCAAACATGTTGCTGGTGATTATAATACCATTTAGGTGGGTTGGGTTCTT
AACTAGGATCATGGCGGCAGAATCAATCAATTGATGTTGAACCTTCAATGTAGGGA
ATTCGTTCTTGATGGTTTCCTCCACAGTTTTTCTCCATAATCTTGAAGAGGCCAAAAC
ATTAGCTTTATCCAAGGACCAAATAGGCAATGGTGGCTCATGTTGTAGGGCCATGAA
AGCGGCCATTCTTGTGATTCTTTGCACTTCTGGAACGGTGTATTGTTCACTATCCCAA
GCGACACCATCACCATCGTCTTCCTTTCTCTTACCAAAGTAAATACCTCCCACTAATT
CTCTGACAACAACGAAGTCAGTACCTTTAGCAAATTGTGGCTTGATTGGAGATAAGT
CTAAAAGAGAGTCGGATGCAAAGTTACATGGTCTTAAGTTGGCGTACAATTGAAGTT
CTTTACGGATTTTTAGTAAACCTTGTTCAGGTCTAACACTACCGGTACCCCATTTAGG
ACCACCCACAGCACCTAACAAAACGGCATCAGCCTTCTTGGAGGCTTCCAGCGCCTC
ATCTGGAAGTGGAACACCTGTAGCATCGATAGCAGCACCACCAATTAAATGATTTC
GAAATCGAACTTGACATTGGAACGAACATCAGAAATAGCTTTAAGAACCTTAATGG
CTTCGGCTGTGATTTCTTGACCAACGTGGTCACCTGGCAAAACGACGATCTTCTTAG
GGGCAGACATTAGAATGGTATATCCTTGAAATATATATATATATATTGCTGAAATGT

AAAAGGTAAGAAAAGTTAGAAAGTAAGACGATTGCTAACCACCTATTGGAAAAAC
AATAGGTCCTTAAATAATATTGTCAACTTCAAGTATTGTGATGCAAGCATTTAGTCA
TGAACGCTTCTCTATTCTATATGAAAAGCCGGTTCCGGCGCTCTCACCTTTCCTTTTT
CTCCCAATTTTTCAGTTGAAAAAGGTATATGCGTCAGGCGACCTCTGAAATTAACAA
AAAATTTCCAGTCATCGAATTTGATTCTGTGCGATAGCGCCCCTGTGTGTTCTCGTTA
TGTTGAGGAAAAAAATAATGGTTGCTAAGAGATTCGAACTCTTGCATCTTACGATAC
CTGAGTATTCCCACAGTTAACTGCGGTCAAGATATTTCTTGAATCAGGCGCCTTAGA
CCGCTCGGCCAAACAACCAATTACTTGTTGAGAAATAGAGTATAATTATCCTATAAA
TATAACGTTTTTGAACACACATGAACAAGGAAGTACAGGACAATTGATTTTGAAGA
GAATGTGGATTTTGATGTAATTGTTGGGATTCCATTTTTAATAAGGCAATAATATTA
GGTATGTAGATATACTAGAAGTTCTCCTCGACCGGTCGATATGCGGTGTGAAATACC
GCACAGATGCGTAAGGAGAAAATACCGCATCAGGAAATTGTAAGCGTTAATATTTT
GTTAAAATTCGCGTTAAATTTTTGTTAAATCAGCTCATTTTTTAACCAATAGGCCGAA
ATCGGCAAAATCCCTTATAAATCAAAAGAATAGACCGAGATAGGGTTGAGTGTTGT
TCCAGTTTGGAACAAGAGTCCACTATTAAAGAACGTGGACTCCAACGTCAAAGGGC
GAAAAACCGTCTATCAGGGCGATGGCCCACTACGTGAACCATCACCCTAATCAAGT
TTTTTGGGGTCGAGGTGCCGTAAAGCACTAAATCGGAACCCTAAAGGGAGCCCCCG
ATTTAGAGCTTGACGGGGAAAGCCGGCGAACGTGGCGAGAAAGGAAGGGAAGAAA
GCGAAAGGAGCGGGCGCTAGGGCGCTGGCAAGTGTAGCGGTCACGCTGCGCGTAAC
CACCACACCCGCCGCGCTTAATGCGCCGCTACAGGGCGCGTCCATTCGCCATTCAGG
CTGCGCAACTGTTGGGAAGGGCGATCGGTGCGGGCCTCTTCGCTATTACGCCAGCTG
GCGAAAGGGGGATGTGCTGCAAGGCGATTAAGTTGGGTAACGCCAGGGTTTTCCCA
GTCACGACGTTGTAAAACGACGGCCAGTGAATTGTAATACGACTCACTATAGGGCG
AATTGGAGCTCCACCGCGGTGGCGGCCGCTCTAGACAATACGAAGGAACGATCCCG
CGTCAGCAGATGTCACAATTTGATTATTTCATTGTCACATGCTTTTTGACTCATCCTT
CCGCGAAAGGAGAACTTTTTACAATGCCGCCGGCTCTGTTAGCGTACCCTTCTACAA
GCAGATAGCAGAACAAACACATGATATATTCAAAAGGTGCAATGTCAGAGAACATA
TATTGCGCCCCTGTCCTGTAGACATCAGTCATTTTTCGCGGGACTTGAATGGCGCAC
CATTATCCTATACAGAAAACATTAAATTTACTGCAAAATTTTGGGCAAACGCTTGGA
TAGACTATGAATCCGTCATGAAGACACTACTTGTAAAATTATATATATCCTTTTTTTT
CATGTGCTAGTAAGAATTAGCAACTACTTTCATTCATCTTCGACGCAACTTCGAGGA
AAGCACCTTTTTACTTGCTGTAGCTCTATTCTCTTCCCCAACCACCTTTTCCTTTATTC
CAAAATTTTTAAAACTTTTTCTGTTACATTATATAATCTTCTGTCTGAAATGTTTGGA
TATAACGCCTCTTGATCCACTTTGTATATGCGTGCTATTTATTTTCAGATTTATAAAG
AGTATGAGCGTCATTTACATAAATAGCTGAAGTTATTCATGGAAAATACGAAGAGC
ACGTATGTGAGCCAACAGAACATTTGACGTAAGACTCTACAATGTGCCAAGAACTG
GACAAGTAGAGGACTGAGAACTTTATTTCAATTCATTGCTCCTTTTTGGTGGCGCTA
CCTTTAGCGAAGGTCAATGATGAATGTGCACATGCTGTCGAAACCAAAAAGCAAAT
TCTAACCAACTTCAAAATGACATAGTCATCTGATATTTCTACTCATTATAGATAGTAT
GGGAGCCTTGAAACGAAAAGTAAGTAAAAAGCTGGATATGAGCAGTATGAGGTAG
ACCTTAGCTACATCATTTCCCCCAATAGCTGCTGCAAATATCTGGTTAAATTTGTGAT
TCCATGAAGAGGATAACAGACTTGTTAAAAAGCATCCTGTCAAAATCTAATTTTTGA
AGGGCAGTATTCAATTCATAATTTACTTTAGCTTAGATCCTTCCAATTTATACATGGT
ATTATAACCAGATCATAAACTACAATCTGTCGCTACCGCATGTACGAAGAATACTTA
AGTCACTTGCTCTCTCATCATTTGTACATTTTTCAGTAATACCGTTTGATAGCGCCGT
CTTTATTACCCGGATTATTCCGTTGACCCTGAATGAAAAATTTTTCAGAAATCCAGT

GCTAAGCGTCAAATCAATGAAATACATCACTGTATTTTTAACTGATATACTGTTGGT
GTGGCCTTAACGACACCTTTATTTCTTAATTCATTTCGGCTTGTCTCCTTATTAAGAC
TACAGAAATAGACAAAGGAAATACTTCAATAATGGATAACGAGGTTGAAAAAAATA
TTGAGATCTGGAAGGTCAAGAAGTTGGTCCAATCTTTAGAAAAAGCT**G**GAGGTAAT
GGTACTTCTATGATTTCCTTAGTTATTCCTCCTAAGGGTCAAATTCCACTGTACCAAA
AAATGTTAACAGATGAATATGGTACTGCCTCGA**G**TATTAAATCTAGGGTTAATCGTC
TTTCCGTTTTATCTGCTATCACTTCCACCCAACAAAAGTTGAAGCTATATAATACTTT
GCCCAAGAACGGTTTAGTTTTATATTGTGGTGATATCATCACTGAAGATGGTAAAGA
AAAAAAGGTCACTTTTGACATCGAACCTTACAAACCTATCAACACATCCTTATATTT
GTGTGATAACAAATTTCATACAGAAGTTCTTTCGGAATTGCTTCAAGCTGACGACAA
GTTCGGTTTTATAGTCATGGACGGTCAAGGTACTTTGTTTGGTTCTGTGTCCGGTAAT
ACGAGAACTGTTTTACATAAATTTACTGTCGATCTGCCAAAAAAGCATGGTAGAGGT
GGTCAATCTGCGCTTCGTTTTGCTCGTTTAAGAGAAGAAAAAAGACATAATTATGTG
AGAAAGGTCGCCGAAGTTGCTGTTCAAAATTTTATTACTAATGACAAAGTCAATGTT
AAGGGTTTAATTTTAGCTGGTTCTGCTGACTTTAAGACCGATTTGGCTAAATCTGAAT
TATTCGATCCAAGACTAGCATGTAAGGTTATTTCCATCGTGGATGTTTCTTATGGTGG
TGAAAACGGTTTCAACCAGGCTATCGAACTTTCTGCCGAAGCGTTGGCCAATGTCAA
GTATGTTCAAGAAAAGAAATTATTGGAGGCATATTT**G**GACGAAATTTCCCAGGACA
CTGGTAAATTCTGTTATGGTATAGATGATACTTTAAAGGCATTGGATTTAGGTGCAG
TCGAAAAATTAATTGTTTTCGAAAATTTGGAAACTATCAGATATACATTTAAAGATG
CCGAGGATAATGAGGTTATAAAATTCGCTGAACCAGAAGCCAAGGACAAGTCGTTT
GCTATTGACAAAGCTACCGGCCAAGAAATGA**A**ACGTTGTCTCCGAAGAACCTTTAATT
GAATGGCTAGCAGCTAACTACAAAAACTTCGGTGCTACCTTGGAATTCATCACAGAC
AAATCTTCAGAAGG**G**GCCCAATTTGTCACAGGTTTTGGTGGTATTGGTGCCATGCTG
CGTTACAAAGTTAATTTTGAACAACTAGTTGATGAA**C**CTGAGGATGAATATTATGAC
GAAGATGAAGGATCCGACTATGATTTCATTTAAATAAATAAAAGGGGGAGAAAAAA
ATCGAATCAAAAAGAATTTAATCACTAGATGCCAGATTTAAATTAAATTCGCTTTTA
ATTTTTTGTACAATATAATATATACTTGGTAAACCTTTTGCTCTATATTGAGCTAATT
CCTTTGTTGAAAGTACATAGTGGGTTTAGAGGACCGTGTATATTACGTAGAAAATAC
AGTGAAAGGAGAGTTTCTCTTCAAAAGCCTCGACGGTATCGATAAGCTTATCGATAC
CGTCGACCTCGAGGGGGGGCCCGGTACCAGCTTTTGTTCCCTTTAGTGAGGGTTAAT
TTCGAGCTTGGCGTAATCATGGTCATAGCTGTTTCCTGTGTGAAATTGTTATCCGCTC
ACAATTCCACACAACATACGAGCCGGAAGCATAAAGTGTAAAGCCTGGGGTGCCTA
ATGAGTGAGCTAACTCACATTAATTGCGTTGCGCTCACTGCCCGCTTTCCAGTCGGG
AAACCTGTCGTGCCAGCTGCATTAATGAATCGGCCAACGCGCGGGGAGAGGCGGTT
TGCGTATTGGGCGCTCTTCCGCTTCCTCGCTCACTGACTCGCTGCGCTCGGTCGTTCG
GCTGCGGCGAGCGGTATCAGCTCACTCAAAGGCGGTAATACGGTTATCCACAGAAT
CAGGGGATAACGCAGGAAAGAACATGTGAGCAAAAGGCCAGCAAAAGGCCAGGAA
CCGTAAAAAGGCCGCGTTGCTGGCGTTTTTCCATAGGCTCCGCCCCCCTGACGAGCA
TCACAAAAATCGACGCTCAAGTCAGAGGTGGCGAAACCCGACAGGACTATAAAGAT
ACCAGGCGTTTCCCCCTGGAAGCTCCCTCGTGCGCTCTCCTGTTCCGACCCTGCCGCT
TACCGGATACCTGTCCGCCTTTCTCCCTTCGGGAAGCGTGGCGCTTTCTCATAGCTCA
CGCTGTAGGTATCTCAGTTCGGTGTAGGTCGTTCGCTCCAAGCTGGGCTGTGTGCAC
GAACCCCCCGTTCAGCCCGACCGCTGCGCCTTATCCGGTAACTATCGTCTTGAGTCC
AACCCGGTAAGACACGACTTATCGCCACTGGCAGCAGCCACTGGTAACAGGATTAG
CAGAGCGAGGTATGTAGGCGGTGCTACAGAGTTCTTGAAGTGGTGGCCTAACTACG

GCTACACTAGAAGAACAGTATTTGGTATCTGCGCTCTGCTGAAGCCAGTTACCTTCG
GAAAAAGAGTTGGTAGCTCTTGATCCGGCAAACAAACCACCGCTGGTAGCGGTGGT
TTTTTTGTTTGCAAGCAGCAGATTACGCGCAGAAAAAAAGGATCTCAAGAAGATCCT
TTGATCTTTTCTACGGGGTCTGACGCTCAGTGGAACGAAAACTCACGTTAAGGGATT
TTGGTCATGAGATTATCAAAAAGGATCTTCACCTAGATCCTTTTAAATTAAAAATGA
AGTTTTAAATCAATCTAAAGTATATATGAGTAAACTTGGTCTGACAGTTACCAATGC
TTAATCAGTGAGGCACCTATCTCAGCGATCTGTCTATTTCGTTCATCCATAGTTGCCT
GACTCCCCGTCGTGTAGATAACTACGATACGGGAGGGCTTACCATCTGGCCCCAGTG
CTGCAATGATACCGCGAGACCCACGCTCACCGGCTCCAGATTTATCAGCAATAAACC
AGCCAGCCGGAAGGGCCGAGCGCAGAAGTGGTCCTGCAACTTTATCCGCCTCCATC
CAGTCTATTAATTGTTGCCGGGAAGCTAGAGTAAGTAGTTCGCCAGTTAATAGTTTG
CGCAACGTTGTTGCCATTGCTACAGGCATCGTGGTGTCACGCTCGTCGTTTGGTATG
GCTTCATTCAGCTCCGGTTCCCAACGATCAAGGCGAGTTACATGATCCCCCATGTTG
TGCAAAAAAGCGGTTAGCTCCTTCGGTCCTCCGATCGTTGTCAGAAGTAAGTTGGCC
GCAGTGTTATCACTCATGGTTATGGCAGCACTGCATAATTCTCTTACTGTCATGCCAT
CCGTAAGATGCTTTTCTGTGACTGGTGAGTACTCAACCAAGTCATTCTGAGAATAGT
GTATGCGGCGACCGAGTTGCTCTTGCCCGGCGTCAATACGGGATAATACCGCGCCAC
ATAGCAGAACTTTAAAAGTGCTCATCATTGGAAAACGTTCTTCGGGGCGAAAACTCT
CAAGGATCTTACCGCTGTTGAGATCCAGTTCGATGTAACCCACTCGTGCACC

**PTH353-SUP45-CR3T.9**
*SUP45* Promoter *SUP45-**CR3T.9***

GGGGAGAGGCGGTTTGCGTATTGGGCGCTCTTCCGCTTCCTCGCTCACTGACTCGCT
GCGCTCGGTCGTTCGGCTGCGGCGAGCGGTATCAGCTCACTCAAAGGCGGTAATAC
GGTTATCCACAGAATCAGGGGATAACGCAGGAAAGAACATGTGAGCAAAAGGCCA
GCAAAAGGCCAGGAACCGTAAAAAGGCCGCGTTGCTGGCGTTTTTCCATAGGCTCC
GCCCCCCTGACGAGCATCACAAAAATCGACGCTCAAGTCAGAGGTGGCGAAACCCG
ACAGGACTATAAAGATACCAGGCGTTTCCCCCTGGAAGCTCCCTCGTGCGCTCTCCT
GTTCCGACCCTGCCGCTTACCGGATACCTGTCCGCCTTTCTCCCTTCGGGAAGCGTG
GCGCTTTCTCATAGCTCACGCTGTAGGTATCTCAGTTCGGTGTAGGTCGTTCGCTCCA
AGCTGGGCTGTGTGCACGAACCCCCCGTTCAGCCCGACCGCTGCGCCTTATCCGGTA
ACTATCGTCTTGAGTCCAACCCGGTAAGACACGACTTATCGCCACTGGCAGCAGCCA
CTGGTAACAGGATTAGCAGAGCGAGGTATGTAGGCGGTGCTACAGAGTTCTTGAAG
TGGTGGCCTAACTACGGCTACACTAGAAGAACAGTATTTGGTATCTGCGCTCTGCTG
AAGCCAGTTACCTTCGGAAAAAGAGTTGGTAGCTCTTGATCCGGCAAACAAACCAC
CGCTGGTAGCGGTGGTTTTTTTGTTTGCAAGCAGCAGATTACGCGCAGAAAAAAAGG
ATCTCAAGAAGATCCTTTGATCTTTTCTACGGGGTCTGACGCTCAGTGGAACGAAAA
CTCACGTTAAGGATTTTGGTCATGAGATTATCAAAAAGGATCTTCACCTAGATCCT
TTTAAATTAAAAATGAAGTTTTAAATCAATCTAAAGTATATATGAGTAAACTTGGTC
TGACAGTTACCAATGCTTAATCAGTGAGGCACCTATCTCAGCGATCTGTCTATTTCGT
TCATCCATAGTTGCCTGACTCCCCGTCGTGTAGATAACTACGATACGGGAGGGCTTA
CCATCTGGCCCCAGTGCTGCAATGATACCGCGAGACCCACGCTCACCGGCTCCAGAT
TTATCAGCAATAAACCAGCCAGCCGGAAGGGCCGAGCGCAGAAGTGGTCCTGCAAC
TTTATCCGCCTCCATCCAGTCTATTAATTGTTGCCGGGAAGCTAGAGTAAGTAGTTC
GCCAGTTAATAGTTTGCGCAACGTTGTTGCCATTGCTACAGGCATCGTGGTGTCACG

CTCGTCGTTTGGTATGGCTTCATTCAGCTCCGGTTCCCAACGATCAAGGCGAGTTAC
ATGATCCCCCATGTTGTGCAAAAAAGCGGTTAGCTCCTTCGGTCCTCCGATCGTTGT
CAGAAGTAAGTTGGCCGCAGTGTTATCACTCATGGTTATGGCAGCACTGCATAATTC
TCTTACTGTCATGCCATCCGTAAGATGCTTTTCTGTGACTGGTGAGTACTCAACCAAG
TCATTCTGAGAATAGTGTATGCGGCGACCGAGTTGCTCTTGCCCGGCGTCAATACGG
GATAATACCGCGCCACATAGCAGAACTTTAAAAGTGCTCATCATTGGAAAACGTTCT
TCGGGGCGAAAACTCTCAAGGATCTTACCGCTGTTGAGATCCAGTTCGATGTAACCC
ACTCGTGCACCCAACTGATCTTCAGCATCTTTTACTTTCACCAGCGTTTCTGGGTGAG
CAAAAACAGGAAGGCAAAATGCCGCAAAAAAGGGAATAAGGGCGACACGGAAATG
TTGAATACTCATACTCTTCCTTTTTCAATATTATTGAAGCATTTATCAGGGTTATTGT
CTCATGAGCGGATACATATTTGAATGTATTTAGAAAAATAAACAAATAGGGGTTCCG
CGCACATTTCCCCGAAAAGTGCCACCTGGGTCCTTTTCATCACGTGCTATAAAAATA
ATTATAATTTAAATTTTTTAATATAAATATATAAATTAAAAATAGAAAGTAAAAAAA
GAAATTAAAGAAAAAATAGTTTTTGTTTTCCGAAGATGTAAAAGACTCTAGGGGGA
TCGCCAACAAATACTACCTTTTATCTTGCTCTTCCTGCTCTCAGGTATTAATGCCGAA
TTGTTTCATCTTGTCTGTGTAGAAGACCACACACGAAAATCCTGTGATTTTACATTTT
ACTTATCGTTAATCGAATGTATATCTATTTAATCTGCTTTTCTTGTCTAATAAATATA
TATGTAAAGTACGCTTTTTGTTGAAATTTTTTAAACCTTTGTTTATTTTTTTTTCTTCAT
TCCGTAACTCTTCTACCTTCTTTATTTACTTTCTAAAATCCAAATACAAAACATAAAA
ATAAATAAACACAGAGTAAATTCCCAAATTATTCCATCATTAAAAGATACGAGGCG
CGTGTAAGTTACAGGCAAGCGATCCGTCCTAAGAAACCATTATTATCATGACATTAA
CCTATAAAAATAGGCGTATCACGAGGCCCTTTCGTCTCGCGCGTTTCGGTGATGACG
GTGAAAACCTCTGACACATGCAGCTCCCGGAGACGGTCACAGCTTGTCTGTAAGCG
GATGCCGGGAGCAGACAAGCCCGTCAGGGCGCGTCAGCGGGTGTTGGCGGGTGTCG
GGGCTGGCTTAACTATGCGGCATCAGAGCAGATTGTACTGAGAGTGCACCATATCG
ACTACGTCGTTAAGGCCGTTTCTGACAGAGTAAAATTCTTGAGGGAACTTTCACCAT
TATGGGAAATGGTTCAAGAAGGTATTGACTTAAACTCCATCAAATGGTCAGGTCATT
GAGTGTTTTTTATTTGTTGTATTTTTTTTTTTTTAGAGAAAATCCTCCAATATATAAATT
AGGAATCATAGTTTCATGATTTTCTGTTACACCTAACTTTTTGTGTGGTGCCCTCCTC
CTTGTCAATATTAATGTTAAAGTGCAATTCTTTTTCCTTATCACGTTGAGCCATTAGT
ATCAATTTGCTTACCTGTATTCCTTTACATCCTCCTTTTTCTCCTTCTTGATAAATGTA
TGTAGATTGCGTATATAGTTTCGTCTACCCTATGAACATATTCCATTTTGTAATTTCG
TGTCGTTTCTATTATGAATTTCATTTATAAAGTTTATGTACAAATATCATAAAAAAAG
AGAATCTTTTTAAGCAAGGATTTTCTTAACTTCTTCGGCGACAGCATCACCGACTTCG
GTGGTACTGTTGGAACCACCTAAATCACCAGTTCTGATACCTGCATCCAAAACCTTT
TTAACTGCATCTTCAATGGCCTTACCTTCTTCAGGCAAGTTCAATGACAATTTCAACA
TCATTGCAGCAGACAAGATAGTGGCGATAGGGTTGACCTTATTCTTTGGCAAATCTG
GAGCAGAACCGTGGCATGGTTCGTACAAACCAAATGCGGTGTTCTTGTCTGGCAAA
GAGGCCAAGGACGCAGATGGCAACAAACCCAAGGAACCTGGGATAACGGAGGCTT
CATCGGAGATGATATCACCAAACATGTTGCTGGTGATTATAATACCATTTAGGTGGG
TTGGGTTCTTAACTAGGATCATGGCGGCAGAATCAATCAATTGATGTTGAACCTTCA
ATGTAGGGAATTCGTTCTTGATGGTTTCCTCCACAGTTTTTCTCCATAATCTTGAAGA
GGCCAAAACATTAGCTTTATCCAAGGACCAAATAGGCAATGGTGGCTCATGTTGTAG
GGCCATGAAAGCGGCCATTCTTGTGATTCTTTGCACTTCTGGAACGGTGTATTGTTCA
CTATCCCAAGCGACACCATCACCATCGTCTTCCTTTCTCTTACCAAAGTAAATACCTC
CCACTAATTCTCTGACAACAACGAAGTCAGTACCTTTAGCAAATTGTGGCTTGATTG

GAGATAAGTCTAAAAGAGAGTCGGATGCAAAGTTACATGGTCTTAAGTTGGCGTAC
AATTGAAGTTCTTTACGGATTTTTAGTAAACCTTGTTCAGGTCTAACACTACCGGTAC
CCCATTTAGGACCACCCACAGCACCTAACAAAACGGCATCAGCCTTCTTGGAGGCTT
CCAGCGCCTCATCTGGAAGTGGAACACCTGTAGCATCGATAGCAGCACCACCAATT
AAATGATTTTCGAAATCGAACTTGACATTGGAACGAACATCAGAAATAGCTTTAAG
AACCTTAATGGCTTCGGCTGTGATTTCTTGACCAACGTGGTCACCTGGCAAAACGAC
GATCTTCTTAGGGGCAGACATTAGAATGGTATATCCTTGAAATATATATATATATAT
TGCTGAAATGTAAAAGGTAAGAAAAGTTAGAAAGTAAGACGATTGCTAACCACCTA
TTGGAAAAAACAATAGGTCCTTAAATAATATTGTCAACTTCAAGTATTGTGATGCAA
GCATTTAGTCATGAACGCTTCTCTATTCTATATGAAAAGCCGGTTCCGGCGCTCTCAC
CTTTCCTTTTTCTCCCAATTTTTCAGTTGAAAAAGGTATATGCGTCAGGCGACCTCTG
AAATTAACAAAAAATTTCCAGTCATCGAATTTGATTCTGTGCGATAGCGCCCCTGTG
TGTTCTCGTTATGTTGAGGAAAAAAATAATGGTTGCTAAGAGATTCGAACTCTTGCA
TCTTACGATACCTGAGTATTCCCACAGTTAACTGCGGTCAAGATATTTCTTGAATCA
GGCGCCTTAGACCGCTCGGCCAAACAACCAATTACTTGTTGAGAAATAGAGTATAAT
TATCCTATAAATATAACGTTTTTGAACACACATGAACAAGGAAGTACAGGACAATTG
ATTTTGAAGAGAATGTGGATTTTGATGTAATTGTTGGGATTCCATTTTTAATAAGGC
AATAATATTAGGTATGTAGATATACTAGAAGTTCTCCTCGACCGGTCGATATGCGGT
GTGAAATACCGCACAGATGCGTAAGGAGAAAATACCGCATCAGGAAATTGTAAGCG
TTAATATTTTGTTAAAATTCGCGTTAAATTTTTGTTAAATCAGCTCATTTTTTAACCA
ATAGGCCGAAATCGGCAAAATCCCTTATAAATCAAAAGAATAGACCGAGATAGGGT
TGAGTGTTGTTCCAGTTTGGAACAAGAGTCCACTATTAAAGAACGTGGACTCCAACG
TCAAAGGGCGAAAAACCGTCTATCAGGGCGATGGCCCACTACGTGAACCATCACCC
TAATCAAGTTTTTTGGGGTCGAGGTGCCGTAAAGCACTAAATCGGAACCCTAAAGG
GAGCCCCCGATTTAGAGCTTGACGGGGAAAGCCGGCGAACGTGGCGAGAAAGGAA
GGGAAGAAAGCGAAAGGAGCGGGCGCTAGGGCGCTGGCAAGTGTAGCGGTCACGC
TGCGCGTAACCACCACACCCGCCGCGCTTAATGCGCCGCTACAGGGCGCGTCCATTC
GCCATTCAGGCTGCGCAACTGTTGGGAAGGGCGATCGGTGCGGGCCTCTTCGCTATT
ACGCCAGCTGGCGAAAGGGGGATGTGCTGCAAGGCGATTAAGTTGGGTAACGCCAG
GGTTTTCCCAGTCACGACGTTGTAAAACGACGGCCAGTGAATTGTAATACGACTCAC
TATAGGGCGAATTGGAGCTCCACCGCGGTGGCGGCCGCTCTAGACAATACGAAGGA
ACGATCCCGCGTCAGCAGATGTCACAATTTGATTATTTCATTGTCACATGCTTTTTGA
CTCATCCTTCCGCGAAAGGAGAACTTTTTACAATGCCGCCGGCTCTGTTAGCGTACC
CTTCTACAAGCAGATAGCAGAACAAACACATGATATATTCAAAAGGTGCAATGTCA
GAGAACATATATTGCGCCCCTGTCCTGTAGACATCAGTCATTTTTCGCGGGACTTGA
ATGGCGCACCATTATCCTATACAGAAAACATTAAATTTACTGCAAAATTTTGGGCAA
ACGCTTGGATAGACTATGAATCCGTCATGAAGACACTACTTGTAAAATTATATATAT
CCTTTTTTTTCATGTGCTAGTAAGAATTAGCAACTACTTTCATTCATCTTCGACGCAA
CTTCGAGGAAAGCACCTTTTTACTTGCTGTAGCTCTATTCTCTTCCCCAACCACCTTT
TCCTTTATTCCAAAATTTTTAAAACTTTTTCTGTTACATTATATAATCTTCTGTCTGAA
ATGTTTGGATATAACGCCTCTTGATCCACTTTGTATATGCGTGCTATTTATTTTCAGA
TTTATAAAGAGTATGAGCGTCATTTACATAAATAGCTGAAGTTATTCATGGAAAATA
CGAAGAGCACGTATGTGAGCCAACAGAACATTTGACGTAAGACTCTACAATGTGCC
AAGAACTGGACAAGTAGAGGACTGAGAACTTTATTTCAATTCATTGCTCCTTTTTGG
TGGCGCTACCTTTAGCGAAGGTCAATGATGAATGTGCACATGCTGTCGAAACCAAA
AAGCAAATTCTAACCAACTTCAAAATGACATAGTCATCTGATATTTCTACTCATTAT

AGATAGTATGGGAGCCTTGAAACGAAAAGTAAGTAAAAAGCTGGATATGAGCAGTA
TGAGGTAGACCTTAGCTACATCATTTCCCCCAATAGCTGCTGCAAATATCTGGTTAA
ATTTGTGATTCCATGAAGAGGATAACAGACTTGTTAAAAAGCATCCTGTCAAAATCT
AATTTTTGAAGGGCAGTATTCAATTCATAATTTACTTTAGCTTAGATCCTTCCAATTT
ATACATGGTATTATAACCAGATCATAAACTACAATCTGTCGCTACCGCATGTACGAA
GAATACTTAAGTCACTTGCTCTCTCATCATTTGTACATTTTTCAGTAATACCGTTTGA
TAGCGCCGTCTTTATTACCCGGATTATTCCGTTGACCCTGAATGAAAAATTTTTTCAG
AAATCCAGTGCTAAGCGTCAAATCAATGAAATACATCACTGTATTTTTAACTGATAT
ACTGTTGGTGTGGCCTTAACGACACCTTTATTTCTTAATTCATTTCGGCTTGTCTCCTT
ATTAAGACTACAGAAATAGACAAAGGAAATACTTCAATAATGGATAACGAGGTTGA
AAAAAATATTGAGATCTGGAAGGTCAAGAAGTTGGTCCAATCTTTAGAAAAAGCT**G**
GAGGTAATGGTACTTCTATGATTTCCTTAGTTATTCCTCCTAAGGGTCAAATTCCACT
GTACCAAAAAATGTTAACAGATGAATATGGTACTGCCTCGAATATTAAATCTAGGGT
TAATCGTCTTTCCG**C**TTTATCTGCTATCACTTCCACCCAACAAAAGTTGAAGCTATAT
AATACTTTGCCCAAGAACGGTTTAGTTTTATATTGTGGTGATATCATCACTGAAGAT
GGTAAAGAAAAAAAGGTCACTTTTGACATCGAACCTTACAAACCTATCAACACATC
CTTATATTTGTGTGATAACAAATTTCATACAGAAGTTCTTTCGGAATTGCTTCAAGCT
GACGACAAGTTCGGTTTTATAGTCATGGACGGTCAAGGTACTTTGTTTGGTTCTGTGT
CCGGTAATACGAGAACTG**C**TTTACATAAATTTACTGTCGATCTGCCAAAAAAGCATG
GTAGAGGTGGTCAATCTGCGCTTCGTTTTGCTCGTTTAAGAGAAGAAAAAAGACATA
ATTATGTGAGAAAGGTCGCCGAAGTTGCTGTTCAAAATTTTATTACTAATGACAAAG
TCAATGTTAAGGGTTTAATTTTAGCTGGTTCTGCTGACTTTAAGACCGATTTGGCTAA
ATCTGAATTATTCGATCCAAGACTAGCATGTAAGGTTATTTCCATCGTGGATGTTTCT
TATGGTGGTGAAAACGGTTTCAACCAGGCTATCGAACTTTCTGCCGAAGCGTTGGCC
AATGTCAAGTATGTTCAAGAAAAGAAATTATTGGAGGCATATTTTGACGAAATTTCC
CAGGACACTGGTAAATTCTGTTATGGTATAGATGATACTTTAAAGGCATTGGATTTA
GGTGCAGTCGAAAAATTAATTGTTTTCGAAAATTTGGAAACTATCAGATATACATTT
AAAGATGCCGAGGATAATGAGGTTATAAAATTCGCTGAACCAGAAGCCAAGGACAA
GTCGTTTGCTATTGACAAAGCTACCGGCCAAGAAATGGACGTTGTCTCCG**G**AGAACC
TTTAATTGAATGGCTAGCAGCTAACTACAAAAACTTCGGTGCTACCTTGGAATTCAT
CACAGACAAATCTTCAGAAGGTGCCCAATTTGTCACAGGTTTTGGTGGTATTGGTGC
CATGCTGCGTTACAAAGTTAATTTTGAACAACTAGTTGATGAATCTGAGGATGAATA
TTATGACGAAGATGAAGGATCCGACTATGATTTCATTTAAATAAATAAAAGGGAGA
GAAAAAAATCGAATCAAAAAGAATTTAATCACTAGATGCCAGATTTAAATTAAATT
CGCTTTTAATTTTTTGTACAATATAATATATACTTGGTAAACCTTTTGCTCTATATTG
AGCTAATTCCTTTGTTGAAAGTACATAGTGGGTTTAGAGGACCGTGTATATTACGTA
GAAAATACAGTGAAAGGAGAGTTTCTCTTCAAAAGCCTCGACGGTATCGATAAGCT
TATCGATACCGTCGACCTCGAGGGGGGGCCCGGTACCAGCTTTTGTTCCCTTTAGTG
AGGGTTAATTTCGAGCTTGGCGTAATCATGGTCATAGCTGTTTCCTGTGTGAAATTGT
TATCCGCTCACAATTCCACACAACATACGAGCCGGAAGCATAAAGTGTAAAGCCTG
GGGTGCCTAATGAGTGAGCTAACTCACATTAATTGCGTTGCGCTCACTGCCCGCTTT
CCAGTCGGGAAACCTGTCGTGCCAGCTGCATTAATGAATCGGCCAACGCGC

**PTH353-SUP45-BR2.2**

*SUP45* Promoter *SUP45****-BR2.2***

CACTCGTGCACCCAACTGATCTTCAGCATCTTTTACTTTCACCAGCGTTTCTGGGTGA
GCAAAAACAGGAAGGCAAAATGCCGCAAAAAAGGGAATAAGGGCGACACGGAAAT
GTTGAATACTCATACTCTTCCTTTTTCAATATTATTGAAGCATTTATCAGGGTTATTG
TCTCATGAGCGGATACATATTTGAATGTATTTAGAAAAATAAACAAATAGGGGTTCC
GCGCACATTTCCCCGAAAAGTGCCACCTGGGTCCTTTTCATCACGTGCTATAAAAAT
AATTATAATTTAAATTTTTTAATATAAATATATAAATTAAAAATAGAAAGTAAAAAA
AGAAATTAAAGAAAAAATAGTTTTTGTTTTCCGAAGATGTAAAAGACTCTAGGGGG
ATCGCCAACAAATACTACCTTTTATCTTGCTCTTCCTGCTCTCAGGTATTAATGCCGA
ATTGTTTCATCTTGTCTGTGTAGAAGACCACACACGAAAATCCTGTGATTTTACATTT
TACTTATCGTTAATCGAATGTATATCTATTTAATCTGCTTTTCTTGTCTAATAAATAT
ATATGTAAAGTACGCTTTTTGTTGAAATTTTTTAAACCTTTGTTTATTTTTTTTTCTTCA
TTCCGTAACTCTTCTACCTTCTTTATTTACTTTCTAAAATCCAAATACAAAACATAAA
AATAAATAAACACAGAGTAAATTCCCAAATTATTCCATCATTAAAAGATACGAGGC
GCGTGTAAGTTACAGGCAAGCGATCCGTCCTAAGAAACCATTATTATCATGACATTA
ACCTATAAAAATAGGCGTATCACGAGGCCCTTTCGTCTCGCGCGTTTCGGTGATGAC
GGTGAAAACCTCTGACACATGCAGCTCCCGGAGACGGTCACAGCTTGTCTGTAAGC
GGATGCCGGGAGCAGACAAGCCCGTCAGGGCGCGTCAGCGGGTGTTGGCGGGTGTC
GGGGCTGGCTTAACTATGCGGCATCAGAGCAGATTGTACTGAGAGTGCACCATATC
GACTACGTCGTTAAGGCCGTTTCTGACAGAGTAAAATTCTTGAGGGAACTTTCACCA
TTATGGGAAATGGTTCAAGAAGGTATTGACTTAAACTCCATCAAATGGTCAGGTCAT
TGAGTGTTTTTTATTTGTTGTATTTTTTTTTTTTTAGAGAAAATCCTCCAATATATAAA
TTAGGAATCATAGTTTCATGATTTTCTGTTACACCTAACTTTTTGTGTGGTGCCCTCC
TCCTTGTCAATATTAATGTTAAAGTGCAATTCTTTTTCCTTATCACGTTGAGCCATTA
GTATCAATTTGCTTACCTGTATTCCTTTACATCCTCCTTTTTCTCCTTCTTGATAAATG
TATGTAGATTGCGTATATAGTTTCGTCTACCCTATGAACATATTCCATTTTGTAATTT
CGTGTCGTTTCTATTATGAATTTCATTTATAAAGTTTATGTACAAATATCATAAAAAA
AGAGAATCTTTTTAAGCAAGGATTTTCTTAACTTCTTCGGCGACAGCATCACCGACT
TCGGTGGTACTGTTGGAACCACCTAAATCACCAGTTCTGATACCTGCATCCAAAACC
TTTTTAACTGCATCTTCAATGGCCTTACCTTCTTCAGGCAAGTTCAATGACAATTTCA
ACATCATTGCAGCAGACAAGATAGTGGCGATAGGGTTGACCTTATTCTTTGGCAAAT
CTGGAGCAGAACCGTGGCATGGTTCGTACAAACCAAATGCGGTGTTCTTGTCTGGCA
AAGAGGCCAAGGACGCAGATGGCAACAAACCCAAGGAACCTGGGATAACGGAGGC
TTCATCGGAGATGATATCACCAAACATGTTGCTGGTGATTATAATACCATTTAGGTG
GGTTGGGTTCTTAACTAGGATCATGGCGGCAGAATCAATCAATTGATGTTGAACCTT
CAATGTAGGGAATTCGTTCTTGATGGTTTCCTCCACAGTTTTTCTCCATAATCTTGAA
GAGGCCAAAACATTAGCTTTATCCAAGGACCAAATAGGCAATGGTGGCTCATGTTGT
AGGGCCATGAAAGCGGCCATTCTTGTGATTCTTTGCACTTCTGGAACGGTGTATTGT
TCACTATCCCAAGCGACACCATCACCATCGTCTTCCTTTCTCTTACCAAAGTAAATAC
CTCCCACTAATTCTCTGACAACAACGAAGTCAGTACCTTTAGCAAATTGTGGCTTGA
TTGGAGATAAGTCTAAAAGAGAGTCGGATGCAAAGTTACATGGTCTTAAGTTGGCG
TACAATTGAAGTTCTTTACGGATTTTTAGTAAACCTTGTTCAGGTCTAACACTACCGG
TACCCCATTTAGGACCACCCACAGCACCTAACAAAACGGCATCAGCCTTCTTGGAGG
CTTCCAGCGCCTCATCTGGAAGTGGAACACCTGTAGCATCGATAGCAGCACCACCA
ATTAAATGATTTTCGAAATCGAACTTGACATTGGAACGAACATCAGAAATAGCTTTA
AGAACCTTAATGGCTTCGGCTGTGATTTCTTGACCAACGTGGTCACCTGGCAAAACG
ACGATCTTCTTAGGGGCAGACATTAGAATGGTATATCCTTGAAATATATATATATAT

ATTGCTGAAATGTAAAAGGTAAGAAAAGTTAGAAAGTAAGACGATTGCTAACCACC
TATTGGAAAAAACAATAGGTCCTTAAATAATATTGTCAACTTCAAGTATTGTGATGC
AAGCATTTAGTCATGAACGCTTCTCTATTCTATATGAAAAGCCGGTTCCGGCGCTCT
CACCTTTCCTTTTTCTCCCAATTTTTCAGTTGAAAAAGGTATATGCGTCAGGCGACCT
CTGAAATTAACAAAAAATTTCCAGTCATCGAATTTGATTCTGTGCGATAGCGCCCCT
GTGTGTTCTCGTTATGTTGAGGAAAAAAATAATGGTTGCTAAGAGATTCGAACTCTT
GCATCTTACGATACCTGAGTATTCCCACAGTTAACTGCGGTCAAGATATTTCTTGAA
TCAGGCGCCTTAGACCGCTCGGCCAAACAACCAATTACTTGTTGAGAAATAGAGTAT
AATTATCCTATAAATATAACGTTTTTGAACACACATGAACAAGGAAGTACAGGACA
ATTGATTTTGAAGAGAATGTGGATTTTGATGTAATTGTTGGGATTCCATTTTTAATAA
GGCAATAATATTAGGTATGTAGATATACTAGAAGTTCTCCTCGACCGGTCGATATGC
GGTGTGAAATACCGCACAGATGCGTAAGGAGAAAATACCGCATCAGGAAATTGTAA
GCGTTAATATTTTGTTAAAATTCGCGTTAAATTTTTGTTAAATCAGCTCATTTTTTAA
CCAATAGGCCGAAATCGGCAAAATCCCTTATAAATCAAAAGAATAGACCGAGATAG
GGTTGAGTGTTGTTCCAGTTTGGAACAAGAGTCCACTATTAAAGAACGTGGACTCCA
ACGTCAAAGGGCGAAAAACCGTCTATCAGGGCGATGGCCCACTACGTGAACCATCA
CCCTAATCAAGTTTTTTGGGGTCGAGGTGCCGTAAAGCACTAAATCGGAACCCTAAA
GGGAGCCCCCGATTTAGAGCTTGACGGGGAAAGCCGGCGAACGTGGCGAGAAAGG
AAGGGAAGAAAGCGAAAGGAGCGGGCGCTAGGGCGCTGGCAAGTGTAGCGGTCAC
GCTGCGCGTAACCACCACACCCGCCGCGCTTAATGCGCCGCTACAGGGCGCGTCCAT
TCGCCATTCAGGCTGCGCAACTGTTGGGAAGGGCGATCGGTGCGGGCCTCTTCGCTA
TTACGCCAGCTGGCGAAAGGGGGATGTGCTGCAAGGCGATTAAGTTGGGTAACGCC
AGGGTTTTCCCAGTCACGACGTTGTAAAACGACGGCCAGTGAATTGTAATACGACTC
ACTATAGGGCGAATTGGAGCTCCACCGCGGTGGCGGCCGCTCTAGACAATACGAAG
GAACGATCCCGCGTCAGCAGATGTCACAATTTGATTATTTCATTGTCACATGCTTTTT
GACTCATCCTTCCGCGAAAGGAGAACTTTTTACAATGCCGCCGGCTCTGTTAGCGTA
CCCTTCTACAAGCAGATAGCAGAACAAACACATGATATATTCAAAAGGTGCAATGT
CAGAGAACATATATTGCGCCCCTGTCCTGTAGACATCAGTCATTTTTCGCGGGACTT
GAATGGCGCACCATTATCCTATACAGAAAACATTAAATTTACTGCAAAATTTTGGGC
AAACGCTTGGATAGACTATGAATCCGTCATGAAGACACTACTTGTAAAATTATATAT
ATCCTTTTTTTTCATGTGCTAGTAAGAATTAGCAACTACTTTCATTCATCTTCGACGC
AACTTCGAGGAAAGCACCTTTTTACTTGCTGTAGCTCTATTCTCTTCCCCAACCACCT
TTTCCTTTATTCCAAAATTTTTAAAACTTTTTCTGTTACATTATATAATCTTCTGTCTG
AAATGTTTGGATATAACGCCTCTTGATCCACTTTGTATATGCGTGCTATTTATTTTCA
GATTTATAAAGAGTATGAGCGTCATTTACATAAATAGCTGAAGTTATTCATGGAAAA
TACGAAGAGCACGTATGTGAGCCAACAGAACATTTGACGTAAGACTCTACAATGTG
CCAAGAACTGGACAAGTAGAGGACTGAGAACTTTATTTCAATTCATTGCTCCTTTTT
GGTGGCGCTACCTTTAGCGAAGGTCAATGATGAATGTGCACATGCTGTCGAAACCA
AAAAGCAAATTCTAACCAACTTCAAAATGACATAGTCATCTGATATTTCTACTCATT
ATAGATAGTATGGGAGCCTTGAAACGAAAAGTAAGTAAAAAGCTGGATATGAGCAG
TATGAGGTAGACCTTAGCTACATCATTTCCCCCAATAGCTGCTGCAAATATCTGGTT
AAATTTGTGATTCCATGAAGAGGATAACAGACTTGTTAAAAAGCATCCTGTCAAAAT
CTAATTTTTGAAGGGCAGTATTCAATTCATAATTTACTTTAGCTTAGATCCTTCCAAT
TTATACATGGTATTATAACCAGATCATAAACTACAATCTGTCGCTACCGCATGTACG
AAGAATACTTAAGTCACTTGCTCTCTCATCATTTGTACATTTTTCAGTAATACCGTTT
GATAGCGCCGTCTTTATTACCCGGATTATTCCGTTGACCCTGAATGAAAAATTTTTC

AGAAATCCAGTGCTAAGCGTCAAATCAATGAAATACATCACTGTATTTTTAACTGAT
ATACTGTTGGTGTGGCCTTAACGACACCTTTATTTCTTAATTCATTTCGGCTTGTCTC
CTTATTAAGACTACAGAAATAGACAAAGGAAATACTTCAATAATGGATAACGAGGT
TGAAAAAAATATTGAGATCTGGAAGGTCAAGAAGTTGGTCCAATCTTTAGA**G**AAAG
CTAGAGGTAATGGTACTTCTATGATTTCCTTAGTTATTCCTCCTAAGGGTCAAATTCC
ACTGTACCAAAAAATGTTAACAGATGAATATGGTACTGCCTCGAATATTAAATCTAG
GGTTAATCGTCTTTCCGTTTTAT**T**TGCTATCACTTCCACCCAACAAAAGTTGAAGCTA
TATAATACTTTGCCCAAGAACGGTTTAGTTTTATATTGTGGTGATATCATCACTGAAG
ATGGTA**CG**GAAAAAAAGGTCACTTTTGACATCGAACC**C**TACAAACCTATCAACACA
TCCTTATATTTGTGTGATAACAAATTTCATACAGAAGTTCTTTCGGAATTGCTTCAAG
CTGACGACAAGTTCGGTTTTATAGTCATGGACGGTCAAGGTACTTTGTTTGGTTCTGT
GTCCGGTAATACGAGAACTGTTTTACATAAATTTACTGTCGATCTGCCAAAAAAGCA
TGGTAGAGGTGGTCAATCTGCGCTTCGTTTTGCTCGTTTAAGAGAAGAAAAAAGACA
TAATTATGTGAGAAAGGTCGCCGAAGTTGCTGTTCAAAATTTTATTACTAATGACAA
AGTCAATGTTAAGGGTTTAATTTTAGCTGGTTCTGCTGACTTTAAGACCGATTTGGCT
AAATCTGAATTATTCGATCCAAGACTAGCATGTAAGGTTATTTCCATCGTGGATGTT
TCTTATGGTGGTGAAAACGGTTTCAACCAGGCTATCGAACTTTCTGCCGAAGCGTTG
GCCAATGTCAAGTATGTTCAAGAAAAGAAATTATTGGAGGC**G**TATTTTGACGAAATT
TCCCAGGACACTGGTAAATTCTGTTATGGTATAGATGATACTTTAAAGGCATTGGAT
TTAGGTGCAGTCGAAAAATTAATTGTTTTCGAAAATTTGGAAACTATCAGATATACA
TTTAAAGATGCCGAGGATAATGAGGTTATAAAATTCGCTGAACCAGAAGCCAAGGA
CAAGTCGTTTGCTATTGACAAAGCTACCGGCCAAGAAATGGACGTTGTCTCCGAAGA
ACCTTTAATTGAATGGCTAGCAGCTAACTACAAAAACTTCGGTGCTACCTTGGAATT
CATCACAGACAAATCTTCAGAAGGTGCCCAATTTGTCACAGGTTTTGGTGGTATTGG
TGCCATGCTGCGTTACAAAGTTAATTTTGAACAACTAGTTGATGAATCTGAGGATGA
ATATTATGACGAAGATGAAGGATCCGACTATGATTTCATTTAAATAAATAAAAGGG
GGAGAAAAAAATCGAATCAAAAAGAATTTAATCACTAGATGCCAGATTTAAATTAA
ATTCGCTTTTAATTTTTTGTACAATATAATATATACTTGGTAAACCTTTTGCTCTATAT
TGAGCTAATTCCTTTGTTGAAAGTACATAGTGGGTTTAGAGGACCGTGTATATTACG
TAGAAAATACAGTGAAAGGAGAGTTTCTCTTCAAAAGCCTCGACGGTATCGATAAG
CTTATCGATACCGTCGACCTCGAGGGGGGGCCCGGTACCAGCTTTTGTTCCCTTTAG
TGAGGGTTAATTTCGAGCTTGGCGTAATCATGGTCATAGCTGTTTCCTGTGTGAAATT
GTTATCCGCTCACAATTCCACACAACATACGAGCCGGAAGCATAAAGTGTAAAGCC
TGGGGTGCCTAATGAGTGAGCTAACTCACATTAATTGCGTTGCGCTCACTGCCCGCT
TTCCAGTCGGGAAACCTGTCGTGCCAGCTGCATTAATGAATCGGCCAACGCGCGGG
GAGAGGCGGTTTGCGTATTGGGCGCTCTTCCGCTTCCTCGCTCACTGACTCGCTGCG
CTCGGTCGTTCGGCTGCGGCGAGCGGTATCAGCTCACTCAAAGGCGGTAATACGGTT
ATCCACAGAATCAGGGGATAACGCAGGAAAGAACATGTGAGCAAAAGGCCAGCAA
AAGGCCAGGAACCGTAAAAAGGCCGCGTTGCTGGCGTTTTTCCATAGGCTCCGCCCC
CCTGACGAGCATCACAAAAATCGACGCTCAAGTCAGAGGTGGCGAAACCCGACAGG
ACTATAAAGATACCAGGCGTTTCCCCCTGGAAGCTCCCTCGTGCGCTCTCCTGTTCC
GACCCTGCCGCTTACCGGATACCTGTCCGCCTTTCTCCCTTCGGGAAGCGTGGCGCT
TTCTCATAGCTCACGCTGTAGGTATCTCAGTTCGGTGTAGGTCGTTCGCTCCAAGCTG
GGCTGTGTGCACGAACCCCCCGTTCAGCCCGACCGCTGCGCCTTATCCGGTAACTAT
CGTCTTGAGTCCAACCCGGTAAGACACGACTTATCGCCACTGGCAGCAGCCACTGGT
AACAGGATTAGCAGAGCGAGGTATGTAGGCGGTGCTACAGAGTTCTTGAAGTGGTG

GCCTAACTACGGCTACACTAGAAGAACAGTATTTGGTATCTGCGCTCTGCTGAAGCC
AGTTACCTTCGGAAAAAGAGTTGGTAGCTCTTGATCCGGCAAACAAACCACCGCTG
GTAGCGGTGGTTTTTTTGTTTGCAAGCAGCAGATTACGCGCAGAAAAAAAGGATCTC
AAGAAGATCCTTTGATCTTTTCTACGGGGTCTGACGCTCAGTGGAACGAAAACTCAC
GTTAAGGGATTTTGGTCATGAGATTATCAAAAAGGATCTTCACCTAGATCCTTTTAA
ATTAAAAATGAAGTTTTAAATCAATCTAAAGTATATATGAGTAAACTTGGTCTGACA
GTTACCAATGCTTAATCAGTGAGGCACCTATCTCAGCGATCTGTCTATTTCGTTCATC
CATAGTTGCCTGACTCCCCGTCGTGTAGATAACTACGATACGGGAGGGCTTACCATC
TGGCCCCAGTGCTGCAATGATACCGCGAGACCCACGCTCACCGGCTCCAGATTTATC
AGCAATAAACCAGCCAGCCGGAAGGGCCGAGCGCAGAAGTGGTCCTGCAACTTTAT
CCGCCTCCATCCAGTCTATTAATTGTTGCCGGGAAGCTAGAGTAAGTAGTTCGCCAG
TTAATAGTTTGCGCAACGTTGTTGCCATTGCTACAGGCATCGTGGTGTCACGCTCGTC
GTTTGGTATGGCTTCATTCAGCTCCGGTTCCCAACGATCAAGGCGAGTTACATGATC
CCCCATGTTGTGCAAAAAAGCGGTTAGCTCCTTCGGTCCTCCGATCGTTGTCAGAAG
TAAGTTGGCCGCAGTGTTATCACTCATGGTTATGGCAGCACTGCATAATTCTCTTACT
GTCATGCCATCCGTAAGATGCTTTTCTGTGACTGGTGAGTACTCAACCAAGTCATTCT
GAGAATAGTGTATGCGGCGACCGAGTTGCTCTTGCCCGGCGTCAATACGGGATAAT
ACCGCGCCACATAGCAGAACTTTAAAAGTGCTCATCATTGGAAAACGTTCTTCGGGG
CGAAAACTCTCAAGGATCTTACCGCTGTTGAGATCCAGTTCGATGTAACC

**PTH353-SUP45-BR2.4**
*SUP45* Promoter *SUP45-**BR2.4***

CTCGTGCACCCAACTGATCTTCAGCATCTTTTACTTTCACCAGCGTTTCTGGGTGAGC
AAAAACAGGAAGGCAAAATGCCGCAAAAAAGGGAATAAGGGCGACACGGAAATGT
TGAATACTCATACTCTTCCTTTTTCAATATTATTGAAGCATTTATCAGGGTTATTGTC
TCATGAGCGGATACATATTTGAATGTATTTAGAAAAATAAACAAATAGGGGTTCCGC
GCACATTTCCCCGAAAAGTGCCACCTGGGTCCTTTTCATCACGTGCTATAAAAATAA
TTATAATTTAAATTTTTTAATATAAATATATAAATTAAAAATAGAAAGTAAAAAAAG
AAATTAAAGAAAAAATAGTTTTTGTTTTCCGAAGATGTAAAAGACTCTAGGGGGATC
GCCAACAAATACTACCTTTTATCTTGCTCTTCCTGCTCTCAGGTATTAATGCCGAATT
GTTTCATCTTGTCTGTGTAGAAGACCACACACGAAAATCCTGTGATTTTACATTTTAC
TTATCGTTAATCGAATGTATATCTATTTAATCTGCTTTTCTTGTCTAATAAATATATAT
GTAAAGTACGCTTTTTGTTGAAATTTTTTAAACCTTTGTTTATTTTTTTTTCTTCATTCC
GTAACTCTTCTACCTTCTTTATTTACTTTCTAAAATCCAAATACAAAACATAAAAATA
AATAAACACAGAGTAAATTCCCAAATTATTCCATCATTAAAAGATACGAGGCGCGT
GTAAGTTACAGGCAAGCGATCCGTCCTAAGAAACCATTATTATCATGACATTAACCT
ATAAAAATAGGCGTATCACGAGGCCCTTTCGTCTCGCGCGTTTCGGTGATGACGGTG
AAAACCTCTGACACATGCAGCTCCCGGAGACGGTCACAGCTTGTCTGTAAGCGGAT
GCCGGGAGCAGACAAGCCCGTCAGGGCGCGTCAGCGGGTGTTGGCGGGTGTCGGGG
CTGGCTTAACTATGCGGCATCAGAGCAGATTGTACTGAGAGTGCACCATATCGACTA
CGTCGTTAAGGCCGTTTCTGACAGAGTAAAATTCTTGAGGGAACTTTCACCATTATG
GGAAATGGTTCAAGAAGGTATTGACTTAAACTCCATCAAATGGTCAGGTCATTGAGT
GTTTTTTATTTGTTGTATTTTTTTTTTTTTAGAGAAAATCCTCCAATATATAAATTAGG
AATCATAGTTTCATGATTTTCTGTTACACCTAACTTTTTGTGTGGTGCCCTCCTCCTTG
TCAATATTAATGTTAAAGTGCAATTCTTTTTCCTTATCACGTTGAGCCATTAGTATCA

ATTTGCTTACCTGTATTCCTTTACATCCTCCTTTTTCTCCTTCTTGATAAATGTATGTA
GATTGCGTATATAGTTTCGTCTACCCTATGAACATATTCCATTTTGTAATTTCGTGTC
GTTTCTATTATGAATTTCATTTATAAAGTTTATGTACAAATATCATAAAAAAAGAGA
ATCTTTTTAAGCAAGGATTTTCTTAACTTCTTCGGCGACAGCATCACCGACTTCGGTG
GTACTGTTGGAACCACCTAAATCACCAGTTCTGATACCTGCATCCAAAACCTTTTTA
ACTGCATCTTCAATGGCCTTACCTTCTTCAGGCAAGTTCAATGACAATTTCAACATCA
TTGCAGCAGACAAGATAGTGGCGATAGGGTTGACCTTATTCTTTGGCAAATCTGGAG
CAGAACCGTGGCATGGTTCGTACAAACCAAATGCGGTGTTCTTGTCTGGCAAAGAG
GCCAAGGACGCAGATGGCAACAAACCCAAGGAACCTGGGATAACGGAGGCTTCATC
GGAGATGATATCACCAAACATGTTGCTGGTGATTATAATACCATTTAGGTGGGTTGG
GTTCTTAACTAGGATCATGGCGGCAGAATCAATCAATTGATGTTGAACCTTCAATGT
AGGGAATTCGTTCTTGATGGTTTCCTCCACAGTTTTTCTCCATAATCTTGAAGAGGCC
AAAACATTAGCTTTATCCAAGGACCAAATAGGCAATGGTGGCTCATGTTGTAGGGCC
ATGAAAGCGGCCATTCTTGTGATTCTTTGCACTTCTGGAACGGTGTATTGTTCACTAT
CCCAAGCGACACCATCACCATCGTCTTCCTTTCTCTTACCAAAGTAAATACCTCCCA
CTAATTCTCTGACAACAACGAAGTCAGTACCTTTAGCAAATTGTGGCTTGATTGGAG
ATAAGTCTAAAAGAGAGTCGGATGCAAAGTTACATGGTCTTAAGTTGGCGTACAATT
GAAGTTCTTTACGGATTTTTAGTAAACCTTGTTCAGGTCTAACACTACCGGTACCCCA
TTTAGGACCACCCACAGCACCTAACAAAACGGCATCAGCCTTCTTGGAGGCTTCCAG
CGCCTCATCTGGAAGTGGAACACCTGTAGCATCGATAGCAGCACCACCAATTAAAT
GATTTTCGAAATCGAACTTGACATTGGAACGAACATCAGAAATAGCTTTAAGAACCT
TAATGGCTTCGGCTGTGATTTCTTGACCAACGTGGTCACCTGGCAAAACGACGATCT
TCTTAGGGGCAGACATTAGAATGGTATATCCTTGAAATATATATATATATATTGCTG
AAATGTAAAAGGTAAGAAAAGTTAGAAAGTAAGACGATTGCTAACCACCTATTGGA
AAAAACAATAGGTCCTTAAATAATATTGTCAACTTCAAGTATTGTGATGCAAGCATT
TAGTCATGAACGCTTCTCTATTCTATATGAAAAGCCGGTTCCGGCGCTCTCACCTTTC
CTTTTTCTCCCAATTTTTCAGTTGAAAAAGGTATATGCGTCAGGCGACCTCTGAAATT
AACAAAAAATTTCCAGTCATCGAATTTGATTCTGTGCGATAGCGCCCCTGTGTGTTC
TCGTTATGTTGAGGAAAAAAATAATGGTTGCTAAGAGATTCGAACTCTTGCATCTTA
CGATACCTGAGTATTCCCACAGTTAACTGCGGTCAAGATATTTCTTGAATCAGGCGC
CTTAGACCGCTCGGCCAAACAACCAATTACTTGTTGAGAAATAGAGTATAATTATCC
TATAAATATAACGTTTTTGAACACACATGAACAAGGAAGTACAGGACAATTGATTTT
GAAGAGAATGTGGATTTTGATGTAATTGTTGGGATTCCATTTTTAATAAGGCAATAA
TATTAGGTATGTAGATATACTAGAAGTTCTCCTCGACCGGTCGATATGCGGTGTGAA
ATACCGCACAGATGCGTAAGGAGAAAATACCGCATCAGGAAATTGTAAGCGTTAAT
ATTTTGTTAAAATTCGCGTTAAATTTTTGTTAAATCAGCTCATTTTTTAACCAATAGG
CCGAAATCGGCAAAATCCCTTATAAATCAAAAGAATAGACCGAGATAGGGTTGAGT
GTTGTTCCAGTTTGGAACAAGAGTCCACTATTAAAGAACGTGGACTCCAACGTCAAA
GGGCGAAAAACCGTCTATCAGGGCGATGGCCCACTACGTGAACCATCACCCTAATC
AAGTTTTTTGGGGTCGAGGTGCCGTAAAGCACTAAATCGGAACCCTAAAGGGAGCC
CCCGATTTAGAGCTTGACGGGGAAAGCCGGCGAACGTGGCGAGAAAGGAAGGGAA
GAAAGCGAAAGGAGCGGGCGCTAGGGCGCTGGCAAGTGTAGCGGTCACGCTGCGC
GTAACCACCACACCCGCCGCGCTTAATGCGCCGCTACAGGGCGCGTCCATTCGCCAT
TCAGGCTGCGCAACTGTTGGGAAGGGCGATCGGTGCGGGCCTCTTCGCTATTACGCC
AGCTGGCGAAAGGGGGATGTGCTGCAAGGCGATTAAGTTGGGTAACGCCAGGGTTT
TCCCAGTCACGACGTTGTAAAACGACGGCCAGTGAATTGTAATACGACTCACTATAG

GGCGAATTGGAGCTCCACCGCGGTGGCGGCCGCTCTAGACAATACGAAGGAACGAT
CCCGCGTCAGCAGATGTCACAATTTGATTATTTCATTGTCACATGCTTTTTGACTCAT
CCTTCCGCGAAAGGAGAACTTTTTACAATGCCGCCGGCTCTGTTAGCGTACCCTTCT
ACAAGCAGATAGCAGAACAAACACATGATATATTCAAAAGGTGCAATGTCAGAGAA
CATATATTGCGCCCCTGTCCTGTAGACATCAGTCATTTTTCGCGGGACTTGAATGGC
GCACCATTATCCTATACAGAAAACATTAAATTTACTGCAAAATTTTGGGCAAACGCT
TGGATAGACTATGAATCCGTCATGAAGACACTACTTGTAAAATTATATATATCCTTT
TTTTTCATGTGCTAGTAAGAATTAGCAACTACTTTCATTCATCTTCGACGCAACTTCG
AGGAAAGCACCTTTTTACTTGCTGTAGCTCTATTCTCTTCCCCAACCACCTTTTCCTTT
ATTCCAAAATTTTTAAAACTTTTTCTGTTACATTATATAATCTTCTGTCTGAAATGTTT
GGATATAACGCCTCTTGATCCACTTTGTATATGCGTGCTATTTATTTTCAGATTTATA
AAGAGTATGAGCGTCATTTACATAAATAGCTGAAGTTATTCATGGAAAATACGAAG
AGCACGTATGTGAGCCAACAGAACATTTGACGTAAGACTCTACAATGTGCCAAGAA
CTGGACAAGTAGAGGACTGAGAACTTTATTTCAATTCATTGCTCCTTTTTGGTGGCG
CTACCTTTAGCGAAGGTCAATGATGAATGTGCACATGCTGTCGAAACCAAAAAGCA
AATTCTAACCAACTTCAAAATGACATAGTCATCTGATATTTCTACTCATTATAGATA
GTATGGGAGCCTTGAAACGAAAAGTAAGTAAAAAGCTGGATATGAGCAGTATGAGG
TAGACCTTAGCTACATCATTTCCCCCAATAGCTGCTGCAAATATCTGGTTAAATTTGT
GATTCCATGAAGAGGATAACAGACTTGTTAAAAAGCATCCTGTCAAAATCTAATTTT
TGAAGGGCAGTATTCAATTCATAATTTACTTTAGCTTAGATCCTTCCAATTTATACAT
GGTATTATAACCAGATCATAAACTACAATCTGTCGCTACCGCATGTACGAAGAATAC
TTAAGTCACTTGCTCTCTCATCATTTGTACATTTTTCAGTAATACCGTTTGATAGCGC
CGTCTTTATTACCCGGATTATTCCGTTGACCCTGAATGAAAAATTTTTTCAGAAATCC
AGTGCTAAGCGTCAAATCAATGAAATACATCACTGTATTTTTAACTGATATACTGTT
GGTGTGGCCTTAACGACACCTTTATTTCTTAATTCATTTCGGCTTGTCTCCTTATTAA
GACTACAGAAATAGACAAAGGAAATACTTCAATAATGGATAACGAGGTTGAAAAAA
ATATTGAGATCTGGAAGGTCAAGAAGTTGGTCCAATCTTTAGAAAAAGCTAGAGGT
AATGGTACT**C**CTATGATTTCCTTAGTTATTCCTCCTAAGGGTCAAATTCCACTGTACC
AAAAAATGTTAACAGATGAATATGGTACTGCCTCGAATATTAAATCTAGGGTTAATC
GTCTTTCCGTTTTATCTGCTATCACTTCCACCCAACAAAAGTTGAAGCTATATAATAC
TTTGCCCAAGAACGGTTTAGTTTTATATTGTGGTGATATCATCACTGAAGATGGTAA
AGAAAAAAAGGTCACTTTTGACATCGAACCTTACAAACCTATCAACACATCCTTATA
TTTGTGTGATAACAAATTTCATACAGAAGTTCTTTCGGAATTGCTTCAAGCTGACGA
CAAGTTCGGTTTTATAGTCATGGACGGTCAAGGTACTTTGTTTGGTTCTGTGTCCGGT
AATACGAGAACTGTTTTACATAAATTTACTGTCGATCTGCCAAAAAAGCATGGTAGA
GGTGGTCAATCTGCGCTTCGTTTTGCTCGTTTAAGAGAAGAAAAAAGACATAATTAT
GTGAGAAAGGTCGCCGAAGTTGCTGTTCAAAATTTTATTACTAATGACAAAGTCAAT
GTTAAGGGTTTA**C**TTTTAGCTGGTTCTGCTGACTTTAAGACCGATTTGGCTAAATCTG
AATTATTCGATCCAAGACTAGCATGTAAGGTTATTTCCATCGTGGATGTTTCTTATGG
TGGTGAAAACGGTTTCAACCAGGCTATCGAACTTTCTGCCGAAGCGTTGGCCAATGT
CAAGTATGTTCAAGAAAAGAAATTATTGGAGGCATATTTTGACGAAATTTCCCAGGA
CACTGGTAAATTCTGTTATGGTATAGATGATACTTTAAAGGCATTGGAT**C**TAGGTGC
AGTCGAAAAATTAATTGTTTTCGAAAATTTGGAAACTATCAGATATACATTTAAAGA
TGCCGAGGATAATGAGGTTATAAAATTCGCTGAACCAGAAGCCAAGGACAAGTCGT
TTGCTATTGACAAAGCTACCGGCCAAGAAATGGACGTTGTCTCCGAAGAACCTTTAA
TTGAATGGCTAGCAGCTAACTACAAAAACTTCGGTGCTACCTTGGAATTCATCACAG

ACAAATCTTCAGAAGGTGCCCAATTTGTCACAGGTTTTGGTGGTATTGGTGCCATGC
TGCGTTACAAAGTTAATTTTGAACAACTAGTTGATGAATCTGAGGATGAATATTATG
ACGAAGATGAAGGATCCGACTATGATTTCATTTAAATAAATAAAAGGGGGAGAAAA
AAATCGAATCAAAAAGAATTTAATCACTAGATGCCAGATTTAAATTAAATTCGCTTT
TAATTTTTTGTACAATATAATATATACTTGGTAAACCTTTTGCTCTATATTGAGCTAA
TTCCTTTGTTGAAAGTACATAGTGGGTTTAGAGGACCGTGTATATTACGTAGAAAAT
ACAGTGAAAGGAGAGTTTCTCTTCAAAAGCCTCGACGGTATCGATAAGCTTATCGAT
ACCGTCGACCTCGAGGGGGGGCCCGGTACCAGCTTTTGTTCCCTTTAGTGAGGGTTA
ATTTCGAGCTTGGCGTAATCATGGTCATAGCTGTTTCCTGTGTGAAATTGTTATCCGC
TCACAATTCCACACAACATACGAGCCGGAAGCATAAAGTGTAAAGCCTGGGGTGCC
TAATGAGTGAGCTAACTCACATTAATTGCGTTGCGCTCACTGCCCGCTTTCCAGTCG
GGAAACCTGTCGTGCCAGCTGCATTAATGAATCGGCCAACGCGCGGGGAGAGGCGG
TTTGCGTATTGGGCGCTCTTCCGCTTCCTCGCTCACTGACTCGCTGCGCTCGGTCGTT
CGGCTGCGGCGAGCGGTATCAGCTCACTCAAAGGCGGTAATACGGTTATCCACAGA
ATCAGGGGATAACGCAGGAAAGAACATGTGAGCAAAAGGCCAGCAAAAGGCCAGG
AACCGTAAAAAGGCCGCGTTGCTGGCGTTTTTCCATAGGCTCCGCCCCCCTGACGAG
CATCACAAAAATCGACGCTCAAGTCAGAGGTAGCGAAACCCGACAGGACTATAAAG
ATACCAGGCGTTTCCCCCTGGAAGCTCCCTCGTGCGCTCTCCTGTTCCGACCCTGCCG
CTTACCGGATACCTGTCCGCCTTTCTCCCTTCGGGAAGCGTGGCGCTTTCTCATAGCT
CACGCTGTAGGTATCTCAGTTCGGTGTAGGTCGTTCGCTCCAAGCTGGGCTGTGTGC
ACGAACCCCCCGTTCAGCCCGACCGCTGCGCCTTATCCGGTAACTATCGTCTTGAGT
CCAACCCGGTAAGACACGACTTATCGCCACTGGCAGCAGCCACTGGTAACAGGATT
AGCAGAGCGAGGTATGTAGGCGGTGCTACAGAGTTCTTGAAGTGGTGGCCTAACTA
CGGCTACACTAGAAGAACAGTATTTGGTATCTGCGCTCTGCTGAAGCCAGTTACCTT
CGGAAAAAGAGTTGGTAGCTCTTGATCCGGCAAACAAACCACCGCTGGTAGCGGTG
GTTTTTTTGTTTGCAAGCAGCAGATTACGCGCAGAAAAAAAGGATCTCAAGAAGATC
CTTTGATCTTTTCTACGGGGTCTGACGCTCAGTGGAACGAAAACTCACGTTAAGGGA
TTTTGGTCATGAGATTATCAAAAAGGATCTTCACCTAGATCCTTTTAAATTAAAAAT
GAAGTTTTAAATCAATCTAAAGTATATATGAGTAAACTTGGTCTGACAGTTACCAAT
GCTTAATCAGTGAGGCACCTATCTCAGCGATCTGTCTATTTCGTTCATCCATAGTTGC
CTGACTCCCCGTCGTGTAGATAACTACGATACGGGAGGGCTTACCATCTGGCCCCAG
TGCTGCAATGATACCGCGAGACCCACGCTCACCGGCTCCAGATTTATCAGCAATAAA
CCAGCCAGCCGGAAGGGCCGAGCGCAGAAGTGGTCCTGCAACTTTATCCGCCTCCA
TCCAGTCTATTAATTGTTGCCGGGAAGCTAGAGTAAGTAGTTCGCCAGTTAATAGTT
TGCGCAACGTTGTTGCCATTGCTACAGGCATCGTGGTGTCACGCTCGTCGTTTGGTAT
GGCTTCATTCAGCTCCGGTTCCCAACGATCAAGGCGAGTTACATGATCCCCCATGTT
GTGCAAAAAAGCGGTTAGCTCCTTCGGTCCTCCGATCGTTGTCAGAAGTAAGTTGGC
CGCAGTGTTATCACTCATGGTTATGGCAGCACTGCATAATTCTCTTACTGTCATGCCA
TCCGTAAGATGCTTTTCTGTGACTGGTGAGTACTCAACCAAGTCATTCTGAGAATAG
TGTATGCGGCGACCGAGTTGCTCTTGCCCGGCGTCAATACGGGATAATACCGCGCCA
CATAGCAGAACTTTAAAAGTGCTCATCATTGGAAAACGTTCTTCGGGGCGAAAACTC
TCAAGGATCTTACCGCTGTTGAGATCCAGTTCGATGTAACCCA

**PTH353-SUP45-BR2.9**

*SUP45* Promoter *SUP45-**BR2.9***

CAGTGTTATCACTCATGGTTATGGCAGCACTGCATAATTCTCTTACTGTCATGCCATC
CGTAAGATGCTTTTCTGTGACTGGTGAGTACTCAACCAAGTCATTCTGAGAATAGTG
TATGCGGCGACCGAGTTGCTCTTGCCCGGCGTCAATACGGGATAATACCGCGCCACA
TAGCAGAACTTTAAAAGTGCTCATCATTGGAAAACGTTCTTCGGGGCGAAAACTCTC
AAGGATCTTACCGCTGTTGAGATCCAGTTCGATGTAACCCACTCGTGCACCCAACTG
ATCTTCAGCATCTTTTACTTTCACCAGCGTTTCTGGGTGAGCAAAAACAGGAAGGCA
AAATGCCGCAAAAAAGGGAATAAGGGCGACACGGAAATGTTGAATACTCATACTCT
TCCTTTTTCAATATTATTGAAGCATTTATCAGGGTTATTGTCTCATGAGCGGATACAT
ATTTGAATGTATTTAGAAAAATAAACAAATAGGGGTTCCGCGCACATTTCCCCGAAA
AGTGCCACCTGGGTCCTTTTCATCACGTGCTATAAAAATAATTATAATTTAAATTTTT
TAATATAAATATATAAATTAAAAATAGAAAGTAAAAAAAGAAATTAAAGAAAAAAT
AGTTTTTGTTTTCCGAAGATGTAAAAGACTCTAGGGGGATCGCCAACAAATACTACC
TTTTATCTTGCTCTTCCTGCTCTCAGGTATTAATGCCGAATTGTTTCATCTTGTCTGTG
TAGAAGACCACACACGAAAATCCTGTGATTTTACATTTTACTTATCGTTAATCGAAT
GTATATCTATTTAATCTGCTTTTCTTGTCTAATAAATATATATGTAAAGTACGCTTTTT
GTTGAAATTTTTTAAACCTTTGTTTATTTTTTTTTCTTCATTCCGTAACTCTTCTACCTTC
TTTATTTACTTTCTAAAATCCAAATACAAAACATAAAAATAAATAAACACAGAGTAA
ATTCCCAAATTATTCCATCATTAAAAGATACGAGGCGCGTGTAAGTTACAGGCAAGC
GATCCGTCCTAAGAAACCATTATTATCATGACATTAACCTATAAAAATAGGCGTATC
ACGAGGCCCTTTCGTCTCGCGCGTTTCGGTGATGACGGTGAAAACCTCTGACACATG
CAGCTCCCGGAGACGGTCACAGCTTGTCTGTAAGCGGATGCCGGGAGCAGACAAGC
CCGTCAGGGCGCGTCAGCGGGTGTTGGCGGGTGTCGGGGCTGGCTTAACTATGCGG
CATCAGAGCAGATTGTACTGAGAGTGCACCATATCGACTACGTCGTTAAGGCCGTTT
CTGACAGAGTAAAATTCTTGAGGGAACTTTCACCATTATGGGAAATGGTTCAAGAA
GGTATTGACTTAAACTCCATCAAATGGTCAGGTCATTGAGTGTTTTTTATTTGTTGTA
TTTTTTTTTTTTTAGAGAAAATCCTCCAATATATAAATTAGGAATCATAGTTTCATGA
TTTTCTGTTACACCTAACTTTTTGTGTGGTGCCCTCCTCCTTGTCAATATTAATGTTAA
AGTGCAATTCTTTTTCCTTATCACGTTGAGCCATTAGTATCAATTTGCTTACCTGTAT
TCCTTTACATCCTCCTTTTTCTCCTTCTTGATAAATGTATGTAGATTGCGTATATAGTT
TCGTCTACCCTATGAACATATTCCATTTTGTAATTTCGTGTCGTTTCTATTATGAATTT
CATTTATAAAGTTTATGTACAAATATCATAAAAAAAGAGAATCTTTTTAAGCAAGGA
TTTTCTTAACTTCTTCGGCGACAGCATCACCGACTTCGGTGGTACTGTTGGAACCACC
TAAATCACCAGTTCTGATACCTGCATCCAAAACCTTTTTAACTGCATCTTCAATGGCC
TTACCTTCTTCAGGCAAGTTCAATGACAATTTCAACATCATTGCAGCAGACAAGATA
GTGGCGATAGGGTTGACCTTATTCTTTGGCAAATCTGGAGCAGAACCGTGGCATGGT
TCGTACAAACCAAATGCGGTGTTCTTGTCTGGCAAAGAGGCCAAGGACGCAGATGG
CAACAAACCCAAGGAACCTGGGATAACGGAGGCTTCATCGGAGATGATATCACCAA
ACATGTTGCTGGTGATTATAATACCATTTAGGTGGGTTGGGTTCTTAACTAGGATCA
TGGCGGCAGAATCAATCAATTGATGTTGAACCTTCAATGTAGGGAATTCGTTCTTGA
TGGTTTCCTCCACAGTTTTTCTCCATAATCTTGAAGAGGCCAAAACATTAGCTTTATC
CAAGGACCAAATAGGCAATGGTGGCTCATGTTGTAGGGCCATGAAAGCGGCCATTC
TTGTGATTCTTTGCACTTCTGGAACGGTGTATTGTTCACTATCCCAAGCGACACCATC
ACCATCGTCTTCCTTTCTCTTACCAAAGTAAATACCTCCCACTAATTCTCTGACAACA
ACGAAGTCAGTACCTTTAGCAAATTGTGGCTTGATTGGAGATAAGTCTAAAAGAGA
GTCGGATGCAAAGTTACATGGTCTTAAGTTGGCGTACAATTGAAGTTCTTTACGGAT
TTTTAGTAAACCTTGTTCAGGTCTAACACTACCGGTACCCCATTTAGGACCACCCAC

AGCACCTAACAAAACGGCATCAGCCTTCTTGGAGGCTTCCAGCGCCTCATCTGGAAG
TGGAACACCTGTAGCATCGATAGCAGCACCACCAATTAAATGATTTTCGAAATCGAA
CTTGACATTGGAACGAACATCAGAAATAGCTTTAAGAACCTTAATGGCTTCGGCTGT
GATTTCTTGACCAACGTGGTCACCTGGCAAAACGACGATCTTCTTAGGGGCAGACAT
TAGAATGGTATATCCTTGAAATATATATATATATATTGCTGAAATGTAAAAGGTAAG
AAAAGTTAGAAAGTAAGACGATTGCTAACCACCTATTGGAAAAAACAATAGGTCCT
TAAATAATATTGTCAACTTCAAGTATTGTGATGCAAGCATTTAGTCATGAACGCTTC
TCTATTCTATATGAAAAGCCGGTTCCGGCGCTCTCACCTTTCCTTTTTCTCCCAATTTT
TCAGTTGAAAAAGGTATATGCGTCAGGCGACCTCTGAAATTAACAAAAAATTTCCA
GTCATCGAATTTGATTCTGTGCGATAGCGCCCCTGTGTGTTCTCGTTATGTTGAGGAA
AAAAATAATGGTTGCTAAGAGATTCGAACTCTTGCATCTTACGATACCTGAGTATTC
CCACAGTTAACTGCGGTCAAGATATTTCTTGAATCAGGCGCCTTAGACCGCTCGGCC
AAACAACCAATTACTTGTTGAGAAATAGAGTATAATTATCCTATAAATATAACGTTT
TTGAACACACATGAACAAGGAAGTACAGGACAATTGATTTTGAAGAGAATGTGGAT
TTTGATGTAATTGTTGGGATTCCATTTTTAATAAGGCAATAATATTAGGTATGTAGAT
ATACTAGAAGTTCTCCTCGACCGGTCGATATGCGGTGTGAAATACCGCACAGATGCG
TAAGGAGAAAATACCGCATCAGGAAATTGTAAGCGTTAATATTTTGTTAAAATTCGC
GTTAAATTTTTGTTAAATCAGCTCATTTTTTAACCAATAGGCCGAAATCGGCAAAAT
CCCTTATAAATCAAAAGAATAGACCGAGATAGGGTTGAGTGTTGTTCCAGTTTGGAA
CAAGAGTCCACTATTAAAGAACGTGGACTCCAACGTCAAAGGGCGAAAAACCGTCT
ATCAGGGCGATGGCCCACTACGTGAACCATCACCCTAATCAAGTTTTTTGGGGTCGA
GGTGCCGTAAAGCACTAAATCGGAACCCTAAAGGGAGCCCCCGATTTAGAGCTTGA
CGGGGAAAGCCGGCGAACGTGGCGAGAAAGGAAGGGAAGAAAGCGAAAGGAGCG
GGCGCTAGGGCGCTGGCAAGTGTAGCGGTCACGCTGCGCGTAACCACCACACCCGC
CGCGCTTAATGCGCCGCTACAGGGCGCGTCCATTCGCCATTCAGGCTGCGCAACTGT
TGGGAAGGGCGATCGGTGCGGGCCTCTTCGCTATTACGCCAGCTGGCGAAAGGGGG
ATGTGCTGCAAGGCGATTAAGTTGGGTAACGCCAGGGTTTTCCCAGTCACGACGTTG
TAAAACGACGGCCAGTGAATTGTAATACGACTCACTATAGGGCGAATTGGAGCTCC
ACCGCGGTGGCGGCCGCTCTAGACAATACGAAGGAACGATCCCGCGTCAGCAGATG
TCACAATTTGATTATTTCATTGTCACATGCTTTTTGACTCATCCTTCCGCGAAAGGAG
AACTTTTTACAATGCCGCCGGCTCTGTTAGCGTACCCTTCTACAAGCAGATAGCAGA
ACAAACACATGATATATTCAAAAGGTGCAATGTCAGAGAACATATATTGCGCCCCT
GTCCTGTAGACATCAGTCATTTTTCGCGGGACTTGAATGGCGCACCATTATCCTATA
CAGAAAACATTAAATTTACTGCAAAATTTTGGGCAAACGCTTGGATAGACTATGAAT
CCGTCATGAAGACACTACTTGTAAAATTATATATATCCTTTTTTTTCATGTGCTAGTA
AGAATTAGCAACTACTTTCATTCATCTTCGACGCAACTTCGAGGAAAGCACCTTTTT
ACTTGCTGTAGCTCTATTCTCTTCCCCAACCACCTTTTCCTTTATTCCAAAATTTTTAA
AACTTTTTCTGTTACATTATATAATCTTCTGTCTGAAATGTTTGGATATAACGCCTCT
TGATCCACTTTGTATATGCGTGCTATTTATTTTCAGATTTATAAAGAGTATGAGCGTC
ATTTACATAAATAGCTGAAGTTATTCATGGAAAATACGAAGAGCACGTATGTGAGC
CAACAGAACATTTGACGTAAGACTCTACAATGTGCCAAGAACTGGACAAGTAGAGG
ACTGAGAACTTTATTTCAATTCATTGCTCCTTTTTGGTGGCGCTACCTTTAGCGAAGG
TCAATGATGAATGTGCACATGCTGTCGAAACCAAAAAGCAAATTCTAACCAACTTCA
AAATGACATAGTCATCTGATATTTCTACTCATTATAGATAGTATGGGAGCCTTGAAA
CGAAAAGTAAGTAAAAAGCTGGATATGAGCAGTATGAGGTAGACCTTAGCTACATC
ATTTCCCCCAATAGCTGCTGCAAATATCTGGTTAAATTTGTGATTCCATGAAGAGGA

TAACAGACTTGTTAAAAAGCATCCTGTCAAAATCTAATTTTTGAAGGGCAGTATTCA
ATTCATAATTTACTTTAGCTTAGATCCTTCCAATTTATACATGGTATTATAACCAGAT
CATAAACTACAATCTGTCGCTACCGCATGTACGAAGAATACTTAAGTCACTTGCTCT
CTCATCATTTGTACATTTTTCAGTAATACCGTTTGATAGCGCCGTCTTTATTACCCGG
ATTATTCCGTTGACCCTGAATGAAAAATTTTTTCAGAAATCCAGTGCTAAGCGTCAA
ATCAATGAAATACATCACTGTATTTTTAACTGATATACTGTTGGTGTGGCCTTAACG
ACACCTTTATTTCTTAATTCATTTCGGCTTGTCTCCTTATTAAGACTACAGAAATAGA
CAAAGGAAATACTTCAATAATGGATAACGAGGTTGAAAAAAATATTGAGATCTGGA
AGGTCAAGAAGTTGGTCCAATCTTTAGAAAAAGCTAGAGGTAATGGTACTTCTATGA
TTTCCTTAGTTATTCCTCCTAAGGGTCAAATTCCACTGTACCAAAAAATGTTAACAG
ATGAATATGGTACTGCCTCGAATATTAAATCTAGGGTTAATCGTCTTTCCGTTT**C**A**C**C
TGCTATCACTTCCACCCAACAAAAG**C**TGAAGCTATATAATACTTTGCCCAAGAACGG
TTTAGTTTTATATTGTGGTGATATCATCACTGAAGATGGTAAAGAAAAAAAGGTCAC
TTTTGACATCGAACCTTACAAACCTATCAACACATCCTTATATTTGTGTGATAACAA
ATTTCATACAGAAGTTCTTTCGGAATTGCTTCAAGCTGACGACAAGTTCGGTTTTATA
GTCATGGACGGTCAAGGTACTTTGTTTGGTTCTGTGTCCGGTAATACGAGAACTGTT
TTACATAAATTTACTGTCGATCTGCCAAAAAAGCATGGTAGAGGTGGTCAATCTGCG
CTTCGTTTTGCTCGTTTAAGAGAAGAAAAAAGACATAATTATGTGAGAAAGGTCGCC
GAAGTTGCTGTTCAAAATTTTATTACTAATGACAAAGTCAATGTTAAGGGTTTAATT
TTAGCTGGTTCTGCTGACTTTAAGACCGATTTGGCTAAATCTGAATTATTCGATCCAA
GACTAGCATGTAAGGTTATTTCCATCGTGGATGTTTCTTATGGTGGTGAAAACGGTT
TCAACCAGGCTATCGAACTTTCTGCCGAAGCGTTGGCCAATGTCAAGTATGTTCAAG
**G**AAAGAAATTATTGGAGGCATATTTTGACGAAATTTCCCAGGACACTGGTAAATTCT
GTTATGGTATAGATGATACTTTAAAGGCATTGGATTTAGGTGCAGTCGAAAAATTAA
TTGTTTTCGAAAATTTGGAAACTATCAGATATACATTTAAAGATGCCGAGGATAATG
AGGTTATAAAATTCGCTGAACCAGAAGCCAAGGACAAGTCGTTTGCTATTGACAAA
GCTACCGGCCAAGAAATGGACGTTGTCTCCGAAGAACCTTTAATTGAATGGCTAGCA
GCTAACTACAAAAACTTCGGTGCTACCTTGGAATTCATCACAGACAAATCTTCAGAA
GGTGCCCAATTTGTCACAGGTTTTGGTGGTATTGGTGCCATGCTGCGTTACAAAG**C**T
AATTTTGAACAACTAGTTGATGAATCTGAGGATGAATATTATGACGAAGATGAAGG
ATCCGACTATGATTTCATTTAAATAAATAAAAGGGGGAGAAAAAAATCGAATCAAA
AAGAATTTAATCACTAGATGCCAGATTTAAATTAAATTCGCTTTTAATTTTTGTACA
ATATAATATATACTTGGTAAACCTTTTGCTCTATATTGAGCTAATTCCTTTGTTGAAA
GTACATAGTGGGTTTAGAGGACCGTGTATATTACGTAGAAAATACAGTGAAAGGAG
AGTTTCTCTTCAAAAGCCTCGACGGTATCGATAAGCTTATCGATACCGTCGACCTCG
AGGGGGGGCCCGGTACCAGCTTTTGTTCCCTTTAGTGAGGGTTAATTTCGAGCTTGG
CGTAATCATGGTCATAGCTGTTTCCTGTGTGAAATTGTTATCCGCTCACAATTCCACA
CAACATACGAGCCGGAAGCATAAAGTGTAAAGCCTGGGGTGCCTAATGAGTGAGCT
AACTCACATTAATTGCGTTGCGCTCACTGCCCGCTTTCCAGTCGGGAAACCTGTCGT
GCCAGCTGCATTAATGAATCGGCCAACGCGCGGGGAGAGGCGGTTTGCGTATTGGG
CGCTCTTCCGCTTCCTCGCTCACTGACTCGCTGCGCTCGGTCGTTCGGCTGCGGCGAG
CGGTATCAGCTCACTCAAAGGCGGTAATACGGTTATCCACAGAATCAGGGGATAAC
GCAGGAAAGAACATGTGAGCAAAAGGCCAGCAAAAGGCCAGGAACCGTAAAAAGG
CCGCGTTGCTGGCGTTTTTCCATAGGCTCCGCCCCCCTGACGAGCATCACAAAAATC
GACGCTCAAGTCAGAGGTGGCGAAACCCGACAGGACTATAAAGATACCAGGCGTTT
CCCCCTGGAAGCTCCCTCGTGCGCTCTCCTGTTCCGACCCTGCCGCTTACCGGATACC

TGTCCGCCTTTCTCCCTTCGGGAAGCGTGGCGCTTTCTCATAGCTCACGCTGTAGGTA
TCTCAGTTCGGTGTAGGTCGTTCGCTCCAAGCTGGGCTGTGTGCACGAACCCCCCGT
TCAGCCCGACCGCTGCGCCTTATCCGGTAACTATCGTCTTGAGTCCAACCCGGTAAG
ACACGACTTATCGCCACTGGCAGCAGCCACTGGTAACAGGATTAGCAGAGCGAGGT
ATGTAGGCGGTGCTACAGAGTTCTTGAAGTGGTGGCCTAACTACGGCTACACTAGAA
GAACAGTATTTGGTATCTGCGCTCTGCTGAAGCCAGTTACCTTCGGAAAAAGAGTTG
GTAGCTCTTGATCCGGCAAACAAACCACCGCTGGTAGCGGTGGTTTTTTTGTTTGCA
AGCAGCAGATTACGCGCAGAAAAAAAGGATCTCAAGAAGATCCTTTGATCTTTTCTA
CGGGGTCTGACGCTCAGTGGAACGAAAACTCACGTTAAGGGATTTTGGTCATGAGA
TTATCAAAAAGGATCTTCACCTAGATCCTTTTAAATTAAAAATGAAGTTTTAAATCA
ATCTAAAGTATATATGAGTAAACTTGGTCTGACAGTTACCAATGCTTAATCAGTGAG
GCACCTATCTCAGCGATCTGTCTATTTCGTTCATCCATAGTTGCCTGACTCCCCGTCG
TGTAGATAACTACGATACGGGAGGGCTTACCATCTGGCCCCAGTGCTGCAATGATAC
CGCGAGACCCACGCTCACCGGCTCCAGATTTATCAGCAATAAACCAGCCAGCCGGA
AGGGCCGAGCGCAGAAGTGGTCCTGCAACTTTATCCGCCTCCATCCAGTCTATTAAT
TGTTGCCGGGAAGCTAGAGTAAGTAGTTCGCCAGTTAATAGTTTGCGCAACGTTGTT
GCCATTGCTACAGGCATCGTGGTGTCACGCTCGTCGTTTGGTATGGCTTCATTCAGCT
CCGGTTCCCAACGATCAAGGCGAGTTACATGATCCCCCATGTTGTGCAAAAAAGCG
GTTAGCTCCTTCGGTCCTCCGATCGTTGTCAGAAGTAAGTTGGCCG

**PTH353-SUP45-BR3T.4**
*SUP45* Promoter *SUP45*-***BR3T.4***

CTCGTGCACCCAACTGATCTTCAGCATCTTTTACTTTCACCAGCGTTTCTGGGTGAGC
AAAAACAGGAAGGCAAAATGCCGCAAAAAAGGGAATAAGGGCGACACGGAAATGT
TGAATACTCATACTCTTCCTTTTTCAATATTATTGAAGCATTTATCGGGGTTATTGTC
TCATGAGCGGATACATATTTGAATGTATTTAGAAAAATAAACAAATAGGGGGTTCC
GCGCACATTTCCCCGAAAAGTGCCACCTGGGTCCTTTTCATCACGTGCTATAAAAAT
AATTATAATTTAAATTTTTTAATATAAATATATAAATTAAAAATAGAAAGTAAAAAA
AGAAATTAAAGAAAAAATAGTTTTTGTTTTCCGAAGATGTAAAAGACTCTAGGGGG
ATCGCCAACAAATACTACCTTTTATCTTGCTCTTCCTGCTCTCAGGTATTAATGCCGA
ATTGTTTCATCTTGTCTGTGTAGAAGACCACACACGAAAATCCTGTGATTTTACATTT
TACTTATCGTTAATCGAATGTATATCTATTTAATCTGCTTTTCTTGTCTAATAAATAT
ATATGTAAAGTACGCTTTTTGTTGAAATTTTTTAAACCTTTGTTTATTTTTTTTTCTTC
ATTCCGTAACTCTTCTACCTTCTTTATTTACTTTCTAAAATCCAAATACAAAACATAA
AAATAAATAAACACAGAGTAAATTCCCAAATTATTCCATCATTAAAAGATACGAGG
CGCGTGTAAGTTACAGGCAAGCGATCCGTCCTAAGAAACCATTATTATCATGACATT
AACCTATAAAAATAGGCGTATCACGAGGCCCTTTCGTCTCGCGCGTTTCGGTGATGA
CGGTGAAAACCTCTGACACATGCAGCTCCCGGAGACGGTCACAGCTTGTCTGTAAG
CGGATGCCGGGAGCAGACAAGCCCGTCAGGGCGCGTCAGCGGGTGTTGGCGGGTGT
CGGGGCTGGCTTAACTATGCGGCATCAGAGCAGATTGTACTGAGAGTGCACCATATC
GACTACGTCGTTAAGGCCGTTTCTGACAGAGTAAAATTCTTGAGGGAACTTTCACCA
TTATGGGAAATGGTTCAAGAAGGTATTGACTTAAACTCCATCAAATGGTCAGGTCAT
TGAGTGTTTTTTATTTGTTGTATTTTTTTTTTTTTAGAGAAAATCCTCCAATATATAAA
TTAGGAATCATAGTTTCATGATTTTCTGTTACACCTAACTTTTGTGTGGTGCCCTCC
TCCTTGTCAATATTAATGTTAAAGTGCAATTCTTTTTCCTTATCACGTTGAGCCATTA

GTATCAATTTGCTTACCTGTATTCCTTTACATCCTCCTTTTTCTCCTTCTTGATAAATG
TATGTAGATTGCGTATATAGTTTCGTCTACCCTATGAACATATTCCATTTTGTAATTT
CGTGTCGTTTCTATTATGAATTTCATTTATAAAGTTTATGTACAAATATCATAAAAAA
AGAGAATCTTTTTAAGCAAGGATTTTCTTAACTTCTTCGGCGACAGCATCACCGACT
TCGGTGGTACTGTTGGAACCACCTAAATCACCAGTTCTGATACCTGCATCCAAAACC
TTTTTAACTGCATCTTCAATGGCCTTACCTTCTTCAGGCAAGTTCAATGACAATTTCA
ACATCATTGCAGCAGACAAGATAGTGGCGATAGGGTTGACCTTATTCTTTGGCAAAT
CTGGAGCAGAACCGTGGCATGGTTCGTACAAACCAAATGCGGTGTTCTTGTCTGGCA
AAGAGGCCAAGGACGCAGATGGCAACAAACCCAAGGAACCTGGGATAACGGAGGC
TTCATCGGAGATGATATCACCAAACATGTTGCTGGTGATTATAATACCATTTAGGTG
GGTTGGGTTCTTAACTAGGATCATGGCGGCAGAATCAATCAATTGATGTTGAACCTT
CAATGTAGGGAATTCGTTCTTGATGGTTTCCTCCACAGTTTTTCTCCATAATCTTGAA
GAGGCCAAAACATTAGCTTTATCCAAGGACCAAATAGGCAATGGTGGCTCATGTTGT
AGGGCCATGAAAGCGGCCATTCTTGTGATTCTTTGCACTTCTGGAACGGTGTATTGT
TCACTATCCCAAGCGACACCATCACCATCGTCTTCCTTTCTCTTACCAAAGTAAATAC
CTCCCACTAATTCTCTGACAACAACGAAGTCAGTACCTTTAGCAAATTGTGGCTTGA
TTGGAGATAAGTCTAAAAGAGAGTCGGATGCAAAGTTACATGGTCTTAAGTTGGCG
TACAATTGAAGTTCTTTACGGATTTTTAGTAAACCTTGTTCAGGTCTAACACTACCGG
TACCCCATTTAGGACCACCCACAGCACCTAACAAAACGGCATCAGCCTTCTTGGAGG
CTTCCAGCGCCTCATCTGGAAGTGGAACACCTGTAGCATCGATAGCAGCACCACCA
ATTAAATGATTTTCGAAATCGAACTTGACATTGGAACGAACATCAGAAATAGCTTTA
AGAACCTTAATGGCTTCGGCTGTGATTTCTTGACCAACGTGGTCACCTGGCAAAACG
ACGATCTTCTTAGGGGCAGACATTAGAATGGTATATCCTTGAAATATATATATATAT
ATTGCTGAAATGTAAAAGGTAAGAAAAGTTAGAAAGTAAGACGATTGCTAACCACC
TATTGGAAAAAACAATAGGTCCTTAAATAATATTGTCAACTTCAAGTATTGTGATGC
AAGCATTTAGTCATGAACGCTTCTCTATTCTATATGAAAAGCCGGTTCCGGCGCTCT
CACCTTTCCTTTTTCTCCCAATTTTTCAGTTGAAAAAGGTATATGCGTCAGGCGACCT
CTGAAATTAACAAAAAATTTCCAGTCATCGAATTTGATTCTGTGCGATAGCGCCCCT
GTGTGTTCTCGTTATGTTGAGGAAAAAAATAATGGTTGCTAAGAGATTCGAACTCTT
GCATCTTACGATACCTGAGTATTCCCACAGTTAACTGCGGTCAAGATATTTCTTGAA
TCAGGCGCCTTAGACCGCTCGGCCAAACAACCAATTACTTGTTGAGAAATAGAGTAT
AATTATCCTATAAATATAACGTTTTTGAACACACATGAACAAGGAAGTACAGGACA
ATTGATTTTGAAGAGAATGTGGATTTTGATGTAATTGTTGGGATTCCATTTTTAATAA
GGCAATAATATTAGGTATGTAGATATACTAGAAGTTCTCCTCGACCGGTCGATATGC
GGTGTGAAATACCGCACAGATGCGTAAGGAGAAAATACCGCATCAGGAAATTGTAA
GCGTTAATATTTTGTTAAAATTCGCGTTAAATTTTTGTTAAATCAGCTCATTTTTTAA
CCAATAGGCCGAAATCGGCAAAATCCCTTATAAATCAAAAGAATAGACCGAGATAG
GGTTGAGTGTTGTTCCAGTTTGGAACAAGAGTCCACTATTAAAGAACGTGGACTCCA
ACGTCAAAGGGCGAAAAACCGTCTATCAGGGCGATGGCCCACTACGTGAACCATCA
CCCTAATCAAGTTTTTTGGGGTCGAGGTGCCGTAAAGCACTAAATCGGAACCCTAAA
GGGAGCCCCCGATTTAGAGCTTGACGGGGAAAGCCGGCGAACGTGGCGAGAAAGG
AAGGGAAGAAAGCGAAAGGAGCGGGCGCTAGGGCGCTGGCAAGTGTAGCGGTCAC
GCTGCGCGTAACCACCACACCCGCCGCGCTTAATGCGCCGCTACAGGGCGCGTCCAT
TCGCCATTCAGGCTGCGCAACTGTTGGGAAGGGCGATCGGTGCGGGCCTCTTCGCTA
TTACGCCAGCTGGCGAAAGGGGGATGTGCTGCAAGGCGATTAAGTTGGGTAACGCC
AGGGTTTTCCCAGTCACGACGTTGTAAAACGACGGCCAGTGAATTGTAATACGACTC

ACTATAGGGCGAATTGGAGCTCCACCGCGGTGGCGGCCGCTCTAGACAATACGAAG
GAACGATCCCGCGTCAGCAGATGTCACAATTTGATTATTTCATTGTCACATGCTTTTT
GACTCATCCTTCCGCGAAAGGAGAACTTTTTACAATGCCGCCGGCTCTGTTAGCGTA
CCCTTCTACAAGCAGATAGCAGAACAAACACATGATATATTCAAAAGGTGCAATGT
CAGAGAACATATATTGCGCCCCTGTCCTGTAGACATCAGTCATTTTTCGCGGGACTT
GAATGGCGCACCATTATCCTATACAGAAAACATTAAATTTACTGCAAAATTTTGGGC
AAACGCTTGGATAGACTATGAATCCGTCATGAAGACACTACTTGTAAAATTATATAT
ATCCTTTTTTTTCATGTGCTAGTAAGAATTAGCAACTACTTTCATTCATCTTCGACGC
AACTTCGAGGAAAGCACCTTTTTACTTGCTGTAGCTCTATTCTCTTCCCCAACCACCT
TTTCCTTTATTCCAAAATTTTTAAAACTTTTTCTGTTACATTATATAATCTTCTGTCTG
AAATGTTTGGATATAACGCCTCTTGATCCACTTTGTATATGCGTGCTATTTATTTTCA
GATTTATAAAGAGTATGAGCGTCATTTACATAAATAGCTGAAGTTATTCATGGAAAA
TACGAAGAGCACGTATGTGAGCCAACAGAACATTTGACGTAAGACTCTACAATGTG
CCAAGAACTGGACAAGTAGAGGACTGAGAACTTTATTTCAATTCATTGCTCCTTTTT
GGTGGCGCTACCTTTAGCGAAGGTCAATGATGAATGTGCACATGCTGTCGAAACCA
AAAAGCAAATTCTAACCAACTTCAAAATGACATAGTCATCTGATATTTCTACTCATT
ATAGATAGTATGGGAGCCTTGAAACGAAAAGTAAGTAAAAAGCTGGATATGAGCAG
TATGAGGTAGACCTTAGCTACATCATTTCCCCCAATAGCTGCTGCAAATATCTGGTT
AAATTTGTGATTCCATGAAGAGGATAACAGACTTGTTAAAAAGCATCCTGTCAAAAT
CTAATTTTTGAAGGGCAGTATTCAATTCATAATTTACTTTAGCTTAGATCCTTCCAAT
TTATACATGGTATTATAACCAGATCATAAACTACAATCTGTCGCTACCGCATGTACG
AAGAATACTTAAGTCACTTGCTCTCTCATCATTTGTACATTTTTCAGTAATACCGTTT
GATAGCGCCGTCTTTATTACCCGGATTATTCCGTTGACCCTGAATGAAAAATTTTTTC
AGAAATCCAGTGCTAAGCGTCAAATCAATGAAATACATCACTGTATTTTTAACTGAT
ATACTGTTGGTGTGGCCTTAACGACACCTTTATTTCTTAATTCATTTCGGCTTGTCTC
CTTATTAAGACTACAGAAATAGACAAAGGAAATACTTCAATAATGGATAACGAGGT
TGAAAAAAATATTGAGATCTGGAAGGTCAAGAAGTTGGTCCAATCTTTAGAAAAAG
CTAGAGGTAATGGTACTTCTATGATTTCCTTAGTTATTCCTCCTAAGGGTCAAATTCC
ACTGTACCAAAAAATGTTAACAGATGAATATGGTACTGCCTCGAATATTAAATCTAG
GGTT**G**ATCGTCTTTCCGTTTTATCTGCTATCACTTCCACCCAACAAAAGTTGAAGCTA
TATAATACTTTGCCCAAGAACGGTTTAGTTTTATATTGTGGTGATATCATCACTGAAG
ATGGTAAAGAAA**G**AAAGGTCACTTTTGACATCGAACCTTACAAAC**T**TATCAACACA
TCCTTATATTTGTGTGATAACAAATTTCATACAGAAGTTCTTTCGGAATTGCTTCAAG
CTGACGACAAGTTCGGTTTTATAGTCATGGACGGTCAAGGTACTTTGTTTGGTTCTGT
GTCCGGTAATACGAGAACTGTTTTACATAAATTTACTGTCGATCTGCCAAAAAAGCA
TGGTAGAGGTGGTCAATCTGCGCTTCGTTTTGCTCGTTTAAGAGAAGAAAAAAGACA
TAATTATGTGAGAAAGGTCGCCGAAGTTGCTGTTCAAAATTTTATTACTAATGACAA
AGTCAATGTTAAGGGTTTAATTTTAGCTGGTTCTGCTGACTTTAAGACCGATTTGGCT
AAATCTGAATTATTCGATCCAAGACTAGCATGTAAGGTTATTTCCATCGTGGATGTT
TCTTATGGTGGTGAAAACGGTTTCAACCAGGCTATCGAACTTTCTGCCGAAGCGTTG
GCCAATGTCAAGTATGTTCAAGAAAAGAAATTATTGGAGGCATATTTTGACGAAATT
TCCCAGGACACTGGTAAATTCTGTTATGGTATAGATGATACTTTAAAGGCATTGGAT
TTAGGTGCAGTCGAAAAATTAATTGTTTTCGAAAATTTGGAAACTATCAGATATACA
TTTAAAGATGCCGAGGATAATGAGGTTATAAAATTCGCTGAACCAGAAGCCAAGGA
CAAGTCGTTTGCTATTGACAAAGCTACCGGCCAAGAAATGGACGTTGTCTCCGAAGA
ACCTTTAATTGAATGGCTAGCAGCTAACTACAAAAACTTCGGTGCTACCTTGGAATT

CATCACAGACAAATCTTCAGAAGGTGCCCAATTTGTCACAGGTTTTGGTGGTATTGG
TGCCATGCTGCGTTACAAAGTTAATTTTGAACAACTAGTTGATGAATCTGAGGATGA
ATATTATGACGAAGATGAAGGATCCGACTATGATTTCATTTAAATAAATAAAAGGG
GGAGAAAAAATCGAATCAAAAAGAATTTAATCACTAGATGCCAGATTTAAATTAAA
TTCGCTTTTAATTTTTTGTACAATATAATATATACTTGGTAAACCTTTTGCTCTATATT
GAGCTAATTCCTTTGTTGAAAGTACATAGTGGGTTTAGAGGACCGTGTATATTACGT
AGAAAATACAGTGAAAGGAGAGTTTCTCTTCAAAAGCCTCGACGGTATCGATAAGC
TTATCGATACCGTCGACCTCGAGGGGGGCCCGGTACCAGCTTTTGTTCCCTTTAGTG
AGGGTTAATTTCGAGCTTGGCGTAATCATGGTCATAGCTGTTTCCTGTGTGAAATTGT
TATCCGCTCACAATTCCACACAACATACGAGCCGGAAGCATAAAGTGTAAAGCCTG
GGGTGCCTAATGAGTGAGCTAACTCACATTAATTGCGTTGCGCTCACTGCCCGCTTT
CCAGTCGGGAAACCTGTCGTGCCAGCTGCATTAATGAATCGGCCAACGCGCGGGGA
GAGGCGGTTTGCGTATTGGGCGCTCTTCCGCTTCCTCGCTCACTGACTCGCTGCGCTC
GGTCGTTCGGCTGCGGCGAGCGGTATCAGCTCACTCAAAGGCGGTAATACGGTTATC
CACAGAATCAGGGGATAACGCAGGAAAGAACATGTGAGCAAAAGGCCAGCAAAAG
GCCAGGAACCGTAAAAAGGCCGCGTTGCTGGCGTTTTTCCATAGGCTCCGCCCCCCT
GACGAGCATCACAAAAATCGACGCTCAAGTCAGAGGTGGCGAAACCCGACAGGACT
ATAAAGATACCAGGCGTTTCCCCCTGGAAGCTCCCTCGTGCGCTCTCCTGTTCCGAC
CCTGCCGCTTACCGGATACCTGTCCGCCTTTCTCCCTTCGGGAAGCGTGGCGCTTTCT
CATAGCTCACGCTGTAGGTATCTCAGTTCGGTGTAGGTCGTTCGCTCCAAGCTGGGC
TGTGTGCACGAACCCCCCGTTCAGCCCGACCGCTGCGCCTTATCCGGTAACTATCGT
CTTGAGTCCAACCCGGTAAGACACGACTTATCGCCACTGGCAGCAGCCACTGGTAA
CAGGATTAGCAGAGCGAGGTATGTAGGCGGTGCTACAGAGTTCTTGAAGTGGTGGC
CTAACTACGGCTACACTAGAAGAACAGTATTTGGTATCTGCGCTCTGCTGAAGCCAG
TTACCTTCGGAAAAAGAGTTGGTAGCTCTTGATCCGGCAAACAAACCACCGCTGGTA
GCGGTGGTTTTTTTGTTTGCAAGCAGCAGATTACGCGCAGAAAAAAAGGATCTCAAG
AAGATCCTTTGATCTTTTCTACGGGGTCTGACGCTCAGTGGAACGAAAACTCACGTT
AAGGGATTTTGGTCATGAGATTATCAAAAAGGATCTTCACCTAGATCCTTTTAAATT
AAAAATGAAGTTTTAAATCAATCTAAAGTATATATGAGTAAACTTGGTCTGACAGTT
ACCAATGCTTAATCAGTGAGGCACCTATCTCAGCGATCTGTCTATTTCGTTCATCCAT
AGTTGCCTGACTCCCCGTCGTGTAGATAACTACGATACGGGAGGGCTTACCATCTGG
CCCCAGTGCTGCAATGATACCGCGAGACCCACGCTCACCGGCTCCAGATTTATCAGC
AATAAACCAGCCAGCCGGAAGGGCCGAGCGCAGAAGTGGTCCTGCAACTTTATCCG
CCTCCATCCAGTCTATTAATTGTTGCCGGGAAGCTAGAGTAAGTAGTTCGCCAGTTA
ATAGTTTGCGCAACGTTGTTGCCATTGCTACAGGCATCGTGGTGTCACGCTCGTCGTT
TGGTATGGCTTCATTCAGCTCCGGTTCCCAACGATCAAGGCGAGTTACATGATCCCC
CATGTTGTGCAAAAAAGCGGTTAGCTCCTTCGGTCCTCCGATCGTTGTCAGAAGTAA
GTTGGCCGCAGTGTTATCACTCATGGTTATGGCAGCACTGCATAATTCTCTTACTGTC
ATGCCATCCGTAAGATGCTTTTCTGTGACTGGTGAGTACTCAACCAAGTCATTCTGA
GAATAGTGTATGCGGCGACCGAGTTGCTCTTGCCCGGCGTCAATACGGGATAATACC
GCGCCACATAGCAGAACTTTAAAAGTGCTCATCATTGGAAAACGTTCTTCGGGGCGA
AAACTCTCAAGGATCTTACCGCTGTTGAGATCCAGTTCGATGTAACCCA

**PRS413-SUP45-WT**

*SUP45* Promoter *SUP45-WT* HIS3 Promoter HIS3

TCGCGCGTTTCGGTGATGACGGTGAAAACCTCTGACACATGCAGCTCCCGGAGACG
GTCACAGCTTGTCTGTAAGCGGATGCCGGGAGCAGACAAGCCCGTCAGGGCGCGTC
AGCGCGTGTTGGCGGGTGTCGGGGCTGGCTTAACTATGCGGCATCAGAGCAGATTGT
ACTGAGAGTGCACCATAAATTCCCGTTTTAAGAGCTTGGTGAGCGCTAGGAGTCACT
GCCAGGTATCGTTTGAACACGGCATTAGTCAGGGAAGTCATAACACAGTCCTTTCCC
GCAATTTTCTTTTTCTATTACTCTTGGCCTCCTCTAGTACACTCTATATTTTTTTATGC
CTCGGTAATGATTTTCATTTTTTTTTTTCCCCTAGCGGATGACTCTTTTTTTTTCTTAG
CGATTGGCATTATCACATAATGAATTATACATTATATAAAGTAATGTGATTTCTTCG
AAGAATATACTAAAAAATGAGCAGGCAAGATAAACGAAGGCAAAGATGACAGAGC
AGAAAGCCCTAGTAAAGCGTATTACAAATGAAACCAAGATTCAGATTGCGATCTCTT
TAAAGGGTGGTCCCCTAGCGATAGAGCACTCGATCTTCCCAGAAAAAGAGGCAGAA
GCAGTAGCAGAACAGGCCACACAATCGCAAGTGATTAACGTCCACACAGGTATAGG
GTTTCTGGACCATATGATACATGCTCTGGCCAAGCATTCCGGCTGGTCGCTAATCGT
TGAGTGCATTGGTGACTTACACATAGACGACCATCACACCACTGAAGACTGCGGGA
TTGCTCTCGGTCAAGCTTTTAAAGAGGCCCTACTGGCGCGTGGAGTAAAAAGGTTTG
GATCAGGATTTGCGCCTTTGGATGAGGCACTTTCCAGAGCGGTGGTAGATCTTTCGA
ACAGGCCGTACGCAGTTGTCGAACTTGGTTTGCAAAGGGAGAAAGTAGGAGATCTC
TCTTGCGAGATGATCCCGCATTTTCTTGAAAGCTTTGCAGAGGCTAGCAGAATTACC
CTCCACGTTGATTGTCTGCGAGGCAAGAATGATCATCACCGTAGTGAGAGTGCGTTC
AAGGCTCTTGCGGTTGCCATAAGAGAAGCCACCTCGCCCAATGGTACCAACGATGTT
CCCTCCACCAAAGGTGTTCTTATGTAGTGACACCGATTATTTAAAGCTGCAGCATAC
GATATATATACATGTGTATATATGTATACCTATGAATGTCAGTAAGTATGTATACGA
ACAGTATGATACTGAAGATGACAAGGTAATGCATCATTCTATACGTGTCATTCTGAA
CGAGGCGCGCTTTCCTTTTTTCTTTTTGCTTTTTCTTTTTTTTTCTCTTGAACTCGACGG
ATCTATGCGGTGTGAAATACCGCACAGATGCGTAAGGAGAAAATACCGCATCAGGA
AATTGTAAACGTTAATATTTTGTTAAAATTCGCGTTAAATTTTTGTTAAATCAGCTCA
TTTTTTAACCAATAGGCCGAAATCGGCAAAATCCCTTATAAATCAAAAGAATAGACC
GAGATAGGGTTGAGTGTTGTTCCAGTTTGGAACAAGAGTCCACTATTAAAGAACGTG
GACTCCAACGTCAAAGGGCGAAAAACCGTCTATCAGGGCGATGGCCCACTACGTGA
ACCATCACCCTAATCAAGTTTTTTGGGGTCGAGGTGCCGTAAAGCACTAAATCGGAA
CCCTAAAGGGAGCCCCCGATTTAGAGCTTGACGGGGAAAGCCGGCGAACGTGGCGA
GAAAGGAAGGGAAGAAAGCGAAAGGAGCGGGCGCTAGGGCGCTGGCAAGTGTAGC
GGTCACGCTGCGCGTAACCACCACACCCGCCGCGCTTAATGCGCCGCTACAGGGCG
CGTCGCGCCATTCGCCATTCAGGCTGCGCAACTGTTGGGAAGGGCGATCGGTGCGG
GCCTCTTCGCTATTACGCCAGCTGGCGAAAGGGGGATGTGCTGCAAGGCGATTAAGT
TGGGTAACGCCAGGGTTTTCCCAGTCACGACGTTGTAAAACGACGGCCAGTGAGCG
CGCGTAATACGACTCACTATAGGGCGAATTGGGTACCGGGCCCCCCCTCGAGGCTA
GACAATACGAAGGAACGATCCCGCGTCAGCAGATGTCACAATTTGATTATTTCATTG
TCACATGCTTTTTGACTCATCCTTCCGCGAAAGGAGAACTTTTTACAATGCCGCCGG
CTCTGTTAGCGTACCCTTCTACAAGCAGATAGCAGAACAAACACATGATATATTCAA
AAGGTGCAATGTCAGAGAACATATATTGCGCCCCTGTCCTGTAGACATCAGTCATTT
TTCGCGGGACTTGAATGGCGCACCATTATCCTATACAGAAAACATTAAATTTACTGC
AAAATTTTGGGCAAACGCTTGGATAGACTATGAATCCGTCATGAAGACACTACTTGT
AAAATTATATATATCCTTTTTTTTCATGTGCTAGTAAGAATTAGCAACTACTTTCATT
CATCTTCGACGCAACTTCGAGGAAAGCACCTTTTTACTTGCTGTAGCTCTATTCTCTT
CCCCAACCACCTTTTCCTTTATTCCAAAATTTTTAAAACTTTTTCTGTTACATTATATA

ATCTTCTGTCTGAAATGTTTGGATATAACGCCTCTTGATCCACTTTGTATATGCGTGC
TATTTATTTTCAGATTTATAAAGAGTATGAGCGTCATTTACATAAATAGCTGAAGTT
ATTCATGGAAAATACGAAGAGCACGTATGTGAGCCAACAGAACATTTGACGTAAGA
CTCTACAATGTGCCAAGAACTGGACAAGTAGAGGACTGAGAACTTTATTTCAATTCA
TTGCTCCTTTTTGGTGGCGCTACCTTTAGCGAAGGTCAATGATGAATGTGCACATGCT
GTCGAAACCAAAAAGCAAATTCTAACCAACTTCAAAATGACATAGTCATCTGATATT
TCTACTCATTATAGATAGTATGGGAGCCTTGAAACGAAAAGTAAGTAAAAAGCTGG
ATATGAGCAGTATGAGGTAGACCTTAGCTACATCATTTCCCCCAATAGCTGCTGCAA
ATATCTGGTTAAATTTGTGATTCCATGAAGAGGATAACAGACTTGTTAAAAAGCATC
CTGTCAAAATCTAATTTTTGAAGGGCAGTATTCAATTCATAATTTACTTTAGCTTAGA
TCCTTCCAATTTATACATGGTATTATAACCAGATCATAAACTACAATCTGTCGCTACC
GCATGTACGAAGAATACTTAAGTCACTTGCTCTCTCATCATTTGTACATTTTTCAGTA
ATACCGTTTGATAGCGCCGTCTTTATTACCCGGATTATTCCGTTGACCCTGAATGAAA
AATTTTTTCAGAAATCCAGTGCTAAGCGTCAAATCAATGAAATACATCACTGTATTT
TTAACTGATATACTGTTGGTGTGGCCTTAACGACACCTTTATTTCTTAATTCATTTCG
GCTTGTCTCCTTATTAAGACTACAGAAATAGACAAAGGAAATACTTCAATAATGGAT
AACGAGGTTGAAAAAAATATTGAGATCTGGAAGGTCAAGAAGTTGGTCCAATCTTT
AGAAAAAGCTAGAGGTAATGGTACTTCTATGATTTCCTTAGTTATTCCTCCTAAGGG
TCAAATTCCACTGTACCAAAAAATGTTAACAGATGAATATGGTACTGCCTCGAATAT
TAAATCTAGGGTTAATCGTCTTTCCGTTTTATCTGCTATCACTTCCACCCAACAAAAG
TTGAAGCTATATAATACTTTGCCCAAGAACGGTTTAGTTTTATATTGTGGTGATATCA
TCACTGAAGATGGTAAAGAAAAAAAGGTCACTTTTGACATCGAACCTTACAAACCT
ATCAACACATCCTTATATTTGTGTGATAACAAATTTCATACAGAAGTTCTTTCGGAAT
TGCTTCAAGCTGACGACAAGTTCGGTTTTATAGTCATGGACGGTCAAGGTACTTTGT
TTGGTTCTGTGTCCGGTAATACGAGAACTGTTTTACATAAATTTACTGTCGATCTGCC
AAAAAAGCATGGTAGAGGTGGTCAATCTGCGCTTCGTTTTGCTCGTTTAAGAGAAGA
AAAAAGACATAATTATGTGAGAAAGGTCGCCGAAGTTGCTGTTCAAAATTTTATTAC
TAATGACAAAGTCAATGTTAAGGGTTTAATTTTAGCTGGTTCTGCTGACTTTAAGAC
CGATTTGGCTAAATCTGAATTATTCGATCCAAGACTAGCATGTAAGGTTATTTCCAT
CGTGGATGTTTCTTATGGTGGTGAAAACGGTTTCAACCAGGCTATCGAACTTTCTGC
CGAAGCGTTGGCCAATGTCAAGTATGTTCAAGAAAAGAAATTATTGGAGGCATATTT
TGACGAAATTTCCCAGGACACTGGTAAATTCTGTTATGGTATAGATGATACTTTAAA
GGCATTGGATTTAGGTGCAGTCGAAAAATTAATTGTTTTCGAAAATTTGGAAACTAT
CAGATATACATTTAAAGATGCCGAGGATAATGAGGTTATAAAATTCGCTGAACCAG
AAGCCAAGGACAAGTCGTTTGCTATTGACAAAGCTACCGGCCAAGAAATGGACGTT
GTCTCCGAAGAACCTTTAATTGAATGGCTAGCAGCTAACTACAAAAACTTCGGTGCT
ACCTTGGAATTCATCACAGACAAATCTTCAGAAGGTGCCCAATTTGTCACAGGTTTT
GGTGGTATTGGTGCCATGCTGCGTTACAAAGTTAATTTTGAACAACTAGTTGATGAA
TCTGAGGATGAATATTATGACGAAGATGAAGGATCCGACTATGATTTCATTTAAATA
AATAAAAGGGGGAGAAAAAAATCGAATCAAAAAGAATTTAATCACTAGATGCCAG
ATTTAAATTAAATTCGCTTTTAATTTTTTGTACAATATAATATATACTTGGTAAACCT
TTTGCTCTATATTGAGCTAATTCCTTTGTTGAAAGTACATAGTGGGTTTAGAGGACCG
TGTATATTACGTAGAAAATACAGTGAAAGGAGAGTTTCTCTTCAAAAGCCTCGACGG
TATCGATAAGCTTATCGATACCGCTAGAGCGGCCGCCACCGCGGTGGAGCTCCAGCT
TTTGTTCCCTTTAGTGAGGGTTAATTGCGCGCTTGGCGTAATCATGGTCATAGCTGTT
TCCTGTGTGAAATTGTTATCCGCTCACAATTCCACACAACATAGGAGCCGGAAGCAT

AAAGTGTAAAGCCTGGGGTGCCTAATGAGTGAGGTAACTCACATTAATTGCGTTGCG
CTCACTGCCCGCTTTCCAGTCGGGAAACCTGTCGTGCCAGCTGCATTAATGAATCGG
CCAACGCGCGGGGAGAGGCGGTTTGCGTATTGGGCGCTCTTCCGCTTCCTCGCTCAC
TGACTCGCTGCGCTCGGTCGTTCGGCTGCGGCGAGCGGTATCAGCTCACTCAAAGGC
GGTAATACGGTTATCCACAGAATCAGGGGATAACGCAGGAAAGAACATGTGAGCAA
AAGGCCAGCAAAAGGCCAGGAACCGTAAAAAGGCCGCGTTGCTGGCGTTTTTCCAT
AGGCTCCGCCCCCCTGACGAGCATCACAAAAATCGACGCTCAAGTCAGAGGTGGCG
AAACCCGACAGGACTATAAAGATACCAGGCGTTTCCCCCTGGAAGCTCCCTCGTGC
GCTCTCCTGTTCCGACCCTGCCGCTTACCGGATACCTGTCCGCCTTTCTCCCTTCGGG
AAGCGTGGCGCTTTCTCATAGCTCACGCTGTAGGTATCTCAGTTCGGTGTAGGTCGT
TCGCTCCAAGCTGGGCTGTGTGCACGAACCCCCCGTTCAGCCCGACCGCTGCGCCTT
ATCCGGTAACTATCGTCTTGAGTCCAACCCGGTAAGACACGACTTATCGCCACTGGC
AGCAGCCACTGGTAACAGGATTAGCAGAGCGAGGTATGTAGGCGGTGCTACAGAGT
TCTTGAAGTGGTGGCCTAACTACGGCTACACTAGAAGGACAGTATTTGGTATCTGCG
CTCTGCTGAAGCCAGTTACCTTCGGAAAAAGAGTTGGTAGCTCTTGATCCGGCAAAC
AAACCACCGCTGGTAGCGGTGGTTTTTTTGTTTGCAAGCAGCAGATTACGCGCAGAA
AAAAAGGATCTCAAGAAGATCCTTTGATCTTTTCTACGGGGTCTGACGCTCAGTGGA
ACGAAAACTCACGTTAAGGGATTTTGGTCATGAGATTATCAAAAAGGATCTTCACCT
AGATCCTTTTAAATTAAAAATGAAGTTTTAAATCAATCTAAAGTATATATGAGTAAA
CTTGGTCTGACAGTTACCAATGCTTAATCAGTGAGGCACCTATCTCAGCGATCTGTC
TATTTCGTTCATCCATAGTTGCCTGACTCCCCGTCGTGTAGATAACTACGATACGGG
AGGGCTTACCATCTGGCCCCAGTGCTGCAATGATACCGCGAGACCCACGCTCACCG
GCTCCAGATTTATCAGCAATAAACCAGCCAGCCGGAAGGGCCGAGCGCAGAAGTGG
TCCTGCAACTTTATCCGCCTCCATCCAGTCTATTAATTGTTGCCGGGAAGCTAGAGTA
AGTAGTTCGCCAGTTAATAGTTTGCGCAACGTTGTTGCCATTGCTACAGGCATCGTG
GTGTCACGCTCGTCGTTTGGTATGGCTTCATTCAGCTCCGGTTCCCAACGATCAAGG
CGAGTTACATGATCCCCCATGTTGTGCAAAAAAGCGGTTAGCTCCTTCGGTCCTCCG
ATCGTTGTCAGAAGTAAGTTGGCCGCAGTGTTATCACTCATGGTTATGGCAGCACTG
CATAATTCTCTTACTGTCATGCCATCCGTAAGATGCTTTTCTGTGACTGGTGAGTACT
CAACCAAGTCATTCTGAGAATAGTGTATGCGGCGACCGAGTTGCTCTTGCCCGGCGT
CAATACGGGATAATACCGCGCCACATAGCAGAACTTTAAAAGTGCTCATCATTGGA
AAACGTTCTTCGGGGCGAAAACTCTCAAGGATCTTACCGCTGTTGAGATCCAGTTCG
ATGTAACCCACTCGTGCACCCAACTGATCTTCAGCATCTTTTACTTTCACCAGCGTTT
CTGGGTGAGCAAAAACAGGAAGGCAAAATGCCGCAAAAAAGGGAATAAGGGCGAC
ACGGAAATGTTGAATACTCATACTCTTCCTTTTTCAATATTATTGAAGCATTTATCAG
GGTTATTGTCTCATGAGCGGATACATATTTGAATGTATTTAGAAAAATAAACAAATA
GGGGTTCCGCGCACATTTCCCCGAAAAGTGCCACCTGGGTCCTTTTCATCACGTGCT
ATAAAAATAATTATAATTTAAATTTTTTAATATAAATATATAAATTAAAAATAGAAA
GTAAAAAAAGAAATTAAAGAAAAAATAGTTTTTGTTTTCCGAAGATGTAAAAGACT
CTAGGGGGATCGCCAACAAATACTACCTTTTATCTTGCTCTTCCTGCTCTCAGGTATT
AATGCCGAATTGTTTCATCTTGTCTGTGTAGAAGACCACACACGAAAATCCTGTGAT
TTTACATTTTACTTATCGTTAATCGAATGTATATCTATTTAATCTGCTTTTCTTGTCTA
ATAAATATATATGTAAAGTACGCTTTTTGTTGAAATTTTTTAAACCTTTGTTTATTTTT
TTTTCTTCATTCCGTAACTCTTCTACCTTCTTTATTTACTTTCTAAAATCCAAATACAA
AACATAAAAATAAATAAACACAGAGTAAATTCCCAAATTATTCCATCATTAAAAGA

TACGAGGCGCGTGTAAGTTACAGGCAAGCGATCCGTCCTAAGAAACCATTATTATCA
TGACATTAACCTATAAAAATAGGCGTATCACGAGGCCCTTTCGTC

**PRS413-SUP45-N58A**
*SUP45* Promoter *SUP45-**N58A*** *HIS3* Promoter *HIS3*

TCGCGCGTTTCGGTGATGACGGTGAAAACCTCTGACACATGCAGCTCCCGGAGACG
GTCACAGCTTGTCTGTAAGCGGATGCCGGGAGCAGACAAGCCCGTCAGGGCGCGTC
AGCGCGTGTTGGCGGGTGTCGGGGCTGGCTTAACTATGCGGCATCAGAGCAGATTGT
ACTGAGAGTGCACCATAAATTCCCGTTTTAAGAGCTTGGTGAGCGCTAGGAGTCACT
GCCAGGTATCGTTTGAACACGGCATTAGTCAGGGAAGTCATAACACAGTCCTTTCCC
GCAATTTTCTTTTTCTATTACTCTTGGCCTCCTCTAGTACACTCTATATTTTTTTATGC
CTCGGTAATGATTTTCATTTTTTTTTTTCCCCTAGCGGATGACTCTTTTTTTTTCTTAG
CGATTGGCATTATCACATAATGAATTATACATTATATAAAGTAATGTGATTTCTTCG
AAGAATATACTAAAAAATGAGCAGGCAAGATAAACGAAGGCAAAGATGACAGAGC
AGAAAGCCCTAGTAAAGCGTATTACAAATGAAACCAAGATTCAGATTGCGATCTCTT
TAAAGGGTGGTCCCCTAGCGATAGAGCACTCGATCTTCCCAGAAAAAGAGGCAGAA
GCAGTAGCAGAACAGGCCACACAATCGCAAGTGATTAACGTCCACACAGGTATAGG
GTTTCTGGACCATATGATACATGCTCTGGCCAAGCATTCCGGCTGGTCGCTAATCGT
TGAGTGCATTGGTGACTTACACATAGACGACCATCACACCACTGAAGACTGCGGGA
TTGCTCTCGGTCAAGCTTTTAAAGAGGCCCTACTGGCGCGTGGAGTAAAAAGGTTTG
GATCAGGATTTGCGCCTTTGGATGAGGCACTTTCCAGAGCGGTGGTAGATCTTTCGA
ACAGGCCGTACGCAGTTGTCGAACTTGGTTTGCAAAGGGAGAAAGTAGGAGATCTC
TCTTGCGAGATGATCCCGCATTTTCTTGAAAGCTTTGCAGAGGCTAGCAGAATTACC
CTCCACGTTGATTGTCTGCGAGGCAAGAATGATCATCACCGTAGTGAGAGTGCGTTC
AAGGCTCTTGCGGTTGCCATAAGAGAAGCCACCTCGCCCAATGGTACCAACGATGTT
CCCTCCACCAAAGGTGTTCTTATGTAGTGACACCGATTATTTAAAGCTGCAGCATAC
GATATATATACATGTGTATATATGTATACCTATGAATGTCAGTAAGTATGTATACGA
ACAGTATGATACTGAAGATGACAAGGTAATGCATCATTCTATACGTGTCATTCTGAA
CGAGGCGCGCTTTCCTTTTTTCTTTTTGCTTTTTCTTTTTTTTTCTCTTGAACTCGACGG
ATCTATGCGGTGTGAAATACCGCACAGATGCGTAAGGAGAAAATACCGCATCAGGA
AATTGTAAACGTTAATATTTTGTTAAAATTCGCGTTAAATTTTTGTTAAATCAGCTCA
TTTTTTAACCAATAGGCCGAAATCGGCAAAATCCCTTATAAATCAAAAGAATAGACC
GAGATAGGGTTGAGTGTTGTTCCAGTTTGGAACAAGAGTCCACTATTAAAGAACGTG
GACTCCAACGTCAAAGGGCGAAAAACCGTCTATCAGGGCGATGGCCCACTACGTGA
ACCATCACCCTAATCAAGTTTTTTGGGGTCGAGGTGCCGTAAAGCACTAAATCGGAA
CCCTAAAGGGAGCCCCCGATTTAGAGCTTGACGGGGAAAGCCGGCGAACGTGGCGA
GAAAGGAAGGGAAGAAAGCGAAAGGAGCGGGCGCTAGGGCGCTGGCAAGTGTAGC
GGTCACGCTGCGCGTAACCACCACACCCGCCGCGCTTAATGCGCCGCTACAGGGCG
CGTCGCGCCATTCGCCATTCAGGCTGCGCAACTGTTGGGAAGGGCGATCGGTGCGG
GCCTCTTCGCTATTACGCCAGCTGGCGAAAGGGGGATGTGCTGCAAGGCGATTAAGT
TGGGTAACGCCAGGGTTTTCCCAGTCACGACGTTGTAAAACGACGGCCAGTGAGCG
CGCGTAATACGACTCACTATAGGGCGAATTGGGTACCGGGCCCCCCCTCGAGGCTA
GACAATACGAAGGAACGATCCCGCGTCAGCAGATGTCACAATTTGATTATTTCATTG
TCACATGCTTTTTGACTCATCCTTCCGCGAAAGGAGAACTTTTTACAATGCCGCCGG
CTCTGTTAGCGTACCCTTCTACAAGCAGATAGCAGAACAAACACATGATATATTCAA

AAGGTGCAATGTCAGAGAACATATATTGCGCCCCTGTCCTGTAGACATCAGTCATTT
TTCGCGGGACTTGAATGGCGCACCATTATCCTATACAGAAAACATTAAATTTACTGC
AAAATTTTGGGCAAACGCTTGGATAGACTATGAATCCGTCATGAAGACACTACTTGT
AAAATTATATATATCCTTTTTTTTCATGTGCTAGTAAGAATTAGCAACTACTTTCATT
CATCTTCGACGCAACTTCGAGGAAAGCACCTTTTTACTTGCTGTAGCTCTATTCTCTT
CCCCAACCACCTTTTCCTTTATTCCAAAATTTTTAAAACTTTTTCTGTTACATTATATA
ATCTTCTGTCTGAAATGTTTGGATATAACGCCTCTTGATCCACTTTGTATATGCGTGC
TATTTATTTTCAGATTTATAAAGAGTATGAGCGTCATTTACATAAATAGCTGAAGTT
ATTCATGGAAAATACGAAGAGCACGTATGTGAGCCAACAGAACATTTGACGTAAGA
CTCTACAATGTGCCAAGAACTGGACAAGTAGAGGACTGAGAACTTTATTTCAATTCA
TTGCTCCTTTTTGGTGGCGCTACCTTTAGCGAAGGTCAATGATGAATGTGCACATGCT
GTCGAAACCAAAAAGCAAATTCTAACCAACTTCAAAATGACATAGTCATCTGATATT
TCTACTCATTATAGATAGTATGGGAGCCTTGAAACGAAAAGTAAGTAAAAAGCTGG
ATATGAGCAGTATGAGGTAGACCTTAGCTACATCATTTCCCCCAATAGCTGCTGCAA
ATATCTGGTTAAATTTGTGATTCCATGAAGAGGATAACAGACTTGTTAAAAAGCATC
CTGTCAAAATCTAATTTTTGAAGGGCAGTATTCAATTCATAATTTACTTTAGCTTAGA
TCCTTCCAATTTATACATGGTATTATAACCAGATCATAAACTACAATCTGTCGCTACC
GCATGTACGAAGAATACTTAAGTCACTTGCTCTCTCATCATTTGTACATTTTTCAGTA
ATACCGTTTGATAGCGCCGTCTTTATTACCCGGATTATTCCGTTGACCCTGAATGAAA
AATTTTTTCAGAAATCCAGTGCTAAGCGTCAAATCAATGAAATACATCACTGTATTT
TTAACTGATATACTGTTGGTGTGGCCTTAACGACACCTTTATTTCTTAATTCATTTCG
GCTTGTCTCCTTATTAAGACTACAGAAATAGACAAAGGAAATACTTCAATAATGGAT
AACGAGGTTGAAAAAAATATTGAGATCTGGAAGGTCAAGAAGTTGGTCCAATCTTT
AGAAAAAGCTAGAGGTAATGGTACTTCTATGATTTCCTTAGTTATTCCTCCTAAGGG
TCAAATTCCACTGTACCAAAAAATGTTAACAGATGAATATGGTACTGCCTCG**GC**TAT
TAAATCTAGGGTTAATCGTCTTTCCGTTTTATCTGCTATCACTTCCACCCAACAAAAG
TTGAAGCTATATAATACTTTGCCCAAGAACGGTTTAGTTTTATATTGTGGTGATATCA
TCACTGAAGATGGTAAAGAAAAAAAGGTCACTTTTGACATCGAACCTTACAAACCT
ATCAACACATCCTTATATTTGTGTGATAACAAATTTCATACAGAAGTTCTTTCGGAAT
TGCTTCAAGCTGACGACAAGTTCGGTTTTATAGTCATGGACGGTCAAGGTACTTTGT
TTGGTTCTGTGTCCGGTAATACGAGAACTGTTTTACATAAATTTACTGTCGATCTGCC
AAAAAAGCATGGTAGAGGTGGTCAATCTGCGCTTCGTTTTGCTCGTTTAAGAGAAGA
AAAAAGACATAATTATGTGAGAAAGGTCGCCGAAGTTGCTGTTCAAAATTTTATTAC
TAATGACAAAGTCAATGTTAAGGGTTTAATTTTAGCTGGTTCTGCTGACTTTAAGAC
CGATTTGGCTAAATCTGAATTATTCGATCCAAGACTAGCATGTAAGGTTATTTCCAT
CGTGGATGTTTCTTATGGTGGTGAAAACGGTTTCAACCAGGCTATCGAACTTTCTGC
CGAAGCGTTGGCCAATGTCAAGTATGTTCAAGAAAAGAAATTATTGGAGGCATATTT
TGACGAAATTTCCCAGGACACTGGTAAATTCTGTTATGGTATAGATGATACTTTAAA
GGCATTGGATTTAGGTGCAGTCGAAAAATTAATTGTTTTCGAAAATTTGGAAACTAT
CAGATATACATTTAAAGATGCCGAGGATAATGAGGTTATAAAATTCGCTGAACCAG
AAGCCAAGGACAAGTCGTTTGCTATTGACAAAGCTACCGGCCAAGAAATGGACGTT
GTCTCCGAAGAACCTTTAATTGAATGGCTAGCAGCTAACTACAAAAACTTCGGTGCT
ACCTTGGAATTCATCACAGACAAATCTTCAGAAGGTGCCCAATTTGTCACAGGTTTT
GGTGGTATTGGTGCCATGCTGCGTTACAAAGTTAATTTTGAACAACTAGTTGATGAA
TCTGAGGATGAATATTATGACGAAGATGAAGGATCCGACTATGATTTCATTTAAATA
AATAAAAGGGGGAGAAAAAAATCGAATCAAAAAGAATTTAATCACTAGATGCCAG

ATTTAAATTAAATTCGCTTTTAATTTTTTGTACAATATAATATATACTTGGTAAACCT
TTTGCTCTATATTGAGCTAATTCCTTTGTTGAAAGTACATAGTGGGTTTAGAGGACCG
TGTATATTACGTAGAAAATACAGTGAAAGGAGAGTTTCTCTTCAAAAGCCTCGACGG
TATCGATAAGCTTATCGATACCGCTAGAGCGGCCGCCACCGCGGTGGAGCTCCAGCT
TTTGTTCCCTTTAGTGAGGGTTAATTGCGCGCTTGGCGTAATCATGGTCATAGCTGTT
TCCTGTGTGAAATTGTTATCCGCTCACAATTCCACACAACATAGGAGCCGGAAGCAT
AAAGTGTAAAGCCTGGGGTGCCTAATGAGTGAGGTAACTCACATTAATTGCGTTGCG
CTCACTGCCCGCTTTCCAGTCGGGAAACCTGTCGTGCCAGCTGCATTAATGAATCGG
CCAACGCGCGGGGAGAGGCGGTTTGCGTATTGGGCGCTCTTCCGCTTCCTCGCTCAC
TGACTCGCTGCGCTCGGTCGTTCGGCTGCGGCGAGCGGTATCAGCTCACTCAAAGGC
GGTAATACGGTTATCCACAGAATCAGGGGATAACGCAGGAAAGAACATGTGAGCAA
AAGGCCAGCAAAAGGCCAGGAACCGTAAAAAGGCCGCGTTGCTGGCGTTTTTCCAT
AGGCTCCGCCCCCCTGACGAGCATCACAAAAATCGACGCTCAAGTCAGAGGTGGCG
AAACCCGACAGGACTATAAAGATACCAGGCGTTTCCCCCTGGAAGCTCCCTCGTGC
GCTCTCCTGTTCCGACCCTGCCGCTTACCGGATACCTGTCCGCCTTTCTCCCTTCGGG
AAGCGTGGCGCTTTCTCATAGCTCACGCTGTAGGTATCTCAGTTCGGTGTAGGTCGT
TCGCTCCAAGCTGGGCTGTGTGCACGAACCCCCCGTTCAGCCCGACCGCTGCGCCTT
ATCCGGTAACTATCGTCTTGAGTCCAACCCGGTAAGACACGACTTATCGCCACTGGC
AGCAGCCACTGGTAACAGGATTAGCAGAGCGAGGTATGTAGGCGGTGCTACAGAGT
TCTTGAAGTGGTGGCCTAACTACGGCTACACTAGAAGGACAGTATTTGGTATCTGCG
CTCTGCTGAAGCCAGTTACCTTCGGAAAAAGAGTTGGTAGCTCTTGATCCGGCAAAC
AAACCACCGCTGGTAGCGGTGGTTTTTTTGTTTGCAAGCAGCAGATTACGCGCAGAA
AAAAAGGATCTCAAGAAGATCCTTTGATCTTTTCTACGGGGTCTGACGCTCAGTGGA
ACGAAAACTCACGTTAAGGGATTTTGGTCATGAGATTATCAAAAAGGATCTTCACCT
AGATCCTTTTAAATTAAAAATGAAGTTTTAAATCAATCTAAAGTATATATGAGTAAA
CTTGGTCTGACAGTTACCAATGCTTAATCAGTGAGGCACCTATCTCAGCGATCTGTC
TATTTCGTTCATCCATAGTTGCCTGACTCCCCGTCGTGTAGATAACTACGATACGGG
AGGGCTTACCATCTGGCCCCAGTGCTGCAATGATACCGCGAGACCCACGCTCACCG
GCTCCAGATTTATCAGCAATAAACCAGCCAGCCGGAAGGGCCGAGCGCAGAAGTGG
TCCTGCAACTTTATCCGCCTCCATCCAGTCTATTAATTGTTGCCGGGAAGCTAGAGTA
AGTAGTTCGCCAGTTAATAGTTTGCGCAACGTTGTTGCCATTGCTACAGGCATCGTG
GTGTCACGCTCGTCGTTTGGTATGGCTTCATTCAGCTCCGGTTCCCAACGATCAAGG
CGAGTTACATGATCCCCCATGTTGTGCAAAAAAGCGGTTAGCTCCTTCGGTCCTCCG
ATCGTTGTCAGAAGTAAGTTGGCCGCAGTGTTATCACTCATGGTTATGGCAGCACTG
CATAATTCTCTTACTGTCATGCCATCCGTAAGATGCTTTTCTGTGACTGGTGAGTACT
CAACCAAGTCATTCTGAGAATAGTGTATGCGGCGACCGAGTTGCTCTTGCCCGGCGT
CAATACGGGATAATACCGCGCCACATAGCAGAACTTTAAAAGTGCTCATCATTGGA
AAACGTTCTTCGGGGCGAAAACTCTCAAGGATCTTACCGCTGTTGAGATCCAGTTCG
ATGTAACCCACTCGTGCACCCAACTGATCTTCAGCATCTTTTACTTTCACCAGCGTTT
CTGGGTGAGCAAAAACAGGAAGGCAAAATGCCGCAAAAAAGGGAATAAGGGCGAC
ACGGAAATGTTGAATACTCATACTCTTCCTTTTTCAATATTATTGAAGCATTTATCAG
GGTTATTGTCTCATGAGCGGATACATATTTGAATGTATTTAGAAAAATAAACAAATA
GGGGTTCCGCGCACATTTCCCCGAAAAGTGCCACCTGGGTCCTTTTCATCACGTGCT
ATAAAAATAATTATAATTTAAATTTTTTAATATAAATATATAAATTAAAAATAGAAA
GTAAAAAAAGAAATTAAAGAAAAAATAGTTTTTGTTTTCCGAAGATGTAAAAGACT
CTAGGGGGATCGCCAACAAATACTACCTTTTATCTTGCTCTTCCTGCTCTCAGGTATT

AATGCCGAATTGTTTCATCTTGTCTGTGTAGAAGACCACACACGAAAATCCTGTGAT
TTTACATTTTACTTATCGTTAATCGAATGTATATCTATTTAATCTGCTTTTCTTGTCTA
ATAAATATATATGTAAAGTACGCTTTTTGTTGAAATTTTTTAAACCTTTGTTTATTTTT
TTTTCTTCATTCCGTAACTCTTCTACCTTCTTTATTTACTTTCTAAAATCCAAATACAA
AACATAAAAATAAATAAACACAGAGTAAATTCCCAAATTATTCCATCATTAAAAGA
TACGAGGCGCGTGTAAGTTACAGGCAAGCGATCCGTCCTAAGAAACCATTATTATCA
TGACATTAACCTATAAAAATAGGCGTATCACGAGGCCCTTTCGTC

**PRS413-SUP45-BR3T.4**

*SUP45* Promoter *SUP45-**BR3T.4*** *HIS3* Promoter *HIS3*

TCGCGCGTTTCGGTGATGACGGTGAAAACCTCTGACACATGCAGCTCCCGGAGACG
GTCACAGCTTGTCTGTAAGCGGATGCCGGGAGCAGACAAGCCCGTCAGGGCGCGTC
AGCGCGTGTTGGCGGGTGTCGGGGCTGGCTTAACTATGCGGCATCAGAGCAGATTGT
ACTGAGAGTGCACCATAAATTCCCGTTTTAAGAGCTTGGTGAGCGCTAGGAGTCACT
GCCAGGTATCGTTTGAACACGGCATTAGTCAGGGAAGTCATAACACAGTCCTTTCCC
GCAATTTTCTTTTTCTATTACTCTTGGCCTCCTCTAGTACACTCTATATTTTTTTATGC
CTCGGTAATGATTTCATTTTTTTTTTTCCCCTAGCGGATGACTCTTTTTTTTTCTTAG
CGATTGGCATTATCACATAATGAATTATACATTATATAAAGTAATGTGATTTCTTCG
AAGAATATACTAAAAATGAGCAGGCAAGATAAACGAAGGCAAAGATGACAGAGC
AGAAAGCCCTAGTAAAGCGTATTACAAATGAAACCAAGATTCAGATTGCGATCTCTT
TAAAGGGTGGTCCCCTAGCGATAGAGCACTCGATCTTCCCAGAAAAAGAGGCAGAA
GCAGTAGCAGAACAGGCCACACAATCGCAAGTGATTAACGTCCACACAGGTATAGG
GTTTCTGGACCATATGATACATGCTCTGGCCAAGCATTCCGGCTGGTCGCTAATCGT
TGAGTGCATTGGTGACTTACACATAGACGACCATCACACCACTGAAGACTGCGGGA
TTGCTCTCGGTCAAGCTTTTAAAGAGGCCCTACTGGCGCGTGGAGTAAAAAGGTTTG
GATCAGGATTTGCGCCTTTGGATGAGGCACTTTCCAGAGCGGTGGTAGATCTTTCGA
ACAGGCCGTACGCAGTTGTCGAACTTGGTTTGCAAAGGGAGAAAGTAGGAGATCTC
TCTTGCGAGATGATCCCGCATTTCTTGAAAGCTTTGCAGAGGCTAGCAGAATTACC
CTCCACGTTGATTGTCTGCGAGGCAAGAATGATCATCACCGTAGTGAGAGTGCGTTC
AAGGCTCTTGCGGTTGCCATAAGAGAAGCCACCTCGCCCAATGGTACCAACGATGTT
CCCTCCACCAAAGGTGTTCTTATGTAGTGACACCGATTATTTAAAGCTGCAGCATAC
GATATATATACATGTGTATATATGTATACCTATGAATGTCAGTAAGTATGTATACGA
ACAGTATGATACTGAAGATGACAAGGTAATGCATCATTCTATACGTGTCATTCTGAA
CGAGGCGCGCTTTCCTTTTTTCTTTTTGCTTTTTCTTTTTTTTTCTCTTGAACTCGACGG
ATCTATGCGGTGTGAAATACCGCACAGATGCGTAAGGAGAAAATACCGCATCAGGA
AATTGTAAACGTTAATATTTTGTTAAAATTCGCGTTAAATTTTTGTTAAATCAGCTCA
TTTTTTAACCAATAGGCCGAAATCGGCAAAATCCCTTATAAATCAAAAGAATAGACC
GAGATAGGGTTGAGTGTTGTTCCAGTTTGGAACAAGAGTCCACTATTAAAGAACGTG
GACTCCAACGTCAAAGGGCGAAAAACCGTCTATCAGGGCGATGGCCCACTACGTGA
ACCATCACCCTAATCAAGTTTTTTGGGGTCGAGGTGCCGTAAAGCACTAAATCGGAA
CCCTAAAGGGAGCCCCCGATTTAGAGCTTGACGGGGAAAGCCGGCGAACGTGGCGA
GAAAGGAAGGGAAGAAAGCGAAAGGAGCGGGCGCTAGGGCGCTGGCAAGTGTAGC
GGTCACGCTGCGCGTAACCACCACACCCGCCGCGCTTAATGCGCCGCTACAGGGCG
CGTCGCGCCATTCGCCATTCAGGCTGCGCAACTGTTGGGAAGGGCGATCGGTGCGG
GCCTCTTCGCTATTACGCCAGCTGGCGAAAGGGGGATGTGCTGCAAGGCGATTAAGT

TGGGTAACGCCAGGGTTTTCCCAGTCACGACGTTGTAAAACGACGGCCAGTGAGCG
CGCGTAATACGACTCACTATAGGGCGAATTGGGTACCGGGCCCCCCCTCGAGGCTA
GACAATACGAAGGAACGATCCCGCGTCAGCAGATGTCACAATTTGATTATTTCATTG
TCACATGCTTTTTGACTCATCCTTCCGCGAAAGGAGAACTTTTTACAATGCCGCCGG
CTCTGTTAGCGTACCCTTCTACAAGCAGATAGCAGAACAAACACATGATATATTCAA
AAGGTGCAATGTCAGAGAACATATATTGCGCCCCTGTCCTGTAGACATCAGTCATTT
TTCGCGGGACTTGAATGGCGCACCATTATCCTATACAGAAAACATTAAATTTACTGC
AAAATTTTGGGCAAACGCTTGGATAGACTATGAATCCGTCATGAAGACACTACTTGT
AAAATTATATATATCCTTTTTTTTCATGTGCTAGTAAGAATTAGCAACTACTTTCATT
CATCTTCGACGCAACTTCGAGGAAAGCACCTTTTTACTTGCTGTAGCTCTATTCTCTT
CCCCAACCACCTTTTCCTTTATTCCAAAATTTTTAAAACTTTTTCTGTTACATTATATA
ATCTTCTGTCTGAAATGTTTGGATATAACGCCTCTTGATCCACTTTGTATATGCGTGC
TATTTATTTTCAGATTTATAAAGAGTATGAGCGTCATTTACATAAATAGCTGAAGTT
ATTCATGGAAAATACGAAGAGCACGTATGTGAGCCAACAGAACATTTGACGTAAGA
CTCTACAATGTGCCAAGAACTGGACAAGTAGAGGACTGAGAACTTTATTTCAATTCA
TTGCTCCTTTTTGGTGGCGCTACCTTTAGCGAAGGTCAATGATGAATGTGCACATGCT
GTCGAAACCAAAAAGCAAATTCTAACCAACTTCAAAATGACATAGTCATCTGATATT
TCTACTCATTATAGATAGTATGGGAGCCTTGAAACGAAAAGTAAGTAAAAAGCTGG
ATATGAGCAGTATGAGGTAGACCTTAGCTACATCATTTCCCCCAATAGCTGCTGCAA
ATATCTGGTTAAATTTGTGATTCCATGAAGAGGATAACAGACTTGTTAAAAAGCATC
CTGTCAAAATCTAATTTTTGAAGGGCAGTATTCAATTCATAATTTACTTTAGCTTAGA
TCCTTCCAATTTATACATGGTATTATAACCAGATCATAAACTACAATCTGTCGCTACC
GCATGTACGAAGAATACTTAAGTCACTTGCTCTCTCATCATTTGTACATTTTTCAGTA
ATACCGTTTGATAGCGCCGTCTTTATTACCCGGATTATTCCGTTGACCCTGAATGAAA
AATTTTTTCAGAAATCCAGTGCTAAGCGTCAAATCAATGAAATACATCACTGTATTT
TTAACTGATATACTGTTGGTGTGGCCTTAACGACACCTTTATTTCTTAATTCATTTCG
GCTTGTCTCCTTATTAAGACTACAGAAATAGACAAAGGAAATACTTCAATAATGGAT
AACGAGGTTGAAAAAAATATTGAGATCTGGAAGGTCAAGAAGTTGGTCCAATCTTT
AGAAAAAGCTAGAGGTAATGGTACTTCTATGATTTCCTTAGTTATTCCTCCTAAGGG
TCAAATTCCACTGTACCAAAAAATGTTAACAGATGAATATGGTACTGCCTCGAATAT
TAAATCTAGGGTT**G**ATCGTCTTTCCGTTTTATCTGCTATCACTTCCACCCAACAAAAG
TTGAAGCTATATAATACTTTGCCCAAGAACGGTTTAGTTTTATATTGTGGTGATATCA
TCACTGAAGATGGTAAAGAAA**G**AAAGGTCACTTTTGACATCGAACCTTACAAAC**T**T
ATCAACACATCCTTATATTTGTGTGATAACAAATTTCATACAGAAGTTCTTTCGGAAT
TGCTTCAAGCTGACGACAAGTTCGGTTTTATAGTCATGGACGGTCAAGGTACTTTGT
TTGGTTCTGTGTCCGGTAATACGAGAACTGTTTTACATAAATTTACTGTCGATCTGCC
AAAAAAGCATGGTAGAGGTGGTCAATCTGCGCTTCGTTTTGCTCGTTTAAGAGAAGA
AAAAAGACATAATTATGTGAGAAAGGTCGCCGAAGTTGCTGTTCAAAATTTTATTAC
TAATGACAAAGTCAATGTTAAGGGTTTAATTTTAGCTGGTTCTGCTGACTTTAAGAC
CGATTTGGCTAAATCTGAATTATTCGATCCAAGACTAGCATGTAAGGTTATTTCCAT
CGTGGATGTTTCTTATGGTGGTGAAAACGGTTTCAACCAGGCTATCGAACTTTCTGC
CGAAGCGTTGGCCAATGTCAAGTATGTTCAAGAAAAGAAATTATTGGAGGCATATTT
TGACGAAATTTCCCAGGACACTGGTAAATTCTGTTATGGTATAGATGATACTTTAAA
GGCATTGGATTTAGGTGCAGTCGAAAAATTAATTGTTTTCGAAAATTTGGAAACTAT
CAGATATACATTTAAAGATGCCGAGGATAATGAGGTTATAAAATTCGCTGAACCAG
AAGCCAAGGACAAGTCGTTTGCTATTGACAAAGCTACCGGCCAAGAAATGGACGTT

GTCTCCGAAGAACCTTTAATTGAATGGCTAGCAGCTAACTACAAAAACTTCGGTGCT
ACCTTGGAATTCATCACAGACAAATCTTCAGAAGGTGCCCAATTTGTCACAGGTTTT
GGTGGTATTGGTGCCATGCTGCGTTACAAAGTTAATTTTGAACAACTAGTTGATGAA
TCTGAGGATGAATATTATGACGAAGATGAAGGATCCGACTATGATTTCATTTAAATA
AATAAAAGGGGGAGAAAAAAATCGAATCAAAAAGAATTTAATCACTAGATGCCAG
ATTTAAATTAAATTCGCTTTTAATTTTTTGTACAATATAATATATACTTGGTAAACCT
TTTGCTCTATATTGAGCTAATTCCTTTGTTGAAAGTACATAGTGGGTTTAGAGGACCG
TGTATATTACGTAGAAAATACAGTGAAAGGAGAGTTTCTCTTCAAAAGCCTCGACGG
TATCGATAAGCTTATCGATACCGCTAGAGCGGCCGCCACCGCGGTGGAGCTCCAGCT
TTTGTTCCCTTTAGTGAGGGTTAATTGCGCGCTTGGCGTAATCATGGTCATAGCTGTT
TCCTGTGTGAAATTGTTATCCGCTCACAATTCCACACAACATAGGAGCCGGAAGCAT
AAAGTGTAAAGCCTGGGGTGCCTAATGAGTGAGGTAACTCACATTAATTGCGTTGCG
CTCACTGCCCGCTTTCCAGTCGGGAAACCTGTCGTGCCAGCTGCATTAATGAATCGG
CCAACGCGCGGGGAGAGGCGGTTTGCGTATTGGGCGCTCTTCCGCTTCCTCGCTCAC
TGACTCGCTGCGCTCGGTCGTTCGGCTGCGGCGAGCGGTATCAGCTCACTCAAAGGC
GGTAATACGGTTATCCACAGAATCAGGGGATAACGCAGGAAAGAACATGTGAGCAA
AAGGCCAGCAAAAGGCCAGGAACCGTAAAAAGGCCGCGTTGCTGGCGTTTTTCCAT
AGGCTCCGCCCCCCTGACGAGCATCACAAAAATCGACGCTCAAGTCAGAGGTGGCG
AAACCCGACAGGACTATAAAGATACCAGGCGTTTCCCCCTGGAAGCTCCCTCGTGC
GCTCTCCTGTTCCGACCCTGCCGCTTACCGGATACCTGTCCGCCTTTCTCCCTTCGGG
AAGCGTGGCGCTTTCTCATAGCTCACGCTGTAGGTATCTCAGTTCGGTGTAGGTCGT
TCGCTCCAAGCTGGGCTGTGTGCACGAACCCCCCGTTCAGCCCGACCGCTGCGCCTT
ATCCGGTAACTATCGTCTTGAGTCCAACCCGGTAAGACACGACTTATCGCCACTGGC
AGCAGCCACTGGTAACAGGATTAGCAGAGCGAGGTATGTAGGCGGTGCTACAGAGT
TCTTGAAGTGGTGGCCTAACTACGGCTACACTAGAAGGACAGTATTTGGTATCTGCG
CTCTGCTGAAGCCAGTTACCTTCGGAAAAAGAGTTGGTAGCTCTTGATCCGGCAAAC
AAACCACCGCTGGTAGCGGTGGTTTTTTTGTTTGCAAGCAGCAGATTACGCGCAGAA
AAAAAGGATCTCAAGAAGATCCTTTGATCTTTTCTACGGGGTCTGACGCTCAGTGGA
ACGAAAACTCACGTTAAGGGATTTTGGTCATGAGATTATCAAAAAGGATCTTCACCT
AGATCCTTTTAAATTAAAAATGAAGTTTTAAATCAATCTAAAGTATATATGAGTAAA
CTTGGTCTGACAGTTACCAATGCTTAATCAGTGAGGCACCTATCTCAGCGATCTGTC
TATTTCGTTCATCCATAGTTGCCTGACTCCCCGTCGTGTAGATAACTACGATACGGG
AGGGCTTACCATCTGGCCCCAGTGCTGCAATGATACCGCGAGACCCACGCTCACCG
GCTCCAGATTTATCAGCAATAAACCAGCCAGCCGGAAGGGCCGAGCGCAGAAGTGG
TCCTGCAACTTTATCCGCCTCCATCCAGTCTATTAATTGTTGCCGGGAAGCTAGAGTA
AGTAGTTCGCCAGTTAATAGTTTGCGCAACGTTGTTGCCATTGCTACAGGCATCGTG
GTGTCACGCTCGTCGTTTGGTATGGCTTCATTCAGCTCCGGTTCCCAACGATCAAGG
CGAGTTACATGATCCCCCATGTTGTGCAAAAAAGCGGTTAGCTCCTTCGGTCCTCCG
ATCGTTGTCAGAAGTAAGTTGGCCGCAGTGTTATCACTCATGGTTATGGCAGCACTG
CATAATTCTCTTACTGTCATGCCATCCGTAAGATGCTTTTCTGTGACTGGTGAGTACT
CAACCAAGTCATTCTGAGAATAGTGTATGCGGCGACCGAGTTGCTCTTGCCCGGCGT
CAATACGGGATAATACCGCGCCACATAGCAGAACTTTAAAAGTGCTCATCATTGGA
AAACGTTCTTCGGGGCGAAAACTCTCAAGGATCTTACCGCTGTTGAGATCCAGTTCG
ATGTAACCCACTCGTGCACCCAACTGATCTTCAGCATCTTTTACTTTCACCAGCGTTT
CTGGGTGAGCAAAAACAGGAAGGCAAAATGCCGCAAAAAAGGGAATAAGGGCGAC
ACGGAAATGTTGAATACTCATACTCTTCCTTTTTCAATATTATTGAAGCATTTATCAG

GGTTATTGTCTCATGAGCGGATACATATTTGAATGTATTTAGAAAAATAAACAAATA
GGGGTTCCGCGCACATTTCCCCGAAAAGTGCCACCTGGGTCCTTTTCATCACGTGCT
ATAAAAATAATTATAATTTAAATTTTTTAATATAAATATATAAATTAAAAATAGAAA
GTAAAAAAAGAAATTAAAGAAAAAATAGTTTTTGTTTTCCGAAGATGTAAAAGACT
CTAGGGGGATCGCCAACAAATACTACCTTTTATCTTGCTCTTCCTGCTCTCAGGTATT
AATGCCGAATTGTTTCATCTTGTCTGTGTAGAAGACCACACACGAAAATCCTGTGAT
TTTACATTTTACTTATCGTTAATCGAATGTATATCTATTTAATCTGCTTTTCTTGTCTA
ATAAATATATATGTAAAGTACGCTTTTTGTTGAAATTTTTTAAACCTTTGTTTATTTTT
TTTTCTTCATTCCGTAACTCTTCTACCTTCTTTATTTACTTTCTAAAATCCAAATACAA
AACATAAAAATAAATAAACACAGAGTAAATTCCCAAATTATTCCATCATTAAAAGA
TACGAGGCGCGTGTAAGTTACAGGCAAGCGATCCGTCCTAAGAAACCATTATTATCA
TGACATTAACCTATAAAAATAGGCGTATCACGAGGCCCTTTCGTC

2 Gould, J. *Morpheus*, <https://software.broadinstitute.org/morpheus> (

3 Geneious version 2025.0.
